# *C. elegans* RIG-I-like receptor DRH-1 signals via CARDs to activate anti-viral immunity in intestinal cells

**DOI:** 10.1101/2024.02.05.578694

**Authors:** Lakshmi E. Batachari, Alyssa Y. Dai, Emily R. Troemel

## Abstract

Upon sensing viral RNA, mammalian RIG-I-like receptors activate downstream signals using caspase activation and recruitment domains (CARDs), which ultimately promote transcriptional immune responses that have been well-studied. In contrast, the downstream signaling mechanisms for invertebrate RIG-I-like receptors are much less clear. For example, the *Caenorhabditis elegans* RIG-I-like receptor DRH-1 lacks annotated CARDs and upregulates the distinct output of RNA interference (RNAi). Here we found that, similar to mammal RIG-I-like receptors, DRH-1 signals through two tandem caspase activation and recruitment domains (2CARD) to induce a transcriptional immune response. Expression of DRH-1(2CARD) alone in the intestine was sufficient to induce immune gene expression, increase viral resistance, and promote thermotolerance, a phenotype previously associated with immune activation. We also found that DRH-1 is required in the intestine to induce immune gene expression, and we demonstrate subcellular colocalization of DRH-1 puncta with double-stranded RNA inside the cytoplasm of intestinal cells upon viral infection. Altogether, our results reveal mechanistic and spatial insights into anti-viral signaling in *C. elegans,* highlighting unexpected parallels in RIG-I-like receptor signaling between *C. elegans* and mammals.

**Significance:** Viruses are ubiquitous pathogens that challenge diverse organisms, from bacteria to killer whales. While anti-viral defense has been well-studied in mammals, less is known about defense in invertebrates, including the roundworm *Caenorhabditis elegans*. Here we show that the *C. elegans* viral sensor DRH-1 shares similarities to a viral sensor in mammals called RIG-I. We find that DRH-1 has a signaling motif resembling the 2CARD motif, which is found in RIG-I and activates anti-viral immunity. We demonstrate that overexpression of DRH-1(2CARD) in *C. elegans* promotes resistance to viral infection, and that DRH-1 forms clusters inside intestinal cells during viral infection, similar to RIG-I in humans. Overall, these findings provide insights into *C. elegans* anti-viral immunity, highlighting similarities with mammalian anti-viral immunity.

## Introduction

Over the last several years, the study of immune systems across diverse hosts has uncovered far broader evolutionary conservation in cytosolic innate immune responses than previously appreciated. In particular, recent findings highlight the striking conservation of anti-viral immune pathways between bacteria and mammals (1). For example, the cyclic GMP-AMP synthase (cGAS)-stimulator of interferon genes (STING) pathway, which activates immune responses upon sensing cytosolic DNA in mammals, was recently identified in bacteria, where it detects bacteriophage infection and activates protective immune responses (2). In another example, cytosolic nucleotide-binding and leucine-rich repeat (NLR) receptors, which have long been known to be important in plant and mammalian immunity, have recently been identified in bacteria, where they also activate innate immune responses (3, 4).

A central tenet of cell-intrinsic innate immunity is the recognition of pathogen- or microbe-associated molecular patterns (PAMPs or MAMPs) by germline-encoded receptors, called pattern recognition receptors (PRRs) (5), such as the cGAS and NLR receptors mentioned above. While great strides have been made in understanding the conservation in PRRs between mammalian and bacterial hosts, less is known about the PRRs of invertebrate hosts. In particular, we have a limited understanding of the specific PAMPs/MAMPs sensed by the nematode *Caenorhabditis elegans* and the PRRs that sense these PAMPs/MAMPs (6–8). In general, *C. elegans* lacks canonical PRRs, such as cGAS-STING and NLR receptors, which recognize broad classes of PAMPs/MAMPs. Arguably, the only known class of PRR that detects a general PAMP/MAMP and has been shown to be conserved between *C. elegans* and mammals is the RIG-I-like receptor (RLR) class. Mammalian RLRs, such as RIG-I and MDA5, sense cytosolic RNA to activate a type-I interferon (IFN-I) response, which is a transcriptional response critical for host defense against viral infection.

*C. elegans* has three RIG-I-like receptors – Dicer-related helicase DRH-1, -2 and -3 – that share homology with mammalian RLRs at the helicase and C-terminal domains (CTD). Early work identified a role for DRH-1 in defense against RNA viruses through directing RNA interference (RNAi) to degrade viral RNA (9–13). The viruses used in these studies included the Orsay virus, a single-stranded positive-sense RNA virus that is the only known natural viral pathogen of *C. elegans* (14). One of these early studies on anti-viral RNAi demonstrated that the human RIG-I helicase and CTD could functionally substitute for the homologous domains in DRH-1, when measuring viral RNA levels (11). In mammals, the helicase domain and CTD bind dsRNA generated during RNA virus infection (15). Thus, the ability of human RIG-I helicase/CTD to function in place of DRH-1 helicase/CTD in *C. elegans* suggests that DRH-1 likely binds double-stranded RNA or other viral replication products. Consistent with this hypothesis, a recent cryo-EM analysis of DRH-1, along with its binding partners DCR-1 and RNA-binding protein RDE-4, indicates binding to blunt-ended double-stranded RNA (16).

Relative to the helicase/CTD, the N-terminal domains (NTDs) of RIG-I and DRH-1 share less sequence similarity (Fig. 1A). The NTD of RIG-I contains two tandem caspase activation and recruitment domains (2CARD), which are homotypic interaction motifs. In RIG-I, the 2CARD binds the helicase/CTD to form an autoinhibited configuration in the absence of infection. Upon viral infection, the RIG-I helicase/CTD binds viral RNA replication products, which releases the N-terminal 2CARD to activate downstream signaling (15). In support of this model, the initial identification of RIG-I found that ectopic expression of RIG-I(2CARD) alone was sufficient to induce type-I interferon (IFN-I) gene expression (17). The RIG-I(2CARD) can directly interact with the CARD found in downstream signaling factor MAVS, which promotes CARD oligomerization and downstream signal transduction to trigger the transcription of IFN-I genes. Because *C. elegans* lacks an obvious MAVS homolog and directs RNAi responses, the prevailing thought was that DRH-1 and mammalian RLRs had distinct signaling mechanisms during viral infection (18).

**Fig. 1.**
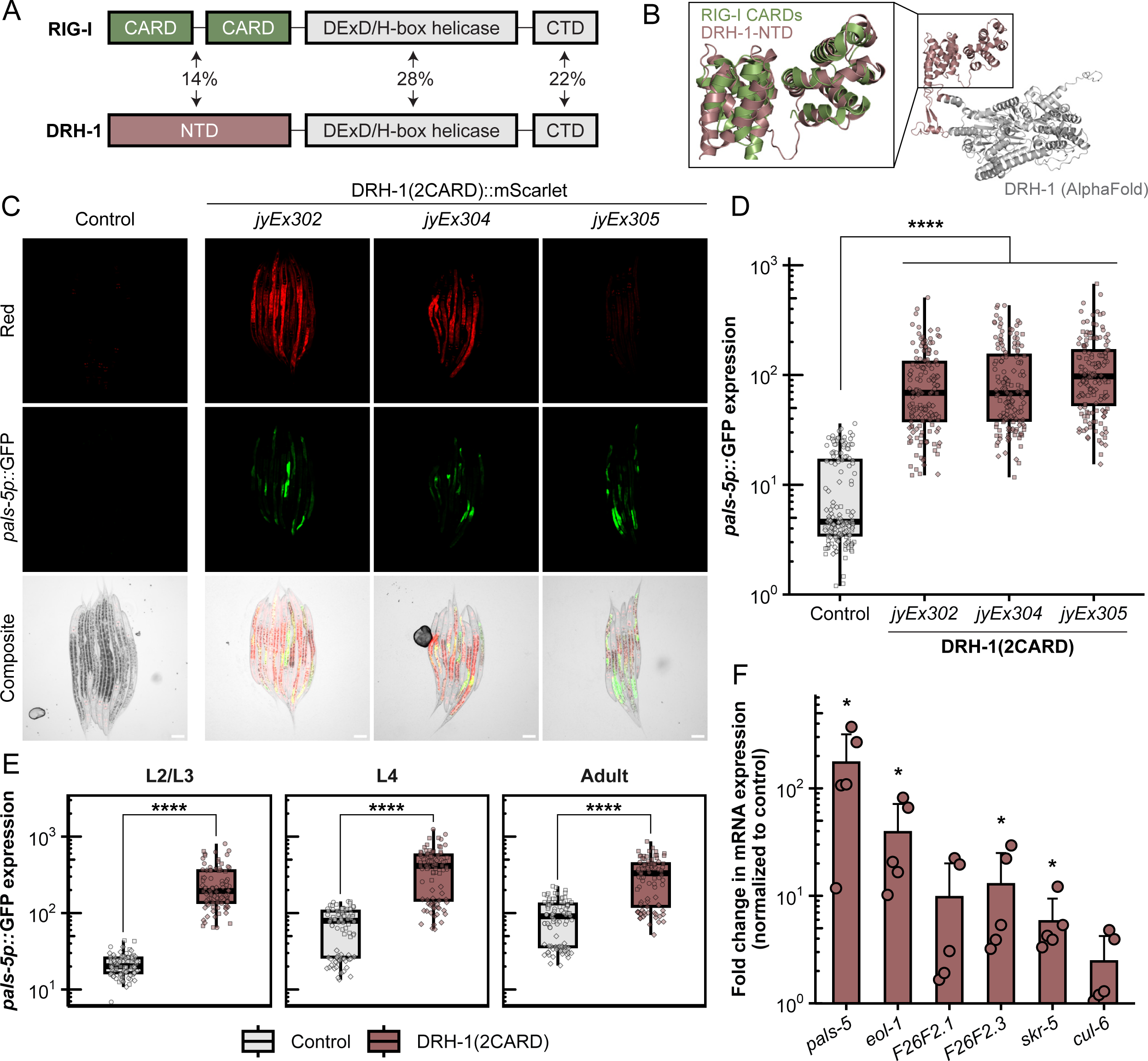
DRH-1 contains two predicted tandem CARD domains that activate *pals-5p::*GFP and IPR gene expression. (A) Comparison of domain architecture and amino acid sequence similarity between RIG-I and DRH-1. Pairwise sequence alignment performed with Clustal Omega. (B) AlphaFold prediction of DRH-1 protein structure (white and dark salmon) with superimposition of RIG-I CARDs (green) at the N-terminus (dark salmon). (C) Intestinal overexpression of DRH-1 NTD, or DRH-1(2CARD), induces *pals-5p::*GFP expression across three independent transgenic lines; *jyEx302, jyEx304*, and *jyEx305*. The designation “*jy<u>Ex</u>”* indicates that the DRH-1(2CARD) transgene is expressed from an <u>ex</u>trachromosomal array. The *myo-3p::*mCherry marker is expressed in body-wall muscle, and is only present in line *jyEx302*, whereas *myo-2p::*mCherry is part of the *jyIs8[pals-5p::gfp]* transgene and is constitutively expressed in the pharynx in all three DRH-1(2CARD) lines and the control. Scale bar = 50 μm. (D) Quantification of *pals-5p*::GFP in the control strain and DRH-1(2CARD) transgenic lines shown in (C). Each dot represents an individual animal; 150 animals were analyzed across three independent experiments for each strain. (E) DRH-1(2CARD) line *jyEx302* shows robust induction of *pals-5p::*GFP expression from larval stage L2 to adults. For each timepoint and genotype, 90 animals were analyzed across three independent experimental replicates. In both (D) and (E), different dot symbols indicate different experimental replicates. Horizontal lines in box-and-whisker plots represent median values, and the box reflects the 25^th^ to 75^th^ percentiles. A Mann-Whitney *U* test was used to determine statistical significance in expression values between each transgenic line and the control; ****p < 0.0001. (F) DRH-1(2CARD) overexpression in line *jyEx305* (without the *pals-5p::*GFP reporter) upregulates endogenous IPR gene expression. qRT-PCR analysis of a mixed-stage population containing both DRH-1(2CARD) transgenic animals and their non-transgenic siblings. Fold change in gene expression was determined relative to an *rde-1* mutant, non-transgenic control strain. Bars represents the mean across experimental replicates; error bars represent the standard deviation. Each dot represents a biological replicate (a plate with a minimum of 2000 animals); four independent experimental replicates were performed. A one-tailed *t*-test was used to calculate p-values; *p < 0.05.

In prior work, we demonstrated that DRH-1 had a separate role from regulating RNAi. Namely, we found that DRH-1 was required for activating a transcriptional immune response in *C. elegans* termed the Intracellular Pathogen Response (IPR) (19). The IPR is triggered by infection with diverse intracellular pathogens that infect the intestine, including the Orsay virus and obligate intracellular fungi called microsporidia (20). While work from other groups indicated that DRH-1 directs an anti-viral RNAi response through interactions with RNAi components, such as DCR-1 and RDE-4, and downstream signaling components RDE-1 and DRH-3 (9, 10, 12), we found that these RNAi factors are dispensable for IPR activation during viral infection (19). However, we did find that heterologous expression of a replication-competent Orsay virus RNA1 genome segment induced most of the IPR genes, dependent on DRH-1, again indicating that DRH-1 senses dsRNA or some other viral replication product (19, 21). Downstream of DRH-1, we have found that the bZIP transcription factor ZIP-1 activates a subset of the IPR genes (22), but otherwise it remains unclear how DRH-1 signals to induce the IPR upon viral infection.

In this study, we show that intestine-specific expression of the DRH-1 NTD alone is sufficient to induce the IPR in a partially ZIP-1-dependent manner. Despite the low primary sequence similarity between DRH-1 NTD and RIG-I(2CARD), we find that AlphaFold (23, 24) predictions indicate high three-dimensional similarity between these two proteins. Ectopic expression of DRH-1(2CARD) (aka DRH-1 NTD) in the intestine increases resistance to viral infection and promotes thermotolerance. Furthermore, we show that DRH-1 is required in the intestine for response to infection. Our subcellular analyses indicate that full-length DRH-1 protein forms puncta inside intestinal cells upon infection, and these puncta co-localize with viral and double-stranded RNA. Overall, these findings advance our understanding of how DRH-1 signals to activate a transcriptional immune response, revealing surprising similarities between mammalian RLRs and *C. elegans* DRH-1.

## Results

### The N-terminus of DRH-1 has two predicted tandem CARDs (2CARD) that activate IPR gene expression

The NTD of mammalian RIG-I contains two tandem CARDs, also known as 2CARD, which are each comprised of six alpha-helices (Fig. 1B). Mammalian CARDs have been extensively studied for their role in innate immune signaling pathways. In particular, CARDs mediate signal transduction through interactions with other CARDs. An example of CARD-CARD-mediated anti-viral signaling involves the interaction of RIG-I CARDs and the CARD found in downstream signaling protein MAVS (15). To investigate structural similarity in the NTDs of DRH-1 and RIG-I, we superimposed the CARDs of human RIG-I (PDB ID: 4p4h) onto the AlphaFold prediction of DRH-1 NTD. The two structures exhibited extensive three-dimensional similarity, including a pair of six anti-parallel alpha-helices that overlaid well between the N-terminal 2CARD of RIG-I and predicted structure for DRH-1 NTD (Fig. 1B). When comparing RIG-I to DRH-1, CARD1 and CARD2 have Root Mean Square Deviation (RMSD) values of 5.6 Å and 5.1 Å, respectively (0 Å is a perfect match). Of note, a comparison of RIG-I CARD1 to CARD2 gives an RMSD value of 5.1 Å, indicating the level of divergence that can be found between two different sequences both annotated as CARDs. Thus, the predicted structure for the NTD of DRH-1 resembles the two tandem CARDs found in the NTD of mammalian RLRs.

In an untargeted approach to identify proteins with structural similarity to the DRH-1 NTD, we used Foldseek (25) to identify matches between the predicted structure of DRH-1 NTD and known structures deposited in the Protein Data Bank (PDB). The top six structural matches corresponded to CARDs, with human and mouse 2CARDs at the top of the list (Table S1A). Next, we used the Dali structure alignment tool. Similar to the results from Foldseek, we found with this analysis that human RIG-I(2CARD) was the top structural match to DRH-1 NTD (Table S1B). Recent structural analysis has revealed that DRH-3, a distinct *C. elegans* RLR with a known role in RNAi (9, 10, 12) but so far not a demonstrated role in transcriptional responses (19), also contains two N-terminal tandem CARDs (26). Notably, the structure for DRH-3 CARDs (PDB ID: 6m6q) was listed as the 7th top structural match for DRH-1 NTD based on the comparisons performed with Dali (Table S1B). This finding lends additional support to the hypothesis that the NTD of DRH-1 contains 2CARD. For these reasons, as well as the RMSD values discussed above, we here-on refer to DRH-1 NTD as DRH-1(2CARD).

Though *C. elegans* lacks obvious IFN and IFN receptor homologs, a collection of findings indicates parallels in the regulation of the IPR in *C. elegans* and the regulation of IFN-I signaling in mammals (27). Given that ectopic expression of RIG-I(2CARD) is sufficient to induce IFN-I signaling in mammals (17), we explored whether DRH-1(2CARD) expression would be sufficient to induce IPR signaling in *C. elegans*. As Orsay virus infects intestinal cells, we used an intestine-specific promoter, *vha-6p*, to drive overexpression of mScarlet-tagged DRH-1(2CARD). Here we found that animals expressing DRH-1(2CARD) exhibited induction of the *pals-5p::*GFP reporter, a commonly used read-out for IPR induction. This effect was seen across three independent extrachromosomal transgenic lines (Fig. 1C, D), and was consistent across developmental stages from the second larval stage through adult (Fig. 1E). As a negative control, we created transgenic animals carrying a construct that lacks DRH-1(2CARD), but contains all other components of the vector, including *vha-6p* and mScarlet. *pals-5p::*GFP expression was not induced in that negative control strain (Fig. S1A). To confirm that the *pals-5p::*GFP reporter reflected endogenous gene expression, we performed qRT-PCR analysis and found that DRH-1(2CARD) significantly upregulated expression of endogenous *pals-5* mRNA, as well as other IPR genes (Fig. 1F and S1B).

The IPR includes hundreds of genes highly upregulated by viral infection, and about one-third of the top 80 genes depend on the bZIP transcription factor ZIP-1 for their expression, including *pals-5* (22, 28–30). To determine whether ZIP-1 was required for IPR induction by DRH-1(2CARD), we analyzed *pals-5p::*GFP in *zip-1* null mutants expressing the DRH-1(2CARD) transgene. Here, we found significantly reduced *pals-5p::*GFP expression in the *zip-1* mutant background compared to the wild-type background (Fig. 2A, B). ZIP-1::GFP localizes to the nucleus in response to known IPR triggers, such as Orsay virus infection and the proteasome blocker bortezomib (Fig. S2A, B) (22). However, DRH-1(2CARD) expression was not sufficient to cause obvious nuclear localization of ZIP-1::GFP. Of note, prior studies have demonstrated that *zip-1* is required to induce early IPR gene expression, and only after prolonged IPR activation does ZIP-1::GFP become visible in the nucleus by fluorescence microscopy (22). Thus, it may be that in the experiments shown in Fig. 2, DRH-1(2CARD) expression is not a potent enough trigger (compared to prolonged infection and proteasome blockade) to promote visible levels of nuclear ZIP-1::GFP. Regardless, our genetic results indicate that *zip-1* plays a significant role in mediating induction of *pals-5p*::GFP upon expression of DRH-1(2CARD).

**Fig. 2.**
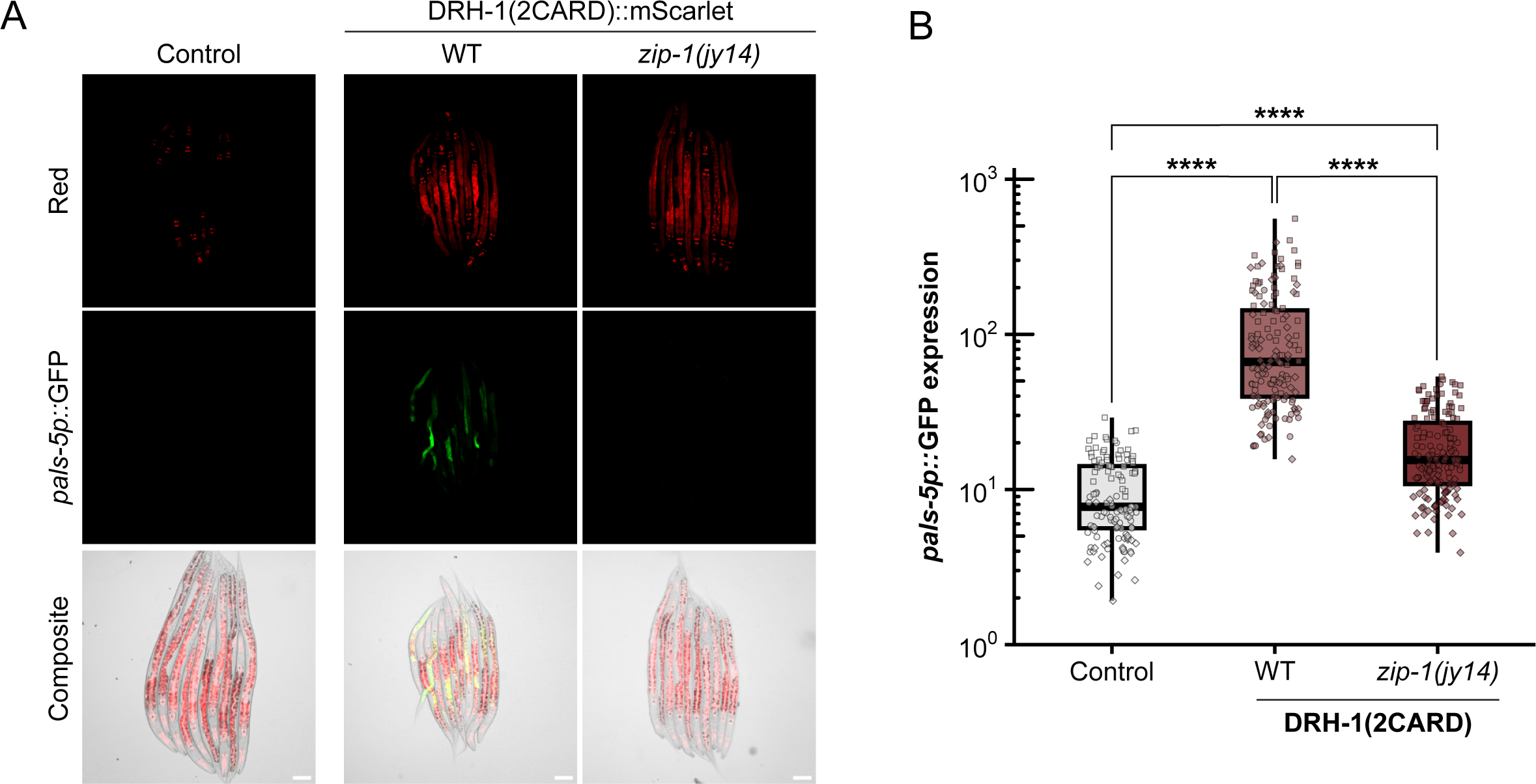
*pals-5p::*GFP upregulation by DRH-1(2CARD) is partially dependent on transcription factor *zip-1.* (A) DRH-1(2CARD) activates *pals-5p::*GFP in a manner that is partially dependent on transcription factor *zip-1. myo-2p::*mCherry is part of the *jyIs8[pals-5p::gfp]* transgene and is constitutively expressed in the pharynx of DRH-1(2CARD) transgenic lines, as well as in a control strain (left) that does express the DRH-1(2CARD) transgene. Scale bar = 50 μm. (B) Quantification of *pals-5p*::GFP in strains shown in panel A. Different dot symbols indicate different experimental replicates. Horizontal lines in box-and-whisker plots represent median values, and the box reflects the 25^th^ to 75^th^ percentiles. A Kruskal-Wallis test with Dunn’s multiple comparisons test was used to calculate p-values; ****p < 0.0001.

### DRH-1(2CARD) expression in the intestine induces several IPR phenotypes, including resistance to viral infection and heat shock

To determine whether the IPR gene induction caused by DRH-1(2CARD) leads to increased resistance to viral infection, we measured the infection rate in a population containing both DRH-1(2CARD) extrachromosomal transgenic animals and their non-transgenic siblings. Across three transgenic lines, animals expressing DRH-1(2CARD) exhibited a decreased infection rate relative to their non-transgenic siblings (Fig. 3A, B, S3A). In contrast to the viral infection rate phenotype, we did not observe significantly increased resistance to the microsporidian intracellular pathogen *Nematocida parisii* upon quantifying pathogen load (Fig. 3C, D). A priori, we would have expected to see increased resistance to *N. parisii*, as constitutive expression of IPR genes in other genetic backgrounds leads to increased resistance against both Orsay virus and *N. parisii* (30–33). Several potential reasons may explain this lack of effect, including the possibility that IPR gene expression is not high enough, or that IPR induction is required in tissues other than the intestine. Nevertheless, DRH-1(2CARD) provided robust protection against viral infection (Fig. 3B).

**Fig. 3.**
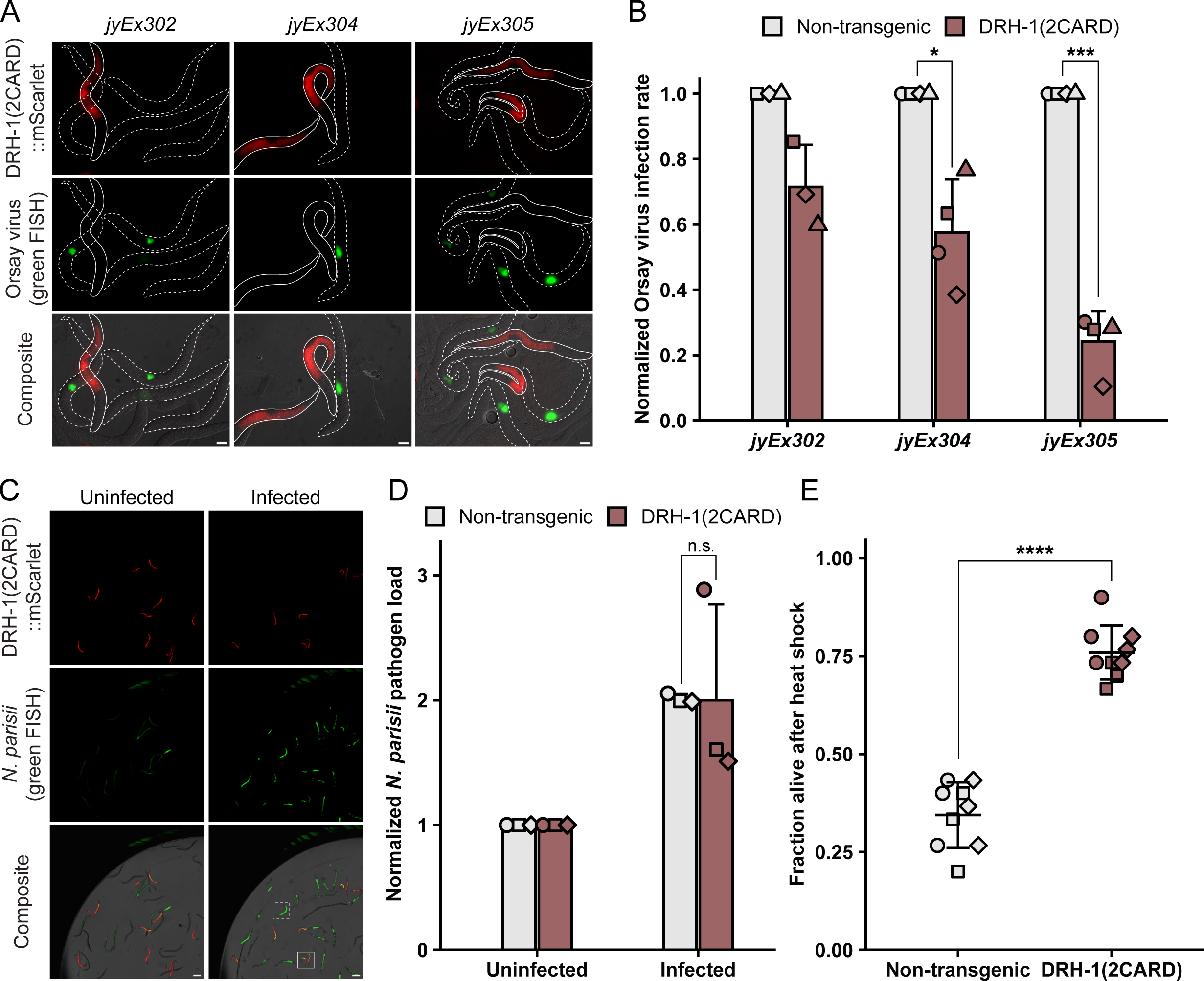
IPR activation by DRH-1(2CARD) reduces viral infection rate and increases thermotolerance, but does not decrease *N. parisii* pathogen load. (A) Representative images of DRH-1(2CARD) transgenic animals (solid white outline) and non-transgenic siblings (dotted white outline) infected with Orsay virus. Viral infection was visualized using fluorescein-conjugated (green) FISH probes targeting the Orsay virus genome. Virus-infected cells are shown in green. Scale bar = 20 μm. (B) DRH-1(2CARD) confers increased resistance to viral infection at 18 hours post inoculation (hpi) relative to non-transgenic siblings across all three transgenic lines. Within each transgenic line, the fraction of the population infected with virus was determined by scoring animals based on the presence of green fluorescence. Normalized infection rate was determined by setting the infection rate of non-transgenic siblings to 1. For transgenic lines *jyEx305* and *jyEx304*, 400 animals were scored across n = 4 independent experimental replicates. For line *jyEx302*, 300 animals were scored across n = 3 independent experimental replicates. A *t-*test was used to calculate p-values; ***p < 0.001; *p < 0.05. (C) Representative images of DRH-1(2CARD) transgenic animals and non-transgenic siblings infected with *N. parisii.* Infection was visualized by staining with green FISH probes targeting *N. parisii* ribosomal RNA. Green fluorescence in uninfected animals is due to autofluorescence. White boxes indicate the presence of *N. parisii* FISH fluorescent signal in DRH-1(2CARD) animals (solid border) and non-transgenic siblings (dashed border). Scale bar = 100 μm. (D) DRH-1(2CARD) expression does not reduce *N. parisii* pathogen load at 30 hpi. Normalized *N. parisii* infection rate was calculated by setting normalized green fluorescence values in uninfected worms to 1. Each dot represents an experimental replicate; three independent experimental replicates were performed. Uninfected: n = 1,011 (non-transgenic animals) or 1,435 (DRH-1(2CARD)). Infected: n = 925 (non-transgenic), 1,292 (DRH-1(2CARD)). A Mann-Whitney *U* test was used to determine significance; n.s. = not significant. (E) DRH-1(2CARD) animals exhibit increased thermotolerance relative to non-transgenic siblings. Survival was scored after a 2 h heat shock at 37.5°C followed by a 24 h incubation at 20°C. Nine biological replicates (n = 9 plates) were scored over three independent experimental replicates. 30 animals were scored per plate. A two-tailed *t-*test was used to calculate p-values; ****p < 0.0001. Mean values are represented by bar height (B, D) or cross bar (E); error bars represent the standard deviation. Different dot symbols indicate different experimental replicates.

In addition to pathogen resistance, activation of the IPR is associated with several other phenotypes, including slowed development and increased resistance to heat shock (29, 34). Indeed, we observed impaired development in DRH-1(2CARD) transgenic animals compared to their non-transgenic siblings (Fig. S3B, C). Furthermore, we found a substantial increase in thermotolerance, or resistance to heat shock. Specifically, ∼75% of DRH-1(2CARD) transgenic animals survived 24 hours (h) after a 2 h 37.5°C heat shock, compared to only ∼35% survival of their non-transgenic siblings (Fig. 3E). Taken together, these observations are consistent with the model that DRH-1(2CARD) expression activates the IPR, resulting in developmental and heat shock phenotypes similar to what has been previously observed in the context of other IPR triggers.

Next, we explored whether intestinal overexpression of full-length DRH-1 would promote the same phenotypes as overexpression of DRH-1(2CARD). However, in our attempts to generate transgenic lines carrying *vha-6p::drh-1::mScarlet*, we found that overexpression of full-length DRH-1 led to complete larval arrest, despite multiple attempts to generate transgenic animals (Fig. S3D). Interestingly, we found that 100% of these transgenic progeny (n = 103 transgenic progeny across four injections) exhibited *pals-5p::*GFP activation, indicative of IPR activation. Therefore, while the intestine-specific full-length DRH-1 appears to activate the IPR, the phenotype of complete larval arrest prevented our ability to assess other IPR phenotypes. Of note, we have previously observed complete larval arrest in genetic backgrounds that cause very strong IPR activation, such as null mutants of *pals-17*, a gene encoding a negative regulator of the IPR that is expressed predominantly in the intestine (33).

### DRH-1 is required in the intestine for IPR induction, and forms puncta upon infection that co-localize with viral RNA

The results above indicated that ectopic expression of DRH-1 alone in the intestine was sufficient to induce the IPR. scRNAseq analysis suggests that *drh-1* is expressed in intestinal tissue (35), and our prior work has shown that IPR activation can occur in either the intestine or the epidermis (29, 32). Therefore, we investigated whether DRH-1 is required in either of these tissues for IPR induction by implementing an experimental system that enables gene silencing in a tissue-specific manner. In particular, we used *C. elegans rde-1(ne300)* mutant strains where a wild-type copy of the RNAi factor *rde-1* is expressed only in the intestine or only in the epidermis, in order to rescue RNAi competency in these tissues (36). [Of note, *rde-1(ne300)* null mutants provide more tissue-specificity compared to the widely used partial loss-of-function *rde-1(ne219)* mutants, which are not fully RNAi-deficient.] Here we found that, similar to systemic RNAi against *drh-1,* intestine-specific knockdown of *drh-1* blocked *pals-5p::*GFP induction upon viral infection (Fig. 4). However, epidermis-specific knockdown of *drh-1* did not block *pals-5p::*GFP induction upon viral infection. Therefore, *drh-1* appears to be required in the intestine to induce the IPR upon viral infection.

**Fig. 4.**
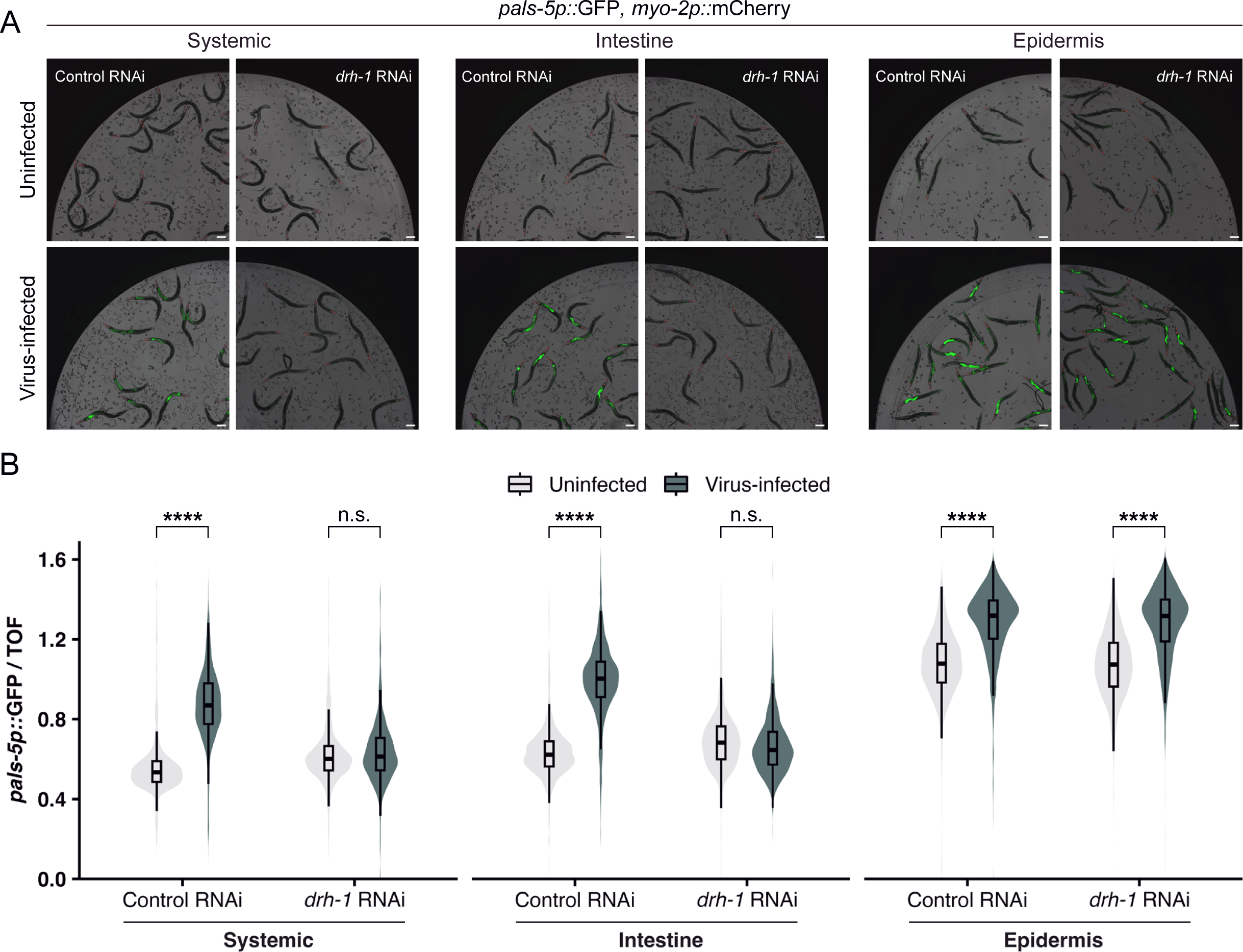
DRH-1 is required in the intestine to upregulate *pals-5p::*GFP expression during viral infection. (A) Representative images of *pals-5p*::GFP expression upon systemic, intestine-specific, or epidermis-specific knockdown of *drh-1* during viral infection. *myo-2p::*mCherry is a part of the *jyIs8[pals-5p::gfp]* transgene and is constitutively expressed in the pharynx. Animals were treated with an empty vector control RNAi (L4440) or *drh-1* RNAi and infected with virus at the L4 stage for 24 h. Scale bar = 100 μm. (B) Quantification of *pals-5p::*GFP fluorescence shown in (A) using a COPAS Biosort instrument. Knockdown of *drh-1* in the intestine, but not the epidermis, blocks the induction of *pals-5p::*GFP upon viral infection. *pals-5p*::GFP fluorescence is normalized to time-of-flight (a measure of worm size). For the epidermis-specific RNAi strain, the overall elevated levels of *pals-5p::*GFP is likely is due to the absence of functional *rde-1* in intestinal cells, resulting in impaired transgene silencing and increased baseline expression of the *pals-5p*::GFP transgene. Dots represent individual animals. Systemic: n = 961 animals (Control RNAi; uninfected), 741 (Control RNAi; infected), 1035 (*drh-1* RNAi; uninfected), or 841 (*drh-1* RNAi; uninfected). Intestine: n = 779 (Control RNAi; uninfected), 784 (Control RNAi; infected), 951 (*drh-1* RNAi; uninfected), or 924 (*drh-1* RNAi; uninfected). Epidermis: n = 1474 (Control RNAi; uninfected), 1564 (Control RNAi; infected), 1961 (*drh-1* RNAi; uninfected), or 1655 (*drh-1* RNAi; uninfected). Horizontal lines in box-and-whisker plots represent median values, and the box reflects the 25^th^ to 75^th^ percentiles. For each tissue-specific knockdown strain, a Mann-Whitney *U* test was used to determine significant differences between uninfected vs. infected worms after treatment with control or *drh-1* RNAi; ****p < 0.0001.

To further understand how DRH-1 signals in intestinal cells, we investigated the subcellular localization of DRH-1(2CARD) and full-length DRH-1 protein upon viral infection. First, we found that DRH-1(2CARD) was expressed in the cytoplasm and often formed large aggregates, both in virus-infected and uninfected animals (Fig. S4A). Given that DRH-1(2CARD) is only a fragment of the normal protein, we next sought to analyze the localization of full-length DRH-1. Because intestine-specific expression of full-length DRH-1 caused larval arrest (Fig. S3D), we turned to a previously generated strain containing a single-copy insertion of the *mScarlet::drh-1* transgene driven by the ubiquitous promoter *rpl-28p* (37). This strain exhibited normal development, possibly due to lower *drh-1* expression levels from a single-copy transgene, compared to higher expression from the previously described multi-copy, extrachromosomal transgene that caused arrest (Fig. S3D). Here, we found that expression of *rpl-28p::mScarlet::drh-1* did not lead to ectopic expression of IPR genes in the absence of infection (Fig. S5A). By crossing the *rpl-28p::mScarlet::drh-1* transgene into a *drh-1* null mutant background, we found that it rescued *drh-1* mutant phenotypes. In particular, we used qRT-PCR to show that *rpl-28p::mScarlet::drh-1* rescued mRNA expression of endogenous IPR genes during viral infection (Fig. S5A). Transgene expression also reduced the increased viral load phenotype of *drh-1* mutants (Fig. S5B). These results indicate that the *rpl-28p::mScarlet::drh-1* transgene in this strain is functional, so we used this strain to further analyze DRH-1 protein localization inside intestinal cells.

First, we infected mScarlet::DRH-1 transgenic animals with Orsay virus and visualized virus-infected cells by fluorescence in situ hybridization (FISH) after 24 h of infection. Uninfected control animals exhibited a homogenous distribution of mScarlet::DRH-1 throughout the intestinal cell cytoplasm (Fig. 5A). In contrast, upon infection, mScarlet::DRH-1 formed discrete puncta in virus-infected intestinal cells (Fig. 5A). When we quantified this effect, we found that 0% of uninfected animals exhibited DRH-1 puncta, while 100% of infected animals exhibited DRH-1 puncta (Fig. 5B). In these experiments, animals often had a mix of infected and uninfected intestinal cells, and in some of these cases we found DRH-1 puncta in uninfected intestinal cells. This result suggests there may be signaling from uninfected to infected intestinal cells to activate DRH-1 signaling (Fig. 5B), although an alternative possibility is that the Orsay virus FISH signal was below the level of detection in the cells we scored as uninfected. Next, we performed the same procedure, but also included an antibody that recognizes dsRNA, in order to assess whether DRH-1 may colocalize with this viral replication product. Indeed, we found that, in contrast to uninfected animals, infected animals exhibited extensive dsRNA antibody staining, which often co-localized with DRH-1 puncta (Fig. 5C). These findings are intriguing in light of studies in mammalian cells, where RIG-I forms puncta in response to viral infection and in the presence of viral RNA (38). RLR signal transduction occurs through CARD-carrying downstream signaling factor MAVS, which also forms puncta inside cells upon infection (39, 40). These mammalian studies in cell culture, paired with oligomerization studies in vitro (41, 42), highlight similarities to our findings here that *C. elegans* DRH-1 forms puncta that co-localize with viral replication products in the intestinal cells of an intact animal (Fig. 5).

**Fig. 5.**
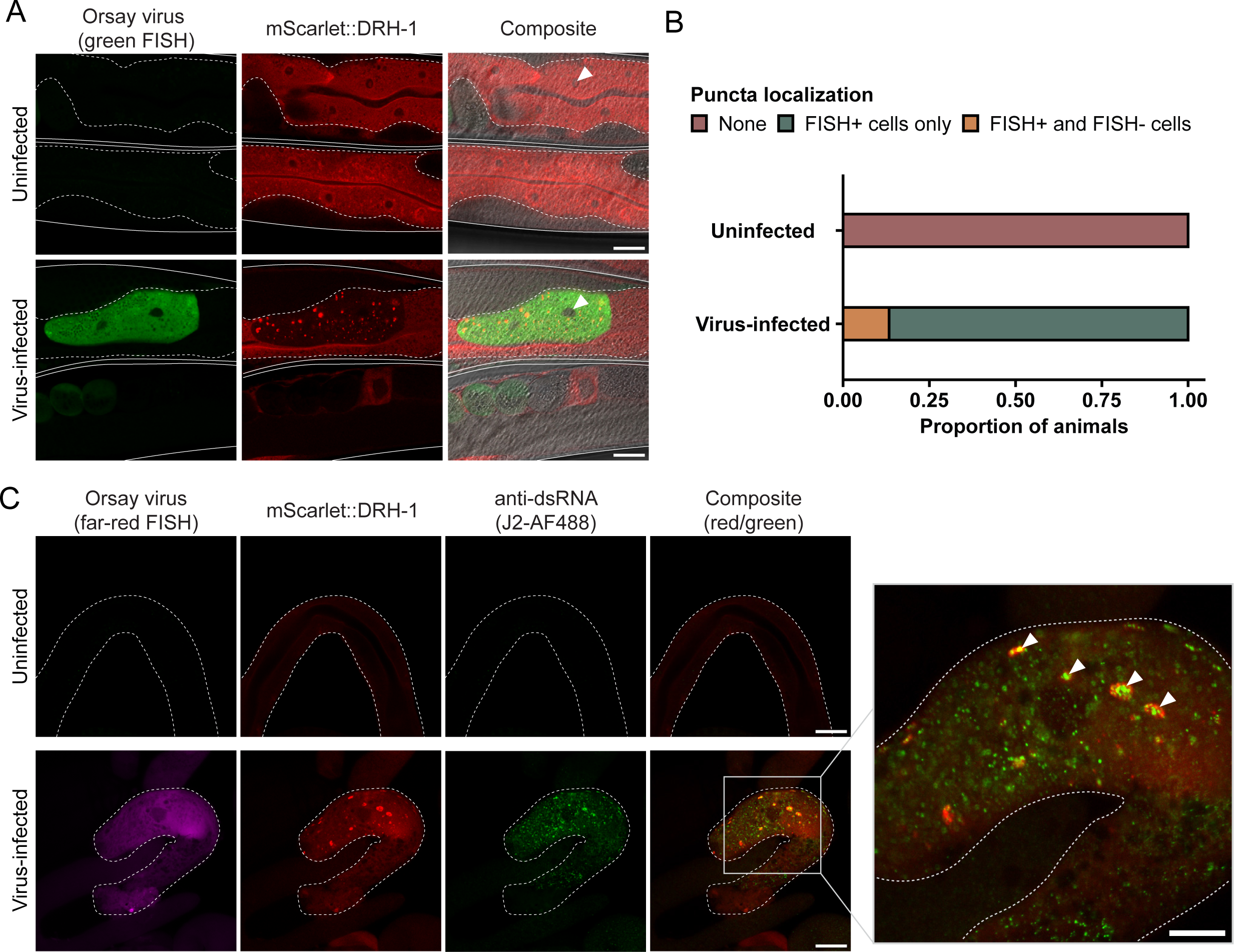
Virus-infected animals exhibit DRH-1 puncta in intestinal cells that co-localize with dsRNA. (A) Representative images of virus-infected adult animals that express mScarlet-tagged DRH-1 (red) as an integrated, single-copy transgene. Viral infection was visualized using fluorescein-conjugated (green) FISH probes targeting the Orsay virus genome. mScarlet::DRH-1 forms puncta (red) in virus-infected cells (green). Individual worms are outlined by solid white lines, and intestines are outlined by dotted white lines. White arrowheads indicate cell nuclei. Scale bar = 25 μm. (B) Quantification of puncta localization in uninfected animals or animals exposed to virus. DRH-1 puncta are only present in virus-infected animals. For each treatment condition, 30 animals were scored across at least three independent experimental replicates. (C) Representative images of dissected intestines (white dotted outline) from uninfected or virus-infected mScarlet::DRH-1 adult animals. Viral infection was visualized using Quasar 670-conjugated (far-red) FISH probes targeting the Orsay virus genome. The J2 antibody was used to visualize dsRNA. White arrowheads indicate co-localization between mScarlet::DRH-1 (red) and dsRNA (green) in virus-infected intestinal cells (magenta). Five worms were scored for each treatment condition. Scale bar = 25 μm.

## Discussion

Integrating in silico predictions with in vivo studies, our work indicates that *C. elegans* DRH-1/RLR contains signaling-competent tandem CARDs that can activate a transcriptional immune response. In the absence of a pathogen trigger, we demonstrated that intestine-specific overexpression of DRH-1(2CARD) promotes IPR activation, partially dependent on the transcription factor *zip-1* (Fig. 1, 2). IPR activation by DRH-1(2CARD) conferred increased resistance to viral infection and increased survival following heat shock (Fig. 3), which are both phenotypes that have been previously linked to IPR activation (43). Prior to this study, the tissue requirement for DRH-1 in the context of Orsay virus infection remained unclear, although recent findings support a role for DRH-1 acting in the intestine to combat age-related pathology (44). Our findings here with both tissue-specific expression (Fig. 1, 3, S3) as well as tissue-specific knock-down (Fig. 4) support a model in which DRH-1 functions in the intestine to induce immune gene activation upon viral infection. The intestine-specific requirement for DRH-1 is consistent with the observation that Orsay virus exhibits tropism for intestinal tissue (45). Furthermore, subcellular analysis of virus-infected intestinal cells reveals the presence of cytoplasmic DRH-1 puncta that co-localize with dsRNA (Fig 5). Taken together, our findings suggest that DRH-1 signals through N-terminal 2CARDs in intestinal cells to induce the IPR during viral infection.

CARD-containing proteins have long been known to be important in mammalian immunity for activating the IFN-I response (e.g. CARDs in RLRs) (15), as well as for activating inflammasome formation and cell death (e.g. CARDs in NLRs) (46), and more recently for coordinating defense in non-mammalian systems. As described in a recent pre-print, CARD-like domains were found to mediate cell death during phage infection as part of an immune defense system in the bacteria *Lysobacter enzymogenes* (4). Results from that study suggest that CARDs may constitute evolutionarily conserved immune signaling modules that likely originated from bacteria and are retained in mammals. In *C. elegans*, structural and functional analyses have described a role for CARDs in facilitating interactions between cell death components CED-3/caspase and CED-4/Apaf-1 to regulate apoptosis (47). Although apoptosis does not seem to have a prominent role in *C. elegans* immunity, CED-3/caspase and CED-4/Apaf-1 have been implicated in the restriction of viral replication in a vaccinia virus infection model (48). In the context of RLRs in *C. elegans*, crystal structures of DRH-3 have recently revealed two tandem CARDs at the N-terminus (26). While DRH-3 is a component of the anti-viral RNAi pathway, the signaling role of DRH-3(2CARD) remains to be determined. Notably, previous domain analysis of DRH-1 demonstrated that the NTD (2CARD) was required for reducing levels of viral RNA (11). In the same study, however, the authors found that expression of DRH-1 NTD was not sufficient to reduce viral RNA levels, which may be due to differing assays and strains used in that study compared to ours. Further investigation into the role of CARDs in *C. elegans* immunity may reveal a better understanding of mechanisms that govern anti-viral signaling, as well as the formation of signaling complexes, in an invertebrate host.

Formation of classical RLR signaling complexes require binding of RNA ligands, such as dsRNA and 5’ triphosphorylated RNA, to initiate protein oligomerization via CARD-CARD interactions (15). Signaling-competent oligomers then interact with CARDs found in MAVS, a mitochondrially localized signaling factor, to initiate a downstream signaling cascade (49). Although *C. elegans* lacks a known MAVS homolog, our subcellular localization analyses here indicate that DRH-1 forms puncta upon viral infection (Fig. 5), which may reflect protein oligomerization at a signaling hub, similar to what is observed in mammals. In addition, the observation that DRH-1 co-localizes with dsRNA antibodies inside viral-infected cells supports a model in which DRH-1 binds viral replication products or modified host RNAs generated during infection. While recent in vitro work has revealed that DRH-1 can bind blunt-end dsRNA, the native ligand for DRH-1 remains unknown (16). Furthermore, whether the dsRNA observed in our study originates from the host or the virus remains to be determined.

Canonical RLR signaling in mammals involves MAVS filament formation, as described above, followed by activation of the TBK-1 kinase, then phosphorylation and nuclear localization of the IRF3 transcription factor, which induces IFN-I gene expression and a systemic innate immune response (15). *C. elegans* lacks known homologs of TBK-1, IRF3 as well as MAVS, so its downstream signaling mechanisms are unknown (43). ZIP-1 is the only signaling factor identified so far that activates transcription downstream of DRH-1. ZIP-1 belongs to a branch of the bZIP transcription factor family that expanded in *C. elegans* (50), and does not have an obvious mammalian ortholog (22). In this study we found that ZIP-1 was partially required for DRH-1(2CARD) signaling, but it did not localize to the nucleus, in contrast to previously described nuclear translocation of ZIP-1 seen upon viral infection or other IPR triggers. In the endogenously tagged GFP strain used for our study, it might be that visible nuclear localization of ZIP-1 may require other triggers, or a trigger stronger than DRH-1(2CARD). Nonetheless, the partial requirement for ZIP-1 is consistent with prior studies (22), including those demonstrating that the IPR can be a systemic immune response in *C. elegans* (32), similar to the anti-viral IFN-I response in mammals.

In recent years, the study of immunity in diverse hosts has revealed remarkable evolutionary conservation in immune signaling pathways from eukaryotes to bacteria (1). Our findings expand upon prior work, which identified a novel role for DRH-1 in activating a transcriptional immune response (19), to uncover previously unappreciated insights into shared characteristics of RLR-mediated signaling between *C. elegans* and mammals. A hallmark of anti-viral immunity in mammals is the induction of systemic immunity triggered by secreted IFN-I ligands signaling through IFN receptors, which are ostensibly absent in *C. elegans*, although other ligand-receptor systems likely mediate systemic immunity as part of the IPR (32). Our work, however, supports the notion that RLR activation of the IFN-I response in mammals, and the IPR in *C. elegans,* resulted from divergent evolution of an ancient immune pathway for sensing cytosolic nucleic acid (43). Proteins involved in co-evolutionary host/pathogen battles commonly undergo amino acid sequence diversification (51). Therfore, the absence of sequence-based homologs does not necessarily indicate the absence of conservation. For example, the *Vibrio cholera* cGAS homolog (*dncV*) and human cGAS share high structural and functional similarity despite only sharing 10% primary sequence identity (52). Future efforts aimed at identifying signaling proteins downstream of DRH-1 in the *C. elegans* RLR pathway, such as determining whether there is a MAVS-like protein or a distinct downstream signal, may reveal novel anti-viral signaling factors involved in innate immune defense.

## Materials and Methods

### *In silico* protein structure analyses

The predicted protein structure of DRH-1 was obtained from the AlphaFold Protein Structure Database (http://alphafold.ebi.ac.uk/) (23, 24). RIG-I CARD domains (PDB ID: 4p4h) were superimposed onto the predicted structure of DRH-1 in PyMOL. PyMOL was also used to isolate the putative DRH-1 NTD (first 276 amino acids), which was queried against known structures in the PDB database using Foldseek (http://search.foldseek.com/search) (25) and Dali (http://ekhidna2.biocenter.helsinki.fi/dali/) (53, 54) protein structure comparison platforms.

### *C. elegans* Maintenance

*C. elegans* strains were maintained on Nematode Growth Media (NGM) agar plates containing streptomycin-resistant *Escherichia coli* OP50-1, unless otherwise specified. S2 Table lists all strains used in this study.

### Molecular cloning and transgenic *C. elegans* strains

Unless indicated otherwise, all constructs are expressed in an RNAi-deficient *rde-1* mutant background to reduce transgene silencing. For cloning *drh-1(2CARD)::mScarlet,* a cDNA sequence encoding the first 276 amino acids of *drh-1*, in addition to a C-terminal sequence-optimized mScarlet tag (also referred to as wrmScarlet) (55), was synthesized as a HiFi gBlock (Integrated DNA Technologies). The gBlock also included 50 base pair (bp) 5’ and 3’ homology to intestine-specific promoter *vha-6p* and *unc-54* 3’utr, respectively. Primers LEB001 and LEB002 were used to amplify off of template plasmid pET636 to generate a vector backbone containing *vha-6p,* an *unc-54* 3’utr, a 3xFLAG tag, and an ampicillin resistance cassette. The *drh-1(2CARD)::mScarlet* sequence was inserted into the vector backbone by Gibson assembly to generate the final construct pET770 [*vha-6p::drh-1(2CARD)::mScarlet::3xFLAG::unc-54 3’utr*]. Gibson assembly was performed using the NEBuilder HiFi DNA Assembly kit (New England Biolabs). 100 ng/μl of pET770 was co-injected along with a body wall muscle co-injection marker (17 ng/μl) into *jyIs8; rde-1* animals to create extrachromosomal array transgenic line *jyEx302* (ERT1076). Because DRH-1(2CARD)::mScarlet fluorescence was visible, additional transgenic lines were created without the use of a co-injection marker. In particular, extrachromosomal array lines *jyEx304* (ERT1207) and *jyEx305* (ERT1182) were generated by injecting 100 ng/μl of pET770 without a co-injection marker.

Figure 1C and 1D demonstrate that DRH-1(2CARD)::mScarlet transgenic lines *jyEx302*, *jyEx304* and *jyEx305* all induce the pals-5p::GFP reporter. However, specific lines were used for specific subsequent experiments for the following reasons. First, developmental analysis of *pals-5p::*GFP expression (Fig. 1E) was performed with transgenic line *jyEx302*, as this was the first line DRH-1(2CARD) line established. Next *jyEx305* was chosen for qRT-PCR (Fig. 1F), because it has the highest rate of transmission of the extrachromosomal array, increasing the percentage of transgenic animals in a large mixed population used to obtain enough RNA for analysis. This strain also has the lowest expression of mScarlet among all three transgenic lines, and for this reason we chose *jyEx305* for Fig. 2 and S2, to minimize red fluorescence bleed-through into the green channel, in order to better image *pals-5p::*GFP and *ZIP-1::*GFP. All three lines increased resistance to viral infection (Fig. 3A, B), but only transgenic line *jyEx304* was used for subsequent *N. parisii* infection and thermotolerance assays, as this line showed significantly decreased viral load, but also exhibited less developmental delay compared to *jyEx305*. Localization of DRH-1(2CARD) was assessed in transgenic line *jyEx304* (Fig S4A, B) as this line showed significantly decreased viral load (Fig. 3B) while still providing a sufficient number of infected transgenic animals for analysis.

To create the empty vector control, the *drh-1(2CARD)* sequence was deleted from pET770 using the Q5 Site-Directed Mutagenesis Kit (New England Biolabs). Primers LEB003 and LEB004 were used to amplify around the *drh-1(2CARD)* sequence in pET770. The linearized fragment was subsequently recircularized and transformed into chemically-comptent NEB DH5α cells (New England Biolabs) to generate the resulting plasmid pET786 [*vha-6p::mScarlet::unc-54 3’utr*]. 10 ng/μl of pET786 was injected into *jyIs8; rde-1* animals. Of note, the negative control construct was initially injected at 100 ng/μl, which is the injection concentration used for pET770 (see above). However, injections at this concentration resulted in extremely high levels of mScarlet expression, in which red fluorescence levels greatly exceeded that of DRH-1(2CARD)::mScarlet lines. Additionally, bleed-through of red fluorescence into the green channel used for imaging was observed in these animals, impairing assessment of *pals-5p::GFP* expression. Therefore, a decreased injection concentration of 10 ng/μl, was necessary to achieve protein expression levels comparable to that of DRH-1(2CARD)::mScarlet, as well as mitigate any off-target effects due to extremely high concentrations of mScarlet. Of note, the mScarlet expression in the negative control line was still higher than in the DRH-1(2CARD) lines. Filler DNA (pBluescript) was added to the injection mix to reach a final DNA concentration of 100 ng/μl.

For intestinal expression of full-length *drh-1,* an intron-containing sequence for *drh-1* was amplified from plasmid Tian233 using primers LEB005 and LEB006, which contain a 40 bp 5’ to *vha-6p* and a 36 bp 3’ homology to *mScarlet*. A vector backbone was generated by linearization of pET770, and PCR products were assembled using the NEBuilder HiFi DNA Assembly kit (New England Biolabs) to generate the final construct pET788. 100 ng/μl of pET788 was injected into *jyIs8; rde-1* animals.

All plasmid sequences were validated by whole-plasmid sequencing. Constructs used in this study can be found in S3 Table. Primer sequences are listed in S4 Table.

### Synchronization of *C. elegans*

To synchronize development, gravid adult animals were washed off plates with M9 media and transferred into a 15 ml conical tube. Worms were pelleted at 3,000 rpm for 30 sec and resuspended in 3 ml of M9 and 1 ml of bleaching solution (500 μl of 5.65–6% sodium hypochlorite solution and 500 μl of 2 M NaOH). After most adults had dissolved, M9 was added to a final volume of 15 ml. Tubes were centrifuged and supernatant was discarded. Released embryos were washed an additional four times with 15ml of M9 and resuspended in a final volume of 3 ml of M9. Embryos were incubated at 20 °C with continual rotation for 20-24 h to hatch L1s.

### *pals-5p::*GFP fluorescence measurements

DRH-1(2CARD) strains were synchronized by bleaching and grown at 20 °C for 24 h (L2/L3 larval stage), 44 h (L4 larval stage), or 68 h (adults). Worms were washed off plates, resuspended in M9, and anesthetized with 10 μM levamisole prior to imaging in a 96-well plate format on a ImageXpress Nano plate reader using the 4x objective (Molecular Devices, LLC, San Jose, CA). Background-corrected mean fluorescence intensity was quantified in Fiji by tracing the worm area and calculating the mean fluorescence intensity (MFI). To correct for background fluorescence, MFI of the well was subtracted from the MFI of each worm. 30-50 animals per genotype were analyzed for each experimental replicate across three independent experiments. For tissue-specific RNAi experiments, GFP fluorescence was measured using a COPAS Biosort instrument (Union Biometrica).

### RNA isolation and qRT-PCR

For DRH-1(2CARD) lines, 40 adults were transferred to 10-cm NGM plates due to smaller brood sizes. For non-transgenic controls, 20 adults were transferred to 10-cm NGM plates. As mentioned above, DRH-1(2CARD) lines were generated in a RNAi-defective *rde-1* mutant background to enhance transgene expression. Thus, non-transgenic control strains are either *rde-1* mutants with (Fig. S1B) or without (Fig. 1F) the *pals-5p::GFP* reporter. Two replicates (two plates) were set up per strain, allowed to have progeny (hundreds of animals) by incubating plates for 96 h at 20 °C. RNA was isolated from these progeny (thousands of animals) using TRI Reagent and 1-bromo-3-chloropropane (BCP) (Molecular Research Center) followed by isopropanol and ethanol washes. Pure RNA was resuspended in nuclease free water. Following cDNA synthesis using the iScript synthesis kit (Bio-Rad), qPCR was performed with iQ SYBR Green Supermix (Bio-Rad) on a CFX Connect Real-Time PCR Detection System (Bio-Rad). Relative gene expression ratios were determined by the Pfaffl method (56). Gene expression values were normalized to the expression of housekeeping gene *snb-1*. qRT-PCR primer sequences are listed in S4 Table.

### Bortezomib treatment

Proteasome inhibition by bortezomib (Selleck Chemicals) was performed as previously described (22). Synchronized L1 animals were plated on 6-cm NGM plates containing OP50-1 and grown for 44 h at 20 °C. Bortezomib in DMSO was top-plated to reach a final concentration in the agar of 20 μM. DMSO was used as a vehicle control. Plates were then incubated for 4 h at 20 °C prior to imaging. To assess ZIP-1::GFP localization, animals were anesthetized with 100 μM levamisole and imaged on a Zeiss LSM700 confocal microscope with Zen 2010 software.

### Orsay virus infections

All infections were performed using virus from the same batch of virus filtrate, which were prepared as previously described (28). Animals were infected with virus filtrate at the L1 or L4 stage. A minimum of two replicates (two plates) were set up per genotype per experiment. At least three independent experiments were performed per timepoint. For L1 infections, developmentally synchronized animals were exposed to a mixture of OP50-1 bacteria, M9, and virus filtrate for 18 h at 20 °C (Fig 3A, B). For L4 infections, synchronized animals were exposed to a mixture of OP50-1 bacteria, M9, and virus filtrate for 24 h at 20 °C (Fig 4, 5). For infection rate measurements and localization analyses, animals were washed in M9 and fixed in 4% paraformaldehyde for 15 min. Following fixation, worms were incubated at 47 °C overnight with fluorescein (FAM)- or Quasar 670-conjugated FISH probes that target Orsay virus RNA1 and RNA2 (Biosearch Technologies). Infection rate was assessed using a Zeiss AxioImager M1 compound microscope. For each experimental replicate, a minimum of 100 animals per genotype were scored for the presence of FISH fluorescent signal. For subcellular localization analyses, animals were imaged using a Zeiss LSM700 confocal microscope with Zen 2010 software. For tissue-specific knockdown experiments, animals were anesthetized with 10 μM levamisole prior to analysis using a COPAS Biosort instrument (Union Biometrica) to measure fluorescence.

### Microsporidia infections

*N. parisii* spores were prepared as previously described (57). A mixed population of DRH-1(2CARD) animals and non-transgenic siblings was synchronized at the L1 stage and plated on 6-cm NGM plates along with a mixture of OP50-1, M9, and one million spores for 30 h at 25°C. A minimum of four replicates (four plates) were set up per treatment. Three independent experiments were performed. To assess the presence of *N. parisii* meronts, animals were washed in M9 and fixed in 4% paraformaldehyde for 15 min prior to an overnight incubation at 47 °C with a FAM-conjugated FISH probe that hybridizes to *N. parisii* ribosomal RNA (Biosearch Technologies). Samples were sorted based on red fluorescence using the COPAS Biosort instrument (Union Biometrica) to obtain homogenous populations of either DRH-1(2CARD) animals or non-transgenic siblings. Green fluorescence was measured for each population using the COPAS Biosort instrument (Union Biometrica). For each genotype, the median green fluorescence of the infected population was normalized to the median fluorescence of the uninfected population.

### Development rate measurements

60 gravid adults were transferred onto a 10-cm NGM plate containing OP50-1 and incubated at 20 °C for 2 h. Adults were gently washed off of plates with M9 such that only eggs remained on the plate, and eggs were incubated at 20 °C for 48 h and 64 h. For each timepoint, development rate was determined by scoring the percentage of L4 stage or older animals. 100 animals were scored per genotype across three independent experimental replicates.

### Thermotolerance assays

Thermotolerance phenotypes were assayed as previously described (34). For each genotype, L4 animals (three plates; 30 L4s per plate) were transferred to NGM plates containing OP50-1. Animals were exposed to heat-shocked for 2 h at 37.5 °C in a dry incubator. Immediately following heat shock, plates transferred to a room temperature for 30 min. followed by incubation at 20 °C for 24 h. Survival was scored by assessing mobility, wherein dead worms were identified by failure to respond to a single touch using a worm pick. Three independent experimental replicates were performed on different days.

### Tissue-specific RNA interference

Systemic, intestine-specific, and epidermis-specific RNAi was performed via the feeding method. RNAi bacterial clones (*drh-1* and L4440 empty vector control) were inoculated in 5 ml LB containing 50 μg/ml carbenicillin and incubated in a 37 °C shaking incubator (250 rpm) for 16 h. Overnight cultures were seeded onto NGM plates supplemented with 5 mM IPTG and 1 mM carbenicillin. Seeded RNAi plates were incubated at room temperature for 4 days. Synchronized L1 animals were transferred to RNAi plates and grown at 20 °C for 44 h. Animals were then exposed to virus for 24 h and then analyzed for *pals-5p::*GFP expression on a COPAS Biosort instrument (Union Biometrica), as described above.

### Immunohistochemistry

For dsRNA localization analyses, synchronized L1s were plated onto 10-cm NGM plates and grown for 44 h at 20 °C to reach the L4 stage. Animals were then exposed to virus filtrate for 24 h at 20 °C, as described above. After 24 h, adults were washed off of plates with M9 and anesthetized in 100 μM levamisole. Intestines were dissected out of 100-200 adults and fixed for in 4% paraformaldehyde for 15 min. To visualize virus-infected cells, samples were incubated with Quasar 670-conjugated FISH probes that hybridize to Orsay virus RNA1 and RNA2 (Biosearch Technologies). Following a 6-8 h incubation at 47 °C, FISH probes were washed off and dissected intestines were incubated in block buffer (PBS, 0.5% Tween-20, 1 mM EDTA, 5% BSA, 0.05% NaN_3_) overnight at 4 °C. Dissected intestines were then stained with a 1:200 dilution of the anti-dsRNA antibody clone rJ2 (Sigma-Aldrich) for 2-4 h, followed by 10 μg/ml of goat anti-mouse IgG(H+L) cross-adsorbed secondary antibody conjugated to Alexa Fluor 488 (Invitrogen) for 1-2 h. Staining was performed in block buffer at room temperature.

### Statistics

All statistical analyses were performed in R. Q-Q plots were used to assess normality of the data, and parametric tests were used when appropriate. A nonparametric test was applied to data that did not meet the assumptions for a parametric test.

## Acknowledgements

We thank Matt Daugherty, Ethan Ewe, Aundrea Koger, Katie Li, Deevya Raman, Oded Rechavi, Alistair Russell, Max Strul, and Nicole Wernet for helpful comments on the manuscript. We thank Mario Bardan Sarmiento for making the Orsay virus filtrate preparations used in this study. We thank Kevin Corbett for help with protein structure alignment and Yating Chang for experimental assistance. We thank Nicole Wernet for crossing the tissue-specifc RNAi strains into a *jyIs8* background. We thank Gary Ruvkun for providing the *rpl-28p::mScarlet::drh-1* strain and expression construct. This work was supported by NIH under R01 AG052622, GM114139, AI176639 and by NSF 2301657 to E.R.T.

## Supporting Information

**S1 Fig.**
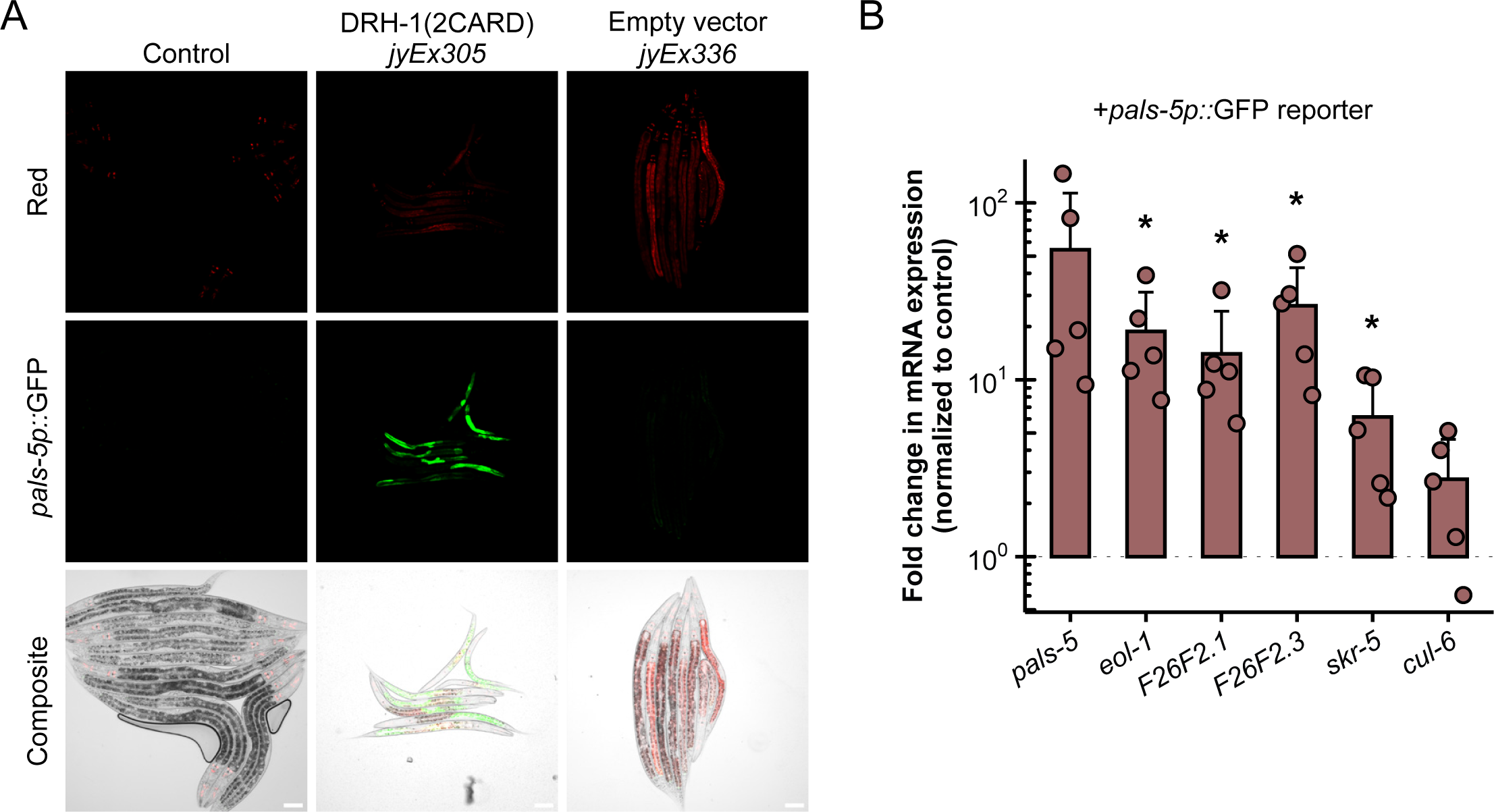
DRH-1(2CARD) specifically induces *pals-5p::*GFP. (A) Representative images showing *pals-5p::*GFP induction in DRH-1(2CARD) transgenic animals (line *jyEx305*) and absence of *pals-5p*::GFP induction in empty vector control (line *jyEx336*). *myo-2p::*mCherry is a part of the *jyIs8[pals-5p::gfp]* transgene and is constitutively expressed in the pharynx. Scale bar = 50 μm. (B) qRT-PCR analysis of DRH-1(2CARD) line *jyEx305* (expresses *pals-5p::*GFP reporter) shown in (A). RNA was extracted from a mixed-stage population containing both DRH-1(2CARD) transgenic animals and their non-transgenic siblings. Fold change in gene expression was determined relative to a non-transgenic control strain (*rde-1* mutant in a *pals-5p::*GFP background). Bars represent the mean across experimental replicates; error bars represent the standard deviation. Each dot represents a biological replicate (a plate with a minimum of 2000 animals); four independent experimental replicates were performed. A one-tailed *t*-test was used to calculate p-values; *p < 0.05.

**S2 Fig.**
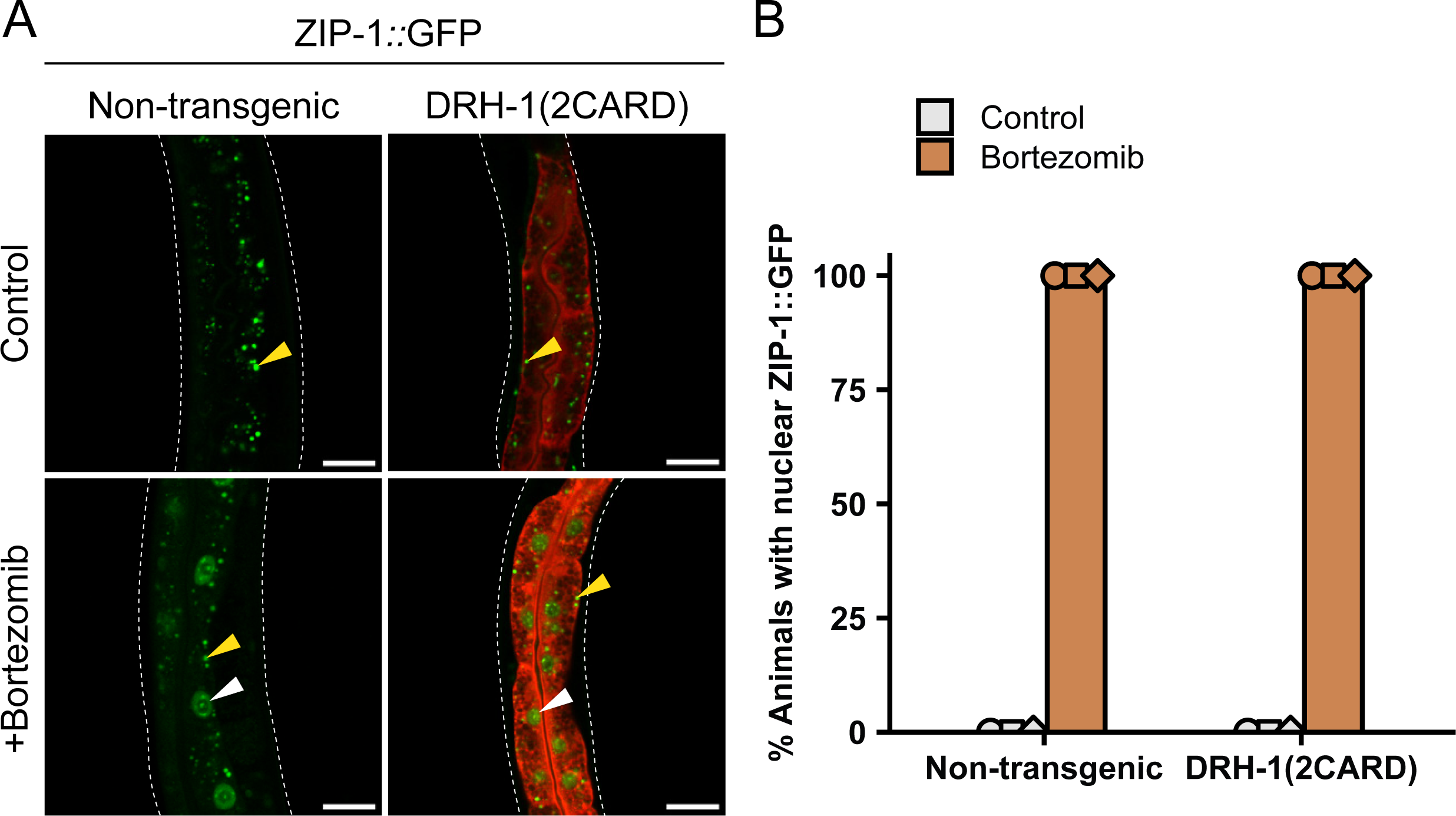
DRH-1(2CARD) expression does not promote nuclear localization of ZIP-1::GFP. Representative images showing ZIP-1::GFP expression in DRH-1(2CARD) transgenic animals and non-transgenic siblings. ZIP-1::GFP is not visible in the nuclei of DRH-1(2CARD) animals. Bortezomib treatment was used as a positive control for nuclear localization of ZIP-1::GFP. White arrowheads indicate ZIP-1::GFP expression in the nucleus. Yellow arrowheads indicate autofluorescence from intestinal gut granules. Scale bar = 25 μm. (B) ZIP-1::GFP is not present in intestinal nuclei of untreated animals, but is expressed in 100% of animals treated with bortezomib. Localization pattern of ZIP-1::GFP is the same for both DRH-1(2CARD) animals and non-transgenic siblings. For each genotype and treatment, 45 total animals were scored. Bars represents the mean across biological replicates; error bars represent the standard deviation. Each dot represents a biological replicate (a plate with 15 animals).

**S3 Fig.**
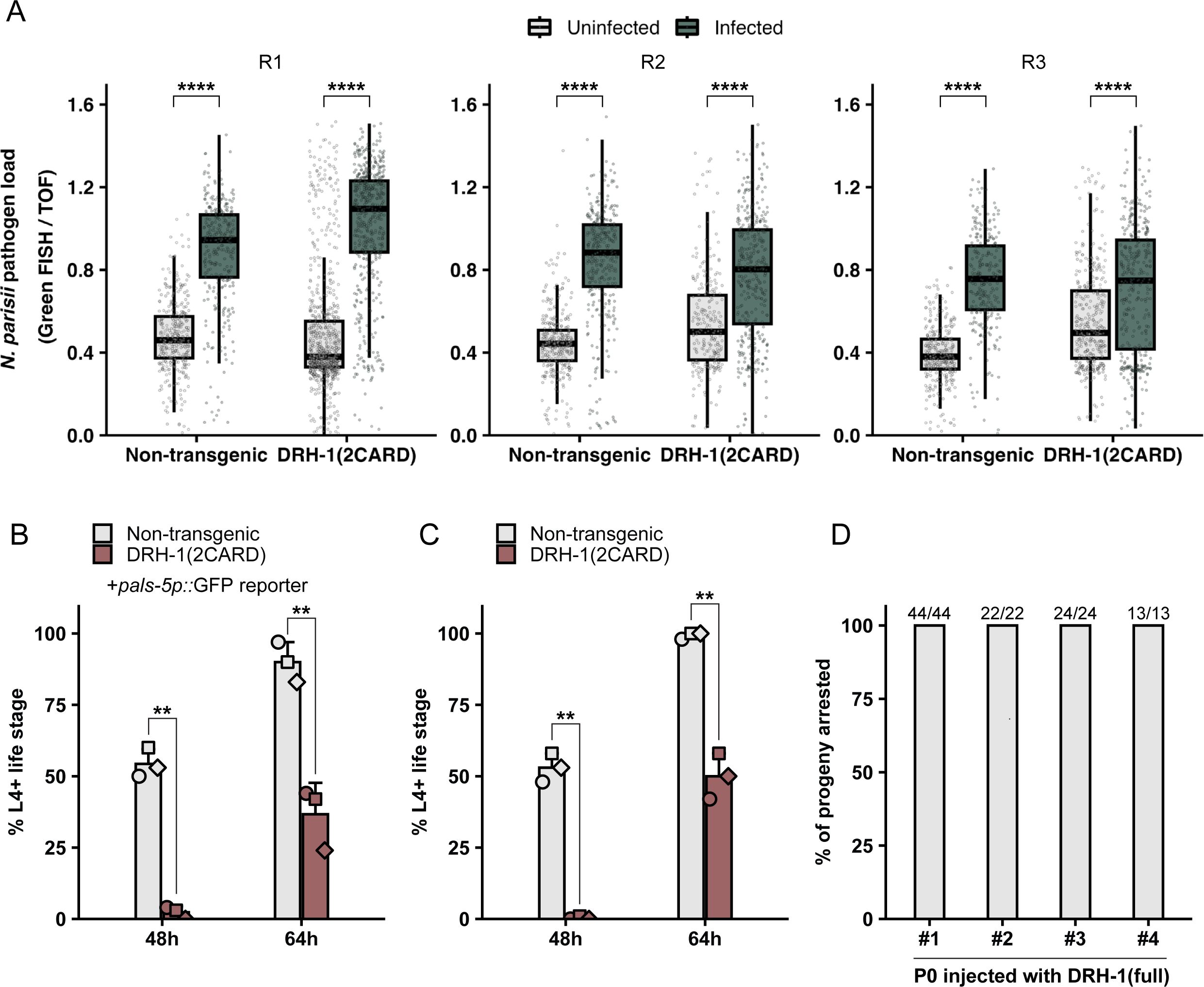
DRH-1(2CARD) expression does not reduce *N. parisii* pathogen load and delays development. (A) Quantification of *N. parisii* pathogen load in individual experiments (combined results shown in Fig. 3D) by FISH using fluorescein-conjugated (green) probes that target *N. parisii* ribosomal RNA. A COPAS Biosort instrument was used to measure green fluorescence normalized to time-of-flight (a measure of worm size). Dots represent individual animals. Left (R1): n = 353 (non-transgenic; uninfected), 317 (non-transgenic; infected), 791 (DRH-1(2CARD); uninfected), or 535 (DRH-1(2CARD); infected). Center (R2): n = 336 (non-transgenic; uninfected), 337 (non-transgenic; infected), 260 (DRH-1(2CARD); uninfected), or 385 (DRH-1(2CARD); infected). Right (R3): n = 322 (non-transgenic; uninfected), 271 (non-transgenic; infected), 384 (DRH-1(2CARD); uninfected), or 372 (DRH-1(2CARD); infected). Horizontal lines in box-and-whisker plots represent median values, and the box reflects the 25^th^ to 75^th^ percentiles. Each panel displays data from an independent experimental replicate. A Mann-Whitney *U* test was used to calculate p-values; ****p < 0.0001. (B) DRH-1(2CARD) animals exhibit delayed development relative to non-transgenic siblings in a strain background with (B) or without (C) the *pals-5p::*GFP reporter. Bars represent the mean across experimental replicates; error bars represent the standard deviation. A two-tailed *t-*test was used to calculate p-values; **p < 0.01. (D) Intestinal overexpression of full-length DRH-1 leads to developmental arrest at L1/L2 larval stages in 100% of transgenic progeny across four injections. The fraction of transgenic progeny exhibiting larval arrest is displayed above each bar.

**S4 Fig.**
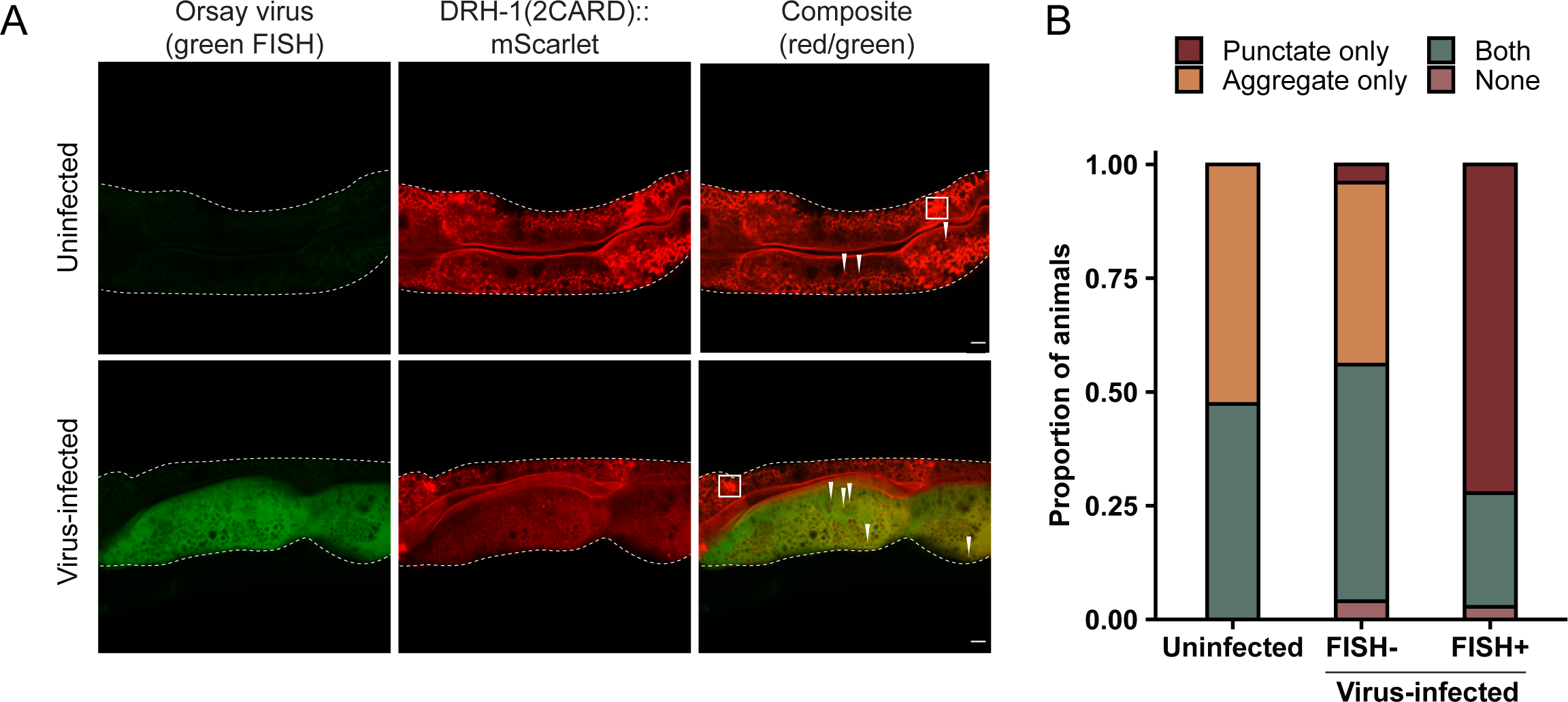
Uninfected intestinal cells predominantly exhibit DRH-1(2CARD) aggregates, whereas virus-infected cells are enriched for DRH-1(2CARD) puncta. In the absence of infection, all animals exhibit DRH-1(2CARD) aggregates in the intestinal cytoplasm (white box), with some animals exhibiting both DRH-1(2CARD) aggregates and puncta (white arrowhead). In infected animals, the majority of virus-infected cells (green FISH staining) contain DRH-1(2CARD) puncta only, with some animals exhibiting both puncta and aggregates. In the same virus-infected animals, most uninfected neighboring cells (no green FISH staining) show DRH-1(2CARD) aggregates. Intestine is outlined by white dashed line. Proportions determined by scoring 38 (uninfected) or 36 (virus-infected) animals across three independent experimental replicates.

**S5 Fig.**
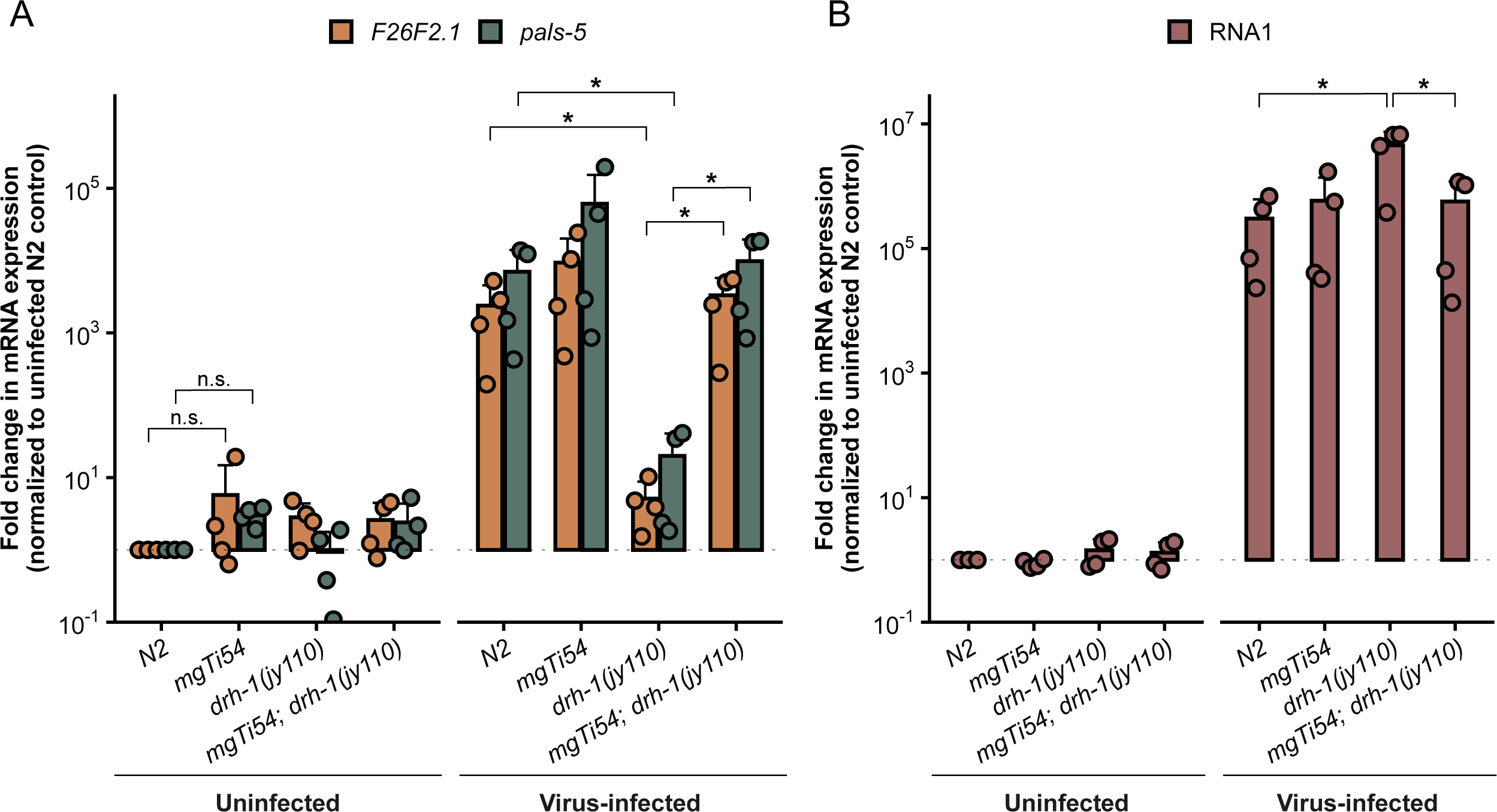
Expression of the *rpl-28p::mScarlet::drh-1* transgene rescues *pals-5* expression and Orsay RNA1 levels upon viral infection. (A) During viral infection, expression of the mScarlet::DRH-1 transgene (*mgTi54)* in a *drh-1(jy110)* deletion mutant is sufficient to rescue mRNA levels of (A) IPR genes *F26F2.1* and *pals-*5, as well as (B) Orsay RNA1 to WT levels. qRT-PCR analysis of *F26F2.1*, *pals-5*, and Orsay RNA1 in WT animals (N2), a *drh-1(jy110)* deletion mutant, and animals that express the *mgTi54* transgene in a *drh-1(jy110)* or WT background. Fold change in gene expression was determined relative to the uninfected WT (N2) control. Bars represent the mean; error bars represent the standard deviation. Each dot represents a biological replicate (a plate with 2000 animals); three independent experimental replicates were performed. A one-sample Wilcoxon signed rank test was used to compare the distribution of values against a hypothetical value of 1 in the uninfected group. A Mann-Whitney *U* test was used to calculate p-values for comparisons between samples in the infected group; *p < 0.05.

## Supplementary Tables – See supporting information

**S1 Table. Foldseek and Dali analyses.**

**S2 Table. List of strains used in this study.**

**S3 Table. Constructs used in this study.**

**S4 Table. Primers used in this study.**

