## Supplemental Tables for "*C. elegans* RIG-I-like receptor DRH-1 signals via CARDs to activate anti-viral immunity in intestinal cells"

**S1A Table.** Foldseek analysis

| No. | Target (PDB ID) | E-value | TM-score | Alignment length | Sequence identity (%) | Description | Organism |
|---|---|---|---|---|---|---|---|
| 1 | 4p4h.cif.gz_C | 3.84E-01 | 3.59E-01 | 204 | 13.7 | Caught-in-action signaling complex of RIG-I 2CARD domain and MAVS CARD domain | *Homo sapiens* |
| 2 | 6e25.cif.gz_MODEL_10_A | 3.52E+00 | 6.28E-01 | 103 | 14.5 | NMR solution structure of the CARD9 CARD bound to zinc | *Homo sapiens* |
| 3 | 6e25.cif.gz_MODEL_3_A | 3.05E+00 | 5.68E-01 | 98 | 14.2 | NMR solution structure of the CARD9 CARD bound to zinc | *Homo sapiens* |
| 4 | 1wxp.cif.gz_MODEL_1_A | 3.69E+00 | 5.66E-01 | 94 | 12.7 | Solution structure of the death domain of nuclear matrix protein p84 (THOC1) | *Homo sapiens* |
| 5 | 6e25.cif.gz_MODEL_4_A | 5.38E+00 | 5.36E-01 | 97 | 14.4 | NMR solution structure of the CARD9 CARD bound to zinc | *Homo sapiens* |
| 6 | 7dni.cif.gz_B | 3.36E+00 | 3.69E-01 | 220 | 8.1 | MDA5 CARDs-MAVS CARD polyUb complex | *Homo sapiens* |
| 7 | 2n7z.cif.gz_MODEL_2_A | 8.61E+00 | 5.89E-01 | 85 | 16.4 | Solution structure of RIP2 CARD | *Homo sapiens* |
| 8 | 6gwm.cif.gz_MODEL_3_A | 9.92E+00 | 5.17E-01 | 90 | 13.3 | Solution structure of rat RIP2 caspase recruitment domain | *Rattus norvegicus* |
| 9 | 5uxv.cif.gz_B | 5.38E+00 | 3.34E-01 | 116 | 11.2 | Crystal Structure of Anti-anti-sigma factor PhyR I40V/S51C mutant from Bartonella quintana | *Bartonella quintana str. Toulouse* |
| 10 | 7ml4.cif.gz_M | 4.89E+00 | 2.59E-01 | 247 | 11.7 | RNA polymerase II initially transcribing complex (ITC) | *Saccharomyces cerevisiae* |
| 11 | 4g97.cif.gz_A | 9.92E+00 | 3.02E-01 | 103 | 11.6 | Crystal structure of the response regulator PhyR from Brucella abortus | *Brucella abortus 2308* |
| 12 | 1h0o.cif.gz_A | 3.69E+00 | 1.52E-01 | 237 | 8 | Cobalt substitution of mouse R2 ribonucleotide reductase to model the reactive diferrous state | *Mus musculus* |
| 13 | 8c3w.cif.gz_A | 8.61E+00 | 1.69E-01 | 241 | 13.2 | Crystal structure of a computationally designed heme binding protein, dnHEM1 | Synthetic construct |
| 14 | 8g0c.cif.gz_d | 6.19E+00 | 1.52E-01 | 258 | 13.5 | Cryo-EM structure of TBAJ-876-bound Mycobacterium smegmatis ATP synthase rotational state 1 (backbone model) | *Mycolicibacterium smegmatis MC2 155* |
| 15 | 3g33.cif.gz_B | 9.92E+00 | 1.53E-01 | 247 | 10.5 | Crystal structure of CDK4/cyclin D3 | *Homo sapiens* |
| 16 | 3v71.cif.gz_A | 8.61E+00 | 1.66E-01 | 212 | 15 | Crystal structure of PUF-6 in complex with 5BE13 RNA | *Caenorhabditis elegans* |
| 17 | 3e4a.cif.gz_B | 9.92E+00 | 1.92E-01 | 85 | 17.6 | Human IDE-inhibitor complex at 2.6 angstrom resolution | *Homo sapiens,* synthetic construct |
| 18 | 5ivw.cif.gz_2 | 7.13E+00 | 1.44E-01 | 270 | 11.1 | Human core TFIIH bound to DNA within the PIC | *Homo sapiens* |
| 19 | 6ivn.cif.gz_C | 8.61E+00 | 1.37E-01 | 217 | 12.9 | Crystal structure of a membrane protein G264A | *Klebsiella pneumoniae* |

**S1B Table.** Dali analysis

| No | Chain | Z | rmsd | lali | nres | %id | Description |
|---|---|---|---|---|---|---|---|
| 1 | 6bzh-B | 8.9 | 6.8 | 104 | 192 | 9 | MOLECULE: PROBABLE ATP-DEPENDENT RNA HELICASE DDX58; |
| 2 | 5juy-B | 8.8 | 13.5 | 112 | 1234 | 10 | MOLECULE: APOPTOTIC PROTEASE-ACTIVATING FACTOR 1; |
| 3 | 6n2m-A | 8.7 | 3.5 | 103 | 142 | 9 | MOLECULE: CASPASE RECRUITMENT DOMAIN-CONTAINING PROTEIN 9; |
| 4 | 7dni-A | 8.7 | 2.6 | 86 | 199 | 13 | MOLECULE: MDA5 CARD; |
| 5 | 6xkj-A | 8.2 | 2.7 | 82 | 85 | 10 | MOLECULE: CASPASE RECRUITMENT DOMAIN-CONTAINING PROTEIN 8; |
| 6 | 2of5-H | 8 | 2.8 | 89 | 100 | 6 | MOLECULE: DEATH DOMAIN-CONTAINING PROTEIN CRADD; |
| 7 | 6m6q-B | 7.9 | 5.1 | 176 | 315 | 9 | MOLECULE: DICER-RELATED HELICASE 3; |
| 8 | 6mks-A | 7.8 | 2.9 | 85 | 90 | 9 | MOLECULE: CHIMERA PROTEIN OF NLR FAMILY CARD DOMAIN-CONTAIN |
| 9 | 4d8o-A | 7.7 | 4.1 | 80 | 509 | 11 | MOLECULE: ANKYRIN-2; |
| 10 | 6gwm-A | 7.6 | 4.3 | 111 | 125 | 11 | MOLECULE: RECEPTOR-INTERACTING SERINE/THREONINE-PROTEIN KIN |
| 11 | 2yqf-A | 7.6 | 5.2 | 96 | 111 | 16 | MOLECULE: ANKYRIN-1; |
| 12 | 4e9m-A | 7.5 | 2.8 | 79 | 97 | 6 | MOLECULE: NUCLEOTIDE-BINDING OLIGOMERIZATION DOMAIN-CONTAIN |
| 13 | 5wvc-B | 7.5 | 2.9 | 89 | 103 | 12 | MOLECULE: APOPTOTIC PROTEASE-ACTIVATING FACTOR 1; |
| 14 | 6ac0-A | 7.5 | 3.8 | 92 | 115 | 11 | MOLECULE: TUMOR NECROSIS FACTOR RECEPTOR TYPE 1-ASSOCIATED |
| 15 | 2mb9-A | 7.2 | 3.4 | 96 | 106 | 8 | MOLECULE: B-CELL LYMPHOMA/LEUKEMIA 10; |
| 16 | 4m9s-B | 7 | 15.5 | 131 | 511 | 9 | MOLECULE: CELL DEATH PROTEIN 4; |
| 17 | 4dwn-A | 6.8 | 2.8 | 87 | 97 | 9 | MOLECULE: BCL10-INTERACTING CARD PROTEIN; |
| 18 | 2gf5-A | 6.8 | 3.8 | 93 | 191 | 12 | MOLECULE: FADD PROTEIN; |
| 19 | 3t6p-A | 6.8 | 3.5 | 95 | 330 | 11 | MOLECULE: BACULOVIRAL IAP REPEAT-CONTAINING PROTEIN 2; |
| 20 | 4uz0-A | 6.7 | 2.7 | 80 | 87 | 11 | MOLECULE: NUCLEOLAR PROTEIN 3; |
| 21 | 1ich-A | 6.5 | 2.9 | 79 | 87 | 15 | MOLECULE: TUMOR NECROSIS FACTOR RECEPTOR-1; |
| 22 | 2o71-A | 6.5 | 3.3 | 86 | 91 | 9 | MOLECULE: DEATH DOMAIN-CONTAINING PROTEIN CRADD; |
| 23 | 1wxp-A | 6.4 | 4.6 | 93 | 110 | 11 | MOLECULE: THO COMPLEX SUBUNIT 1; |
| 24 | 7cmg-A | 5.6 | 10.8 | 83 | 487 | 8 | MOLECULE: POR SECRETION SYSTEM PROTEIN PORM/GLDM; |
| 25 | 1ddf-A | 5.5 | 4.8 | 106 | 127 | 12 | MOLECULE: FAS; |
| 26 | 7k3r-A | 5.5 | 3 | 78 | 95 | 13 | MOLECULE: INTERFERON-INDUCIBLE PROTEIN AIM2; |
| 27 | 3mop-K | 5.4 | 2.9 | 77 | 93 | 9 | MOLECULE: MYELOID DIFFERENTIATION PRIMARY RESPONSE PROTEIN |
| 28 | 2km6-A | 5.4 | 3.1 | 79 | 96 | 13 | MOLECULE: NACHT, LRR AND PYD DOMAINS-CONTAINING PROTEIN 7; |
| 29 | 1ygo-A | 5.2 | 3.1 | 86 | 110 | 7 | MOLECULE: PROBABLE SERINE/THREONINE-PROTEIN KINASE PELLE; |
| 30 | 7bso-A | 5.2 | 3.1 | 78 | 90 | 4 | MOLECULE: NACHT, LRR AND PYD DOMAINS-CONTAINING PROTEIN 9; |
| 31 | 2mjm-A | 5.2 | 3.4 | 82 | 101 | 11 | MOLECULE: PROTEIN NLRC5; |
| 32 | 6jcc-A | 5.2 | 3.3 | 79 | 97 | 13 | MOLECULE: COMPUTATIONAL DESIGNED PROTEIN BASED ON EVOLUTION |

| | | | | | | | |
|---|---|---|---|---|---|---|---|
| 33 | 3mop-A | 5 | 3 | 80 | 105 | 9 | MOLECULE: MYELOID DIFFERENTIATION PRIMARY RESPONSE PROTEIN |
| 34 | 3qf2-B | 4.9 | 2.9 | 73 | 106 | 12 | MOLECULE: NACHT, LRR AND PYD DOMAINS-CONTAINING PROTEIN 3; |
| 35 | 6ncv-A | 4.9 | 2.8 | 72 | 91 | 13 | MOLECULE: NACHT, LRR AND PYD DOMAINS-CONTAINING PROTEIN 6; |
| 36 | 2dbg-A | 4.8 | 3.3 | 82 | 103 | 6 | MOLECULE: MYELOID CELL NUCLEAR DIFFERENTIATION ANTIGEN; |
| 37 | 2ms7-A | 4.8 | 3 | 82 | 102 | 13 | MOLECULE: MITOCHONDRIAL ANTIVIRAL-SIGNALING PROTEIN; |
| 38 | 4zbw-A | 4.7 | 4.1 | 83 | 183 | 5 | MOLECULE: CASPASE-8; |
| 39 | 6eqt-A | 4.6 | 7.8 | 79 | 207 | 9 | MOLECULE: CENTROMERE PROTEIN N; |
| 40 | 2dbh-A | 4.6 | 3.6 | 84 | 102 | 8 | MOLECULE: TUMOR NECROSIS FACTOR RECEPTOR SUPERFAMILY |
| 41 | 5z8q-A | 4.6 | 6.2 | 67 | 101 | 4 | MOLECULE: HEAT SHOCK PROTEIN SSA1; |
| 42 | 6hj7-A | 4.5 | 4 | 83 | 102 | 10 | MOLECULE: DEATH DOMAIN-CONTAINING MEMBRANE PROTEIN NRADD; |
| 43 | 6ajf-A | 4.5 | 5.3 | 100 | 901 | 7 | MOLECULE: DRUG EXPORTERS OF THE RND SUPERFAMILY-LIKE PROTEI |
| 44 | 1e7p-C | 4.5 | 9.9 | 72 | 254 | 8 | MOLECULE: FUMARATE REDUCTASE FLAVOPROTEIN SUBUNIT; |
| 45 | 5xn7-B | 4.4 | 10.9 | 90 | 551 | 8 | MOLECULE: PUTATIVE RTX-TOXIN; |
| 46 | 2n00-A | 4.4 | 2.9 | 80 | 95 | 10 | MOLECULE: INTERFERON-INDUCIBLE PROTEIN AIM2; |
| 47 | 6irw-A | 4.4 | 12.9 | 78 | 495 | 8 | MOLECULE: PHOSPHORYLATED CTD-INTERACTING FACTOR 1; |
| 48 | 3crd-A | 4.4 | 3.8 | 89 | 100 | 12 | MOLECULE: RAIDD; |
| 49 | 6if2-A | 4.3 | 4.4 | 101 | 195 | 8 | MOLECULE: RAS-RELATED PROTEIN RAB-35; |
| 50 | 1irx-B | 4.3 | 8.9 | 73 | 508 | 10 | MOLECULE: LYSYL-TRNA SYNTHETASE; |
| 51 | 1sxj-D | 4.3 | 10.9 | 76 | 328 | 12 | MOLECULE: ACTIVATOR 1 95 KDA SUBUNIT; |
| 52 | 1gzs-B | 4.3 | 10.9 | 69 | 165 | 1 | MOLECULE: SOPE; |
| 53 | 6e6y-A | 4.3 | 20.1 | 79 | 268 | 10 | MOLECULE: DIECKMANN CYCLASE, NCMC; |
| 54 | 1xfi-A | 4.3 | 5.3 | 79 | 343 | 10 | MOLECULE: UNKNOWN PROTEIN; |
| 55 | 6ybw-z | 4.3 | 3.5 | 95 | 160 | 11 | MOLECULE: 40S RIBOSOMAL PROTEIN S4, X ISOFORM; |
| 56 | 1k30-A | 4.3 | 8.7 | 60 | 363 | 7 | MOLECULE: GLYCEROL-3-PHOSPHATE ACYLTRANSFERASE; |
| 57 | 4xom-A | 4.3 | 10.3 | 66 | 206 | 14 | MOLECULE: COENZYME F420:L-GLUTAMATE LIGASE; |
| 58 | 7lxc-A | 4.3 | 2.8 | 70 | 88 | 10 | MOLECULE: DED1CH; |
| 59 | 2bbz-A | 4.3 | 4.7 | 85 | 190 | 8 | MOLECULE: VIRAL CASP8 AND FADD-LIKE APOPTOSIS REGULATOR; |
| 60 | 6poo-A | 4.2 | 10.9 | 108 | 273 | 4 | MOLECULE: BIBA; |
| 61 | 5b4s-B | 4.2 | 10.8 | 56 | 310 | 9 | MOLECULE: CHITOSANASE; |
| 62 | 3p8c-B | 4.1 | 11.3 | 87 | 1086 | 5 | MOLECULE: CYTOPLASMIC FMR1-INTERACTING PROTEIN 1; |
| 63 | 5v8f-D | 4.1 | 12.5 | 87 | 456 | 5 | MOLECULE: DNA REPLICATION LICENSING FACTOR MCM2; |
| 64 | 6jal-A | 4.1 | 11 | 69 | 416 | 10 | MOLECULE: ABC TRANSPORTER, PERIPLASMIC SUBSTRATE-BINDING PR |
| 65 | 5yet-A | 4.1 | 10.6 | 62 | 397 | 15 | MOLECULE: UNCHARACTERIZED PROTEIN R354; |
| 66 | 2g8l-B | 4.1 | 3.7 | 70 | 286 | 13 | MOLECULE: 287AA LONG HYPOTHETICAL PROTEIN; |
| 67 | 6vvo-A | 4.1 | 13.5 | 62 | 448 | 6 | MOLECULE: REPLICATION FACTOR C SUBUNIT 1; |

| | | | | | | | |
|---|---|---|---|---|---|---|---|
| 68 | 6pie-A | 4.1 | 2.8 | 54 | 162 | 13 | MOLECULE: HEMERYTHRIN HHE CATION BINDING DOMAIN PROTEIN; |
| 69 | 7bvf-A | 4.1 | 10 | 92 | 1087 | 5 | MOLECULE: PROBABLE ARABINOSYLTRANSFERASE B; |
| 70 | 5izs-D | 4.1 | 4.1 | 47 | 82 | 9 | MOLECULE: DESIGNED PROTEIN 5L6HC3_1; |
| 71 | 2d96-A | 4 | 4.6 | 84 | 109 | 13 | MOLECULE: NUCLEAR FACTOR NF-KAPPA-B P100 SUBUNIT; |
| 72 | 4elj-A | 4 | 11 | 76 | 590 | 5 | MOLECULE: RETINOBLASTOMA-ASSOCIATED PROTEIN; |
| 73 | 6zg3-D | 4 | 3.3 | 53 | 260 | 4 | MOLECULE: ENERGY-COUPLING FACTOR TRANSPORTER ATP-BINDING PR |
| 74 | 2g7r-A | 4 | 3.3 | 66 | 86 | 11 | MOLECULE: MUCOSA-ASSOCIATED LYMPHOID TISSUE LYMPHOMA TRANSL |
| 75 | 6j72-A | 4 | 3.3 | 57 | 561 | 7 | MOLECULE: ISONIAZID INDUCIBLE GENE PROTEIN INIA; |
| 76 | 3u5z-B | 4 | 13.3 | 66 | 320 | 9 | MOLECULE: DNA POLYMERASE ACCESSORY PROTEIN 44; |
| 77 | 6ny2-Y | 4 | 13.1 | 123 | 915 | 5 | MOLECULE: DNA TARGET STRAND; |
| 78 | 6hsy-A | 4 | 5 | 58 | 190 | 12 | MOLECULE: TOLUENE TOLERANCE PROTEIN TTG2D; |
| 79 | 1wh4-A | 4 | 3.6 | 87 | 127 | 8 | MOLECULE: INTERLEUKIN-1 RECEPTOR-ASSOCIATED KINASE 4; |
| 80 | 1d2z-D | 3.9 | 8.3 | 94 | 153 | 10 | MOLECULE: DEATH DOMAIN OF PELLE; |
| 81 | 2hjm-A | 3.9 | 2.9 | 60 | 91 | 8 | MOLECULE: HYPOTHETICAL PROTEIN PF1176; |
| 82 | 5way-A | 3.9 | 5.3 | 64 | 482 | 5 | MOLECULE: TRANSCRIPTIONAL ACTIVATOR; |
| 83 | 5f3o-A | 3.9 | 3.5 | 59 | 194 | 5 | MOLECULE: PUTATIVE UNCHARACTERIZED PROTEIN; |
| 84 | 7c6g-A | 3.9 | 10.2 | 60 | 420 | 3 | MOLECULE: SUGAR ABC TRANSPORTER, PERIPLASMIC SUGAR-BINDING |
| 85 | 5wb2-A | 3.9 | 5.9 | 75 | 421 | 5 | MOLECULE: ENVELOPE PROTEIN US28, NANOBODY 7 FUSION PROTEIN; |
| 86 | 6jpq-A | 3.9 | 9.4 | 60 | 577 | 12 | MOLECULE: UNCHARACTERIZED AAA DOMAIN-CONTAINING PROTEIN C31 |
| 87 | 4mhq-A | 3.9 | 10.9 | 102 | 400 | 7 | MOLECULE: DNA PRIMASE SMALL SUBUNIT; |
| 88 | 6fze-A | 3.9 | 4.9 | 59 | 209 | 5 | MOLECULE: PUTATIVE SURFACE PROTEIN; |
| 89 | 3add-A | 3.9 | 12.5 | 57 | 251 | 14 | MOLECULE: L-SERYL-TRNA(SEC) KINASE; |
| 90 | 4kbq-C | 3.9 | 8.3 | 63 | 91 | 5 | MOLECULE: E3 UBIQUITIN-PROTEIN LIGASE CHIP; |
| 91 | 6uuw-A | 3.9 | 4.4 | 68 | 1034 | 4 | MOLECULE: ATP-CITRATE SYNTHASE; |
| 92 | 1jgt-B | 3.9 | 11.1 | 68 | 501 | 7 | MOLECULE: BETA-LACTAM SYNTHETASE; |
| 93 | 4aur-A | 3.9 | 5.4 | 68 | 564 | 15 | MOLECULE: LEOA; |
| 94 | 3dpn-A | 3.8 | 4.1 | 72 | 537 | 8 | MOLECULE: PROTEIN CT_858; |
| 95 | 5cwc-A | 3.8 | 7.5 | 61 | 196 | 11 | MOLECULE: DESIGNED HELICAL REPEAT PROTEIN; |
| 96 | 5vyz-A | 3.8 | 13 | 95 | 1144 | 6 | MOLECULE: PYRUVATE CARBOXYLASE; |
| 97 | 2gut-A | 3.8 | 2.7 | 48 | 77 | 4 | MOLECULE: ARC/MEDIATOR, POSITIVE COFACTOR 2 GLUTAMINE/Q- |
| 98 | 5che-A | 3.8 | 3.7 | 56 | 427 | 9 | MOLECULE: GLUTAMYL-TRNA REDUCTASE 1, CHLOROPLASTIC; |
| 99 | 2kn6-A | 3.8 | 3.4 | 67 | 195 | 10 | MOLECULE: APOPTOSIS-ASSOCIATED SPECK-LIKE PROTEIN CONTAININ |
| 100 | 3zjc-E | 3.8 | 12.4 | 64 | 286 | 17 | MOLECULE: GTPASE IMAP FAMILY MEMBER 7; |
| 101 | 4mk6-A | 3.8 | 11.1 | 67 | 188 | 6 | MOLECULE: PROBABLE DIHYDROXYACETONE KINASE REGULATOR DHSK_R |
| 102 | 7ltq-A | 3.8 | 5.8 | 62 | 256 | 6 | MOLECULE: HISTIDINE KINASE; |

| | | | | | | | |
|---|---|---|---|---|---|---|---|
| 103 | 6gvw-B | 3.8 | 6.9 | 55 | 262 | 9 | MOLECULE: BRCA1-A COMPLEX SUBUNIT ABRAXAS 1; |
| 104 | 7jzl-E | 3.8 | 3.1 | 49 | 55 | 12 | MOLECULE: SPIKE GLYCOPROTEIN; |
| 105 | 6jbq-F | 3.8 | 4.9 | 59 | 186 | 8 | MOLECULE: DNA-DIRECTED RNA POLYMERASE SUBUNIT ALPHA; |
| 106 | 6gc1-C | 3.8 | 11.7 | 78 | 551 | 10 | MOLECULE: NHL REPEAT-CONTAINING PROTEIN 2; |
| 107 | 7ew1-A | 3.8 | 5.4 | 68 | 269 | 6 | MOLECULE: GUANINE NUCLEOTIDE-BINDING PROTEIN G(I)/G(S)/G(T) |
| 108 | 3kdw-A | 3.8 | 6.2 | 75 | 206 | 7 | MOLECULE: PUTATIVE SUGAR BINDING PROTEIN; |
| 109 | 7kfu-C | 3.8 | 10.4 | 94 | 909 | 6 | MOLECULE: CAS2; |
| 110 | 1nxh-A | 3.8 | 10.8 | 73 | 124 | 14 | MOLECULE: MTH396 PROTEIN; |
| 111 | 3h0d-A | 3.8 | 5.2 | 72 | 155 | 4 | MOLECULE: CTSR; |
| 112 | 3l32-B | 3.8 | 3.2 | 44 | 45 | 7 | MOLECULE: PHOSPHOPROTEIN; |
| 113 | 3cym-A | 3.8 | 7.5 | 78 | 415 | 5 | MOLECULE: UNCHARACTERIZED PROTEIN BAD_0989; |
| 114 | 7s9y-A | 3.8 | 9.4 | 86 | 901 | 3 | MOLECULE: CYTOCHROME C BIOGENESIS PROTEIN; |
| 115 | 1l1l-A | 3.7 | 9.7 | 85 | 717 | 7 | MOLECULE: RIBONUCLEOSIDE TRIPHOSPHATE REDUCTASE; |
| 116 | 2efl-A | 3.7 | 6.9 | 76 | 281 | 3 | MOLECULE: FORMIN-BINDING PROTEIN 1; |
| 117 | 4ayb-Q | 3.7 | 4.2 | 45 | 50 | 11 | MOLECULE: DNA-DIRECTED RNA POLYMERASE; |
| 118 | 2o7g-A | 3.7 | 3.3 | 47 | 88 | 2 | MOLECULE: PROBABLE RNA POLYMERASE SIGMA-C FACTOR; |
| 119 | 6ocv-A | 3.7 | 4.3 | 76 | 190 | 5 | MOLECULE: HEME NO BINDING DOMAIN PROTEIN; |
| 120 | 7cun-E | 3.7 | 12.5 | 67 | 796 | 9 | MOLECULE: INTEGRATOR COMPLEX SUBUNIT 1; |
| 121 | 4akx-B | 3.7 | 10.8 | 68 | 534 | 6 | MOLECULE: SPCU; |
| 122 | 6ibl-A | 3.7 | 4.2 | 65 | 397 | 11 | MOLECULE: THIOREDOXIN 1,BETA-1 ADRENERGIC RECEPTOR; |
| 123 | 6ptm-A | 3.7 | 4.6 | 66 | 726 | 8 | MOLECULE: UNCHARACTERIZED PROTEIN; |
| 124 | 6rmg-A | 3.7 | 10.6 | 108 | 1026 | 10 | MOLECULE: PROTEIN PATCHED HOMOLOG 1,GFP-LIKE FLUORESCENT |
| 125 | 2d1l-A | 3.7 | 8.5 | 70 | 249 | 4 | MOLECULE: METASTASIS SUPPRESSOR PROTEIN 1; |
| 126 | 2y7h-B | 3.7 | 2.4 | 42 | 530 | 2 | MOLECULE: TYPE-1 RESTRICTION ENZYME ECOKI SPECIFICITY PROTE |
| 127 | 6ebu-A | 3.7 | 4.7 | 77 | 165 | 9 | MOLECULE: LPXE; |
| 128 | 7lt5-B | 3.7 | 11.5 | 65 | 577 | 12 | MOLECULE: SITE-SPECIFIC DNA-METHYLTRANSFERASE (ADENINE-SPEC |
| 129 | 6tnt-A | 3.7 | 17.2 | 93 | 1583 | 4 | MOLECULE: ANAPHASE-PROMOTING COMPLEX SUBUNIT 1; |
| 130 | 6mtz-B | 3.7 | 7.2 | 70 | 347 | 9 | MOLECULE: HEPC.19480.A.B1; |
| 131 | 4wl5-A | 3.7 | 5.1 | 94 | 594 | 5 | MOLECULE: HETERODIMERIC RESTRICTION ENDONUCLEASE R.BSPD6I L |
| 132 | 6qgi-A | 3.7 | 2 | 40 | 498 | 10 | MOLECULE: VP5; |
| 133 | 6cng-A | 3.7 | 4.3 | 63 | 281 | 6 | MOLECULE: FATTY ACID KINASE (FAK) B3 PROTEIN; |
| 134 | 1r5i-D | 3.7 | 4.5 | 67 | 214 | 4 | MOLECULE: HLA CLASS II HISTOCOMPATIBILITY ANTIGEN, DR |
| 135 | 6plm-B | 3.7 | 13.5 | 91 | 752 | 8 | MOLECULE: SIDJ PROTEIN; |
| 136 | 3bes-R | 3.6 | 6.2 | 45 | 250 | 13 | MOLECULE: INTERFERON GAMMA; |
| 137 | 5mps-R | 3.6 | 4.5 | 46 | 108 | 15 | MOLECULE: YEAST UBC4 GENE FOR UBIQUITIN-CONJUGATING ENZYME; |

| | | | | | | | |
|---|---|---|---|---|---|---|---|
| 138 | 6ep3-B | 3.6 | 9.6 | 48 | 218 | 4 | MOLECULE: LMO0651 PROTEIN; |
| 139 | 4ywz-A | 3.6 | 4.5 | 70 | 147 | 13 | MOLECULE: SENSOR PROTEIN KINASE WALK; |
| 140 | 6ssk-B | 3.6 | 8.4 | 91 | 223 | 8 | MOLECULE: ENDOGENOUS RETROVIRUS GROUP K MEMBER 24 GAG POLYP |
| 141 | 6bog-A | 3.6 | 5.5 | 83 | 967 | 10 | MOLECULE: RNA POLYMERASE-ASSOCIATED PROTEIN RAPA; |
| 142 | 2ygu-F | 3.6 | 3.8 | 64 | 118 | 6 | MOLECULE: VENOM ALLERGEN 2; |
| 143 | 7mca-F | 3.6 | 6.3 | 71 | 164 | 11 | MOLECULE: ORIGIN RECOGNITION COMPLEX SUBUNIT 1; |
| 144 | 6f1t-f | 3.6 | 17.8 | 95 | 929 | 7 | MOLECULE: ARP1 ACTIN RELATED PROTEIN 1 HOMOLOG A; |
| 145 | 2hh6-A | 3.6 | 8.2 | 57 | 112 | 18 | MOLECULE: BH3980 PROTEIN; |
| 146 | 1ywm-A | 3.6 | 11.5 | 82 | 180 | 10 | MOLECULE: C PROTEIN ALPHA-ANTIGEN; |
| 147 | 5yx4-A | 3.6 | 9 | 59 | 232 | 3 | MOLECULE: CHALCONE-FLAVONONE ISOMERASE FAMILY PROTEIN; |
| 148 | 7eeb-C | 3.6 | 12.7 | 69 | 278 | 10 | MOLECULE: ENHANCED GREEN FLUORESCENT PROTEIN,CATION CHANNEL |
| 149 | 1pn5-A | 3.6 | 3.1 | 74 | 93 | 11 | MOLECULE: NACHT-, LRR- AND PYD-CONTAINING PROTEIN 2; |
| 150 | 6f1y-f | 3.6 | 5.9 | 83 | 280 | 11 | MOLECULE: CYTOPLASMIC DYNEIN 1 HEAVY CHAIN 1,DYNEIN HEAVY C |
| 151 | 5w7g-A | 3.6 | 5.6 | 56 | 131 | 7 | MOLECULE: ORF140; |
| 152 | 3jcm-I | 3.5 | 16.6 | 85 | 416 | 7 | MOLECULE: PRE-MRNA-SPLICING FACTOR 8; |
| 153 | 5nx9-D | 3.5 | 11.3 | 83 | 477 | 8 | MOLECULE: ADENYLOSUCCINATE LYASE; |
| 154 | 7jz2-A | 3.5 | 11.7 | 74 | 481 | 4 | MOLECULE: SUCCINATE DEHYDROGENASE FLAVOPROTEIN SUBUNIT; |
| 155 | 6dg6-A | 3.5 | 5 | 55 | 100 | 5 | MOLECULE: NEOLEUKIN-2/15; |
| 156 | 3hmj-A | 3.5 | 9.4 | 65 | 1750 | 6 | MOLECULE: FATTY ACID SYNTHASE SUBUNIT ALPHA; |
| 157 | 2ogi-B | 3.5 | 8.1 | 67 | 194 | 7 | MOLECULE: HYPOTHETICAL PROTEIN SAG1661; |
| 158 | 5aqd-M | 3.5 | 3.8 | 55 | 184 | 2 | MOLECULE: PHYCOERYTHRIN ALPHA SUBUNIT; |
| 159 | 1sb0-A | 3.5 | 2.7 | 47 | 87 | 4 | MOLECULE: PROTEIN CBP; |
| 160 | 7plh-A | 3.5 | 5 | 48 | 260 | 10 | MOLECULE: SHTNSC; |
| 161 | 3u5z-A | 3.5 | 18.5 | 84 | 186 | 6 | MOLECULE: DNA POLYMERASE ACCESSORY PROTEIN 44; |
| 162 | 5he9-E | 3.5 | 2.5 | 42 | 56 | 7 | MOLECULE: HELICASE LOADER; |
| 163 | 4gzr-C | 3.5 | 2.8 | 40 | 61 | 13 | MOLECULE: ESAT-6-LIKE PROTEIN 6; |
| 164 | 6vxm-A | 3.5 | 9.3 | 52 | 277 | 15 | MOLECULE: MECHANOSENSITIVE ION CHANNEL PROTEIN 1, MITOCHOND |
| 165 | 6w6w-A | 3.5 | 14 | 79 | 905 | 13 | MOLECULE: CST COMPLEX SUBUNIT CTC1; |
| 166 | 2dd4-H | 3.5 | 5.9 | 67 | 156 | 6 | MOLECULE: THIOCYANATE HYDROLASE ALPHA SUBUNIT; |
| 167 | 3jc5-A | 3.5 | 10.6 | 64 | 208 | 6 | MOLECULE: DNA REPLICATION LICENSING FACTOR MCM2; |
| 168 | 7jve-A | 3.5 | 18.2 | 81 | 253 | 5 | MOLECULE: DSBA FAMILY PROTEIN; |
| 169 | 6lu0-A | 3.5 | 17.7 | 92 | 937 | 12 | MOLECULE: CAS12I2; |
| 170 | 6myo-D | 3.5 | 10.6 | 54 | 102 | 4 | MOLECULE: SUCCINATE DEHYDROGENASE [UBIQUINONE] FLAVOPROTEIN |
| 171 | 3if8-B | 3.5 | 11.5 | 60 | 237 | 8 | MOLECULE: PROTEIN ZWILCH HOMOLOG; |
| 172 | 3w1h-A | 3.5 | 9.7 | 64 | 452 | 6 | MOLECULE: L-SERYL-TRNA(SEC) SELENIUM TRANSFERASE; |

| | | | | | | | |
|---|---|---|---|---|---|---|---|
| 173 | 6nsj-A | 3.5 | 6.5 | 64 | 182 | 8 | MOLECULE: ACID-ACTIVATED UREA CHANNEL; |
| 174 | 6vvo-E | 3.5 | 9.4 | 81 | 346 | 6 | MOLECULE: REPLICATION FACTOR C SUBUNIT 1; |
| 175 | 7dpa-A | 3.5 | 11.6 | 90 | 1642 | 12 | MOLECULE: DEDICATOR OF CYTOKINESIS PROTEIN 5; |
| 176 | 5ziq-A | 3.5 | 2.7 | 63 | 155 | 5 | MOLECULE: GLOBIN PROTEIN; |
| 177 | 5ziy-A | 3.5 | 5.2 | 62 | 195 | 5 | MOLECULE: FLAGELLAR HOOK-ASSOCIATED PROTEIN 3; |
| 178 | 7m2w-F | 3.5 | 11.1 | 92 | 674 | 8 | MOLECULE: TUBULIN GAMMA CHAIN; |
| 179 | 3caz-B | 3.5 | 8.9 | 73 | 210 | 7 | MOLECULE: BAR PROTEIN; |
| 180 | 5a9e-A | 3.5 | 10.5 | 102 | 254 | 8 | MOLECULE: DELTAMBD GAG PROTEIN; |
| 181 | 6d03-E | 3.5 | 4.4 | 75 | 466 | 4 | MOLECULE: TRANSFERRIN RECEPTOR PROTEIN 1; |
| 182 | 2px0-A | 3.4 | 6.5 | 68 | 258 | 4 | MOLECULE: FLAGELLAR BIOSYNTHESIS PROTEIN FLHF; |
| 183 | 6efw-A | 3.4 | 5.3 | 55 | 345 | 5 | MOLECULE: ATP-DEPENDENT (S)-NAD(P)H-HYDRATE DEHYDRATASE; |
| 184 | 3l9w-B | 3.4 | 13.1 | 91 | 357 | 9 | MOLECULE: GLUTATHIONE-REGULATED POTASSIUM-EFFLUX SYSTEM PRO |
| 185 | 5n2j-B | 3.4 | 23.1 | 76 | 1381 | 7 | MOLECULE: UDP-GLUCOSE-GLYCOPROTEIN GLUCOSYLTRANSFERASE- |
| 186 | 1n3k-A | 3.4 | 4.4 | 94 | 130 | 12 | MOLECULE: ASTROCYTIC PHOSPHOPROTEIN PEA-15; |
| 187 | 6izw-B | 3.4 | 8.6 | 49 | 145 | 8 | MOLECULE: MUTUAL GLIDING-MOTILITY PROTEIN MGLA; |
| 188 | 4lnb-A | 3.4 | 6.3 | 60 | 339 | 5 | MOLECULE: CAAX FARNESYLTRANSFERASE ALPHA SUBUNIT RAM2; |
| 189 | 2nup-C | 3.4 | 5.4 | 46 | 137 | 11 | MOLECULE: PROTEIN TRANSPORT PROTEIN SEC23A; |
| 190 | 2hxo-A | 3.4 | 12.6 | 67 | 207 | 7 | MOLECULE: PUTATIVE TETR-FAMILY TRANSCRIPTIONAL REGULATOR; |
| 191 | 6yxp-B | 3.4 | 3.6 | 65 | 272 | 9 | MOLECULE: SWI/SNF COMPLEX SUBUNIT SMARCC1; |
| 192 | 6n1l-A | 3.4 | 3.3 | 53 | 141 | 8 | MOLECULE: FIBRONECTIN-BINDING PROTEIN BBK32; |
| 193 | 7dfe-A | 3.4 | 7.1 | 97 | 142 | 5 | MOLECULE: B6 PROTEIN; |
| 194 | 7obq-u | 3.4 | 5.4 | 64 | 441 | 5 | MOLECULE: SRP RNA; |
| 195 | 3p8c-F | 3.4 | 6.8 | 60 | 156 | 7 | MOLECULE: CYTOPLASMIC FMR1-INTERACTING PROTEIN 1; |
| 196 | 4our-B | 3.4 | 9.9 | 84 | 445 | 4 | MOLECULE: PHYTOCHROME B; |
| 197 | 4m0m-A | 3.4 | 4.3 | 75 | 721 | 11 | MOLECULE: PUTATIVE UNCHARACTERIZED PROTEIN; |
| 198 | 7lxt-J | 3.4 | 7.2 | 64 | 215 | 8 | MOLECULE: 20S PROTEASOME ALPHA-1 SUBUNIT; |
| 199 | 3oao-A | 3.4 | 8.2 | 58 | 140 | 10 | MOLECULE: UNCHARACTERIZED PROTEIN FROM DUF2059 FAMILY; |
| 200 | 5wp4-A | 3.4 | 10.7 | 87 | 487 | 9 | MOLECULE: PHOSPHOETHANOLAMINE N-METHYLTRANSFERASE 1; |
| 201 | 6l1q-A | 3.4 | 10 | 66 | 267 | 6 | MOLECULE: CBBQ PROTEIN; |
| 202 | 6ueb-A | 3.4 | 13.6 | 78 | 2099 | 6 | MOLECULE: LARGE STRUCTURAL PROTEIN; |
| 203 | 5dvi-B | 3.4 | 10 | 49 | 396 | 4 | MOLECULE: BINDING PROTEIN COMPONENT OF ABC SUGAR TRANSPORTE |
| 204 | 2c5q-E | 3.4 | 3.2 | 46 | 233 | 4 | MOLECULE: RRAA-LIKE PROTEIN YER010C; |
| 205 | 6kp3-B | 3.4 | 3.3 | 64 | 112 | 6 | MOLECULE: PROGRAMMED CELL DEATH 6-INTERACTING PROTEIN; |
| 206 | 7s0r-A | 3.4 | 4.4 | 56 | 85 | 9 | MOLECULE: C PROTEIN BETA ANTIGEN; |
| 207 | 4uzz-B | 3.4 | 3.5 | 55 | 65 | 5 | MOLECULE: INTRAFLAGELLAR TRANSPORT COMPLEX B PROTEIN 46 |

| | | | | | | | |
|---|---|---|---|---|---|---|---|
| 208 | 3fjv-A | 3.4 | 7.5 | 67 | 193 | 10 | MOLECULE: UNCHARACTERIZED NOVEL PROTEIN; |
| 209 | 1lq7-A | 3.4 | 3.8 | 52 | 67 | 6 | MOLECULE: ALPHA3W; |
| 210 | 7dvq-7 | 3.4 | 4.3 | 42 | 81 | 5 | MOLECULE: PRE-MRNA-PROCESSING-SPLICING FACTOR 8; |
| 211 | 6nes-A | 3.4 | 7 | 57 | 431 | 0 | MOLECULE: FAD-DEPENDENT MONOOXYGENASE TROPB; |
| 212 | 5w7g-B | 3.4 | 4.8 | 56 | 126 | 7 | MOLECULE: ORF140; |
| 213 | 6ks0-A | 3.4 | 13.7 | 85 | 286 | 8 | MOLECULE: ADIPONECTIN RECEPTOR PROTEIN 1; |
| 214 | 6en3-A | 3.4 | 12.5 | 82 | 934 | 7 | MOLECULE: ENDO-BETA-N-ACETYLGLUCOSAMINIDASE F2,MULTIFUNCTIO |
| 215 | 6zyd-A | 3.4 | 9.5 | 67 | 323 | 10 | MOLECULE: LOW CONDUCTANCE MECHANOSENSITIVE CHANNEL YNAI,LOW |
| 216 | 4bt9-B | 3.4 | 5.4 | 52 | 238 | 8 | MOLECULE: PROLYL 4-HYDROXYLASE SUBUNIT ALPHA-1; |
| 217 | 2nn4-A | 3.4 | 7.5 | 47 | 62 | 6 | MOLECULE: HYPOTHETICAL PROTEIN YQGQ; |
| 218 | 5eqz-A | 3.4 | 7 | 72 | 138 | 10 | MOLECULE: REV PROTEIN; |
| 219 | 4heo-B | 3.4 | 3.3 | 48 | 60 | 6 | MOLECULE: PHOSPHOPROTEIN; |
| 220 | 3j5s-D | 3.4 | 10.3 | 64 | 554 | 9 | MOLECULE: 16S RIBOSOMAL RNA; |
| 221 | 6lo8-D | 3.4 | 9.7 | 53 | 119 | 8 | MOLECULE: MITOCHONDRIAL IMPORT INNER MEMBRANE TRANSLOCASE S |
| 222 | 3bbz-A | 3.4 | 2.7 | 45 | 48 | 2 | MOLECULE: P PROTEIN; |
| 223 | 5gza-A | 3.4 | 11.6 | 75 | 290 | 9 | MOLECULE: PROTEIN O-MANNOSE KINASE; |
| 224 | 5oxf-C | 3.4 | 3.6 | 61 | 601 | 10 | MOLECULE: GTP-BINDING PROTEIN; |
| 225 | 6vbk-B | 3.4 | 2.8 | 46 | 210 | 7 | MOLECULE: LON211; |
| 226 | 2do9-A | 3.4 | 5.6 | 80 | 115 | 9 | MOLECULE: NACHT-, LRR- AND PYD-CONTAINING PROTEIN 10; |
| 227 | 3ezh-A | 3.3 | 5.9 | 58 | 114 | 10 | MOLECULE: NITRATE/NITRITE SENSOR PROTEIN NARX; |
| 228 | 6jiv-A | 3.3 | 10.9 | 83 | 560 | 8 | MOLECULE: SSPE PROTEIN; |
| 229 | 4m70-B | 3.3 | 3.3 | 51 | 91 | 14 | MOLECULE: RX PROTEIN; |
| 230 | 5ulm-B | 3.3 | 10.8 | 79 | 376 | 10 | MOLECULE: MITOGEN-ACTIVATED PROTEIN KINASE KINASE KINASE 5; |
| 231 | 6gw5-A | 3.3 | 2.9 | 54 | 393 | 6 | MOLECULE: PUTATIVE OUTER MEMBRANE PROTEIN; |
| 232 | 3dkq-A | 3.3 | 11.9 | 50 | 230 | 4 | MOLECULE: PKHD-TYPE HYDROXYLASE SBAL_3634; |
| 233 | 1sg9-A | 3.3 | 10.1 | 64 | 276 | 5 | MOLECULE: HEMK PROTEIN; |
| 234 | 4kis-A | 3.3 | 7.2 | 60 | 315 | 7 | MOLECULE: PUTATIVE INTEGRASE [BACTERIOPHAGE A118]; |
| 235 | 6tv6-A | 3.3 | 11.9 | 80 | 362 | 10 | MOLECULE: PROTEIN-ARGININE KINASE; |
| 236 | 2fsf-B | 3.3 | 4.8 | 68 | 723 | 4 | MOLECULE: PREPROTEIN TRANSLOCASE SECA SUBUNIT; |
| 237 | 6qfo-A | 3.3 | 3.8 | 83 | 1348 | 6 | MOLECULE: PEGA DOMAIN-CONTAINING PROTEIN,PEGA DOMAIN-CONTAI |
| 238 | 6rlb-A | 3.3 | 11.2 | 95 | 969 | 8 | MOLECULE: O6-ALKYLGUANINE-DNA ALKYLTRANSFERASE MUTANT,DYNC2 |
| 239 | 3viu-A | 3.3 | 17.4 | 78 | 703 | 5 | MOLECULE: PHOSPHORIBOSYLFORMYLGLYCINAMIDINE SYNTHASE 2; |
| 240 | 3ugm-A | 3.3 | 8.5 | 72 | 854 | 6 | MOLECULE: TAL EFFECTOR AVRBS3/PTHA; |
| 241 | 4dlq-A | 3.3 | 7.7 | 83 | 353 | 5 | MOLECULE: LATROPHILIN-1; |
| 242 | 1szi-A | 3.3 | 5.1 | 53 | 194 | 4 | MOLECULE: MANNOSE-6-PHOSPHATE RECEPTOR BINDING PROTEIN 1; |

| | | | | | | | |
|---|---|---|---|---|---|---|---|
| 243 | 6q7p-A | 3.3 | 22.6 | 66 | 230 | 6 | MOLECULE: OE1.2; |
| 244 | 2e5t-A | 3.3 | 4.2 | 39 | 46 | 15 | MOLECULE: ATP SYNTHASE EPSILON CHAIN; |
| 245 | 2jxn-A | 3.3 | 9.8 | 63 | 116 | 10 | MOLECULE: UNCHARACTERIZED PROTEIN YMR074C; |
| 246 | 4ddg-A | 3.3 | 6.9 | 62 | 399 | 13 | MOLECULE: UBIQUITIN-CONJUGATING ENZYME E2 D2, UBIQUITIN THI |
| 247 | 4ae4-B | 3.3 | 2.8 | 42 | 115 | 7 | MOLECULE: UBIQUITIN-ASSOCIATED PROTEIN 1; |
| 248 | 2xa6-A | 3.3 | 4.3 | 37 | 37 | 11 | MOLECULE: KH DOMAIN-CONTAINING\,RNA-BINDING\,SIGNAL |
| 249 | 6erp-A | 3.3 | 4.5 | 84 | 1011 | 6 | MOLECULE: TRANSCRIPTION FACTOR A, MITOCHONDRIAL; |
| 250 | 2m6u-A | 3.3 | 6.6 | 51 | 82 | 8 | MOLECULE: CHOLINE BINDING PROTEIN A; |
| 251 | 4qsz-A | 3.3 | 11.6 | 85 | 686 | 11 | MOLECULE: MALTOSE-BINDING PERIPLASMIC PROTEIN, JMJC DOMAIN- |
| 252 | 6s7o-E | 3.3 | 4.1 | 47 | 560 | 6 | MOLECULE: DOLICHYL-DIPHOSPHOOLIGOSACCHARIDE--PROTEIN |
| 253 | 3dye-A | 3.3 | 5.3 | 88 | 302 | 10 | MOLECULE: D7 PROTEIN; |
| 254 | 6es4-A | 3.3 | 2.3 | 42 | 207 | 7 | MOLECULE: SYNCRIP, ISOFORM K; |
| 255 | 1ej5-A | 3.3 | 4.2 | 63 | 107 | 2 | MOLECULE: WISKOTT-ALDRICH SYNDROME PROTEIN; |
| 256 | 7d7c-F | 3.3 | 6.7 | 79 | 137 | 9 | MOLECULE: DNA-DIRECTED RNA POLYMERASE SUBUNIT ALPHA; |
| 257 | 5fqf-A | 3.3 | 7.5 | 72 | 583 | 11 | MOLECULE: BETA-N-ACETYLGALACTOSAMINIDASE; |
| 258 | 6djy-C | 3.3 | 4.4 | 71 | 1041 | 7 | MOLECULE: CLAMP PROTEIN; |
| 259 | 7lt2-A | 3.3 | 12.6 | 79 | 387 | 6 | MOLECULE: MAB-21 DOMAIN-CONTAINING PROTEIN; |
| 260 | 6m9a-C | 3.3 | 4.9 | 54 | 157 | 6 | MOLECULE: SIGNALING PROTEIN; |
| 261 | 4wvm-B | 3.3 | 11.4 | 88 | 616 | 10 | MOLECULE: STONUSTOXIN SUBUNIT ALPHA; |
| 262 | 5k47-A | 3.3 | 10.8 | 52 | 484 | 6 | MOLECULE: POLYCYSTIN-2; |
| 263 | 4wbd-A | 3.3 | 11.2 | 71 | 540 | 3 | MOLECULE: BSHC; |
| 264 | 4a4z-A | 3.3 | 14.3 | 71 | 874 | 8 | MOLECULE: ANTIVIRAL HELICASE SKI2; |
| 265 | 6rjw-A | 3.3 | 5.2 | 83 | 161 | 12 | MOLECULE: LYSM DOMAIN PROTEIN; |
| 266 | 5ckw-B | 3.3 | 4.8 | 68 | 400 | 13 | MOLECULE: LEGK4; |
| 267 | 1nek-C | 3.3 | 4.3 | 59 | 129 | 7 | MOLECULE: SUCCINATE DEHYDROGENASE FLAVOPROTEIN SUBUNIT; |
| 268 | 4ams-A | 3.3 | 10.9 | 88 | 367 | 13 | MOLECULE: MG662; |
| 269 | 3bni-A | 3.2 | 12.4 | 74 | 174 | 14 | MOLECULE: PUTATIVE TETR-FAMILY TRANSCRIPTIONAL REGULATOR; |
| 270 | 4afi-A | 3.2 | 2.7 | 46 | 155 | 7 | MOLECULE: AP-3 COMPLEX SUBUNIT DELTA-1, VESICLE-ASSOCIATED |
| 271 | 5mpd-Q | 3.2 | 12.3 | 77 | 434 | 5 | MOLECULE: 26S PROTEASOME REGULATORY SUBUNIT RPN10; |
| 272 | 4abn-B | 3.2 | 6.8 | 55 | 426 | 9 | MOLECULE: TETRATRICOPEPTIDE REPEAT PROTEIN 5; |
| 273 | 2qq8-A | 3.2 | 8.5 | 74 | 288 | 5 | MOLECULE: TBC1 DOMAIN FAMILY MEMBER 14; |
| 274 | 6e11-E | 3.2 | 4 | 72 | 210 | 3 | MOLECULE: UNKNOWN (CLAW); |
| 275 | 7obq-y | 3.2 | 15.6 | 71 | 454 | 8 | MOLECULE: SRP RNA; |
| 276 | 6ekk-B | 3.2 | 11.4 | 74 | 381 | 8 | MOLECULE: DENN DOMAIN-CONTAINING PROTEIN 1A; |
| 277 | 2w4m-A | 3.2 | 23.2 | 67 | 250 | 1 | MOLECULE: N-ACYLNEURAMINATE-9-PHOSPHATASE; |

| | | | | | | | |
|---|---|---|---|---|---|---|---|
| 278 | 3htk-C | 3.2 | 13.1 | 52 | 254 | 8 | MOLECULE: STRUCTURAL MAINTENANCE OF CHROMOSOMES PROTEIN 5; |
| 279 | 5exr-B | 3.2 | 4.4 | 57 | 434 | 18 | MOLECULE: DNA PRIMASE SMALL SUBUNIT; |
| 280 | 6swg-B | 3.2 | 4.2 | 45 | 74 | 13 | MOLECULE: PERIPHILIN-1; |
| 281 | 6exn-a | 3.2 | 5.8 | 54 | 171 | 9 | MOLECULE: U2 SNRNA; |
| 282 | 6e5o-Y | 3.2 | 22.2 | 96 | 898 | 8 | MOLECULE: CASX; |
| 283 | 4oe8-C | 3.2 | 3.8 | 57 | 87 | 11 | MOLECULE: INTERLEUKIN-12 SUBUNIT BETA; |
| 284 | 6ogd-E | 3.2 | 12.5 | 78 | 1078 | 14 | MOLECULE: TOXIN SUBUNIT YENA1; |
| 285 | 2pv4-A | 3.2 | 5.2 | 49 | 145 | 12 | MOLECULE: UNCHARACTERIZED PROTEIN; |
| 286 | 5wx8-A | 3.2 | 3.8 | 61 | 165 | 10 | MOLECULE: IMMEDIATE-EARLY PROTEIN 2; |
| 287 | 6s3k-A | 3.2 | 5.5 | 71 | 573 | 10 | MOLECULE: APC FAMILY PERMEASE; |
| 288 | 4zil-A | 3.2 | 11.8 | 72 | 210 | 10 | MOLECULE: DSBA OXIDOREDUCTASE; |
| 289 | 6tdv-D | 3.2 | 3.7 | 79 | 186 | 5 | MOLECULE: ATPTB1; |
| 290 | 6ikn-A | 3.2 | 10.4 | 87 | 305 | 11 | MOLECULE: GROWTH ARREST-SPECIFIC PROTEIN 7; |
| 291 | 6nr8-6 | 3.2 | 4.6 | 46 | 102 | 11 | MOLECULE: PREFOLDIN SUBUNIT 1; |
| 292 | 6wpg-A | 3.2 | 3.3 | 73 | 119 | 4 | MOLECULE: REGULATORY PROTEIN NPR4; |
| 293 | 2xvt-C | 3.2 | 3.2 | 46 | 79 | 11 | MOLECULE: RECEPTOR ACTIVITY-MODIFYING PROTEIN 2; |
| 294 | 6o0i-A | 3.2 | 4.6 | 37 | 98 | 5 | MOLECULE: DESIGN CONSTRUCT XAA; |
| 295 | 6dan-B | 3.2 | 13.9 | 76 | 328 | 12 | MOLECULE: PHDJ; |
| 296 | 6vhf-A | 3.2 | 4.4 | 41 | 140 | 10 | MOLECULE: PHD-TYPE DOMAIN-CONTAINING PROTEIN; |
| 297 | 6eu1-O | 3.2 | 3.9 | 47 | 551 | 9 | MOLECULE: DNA-DIRECTED RNA POLYMERASE III SUBUNIT RPC1; |
| 298 | 4wvz-D | 3.2 | 4.6 | 59 | 203 | 7 | MOLECULE: THIOL DIOXYGENASE; |
| 299 | 7jpp-E | 3.2 | 12.9 | 71 | 406 | 6 | MOLECULE: ORIGIN RECOGNITION COMPLEX SUBUNIT 1; |
| 300 | 4etx-A | 3.2 | 11 | 68 | 300 | 10 | MOLECULE: PELD; |
| 301 | 6wgm-A | 3.2 | 8 | 58 | 305 | 3 | MOLECULE: TRAP-TYPE C4-DICARBOXYLATE TRANSPORT SYSTEM, PERI |
| 302 | 6uv7-B | 3.2 | 11.4 | 70 | 154 | 10 | MOLECULE: ALR1298 PROTEIN; |
| 303 | 1jr3-D | 3.2 | 10.5 | 75 | 338 | 8 | MOLECULE: DNA POLYMERASE III SUBUNIT GAMMA; |
| 304 | 6cfw-B | 3.2 | 4.5 | 47 | 82 | 4 | MOLECULE: MONOVALENT CATION/H+ ANTIPORTER SUBUNIT D; |
| 305 | 2dwk-A | 3.2 | 7.1 | 99 | 162 | 9 | MOLECULE: PROTEIN RUFY3; |
| 306 | 2xv9-A | 3.2 | 3 | 69 | 134 | 13 | MOLECULE: ABA-1A1 REPEAT UNIT; |
| 307 | 1tiq-A | 3.2 | 9 | 54 | 173 | 6 | MOLECULE: PROTEASE SYNTHASE AND SPORULATION NEGATIVE REGULA |
| 308 | 6l7o-G | 3.2 | 8.2 | 48 | 192 | 10 | MOLECULE: NAD(P)H-QUINONE OXIDOREDUCTASE SUBUNIT 1; |
| 309 | 2b3t-A | 3.2 | 10.8 | 67 | 277 | 9 | MOLECULE: PROTEIN METHYLTRANSFERASE HEMK; |
| 310 | 7k3h-A | 3.2 | 4.9 | 57 | 104 | 11 | MOLECULE: NETWORK HALLUCINATED PROTEIN 0217; |
| 311 | 6r6b-F | 3.2 | 9.1 | 59 | 245 | 10 | MOLECULE: SURFACE PRESENTATION OF ANTIGENS PROTEIN SPAP; |
| 312 | 5l4k-S | 3.2 | 14.8 | 78 | 491 | 8 | MOLECULE: 26S PROTEASOME NON-ATPASE REGULATORY SUBUNIT 4; |

| | | | | | | | |
|---|---|---|---|---|---|---|---|
| 313 | 6cfw-A | 3.2 | 2.8 | 43 | 165 | 0 | MOLECULE: MONOVALENT CATION/H+ ANTIPORTER SUBUNIT D; |
| 314 | 6lo8-C | 3.2 | 3.9 | 50 | 108 | 12 | MOLECULE: MITOCHONDRIAL IMPORT INNER MEMBRANE TRANSLOCASE S |
| 315 | 6umm-A | 3.2 | 6.3 | 61 | 285 | 8 | MOLECULE: ESX-3 SECRETION SYSTEM PROTEIN ECCE3; |
| 316 | 6jt0-B | 3.2 | 12.1 | 88 | 576 | 7 | MOLECULE: GUANYLATE CYCLASE SOLUBLE SUBUNIT ALPHA-1; |
| 317 | 6h4j-A | 3.2 | 10.9 | 55 | 488 | 9 | MOLECULE: UBIQUITIN CARBOXYL-TERMINAL HYDROLASE 25; |
| 318 | 5sy1-A | 3.2 | 10.5 | 82 | 582 | 9 | MOLECULE: CALMODULIN; |
| 319 | 3vp5-A | 3.2 | 12.8 | 61 | 185 | 11 | MOLECULE: TRANSCRIPTIONAL REGULATOR; |
| 320 | 6sty-A | 3.2 | 9.7 | 71 | 198 | 11 | MOLECULE: OLIGORIBONUCLEASE, MITOCHONDRIAL; |
| 321 | 3ufe-A | 3.2 | 4 | 49 | 109 | 14 | MOLECULE: TRANSCRIPTIONAL ANTITERMINATOR (BGLG FAMILY); |
| 322 | 3d5l-A | 3.2 | 3.3 | 47 | 203 | 9 | MOLECULE: REGULATORY PROTEIN RECX; |
| 323 | 7t4x-B | 3.2 | 5.1 | 58 | 450 | 9 | MOLECULE: POTASSIUM CHANNEL AKT1; |
| 324 | 1kw2-A | 3.2 | 7 | 76 | 455 | 9 | MOLECULE: VITAMIN D-BINDING PROTEIN; |
| 325 | 6ulg-N | 3.2 | 11.5 | 85 | 483 | 6 | MOLECULE: FOLLICULIN; |
| 326 | 2x0s-A | 3.2 | 6.9 | 74 | 899 | 8 | MOLECULE: PYRUVATE PHOSPHATE DIKINASE; |
| 327 | 7aqq-N | 3.2 | 7.5 | 84 | 488 | 10 | MOLECULE: NADH-UBIQUINONE OXIDOREDUCTASE CHAIN 3; |
| 328 | 7b8s-C | 3.2 | 12.5 | 59 | 589 | 10 | MOLECULE: MULTIDRUG EFFLUX PUMP SUBUNIT ACRB,MULTIDRUG EFFL |
| 329 | 6swy-1 | 3.2 | 15.5 | 87 | 542 | 10 | MOLECULE: VACUOLAR IMPORT AND DEGRADATION PROTEIN 28; |
| 330 | 6hqv-A | 3.2 | 5.2 | 127 | 1556 | 9 | MOLECULE: PENTAFUNCTIONAL AROM POLYPEPTIDE; |
| 331 | 4b3n-A | 3.2 | 13.3 | 86 | 557 | 9 | MOLECULE: MALTOSE-BINDING PERIPLASMIC PROTEIN, TRIPARTITE M |
| 332 | 4p96-A | 3.2 | 7.5 | 58 | 278 | 10 | MOLECULE: FATTY ACID METABOLISM REGULATOR PROTEIN; |
| 333 | 2ip6-A | 3.2 | 2.6 | 50 | 87 | 6 | MOLECULE: PAPB; |
| 334 | 6dlu-P | 3.2 | 8.5 | 72 | 747 | 8 | MOLECULE: DYNAMIN-1; |
| 335 | 1wgz-A | 3.2 | 13.9 | 84 | 510 | 5 | MOLECULE: CARBOXYPEPTIDASE 1; |
| 336 | 7mez-B | 3.2 | 17.1 | 89 | 547 | 7 | MOLECULE: PHOSPHATIDYLINOSITOL 4,5-BISPHOSPHATE 3-KINASE CA |
| 337 | 6ryp-A | 3.1 | 4.3 | 47 | 164 | 11 | MOLECULE: LIPOPROTEIN SIGNAL PEPTIDASE; |
| 338 | 5y2g-A | 3.1 | 10.4 | 74 | 581 | 7 | MOLECULE: MALTOSE-BINDING PERIPLASMIC PROTEIN,PROTEIN B; |
| 339 | 3u2r-A | 3.1 | 4.4 | 46 | 135 | 15 | MOLECULE: REGULATORY PROTEIN MARR; |
| 340 | 6z5l-A | 3.1 | 7.8 | 62 | 251 | 5 | MOLECULE: MATRIX PROTEIN 1; |
| 341 | 2j68-A | 3.1 | 6 | 78 | 680 | 4 | MOLECULE: BACTERIAL DYNAMIN-LIKE PROTEIN; |
| 342 | 6vrb-C | 3.1 | 10.4 | 57 | 229 | 18 | MOLECULE: RNA (52-MER); |
| 343 | 3dy5-A | 3.1 | 11.6 | 57 | 1002 | 7 | MOLECULE: ALLENE OXIDE SYNTHASE-LIPOXYGENASE PROTEIN; |
| 344 | 4ihq-A | 3.1 | 15.1 | 63 | 512 | 10 | MOLECULE: FLAI ATPASE; |
| 345 | 3pvz-A | 3.1 | 13.9 | 61 | 372 | 7 | MOLECULE: UDP-N-ACETYLGLUCOSAMINE 4,6-DEHYDRATASE; |
| 346 | 3zx6-A | 3.1 | 7.2 | 57 | 303 | 5 | MOLECULE: HAMP, METHYL-ACCEPTING CHEMOTAXIS PROTEIN I; |
| 347 | 3dxj-D | 3.1 | 11.5 | 84 | 1504 | 6 | MOLECULE: DNA-DIRECTED RNA POLYMERASE SUBUNIT ALPHA; CHAIN |

| | | | | | | | |
|---|---|---|---|---|---|---|---|
| 348 | 1xqo-A | 3.1 | 8.8 | 75 | 253 | 9 | MOLECULE: 8-OXOGUANINE DNA GLYCOSYLASE; |
| 349 | 6qd5-A | 3.1 | 11.7 | 77 | 356 | 10 | MOLECULE: UREA TRANSPORTER 1; |
| 350 | 4nfu-B | 3.1 | 13.8 | 60 | 524 | 8 | MOLECULE: EDS1; |
| 351 | 6gw6-B | 3.1 | 3.1 | 61 | 149 | 8 | MOLECULE: RES TOXIN; |
| 352 | 3s1b-A | 3.1 | 2.5 | 34 | 34 | 3 | MOLECULE: VASCULAR ENDOTHELIAL GROWTH FACTOR A; |
| 353 | 6sih-A | 3.1 | 8.5 | 53 | 488 | 8 | MOLECULE: FLAGELLAR HOOK-ASSOCIATED PROTEIN 2; |
| 354 | 6uun-R | 3.1 | 3 | 58 | 354 | 2 | MOLECULE: GUANINE NUCLEOTIDE-BINDING PROTEIN G(S) SUBUNIT A |
| 355 | 5jqk-A | 3.1 | 4.3 | 66 | 631 | 8 | MOLECULE: PEPTIDASE, PUTATIVE; |
| 356 | 7nsb-a | 3.1 | 13.1 | 65 | 568 | 11 | MOLECULE: VACUOLAR IMPORT AND DEGRADATION PROTEIN 30; |
| 357 | 7mrw-B | 3.1 | 6.5 | 86 | 972 | 10 | MOLECULE: CYTOADHERENCE LINKED ASEXUAL PROTEIN 3.1; |
| 358 | 5wgr-A | 3.1 | 17.4 | 94 | 664 | 4 | MOLECULE: FLAVIN-DEPENDENT HALOGENASE; |
| 359 | 5im8-A | 3.1 | 2.8 | 41 | 60 | 5 | MOLECULE: E3 UBIQUITIN-PROTEIN LIGASE MIDLINE-1; |
| 360 | 4nqj-A | 3.1 | 7.7 | 63 | 177 | 11 | MOLECULE: E3 UBIQUITIN-PROTEIN LIGASE TRIM69; |
| 361 | 6xss-A | 3.1 | 5.7 | 59 | 260 | 12 | MOLECULE: C4_NAT_HFUSE-7900; |
| 362 | 5o5e-A | 3.1 | 8.4 | 77 | 381 | 6 | MOLECULE: UDP-N-ACETYLGLUCOSAMINE--DOLICHYL-PHOSPHATE N- |
| 363 | 6mag-A | 3.1 | 4 | 67 | 282 | 3 | MOLECULE: BBVCI ENDONUCLEASE SUBUNIT 2; |
| 364 | 2i5u-A | 3.1 | 3.4 | 46 | 77 | 4 | MOLECULE: DNAD DOMAIN PROTEIN; |
| 365 | 6tmh-d | 3.1 | 8.4 | 59 | 143 | 7 | MOLECULE: INHIBITOR OF F1; |
| 366 | 3o6x-A | 3.1 | 7 | 83 | 639 | 7 | MOLECULE: GLUTAMINE SYNTHETASE; |
| 367 | 4kp3-D | 3.1 | 5.5 | 55 | 82 | 7 | MOLECULE: UNCONVENTIONAL MYOSIN-VA; |
| 368 | 3hjl-A | 3.1 | 3.8 | 57 | 321 | 7 | MOLECULE: FLAGELLAR MOTOR SWITCH PROTEIN FLIG; |
| 369 | 3zt9-A | 3.1 | 11.9 | 62 | 192 | 5 | MOLECULE: SERINE PHOSPHATASE; |
| 370 | 6v3z-A | 3.1 | 3.9 | 55 | 178 | 7 | MOLECULE: SEN1395; |
| 371 | 5ihf-B | 3.1 | 2.7 | 53 | 109 | 4 | MOLECULE: VIRG-LIKE PROTEIN; |
| 372 | 6vm0-A | 3.1 | 7.8 | 64 | 363 | 13 | MOLECULE: GLYCINE RECEPTOR SUBUNIT ALPHAZ1; |
| 373 | 4xxk-A | 3.1 | 7.6 | 58 | 149 | 3 | MOLECULE: PHYCOBILIPROTEIN APCE; |
| 374 | 5xyv-C | 3.1 | 6.5 | 45 | 52 | 0 | MOLECULE: RHINO; |
| 375 | 6z6o-D | 3.1 | 7.5 | 60 | 542 | 7 | MOLECULE: HISTONE DEACETYLASE HDA1; |
| 376 | 4paf-A | 3.1 | 9.8 | 54 | 308 | 9 | MOLECULE: TRAP DICARBOXYLATE TRANSPORTER, DCTP SUBUNIT, PUT |
| 377 | 7bl5-8 | 3.1 | 4.4 | 68 | 157 | 9 | MOLECULE: 50S RIBOSOMAL PROTEIN L2; |
| 378 | 3crl-B | 3.1 | 6.2 | 83 | 383 | 6 | MOLECULE: PYRUVATE DEHYDROGENASE [LIPOAMIDE] KINASE ISOZYME |
| 379 | 5img-A | 3.1 | 12.6 | 95 | 467 | 8 | MOLECULE: DIPEPTIDASE; |
| 380 | 3r2x-C | 3.1 | 2.6 | 46 | 82 | 7 | MOLECULE: HEMAGGLUTININ; |
| 381 | 4neo-A | 3.1 | 2.6 | 51 | 83 | 8 | MOLECULE: PEPTIDE SYNTHETASE NRPS TYPE II-PCP; |
| 382 | 7js4-A | 3.1 | 16.6 | 91 | 953 | 10 | MOLECULE: F5/8 TYPE C DOMAIN PROTEIN; |

| | | | | | | | |
|---|---|---|---|---|---|---|---|
| 383 | 4wat-A | 3.1 | 5.1 | 71 | 332 | 6 | MOLECULE: PFRH5; |
| 384 | 5nwf-A | 3.1 | 3 | 38 | 203 | 5 | MOLECULE: FIC FAMILY PROTEIN; |
| 385 | 6veo-A | 3.1 | 4.8 | 57 | 136 | 9 | MOLECULE: ATPASE FAMILY AAA DOMAIN-CONTAINING PROTEIN 2B; |
| 386 | 3jbz-A | 3.1 | 14.4 | 82 | 960 | 4 | MOLECULE: SERINE/THREONINE-PROTEIN KINASE MTOR; |
| 387 | 6z5s-W | 3.1 | 5.4 | 44 | 94 | 0 | MOLECULE: LIGHT HARVESTING COMPLEX 1 PROTEIN W; |
| 388 | 1ufi-A | 3.1 | 5.4 | 40 | 48 | 15 | MOLECULE: MAJOR CENTROMERE AUTOANTIGEN B; |
| 389 | 6hvn-A | 3.1 | 2.6 | 42 | 175 | 7 | MOLECULE: DIADENYLATE CYCLASE; |
| 390 | 2lse-A | 3.1 | 3.3 | 60 | 101 | 10 | MOLECULE: FOUR HELIX BUNDLE PROTEIN; |
| 391 | 4wd9-A | 3.1 | 11.9 | 67 | 965 | 9 | MOLECULE: NISIN BIOSYNTHESIS PROTEIN NISB; |
| 392 | 4fcg-A | 3.1 | 15.9 | 70 | 296 | 11 | MOLECULE: UNCHARACTERIZED PROTEIN; |
| 393 | 2qw6-C | 3.1 | 3 | 55 | 83 | 13 | MOLECULE: AAA ATPASE, CENTRAL REGION; |
| 394 | 5fia-B | 3.1 | 12.2 | 102 | 397 | 4 | MOLECULE: LPIR1; |
| 395 | 6c48-D | 3.1 | 5.5 | 53 | 87 | 13 | MOLECULE: PROTEIN LIN-9 HOMOLOG; |
| 396 | 5cwf-A | 3.1 | 7 | 66 | 177 | 12 | MOLECULE: DESIGNED HELICAL REPEAT PROTEIN; |
| 397 | 6prk-A | 3.1 | 10.8 | 62 | 115 | 11 | MOLECULE: RICF; |
| 398 | 4cc9-B | 3.1 | 4.9 | 65 | 98 | 2 | MOLECULE: PROTEIN VPRBP; |
| 399 | 6yrf-A | 3.1 | 15.9 | 84 | 777 | 7 | MOLECULE: VEGETATIVE INSECTICIDAL PROTEIN; |
| 400 | 1kmi-Z | 3.1 | 4.8 | 45 | 177 | 11 | MOLECULE: CHEMOTAXIS PROTEIN CHEY; |
| 401 | 6jcx-F | 3.1 | 5.4 | 67 | 186 | 6 | MOLECULE: DNA-DIRECTED RNA POLYMERASE SUBUNIT ALPHA; |
| 402 | 7a5p-S | 3.1 | 3.1 | 41 | 46 | 12 | MOLECULE: U2 SNRNA; |
| 403 | 3gnl-A | 3.1 | 11.6 | 70 | 234 | 6 | MOLECULE: UNCHARACTERIZED PROTEIN, DUF633, LMOF2365_1472; |
| 404 | 6m0r-M | 3.1 | 5.9 | 57 | 71 | 11 | MOLECULE: V-TYPE PROTON ATPASE SUBUNIT C'; |
| 405 | 6sum-A | 3.1 | 4.3 | 58 | 335 | 7 | MOLECULE: AMICOUMACIN KINASE; |
| 406 | 3wdm-A | 3.1 | 8.1 | 61 | 259 | 7 | MOLECULE: 4-PHOSPHOPANTOATE--BETA-ALANINE LIGASE; |
| 407 | 7et3-i | 3.1 | 2.9 | 47 | 63 | 17 | MOLECULE: TRIPLEX CAPSID PROTEIN 2; |
| 408 | 5xef-A | 3.1 | 5 | 43 | 123 | 7 | MOLECULE: FLAGELLAR PROTEIN FLIS; |
| 409 | 1yhd-A | 3.1 | 4.2 | 55 | 92 | 11 | MOLECULE: UPF0269 PROTEIN YGGX; |
| 410 | 5iwz-A | 3.1 | 5.7 | 78 | 372 | 9 | MOLECULE: SYNAPTONEMAL COMPLEX PROTEIN 2; |
| 411 | 6e3y-E | 3.1 | 4.2 | 46 | 115 | 7 | MOLECULE: CALCITONIN GENE-RELATED PEPTIDE 1; |
| 412 | 6vm6-B | 3.1 | 6.2 | 60 | 440 | 5 | MOLECULE: SAVED DOMAIN-CONTAINING PROTEIN; |
| 413 | 3rgu-A | 3.1 | 3.7 | 57 | 88 | 5 | MOLECULE: FIMBRIAE-ASSOCIATED PROTEIN FAP1; |
| 414 | 7bwk-E | 3 | 3.3 | 60 | 116 | 8 | MOLECULE: ICMO (DOTL); |
| 415 | 3glf-G | 3 | 16.9 | 69 | 378 | 6 | MOLECULE: DNA POLYMERASE III SUBUNIT DELTA; |
| 416 | 4gae-A | 3 | 13 | 75 | 416 | 5 | MOLECULE: 1-DEOXY-D-XYLULOSE 5-PHOSPHATE REDUCTOISOMERASE, |
| 417 | 2x2v-A | 3 | 3.2 | 49 | 69 | 8 | MOLECULE: ATP SYNTHASE SUBUNIT C; |

| | | | | | | | |
|---|---|---|---|---|---|---|---|
| 418 | 7jgr-D | 3 | 11.3 | 73 | 441 | 5 | MOLECULE: ORIGIN RECOGNITION COMPLEX SUBUNIT 2; |
| 419 | 6wmk-A | 3 | 2.7 | 45 | 65 | 9 | MOLECULE: BETA SHEET HETERODIMER LHD29 - CHAIN A; |
| 420 | 3ilk-A | 3 | 10.7 | 48 | 239 | 8 | MOLECULE: UNCHARACTERIZED TRNA/RRNA METHYLTRANSFERASE HI038 |
| 421 | 6maf-A | 3 | 4.8 | 70 | 259 | 11 | MOLECULE: BBVCI ENDONUCLEASE SUBUNIT 1; |
| 422 | 6vzd-E | 3 | 5.9 | 51 | 85 | 6 | MOLECULE: PULMONARY SURFACTANT-ASSOCIATED PROTEIN B; |
| 423 | 5utg-A | 3 | 5.9 | 59 | 134 | 7 | MOLECULE: EGG-LYSIN; |
| 424 | 4okv-F | 3 | 4.3 | 45 | 66 | 9 | MOLECULE: HEAVY CHAIN OF 8H7 MAB; |
| 425 | 5h79-D | 3 | 9.1 | 53 | 142 | 4 | MOLECULE: IMMUNOGLOBULIN G-BINDING PROTEIN A; |
| 426 | 5x41-F | 3 | 14.5 | 77 | 244 | 10 | MOLECULE: COBALT ABC TRANSPORTER ATP-BINDING PROTEIN; |
| 427 | 1yns-A | 3 | 7.3 | 96 | 254 | 7 | MOLECULE: E-1 ENZYME; |
| 428 | 3kdq-C | 3 | 5.1 | 54 | 153 | 7 | MOLECULE: UNCHARACTERIZED CONSERVED PROTEIN; |
| 429 | 5h75-D | 3 | 3 | 45 | 225 | 9 | MOLECULE: MERSACIDIN DECARBOXYLASE,IMMUNOGLOBULIN G-BINDING |
| 430 | 5fb1-A | 3 | 3.6 | 72 | 174 | 11 | MOLECULE: NUCLEAR AUTOANTIGEN SP-100; |
| 431 | 5wp0-B | 3 | 14.7 | 63 | 274 | 6 | MOLECULE: NH(3)-DEPENDENT NAD(+) SYNTHETASE; |
| 432 | 6w38-A | 3 | 4.8 | 50 | 331 | 12 | MOLECULE: TERMINAL NUCLEOTIDYLTRANSFERASE 5C; |
| 433 | 7f1t-A | 3 | 5.1 | 71 | 423 | 7 | MOLECULE: C-C MOTIF CHEMOKINE 3,C-C CHEMOKINE RECEPTOR TYPE |
| 434 | 2kyz-A | 3 | 2.8 | 42 | 67 | 10 | MOLECULE: HEAVY METAL BINDING PROTEIN; |
| 435 | 6upn-A | 3 | 9.7 | 54 | 245 | 17 | MOLECULE: ENDOPHILIN-B1; |
| 436 | 7t62-A | 3 | 9.1 | 91 | 561 | 7 | MOLECULE: GLYPICAN-2; |
| 437 | 5c0x-G | 3 | 10 | 67 | 237 | 7 | MOLECULE: EXOSOME COMPLEX COMPONENT RRP45; |
| 438 | 4ifd-H | 3 | 12.3 | 61 | 295 | 2 | MOLECULE: EXOSOME COMPLEX COMPONENT RRP45; |
| 439 | 4bbp-A | 3 | 7.6 | 72 | 266 | 15 | MOLECULE: ZINC ABC TRANSPORTER, PERIPLASMIC ZINC-BINDING PR |
| 440 | 6mur-B | 3 | 7.9 | 56 | 162 | 11 | MOLECULE: UNCHARACTERIZED PROTEIN; |
| 441 | 7eu3-2 | 3 | 5.8 | 52 | 129 | 6 | MOLECULE: NAD(P)H-QUINONE OXIDOREDUCTASE SUBUNIT 1, CHLOROP |
| 442 | 4hqo-B | 3 | 11.1 | 63 | 260 | 10 | MOLECULE: SPOROZOITE SURFACE PROTEIN 2; |
| 443 | 2etd-A | 3 | 6.6 | 72 | 141 | 4 | MOLECULE: LEMA PROTEIN; |
| 444 | 4u8u-G | 3 | 4.6 | 62 | 151 | 2 | MOLECULE: GLOBIN A CHAIN; |
| 445 | 3wx4-A | 3 | 8.7 | 63 | 98 | 14 | MOLECULE: ANTI-RESTRICTION ENDONUCLEASE; |
| 446 | 3hw2-A | 3 | 7.1 | 64 | 308 | 3 | MOLECULE: PROTEIN SIFA; |
| 447 | 6yj4-L | 3 | 8.3 | 82 | 655 | 7 | MOLECULE: NADH-UBIQUINONE OXIDOREDUCTASE CHAIN 3; |
| 448 | 2dgj-A | 3 | 7.7 | 71 | 246 | 7 | MOLECULE: HYPOTHETICAL PROTEIN EBHA; |
| 449 | 3iym-A | 3 | 8.6 | 73 | 396 | 10 | MOLECULE: CAPSID PROTEIN; |
| 450 | 3odn-A | 3 | 9.3 | 94 | 366 | 1 | MOLECULE: DALLY-LIKE PROTEIN; |
| 451 | 2d5w-A | 3 | 9.7 | 75 | 602 | 5 | MOLECULE: PEPTIDE ABC TRANSPORTER, PEPTIDE-BINDING PROTEIN; |
| 452 | 1nu7-D | 3 | 6.2 | 77 | 282 | 6 | MOLECULE: THROMBIN LIGHT CHAIN; |

| | | | | | | | |
|---|---|---|---|---|---|---|---|
| 453 | 6urt-A | 3 | 10.1 | 68 | 331 | 4 | MOLECULE: LOW CONDUCTANCE MECHANOSENSITIVE CHANNEL YNAI; |
| 454 | 4x4w-A | 3 | 10.1 | 63 | 398 | 10 | MOLECULE: CCA TRNA NUCLEOTIDYLTRANSFERASE 1, MITOCHONDRIAL; |
| 455 | 3oov-A | 3 | 5.2 | 41 | 164 | 12 | MOLECULE: METHYL-ACCEPTING CHEMOTAXIS PROTEIN, PUTATIVE; |
| 456 | 5fno-A | 3 | 9.8 | 65 | 568 | 9 | MOLECULE: MANGANESE LIPOXYGENASE; |
| 457 | 6xqk-A | 3 | 2.1 | 37 | 47 | 11 | MOLECULE: CAMP-DEPENDENT PROTEIN KINASE REGULATORY SUBUNIT; |
| 458 | 7kgm-A | 3 | 4.9 | 82 | 512 | 6 | MOLECULE: PUTATIVE EXPORTED PROTEIN; |
| 459 | 6dgc-A | 3 | 10.5 | 42 | 147 | 0 | MOLECULE: ISC1926 TNPA C-TERMINAL CATALYTIC DOMAIN; |
| 460 | 6gse-A | 3 | 10.8 | 73 | 159 | 8 | MOLECULE: ACTIVITY-REGULATED CYTOSKELETON-ASSOCIATED PROTEI |
| 461 | 7nza-A | 3 | 7.7 | 64 | 141 | 5 | MOLECULE: ODORANT BINDING PROTEIN FROM VARROA DESTRUCTOR, F |
| 462 | 6xtt-B | 3 | 11.1 | 61 | 116 | 7 | MOLECULE: NTTA; |
| 463 | 5n8k-C | 3 | 8.2 | 49 | 80 | 4 | MOLECULE: GALACTOCEREBROSIDASE; |
| 464 | 3bjd-C | 3 | 6.8 | 78 | 324 | 10 | MOLECULE: PUTATIVE 3-OXOACYL-(ACYL-CARRIER-PROTEIN) SYNTHAS |
| 465 | 5lfj-B | 3 | 5.1 | 60 | 113 | 8 | MOLECULE: BACTERIAL PROTEASOME ACTIVATOR; |
| 466 | 6oiw-A | 3 | 4.9 | 100 | 504 | 7 | MOLECULE: DEOXYGUANOSINETRIPHOSPHATE TRIPHOSPHOHYDROLASE; |
| 467 | 6br8-A | 3 | 13.8 | 79 | 249 | 5 | MOLECULE: PROTEIN A6 HOMOLOG; |
| 468 | 2c2l-A | 3 | 10.1 | 68 | 281 | 6 | MOLECULE: CARBOXY TERMINUS OF HSP70-INTERACTING PROTEIN; |
| 469 | 6z11-H | 3 | 17 | 56 | 193 | 2 | MOLECULE: DNA-DIRECTED RNA POLYMERASE SUBUNIT ALPHA; |
| 470 | 6c6r-A | 3 | 8.6 | 68 | 451 | 6 | MOLECULE: SQUALENE MONOOXYGENASE; |
| 471 | 2es4-E | 3 | 2.6 | 46 | 278 | 7 | MOLECULE: LIPASE; |
| 472 | 3jrt-A | 3 | 5.9 | 65 | 166 | 9 | MOLECULE: INTEGRON CASSETTE PROTEIN VPC_CASS2; |
| 473 | 6cgh-A | 3 | 3.7 | 54 | 89 | 9 | MOLECULE: DNAJ HOMOLOG SUBFAMILY C MEMBER 2; |
| 474 | 6ysg-A | 3 | 17.8 | 84 | 1264 | 11 | MOLECULE: MG-CHELATASE SUBUNIT CHLH; |
| 475 | 1jj2-O | 3 | 7.2 | 65 | 143 | 9 | MOLECULE: 23S RRNA; |
| 476 | 6hyd-A | 3 | 13.8 | 86 | 1574 | 10 | MOLECULE: MIDASIN,MIDASIN,MIDASIN; |
| 477 | 7jsj-A | 3 | 10.6 | 77 | 468 | 9 | MOLECULE: SOLUTE CARRIER FAMILY 13 MEMBER 5; |
| 478 | 6ah0-w | 3 | 5.4 | 53 | 443 | 4 | MOLECULE: U5SNRNA; |
| 479 | 6fwr-A | 3 | 9.6 | 82 | 699 | 4 | MOLECULE: ATP-DEPENDENT DNA HELICASE DING; |
| 480 | 4wpe-A | 3 | 6.1 | 79 | 275 | 13 | MOLECULE: CYTOKINESIS PROTEIN 2; |
| 481 | 6em5-R | 3 | 3.6 | 45 | 120 | 18 | MOLECULE: 5.8S RIBOSOMAL RNA; |
| 482 | 5vvr-B | 3 | 8 | 80 | 1207 | 14 | MOLECULE: DNA-DIRECTED RNA POLYMERASE II SUBUNIT RPB1; |
| 483 | 2e62-A | 3 | 2.8 | 43 | 61 | 9 | MOLECULE: PROTEIN AT5G25060; |
| 484 | 4z7x-A | 3 | 7.7 | 67 | 211 | 4 | MOLECULE: MDBA; |
| 485 | 2khv-A | 3 | 3.7 | 41 | 106 | 0 | MOLECULE: PHAGE INTEGRASE; |
| 486 | 6xi6-A | 3 | 5.7 | 53 | 269 | 8 | MOLECULE: HELICAL FUSION DESIGN; |
| 487 | 3bpt-A | 3 | 17.6 | 80 | 362 | 5 | MOLECULE: 3-HYDROXYISOBUTYRYL-COA HYDROLASE; |

| | | | | | | | |
|---|---|---|---|---|---|---|---|
| 488 | 3owg-A | 3 | 16.6 | 102 | 457 | 5 | MOLECULE: POLY(A) POLYMERASE CATALYTIC SUBUNIT; |
| 489 | 6w17-F | 3 | 14.1 | 51 | 168 | 8 | MOLECULE: ACTIN-RELATED PROTEIN 3; |
| 490 | 5ztc-A | 3 | 11.8 | 71 | 197 | 14 | MOLECULE: LMO2088 PROTEIN; |
| 491 | 6vjr-A | 3 | 5 | 61 | 502 | 11 | MOLECULE: VINYLACETYL-COA DELTA-ISOMERASE; |
| 492 | 5j0l-A | 3 | 3.9 | 65 | 130 | 11 | MOLECULE: DESIGNED PROTEIN 3L6HC2_2; |
| 493 | 4zs9-B | 3 | 3.2 | 50 | 378 | 8 | MOLECULE: SUGAR BINDING PROTEIN OF ABC TRANSPORTER SYSTEM; |
| 494 | 6tdz-C | 3 | 11.9 | 76 | 530 | 5 | MOLECULE: OLIGOMYCIN SENSITIVITY CONFERRING PROTEIN (OSCP); |
| 495 | 6z5u-F | 3 | 9.9 | 53 | 193 | 9 | MOLECULE: ABC TRANSPORTER PERMEASE; |
| 496 | 7b93-L | 3 | 9.6 | 92 | 607 | 10 | MOLECULE: NADH-UBIQUINONE OXIDOREDUCTASE CHAIN 3; |
| 497 | 5i04-A | 3 | 13.4 | 81 | 668 | 6 | MOLECULE: MALTOSE-BINDING PERIPLASMIC PROTEIN,ENDOGLIN; |
| 498 | 6nd4-J | 3 | 8.4 | 60 | 493 | 3 | MOLECULE: ETS RRNA; |
| 499 | 7a6h-O | 3 | 10.3 | 54 | 512 | 7 | MOLECULE: DNA-DIRECTED RNA POLYMERASE III SUBUNIT RPC1; |
| 500 | 6z16-d | 3 | 4.9 | 70 | 489 | 7 | MOLECULE: MULTISUBUNIT NA+/H+ ANTIPORTER, A SUBUNIT; |
| 501 | 4x2c-A | 3 | 5.6 | 54 | 207 | 9 | MOLECULE: FIC FAMILY PROTEIN PUTATIVE FILAMENTATION INDUCED |
| 502 | 2w4s-A | 3 | 3 | 56 | 86 | 13 | MOLECULE: ANKYRIN-REPEAT PROTEIN; |
| 503 | 4dn7-A | 2.9 | 4.1 | 82 | 406 | 4 | MOLECULE: ABC TRANSPORTER, ATP-BINDING PROTEIN; |
| 504 | 7jrl-A | 2.9 | 10.2 | 94 | 457 | 11 | MOLECULE: F5/8 TYPE C DOMAIN PROTEIN; |
| 505 | 1fx2-A | 2.9 | 3.4 | 59 | 235 | 10 | MOLECULE: RECEPTOR-TYPE ADENYLATE CYCLASE GRESAG 4.1; |
| 506 | 3g5b-A | 2.9 | 2.8 | 77 | 383 | 6 | MOLECULE: NETRIN RECEPTOR UNC5B; |
| 507 | 6rie-A | 2.9 | 19.4 | 133 | 1285 | 9 | MOLECULE: DNA-DEPENDENT RNA POLYMERASE SUBUNIT RPO147; |
| 508 | 5anz-A | 2.9 | 8.6 | 81 | 375 | 5 | MOLECULE: SOLUBLE LYTIC TRANSGLYCOSYLASE B3; |
| 509 | 3mkw-B | 2.9 | 3.3 | 47 | 115 | 6 | MOLECULE: DNA (5'- |
| 510 | 7o8c-A | 2.9 | 3.4 | 49 | 582 | 6 | MOLECULE: SURFACE GLYCAN-BINDING PROTEIN BO2743; |
| 511 | 6ks5-B | 2.9 | 6.5 | 85 | 313 | 12 | MOLECULE: TYPE IV SECRETION PROTEIN DOT; |
| 512 | 5byh-D | 2.9 | 11.1 | 86 | 1372 | 2 | MOLECULE: DNA-DIRECTED RNA POLYMERASE SUBUNIT ALPHA; |
| 513 | 5z7b-B | 2.9 | 9.8 | 88 | 197 | 5 | MOLECULE: PADR FAMILY TRANSCRIPTIONAL REGULATOR; |
| 514 | 1k32-A | 2.9 | 9.9 | 96 | 1023 | 3 | MOLECULE: TRICORN PROTEASE; |
| 515 | 6snr-A | 2.9 | 9.5 | 58 | 402 | 10 | MOLECULE: LIPID II:GLYCINE GLYCYLTRANSFERASE; |
| 516 | 6ird-B | 2.9 | 3.8 | 57 | 254 | 5 | MOLECULE: 1-PHOSPHATIDYLINOSITOL 4,5-BISPHOSPHATE PHOSPHODI |
| 517 | 4g33-A | 2.9 | 14.2 | 86 | 661 | 9 | MOLECULE: 15S-LIPOXYGENASE; |
| 518 | 2qe9-A | 2.9 | 8.1 | 68 | 165 | 4 | MOLECULE: UNCHARACTERIZED PROTEIN YIZA; |
| 519 | 6uva-E | 2.9 | 15 | 51 | 111 | 4 | MOLECULE: GUANINE NUCLEOTIDE-BINDING PROTEIN G(S) SUBUNIT A |
| 520 | 6of3-A | 2.9 | 5.1 | 75 | 187 | 7 | MOLECULE: RIBONUCLEASE; |
| 521 | 5jw9-B | 2.9 | 8.4 | 47 | 115 | 4 | MOLECULE: AF4/FMR2 FAMILY MEMBER 4; |
| 522 | 4m70-R | 2.9 | 2.6 | 41 | 56 | 0 | MOLECULE: RX PROTEIN; |

| | | | | | | | |
|---|---|---|---|---|---|---|---|
| 523 | 1hlv-A | 2.9 | 13 | 57 | 131 | 11 | MOLECULE: CENP-B BOX DNA; |
| 524 | 1ysm-A | 2.9 | 4.7 | 42 | 55 | 10 | MOLECULE: CALCYCLIN-BINDING PROTEIN; |
| 525 | 2pyw-A | 2.9 | 10.6 | 77 | 417 | 6 | MOLECULE: UNCHARACTERIZED PROTEIN; |
| 526 | 6ff1-A | 2.9 | 2.5 | 40 | 69 | 18 | MOLECULE: CSOZ; |
| 527 | 6v85-F | 2.9 | 3.1 | 44 | 47 | 5 | MOLECULE: RNA-DIRECTED RNA POLYMERASE L; |
| 528 | 6vls-D | 2.9 | 9.2 | 74 | 963 | 7 | MOLECULE: MALTOSE/MALTODEXTRIN-BINDING PERIPLASMIC PROTEIN, |
| 529 | 6z16-f | 2.9 | 4.7 | 45 | 88 | 11 | MOLECULE: MULTISUBUNIT NA+/H+ ANTIPORTER, A SUBUNIT; |
| 530 | 5v8f-6 | 2.9 | 6.7 | 89 | 692 | 7 | MOLECULE: DNA REPLICATION LICENSING FACTOR MCM2; |
| 531 | 4ykn-A | 2.9 | 12.9 | 65 | 1204 | 9 | MOLECULE: PHOSPHATIDYLINOSITOL 3-KINASE REGULATORY SUBUNIT |
| 532 | 7m4p-A | 2.9 | 10.9 | 101 | 1049 | 6 | MOLECULE: EFFLUX PUMP MEMBRANE TRANSPORTER; |
| 533 | 2yia-A | 2.9 | 12.7 | 73 | 790 | 7 | MOLECULE: RNA-DIRECTED RNA POLYMERASE; |
| 534 | 3ksy-A | 2.9 | 4.9 | 96 | 1007 | 9 | MOLECULE: SON OF SEVENLESS HOMOLOG 1; |
| 535 | 1yre-B | 2.9 | 9.3 | 45 | 187 | 0 | MOLECULE: HYPOTHETICAL PROTEIN PA3270; |
| 536 | 7ns4-i | 2.9 | 4.4 | 53 | 358 | 6 | MOLECULE: E3 UBIQUITIN-PROTEIN LIGASE RMD5; |
| 537 | 3hj1-B | 2.9 | 8.5 | 77 | 387 | 6 | MOLECULE: MINOR EDITOSOME-ASSOCIATED TUTASE; |
| 538 | 6bx3-F | 2.9 | 5.6 | 54 | 215 | 11 | MOLECULE: HISTONE-LYSINE N-METHYLTRANSFERASE, H3 LYSINE-4 S |
| 539 | 4n4u-A | 2.9 | 10.3 | 51 | 304 | 10 | MOLECULE: PUTATIVE ABC TRANSPORTER PERIPLASMIC SOLUTE-BINDI |
| 540 | 7ebe-G | 2.9 | 21 | 88 | 535 | 8 | MOLECULE: ISOCITRATE LYASE; |
| 541 | 4jvy-A | 2.9 | 5.6 | 66 | 191 | 11 | MOLECULE: FEMALE GERMLINE-SPECIFIC TUMOR SUPPRESSOR GLD-1; |
| 542 | 4xc6-A | 2.9 | 12.2 | 98 | 1067 | 6 | MOLECULE: ISOBUTYRYL-COA MUTASE FUSED; |
| 543 | 6cfw-G | 2.9 | 4.6 | 49 | 113 | 6 | MOLECULE: MONOVALENT CATION/H+ ANTIPORTER SUBUNIT D; |
| 544 | 3g06-A | 2.9 | 5.9 | 77 | 601 | 6 | MOLECULE: SSPH2 (LEUCINE-RICH REPEAT PROTEIN); |
| 545 | 6qm7-H | 2.9 | 9.2 | 72 | 229 | 3 | MOLECULE: PROTEASOME ALPHA1 CHAIN; |
| 546 | 5ue8-A | 2.9 | 11.3 | 90 | 847 | 8 | MOLECULE: PROTEIN UNC-13 HOMOLOG A; |
| 547 | 7eeb-K | 2.9 | 4.3 | 52 | 143 | 10 | MOLECULE: ENHANCED GREEN FLUORESCENT PROTEIN,CATION CHANNEL |
| 548 | 6ln2-A | 2.9 | 5.3 | 73 | 437 | 3 | MOLECULE: GLUCAGON-LIKE PEPTIDE 1 RECEPTOR,RUBREDOXIN,GLUCA |
| 549 | 6ezn-B | 2.9 | 8.9 | 46 | 110 | 9 | MOLECULE: DOLICHYL-DIPHOSPHOOLIGOSACCHARIDE--PROTEIN |
| 550 | 5y08-A | 2.9 | 4.7 | 60 | 205 | 10 | MOLECULE: 4-HYDROXY-2,2'-BIPYRROLE-5-METHANOL SYNTHASE PIGH |
| 551 | 6t1z-A | 2.9 | 6 | 65 | 393 | 6 | MOLECULE: LMRP INTEGRAL MEMBRANE PROTEIN; |
| 552 | 3f44-A | 2.9 | 14.7 | 62 | 210 | 13 | MOLECULE: PUTATIVE MONOOXYGENASE; |
| 553 | 3mp8-A | 2.9 | 14.3 | 74 | 513 | 15 | MOLECULE: MALTOSE-BINDING PERIPLASMIC PROTEIN,LINKER,SAGA-A |
| 554 | 7sr1-A | 2.9 | 3 | 42 | 123 | 7 | MOLECULE: SORTING NEXIN-25; |
| 555 | 2rno-A | 2.9 | 3.5 | 71 | 110 | 7 | MOLECULE: PUTATIVE DNA-BINDING PROTEIN; |
| 556 | 4eyy-R | 2.9 | 4.8 | 45 | 59 | 7 | MOLECULE: ICMR; |
| 557 | 3t38-A | 2.9 | 8.4 | 59 | 199 | 7 | MOLECULE: ARSENATE REDUCTASE; |

| | | | | | | | |
|---|---|---|---|---|---|---|---|
| 558 | 6v3f-A | 2.9 | 8 | 96 | 1234 | 3 | MOLECULE: NPC1-LIKE INTRACELLULAR CHOLESTEROL TRANSPORTER 1 |
| 559 | 2oxl-A | 2.9 | 9.4 | 52 | 62 | 15 | MOLECULE: HYPOTHETICAL PROTEIN YMGB; |
| 560 | 6soy-A | 2.9 | 10.1 | 77 | 323 | 8 | MOLECULE: ESAG6, SUBUNIT OF HETERODIMERIC TRANSFERRIN RECEP |
| 561 | 2qpx-A | 2.9 | 4.3 | 47 | 376 | 9 | MOLECULE: PREDICTED METAL-DEPENDENT HYDROLASE OF THE TIM-BA |
| 562 | 5ukv-A | 2.9 | 4.2 | 49 | 74 | 12 | MOLECULE: ATP-BINDING PROTEIN; |
| 563 | 2mpn-A | 2.9 | 4.6 | 45 | 68 | 4 | MOLECULE: INNER MEMBRANE PROTEIN YGAP; |
| 564 | 6idp-A | 2.9 | 8.5 | 77 | 438 | 3 | MOLECULE: MATE FAMILY EFFLUX TRANSPORTER; |
| 565 | 4oo8-D | 2.9 | 11.4 | 87 | 1163 | 7 | MOLECULE: CRISPR-ASSOCIATED ENDONUCLEASE CAS9/CSN1; |
| 566 | 3bqy-A | 2.9 | 10.8 | 69 | 205 | 6 | MOLECULE: PUTATIVE TETR FAMILY TRANSCRIPTIONAL REGULATOR; |
| 567 | 1pjq-B | 2.9 | 12.4 | 69 | 456 | 4 | MOLECULE: SIROHEME SYNTHASE; |
| 568 | 5m41-A | 2.9 | 4.6 | 115 | 712 | 11 | MOLECULE: NIGRITOXINE; |
| 569 | 6lum-G | 2.9 | 4 | 58 | 123 | 7 | MOLECULE: SUCCINATE DEHYDROGENASE SUBUNIT C; |
| 570 | 4fzo-A | 2.9 | 5 | 54 | 82 | 9 | MOLECULE: URANYL BINDING PROTEIN; |
| 571 | 7mi6-A | 2.9 | 13.2 | 97 | 2419 | 5 | MOLECULE: FUSION PROTEIN OF DYNEIN AND ENDOLYSIN; |
| 572 | 2ols-A | 2.9 | 15.1 | 59 | 725 | 7 | MOLECULE: PHOSPHOENOLPYRUVATE SYNTHASE; |
| 573 | 6wtw-A | 2.9 | 9.7 | 66 | 489 | 9 | MOLECULE: DASS FAMILY SODIUM-COUPLED ANION SYMPORTER; |
| 574 | 5udb-7 | 2.9 | 16.3 | 94 | 726 | 5 | MOLECULE: DNA REPLICATION LICENSING FACTOR MCM2; |
| 575 | 5tj4-H | 2.9 | 11.7 | 87 | 537 | 10 | MOLECULE: SUGAR ABC TRANSPORTER SUBSTRATE-BINDING PROTEIN,G |
| 576 | 4lws-A | 2.9 | 7 | 58 | 100 | 7 | MOLECULE: UNCHARACTERIZED PROTEIN; |
| 577 | 6khn-A | 2.9 | 11.2 | 71 | 490 | 8 | MOLECULE: TRYPTOPHAN DECARBOXYLASE 1; |
| 578 | 5vyl-A | 2.9 | 10.8 | 80 | 502 | 1 | MOLECULE: INNER TEGUMENT PROTEIN; |
| 579 | 1iru-G | 2.9 | 5 | 65 | 245 | 8 | MOLECULE: 20S PROTEASOME; |
| 580 | 4d7x-A | 2.9 | 3.6 | 48 | 86 | 17 | MOLECULE: MEDIATOR OF RNA POLYMERASE II TRANSCRIPTION SUBUN |
| 581 | 5jm8-A | 2.9 | 9.6 | 77 | 556 | 4 | MOLECULE: AEROBACTIN SYNTHASE IUCA; |
| 582 | 7aoz-I | 2.9 | 3.9 | 71 | 464 | 11 | MOLECULE: DNA-DIRECTED RNA POLYMERASE 147 KDA POLYPEPTIDE; |
| 583 | 5c9e-A | 2.9 | 10.5 | 63 | 263 | 13 | MOLECULE: SEPL; |
| 584 | 3t58-B | 2.9 | 5.8 | 67 | 504 | 3 | MOLECULE: SULFHYDRYL OXIDASE 1; |
| 585 | 4cti-D | 2.9 | 8.2 | 57 | 222 | 5 | MOLECULE: OSMOLARITY SENSOR PROTEIN ENVZ, AF1503; |
| 586 | 4z39-A | 2.9 | 3.1 | 58 | 121 | 7 | MOLECULE: ODORANT-BINDING PROTEIN; |
| 587 | 1kyq-B | 2.9 | 9.8 | 73 | 273 | 8 | MOLECULE: SIROHEME BIOSYNTHESIS PROTEIN MET8; |
| 588 | 5j10-A | 2.9 | 4.4 | 39 | 70 | 23 | MOLECULE: PEPTIDE DESIGN 2L4HC2_24; |
| 589 | 5a1q-A | 2.9 | 3.2 | 57 | 69 | 11 | MOLECULE: AF1502; |
| 590 | 5i5k-B | 2.9 | 14.7 | 95 | 1634 | 9 | MOLECULE: COMPLEMENT C5; |
| 591 | 5lbm-C | 2.9 | 2.7 | 41 | 90 | 5 | MOLECULE: TRANSCRIPTIONAL REPRESSOR FRMR; |
| 592 | 7sh3-A | 2.9 | 3 | 49 | 63 | 12 | MOLECULE: SYNTHETIC VIRB8 MINIPROTEIN BINDER; |

| | | | | | | | |
|---|---|---|---|---|---|---|---|
| 593 | 5xsv-A | 2.9 | 4.9 | 82 | 486 | 7 | MOLECULE: CHITINASE; |
| 594 | 7but-A | 2.9 | 5.9 | 77 | 142 | 6 | MOLECULE: ACINIFORM SPIDROIN; |
| 595 | 5bq9-B | 2.9 | 9.6 | 67 | 437 | 9 | MOLECULE: UNCHARACTERIZED PROTEIN; |
| 596 | 6yvu-B | 2.9 | 8.4 | 75 | 1191 | 5 | MOLECULE: STRUCTURAL MAINTENANCE OF CHROMOSOMES PROTEIN 2,S |
| 597 | 1r4g-A | 2.9 | 3 | 46 | 53 | 9 | MOLECULE: RNA POLYMERASE ALPHA SUBUNIT; |
| 598 | 7drt-B | 2.9 | 4.3 | 65 | 496 | 6 | MOLECULE: PROTEIN WNT-3A; |
| 599 | 4f4h-A | 2.9 | 17 | 87 | 540 | 10 | MOLECULE: GLUTAMINE DEPENDENT NAD+ SYNTHETASE; |
| 600 | 4bk0-B | 2.9 | 4.1 | 38 | 97 | 8 | MOLECULE: ATP-DEPENDENT DNA HELICASE Q5; |
| 601 | 4gcz-B | 2.9 | 8.4 | 60 | 378 | 2 | MOLECULE: BLUE-LIGHT PHOTORECEPTOR, SENSOR PROTEIN FIXL; |
| 602 | 4uxv-A | 2.9 | 5.8 | 72 | 522 | 7 | MOLECULE: SEPTATION RING FORMATION REGULATOR EZRA; |
| 603 | 4zel-A | 2.9 | 9.7 | 60 | 550 | 12 | MOLECULE: DOPAMINE BETA-HYDROXYLASE; |
| 604 | 6hwj-A | 2.9 | 9.9 | 69 | 428 | 10 | MOLECULE: MALTOKINASE; |
| 605 | 5ogs-A | 2.9 | 10 | 77 | 403 | 4 | MOLECULE: WD REPEAT AND HMG-BOX DNA-BINDING PROTEIN 1; |
| 606 | 2lqx-A | 2.9 | 3.2 | 38 | 41 | 5 | MOLECULE: TRYPSIN INHIBITOR BWI-2C; |
| 607 | 6elu-A | 2.9 | 4.3 | 62 | 193 | 8 | MOLECULE: SERUM RESISTANCE ASSOCIATED; VSG PROTEIN; |
| 608 | 5jje-B | 2.9 | 3.9 | 44 | 63 | 0 | MOLECULE: SENSORY RHODOPSIN-2; |
| 609 | 7lt1-A | 2.9 | 3.7 | 67 | 403 | 3 | MOLECULE: PROTEIN MB21D2; |
| 610 | 6t58-A | 2.9 | 9.7 | 67 | 524 | 7 | MOLECULE: CELLULAR TUMOR ANTIGEN P53,PROTEIN S100-A4,PROTEI |
| 611 | 5t58-A | 2.9 | 4.4 | 65 | 233 | 5 | MOLECULE: KLLA0F02343P; |
| 612 | 3wfw-A | 2.9 | 8.2 | 61 | 138 | 10 | MOLECULE: HEMOGLOBIN-LIKE FLAVOPROTEIN FUSED TO ROADBLOCK/L |
| 613 | 2zy4-F | 2.9 | 16.3 | 71 | 535 | 4 | MOLECULE: L-ASPARTATE BETA-DECARBOXYLASE; |
| 614 | 6gho-B | 2.8 | 13.4 | 83 | 274 | 5 | MOLECULE: REGULATORY PROTEIN SPX; |
| 615 | 1ysy-A | 2.8 | 3.4 | 52 | 85 | 4 | MOLECULE: REPLICASE POLYPROTEIN 1AB (PP1AB) (ORF1AB); |
| 616 | 2win-M | 2.8 | 6.8 | 64 | 84 | 8 | MOLECULE: COMPLEMENT C3 BETA CHAIN; |
| 617 | 3s8r-B | 2.8 | 10.5 | 71 | 686 | 4 | MOLECULE: GLUTARYL-7-AMINOCEPHALOSPORANIC-ACID ACYLASE; |
| 618 | 6fhs-C | 2.8 | 8.3 | 67 | 459 | 6 | MOLECULE: RUVB-LIKE HELICASE; |
| 619 | 3omd-A | 2.8 | 4 | 67 | 145 | 9 | MOLECULE: UNCHARACTERIZED PROTEIN; |
| 620 | 3hai-A | 2.8 | 8.6 | 82 | 294 | 1 | MOLECULE: HUMAN PACSIN1 F-BAR; |
| 621 | 7kpx-C | 2.8 | 15.6 | 68 | 1352 | 1 | MOLECULE: MEIOTIC MRNA STABILITY PROTEIN KINASE SSN3; |
| 622 | 6uz8-A | 2.8 | 12.3 | 65 | 734 | 3 | MOLECULE: SHORT TRANSIENT RECEPTOR POTENTIAL CHANNEL 6; |
| 623 | 7abw-B | 2.8 | 8.6 | 59 | 365 | 3 | MOLECULE: PEPSY DOMAIN-CONTAINING PROTEIN; |
| 624 | 3ats-A | 2.8 | 8 | 76 | 352 | 8 | MOLECULE: PUTATIVE UNCHARACTERIZED PROTEIN; |
| 625 | 6c0f-D | 2.8 | 10.6 | 82 | 190 | 10 | MOLECULE: SACCHAROMYCES CEREVISIAE S288C 35S PRE-RIBOSOMAL |
| 626 | 6xqj-A | 2.8 | 5.2 | 71 | 173 | 8 | MOLECULE: PROTEIN VPR,UV EXCISION REPAIR PROTEIN RAD23 HOMO |
| 627 | 6y37-C | 2.8 | 3.1 | 56 | 118 | 7 | MOLECULE: HAPB; |

| | | | | | | | |
|---|---|---|---|---|---|---|---|
| 628 | 1sgm-A | 2.8 | 12.1 | 63 | 184 | 5 | MOLECULE: PUTATIVE HTH-TYPE TRANSCRIPTIONAL REGULATOR YXAF; |
| 629 | 5xfs-A | 2.8 | 3.8 | 42 | 78 | 2 | MOLECULE: PE FAMILY PROTEIN PE8; |
| 630 | 6ei9-A | 2.8 | 4.5 | 58 | 306 | 10 | MOLECULE: TRNA-DIHYDROURIDINE SYNTHASE B; |
| 631 | 7oca-E | 2.8 | 4.7 | 57 | 158 | 21 | MOLECULE: GLUTAMATE RECEPTOR 1; |
| 632 | 6k4e-B | 2.8 | 2.9 | 54 | 246 | 13 | MOLECULE: HAMP DOMAIN-CONTAINING PROTEIN; |
| 633 | 2o7t-A | 2.8 | 11.2 | 59 | 188 | 10 | MOLECULE: TRANSCRIPTIONAL REGULATOR; |
| 634 | 5gvv-A | 2.8 | 12.5 | 81 | 392 | 2 | MOLECULE: GLYCOSYL TRANSFERASE FAMILY 8; |
| 635 | 6nr8-4 | 2.8 | 4.7 | 43 | 104 | 9 | MOLECULE: PREFOLDIN SUBUNIT 1; |
| 636 | 2yfb-B | 2.8 | 7 | 62 | 238 | 8 | MOLECULE: METHYL-ACCEPTING CHEMOTAXIS TRANSDUCER; |
| 637 | 2p22-A | 2.8 | 5 | 44 | 168 | 9 | MOLECULE: SUPPRESSOR PROTEIN STP22 OF TEMPERATURE-SENSITIVE |
| 638 | 4he8-D | 2.8 | 4.3 | 45 | 160 | 11 | MOLECULE: NADH-QUINONE OXIDOREDUCTASE SUBUNIT 7; |
| 639 | 6hd8-B | 2.8 | 6.4 | 59 | 232 | 10 | MOLECULE: NANOBODY,MALTOSE/MALTODEXTRIN-BINDING PERIPLASMIC |
| 640 | 6n7o-B | 2.8 | 2.8 | 37 | 39 | 16 | MOLECULE: GIL01 GP7; |
| 641 | 5z9w-A | 2.8 | 11.5 | 50 | 388 | 10 | MOLECULE: EBOLAVIRUS NUCLEOPROTEIN (RESIDUES 19-406); |
| 642 | 6r24-A | 2.8 | 12.7 | 79 | 159 | 5 | MOLECULE: TRANSPOSON TY3-I GAG-POL POLYPROTEIN; |
| 643 | 2bma-A | 2.8 | 5 | 60 | 467 | 8 | MOLECULE: GLUTAMATE DEHYDROGENASE (NADP+); |
| 644 | 1te5-A | 2.8 | 4.1 | 59 | 253 | 8 | MOLECULE: CONSERVED HYPOTHETICAL PROTEIN; |
| 645 | 7rsf-A | 2.8 | 11.7 | 68 | 380 | 10 | MOLECULE: ACETYLORNITHINE DEACETYLASE; |
| 646 | 2qeb-B | 2.8 | 4.2 | 82 | 145 | 7 | MOLECULE: D7R4 PROTEIN; |
| 647 | 5an6-A | 2.8 | 10.7 | 67 | 123 | 1 | MOLECULE: CRISPR-ASSOCIATED PROTEIN, CSM2 FAMILY; |
| 648 | 5wwl-N | 2.8 | 5.7 | 59 | 155 | 2 | MOLECULE: CENTROMERE PROTEIN MIS12; |
| 649 | 6thk-A | 2.8 | 4.6 | 51 | 466 | 6 | MOLECULE: PYOCIN S5; |
| 650 | 1ixs-B | 2.8 | 7.8 | 68 | 315 | 10 | MOLECULE: HOLLIDAY JUNCTION DNA HELICASE RUVA; |
| 651 | 2j5y-A | 2.8 | 2.2 | 38 | 61 | 3 | MOLECULE: PEPTOSTREPTOCOCCAL ALBUMIN-BINDING PROTEIN; |
| 652 | 7ogp-E | 2.8 | 7.7 | 60 | 377 | 7 | MOLECULE: PHIKZ055,PHIKZ056.1; |
| 653 | 2peg-B | 2.8 | 3.3 | 63 | 139 | 3 | MOLECULE: HEMOGLOBIN SUBUNIT ALPHA; |
| 654 | 1lrz-A | 2.8 | 13.1 | 77 | 400 | 5 | MOLECULE: FACTOR ESSENTIAL FOR EXPRESSION OF METHICILLIN |
| 655 | 2luh-A | 2.8 | 7.3 | 59 | 167 | 7 | MOLECULE: VACUOLAR PROTEIN SORTING-ASSOCIATED PROTEIN VTA1; |
| 656 | 6rd4-1 | 2.8 | 13.2 | 82 | 595 | 6 | MOLECULE: ASA-10: POLYTOMELLA F-ATP SYNTHASE ASSOCIATED SUB |
| 657 | 7emf-D | 2.8 | 6.9 | 60 | 158 | 13 | MOLECULE: MEDIATOR OF RNA POLYMERASE II TRANSCRIPTION SUBUN |
| 658 | 4eyw-B | 2.8 | 4.1 | 46 | 634 | 4 | MOLECULE: CARNITINE O-PALMITOYLTRANSFERASE 2, MITOCHONDRIAL |
| 659 | 6ynw-H | 2.8 | 3.6 | 44 | 75 | 5 | MOLECULE: SUBUNIT C; |
| 660 | 4ghl-A | 2.8 | 3.3 | 62 | 132 | 8 | MOLECULE: SHORT PALINDROMIC RNA AGACAGCAUAUGCUGUCU; |
| 661 | 5y9o-A | 2.8 | 6.3 | 61 | 401 | 8 | MOLECULE: WIPA; |
| 662 | 4bx9-C | 2.8 | 5.2 | 53 | 95 | 11 | MOLECULE: VACUOLAR PROTEIN SORTING-ASSOCIATED PROTEIN 33A; |

| | | | | | | | |
|---|---|---|---|---|---|---|---|
| 663 | 6ae9-A | 2.8 | 2.9 | 53 | 253 | 11 | MOLECULE: PROBABLE PROTEIN PHOSPHATASE 2C 1; |
| 664 | 6f1e-A | 2.8 | 6.3 | 60 | 143 | 3 | MOLECULE: INTERFERON GAMMA; |
| 665 | 6lqo-L | 2.8 | 5.7 | 49 | 94 | 10 | MOLECULE: CYTOPLASMIC ENVELOPMENT PROTEIN 1; |
| 666 | 3rpm-A | 2.8 | 11.1 | 85 | 450 | 13 | MOLECULE: BETA-N-ACETYL-HEXOSAMINIDASE; |
| 667 | 2hl7-A | 2.8 | 2.5 | 39 | 82 | 8 | MOLECULE: CYTOCHROME C-TYPE BIOGENESIS PROTEIN CCMH; |
| 668 | 3h36-A | 2.8 | 3.2 | 47 | 78 | 11 | MOLECULE: POLYRIBONUCLEOTIDE NUCLEOTIDYLTRANSFERASE; |
| 669 | 2i1k-A | 2.8 | 7.9 | 58 | 545 | 7 | MOLECULE: MOESIN; |
| 670 | 1wxq-A | 2.8 | 10.1 | 78 | 344 | 14 | MOLECULE: GTP-BINDING PROTEIN; |
| 671 | 2vj4-A | 2.8 | 9.9 | 62 | 292 | 13 | MOLECULE: PROTEIN MXIC; |
| 672 | 5hx2-A | 2.8 | 13.5 | 82 | 1030 | 12 | MOLECULE: BASEPLATE WEDGE PROTEIN GP7; |
| 673 | 2klz-A | 2.8 | 5.5 | 41 | 46 | 12 | MOLECULE: ATAXIN-3; |
| 674 | 6m9k-D | 2.8 | 3.1 | 50 | 67 | 8 | MOLECULE: EXONUCLEASE; |
| 675 | 2khq-A | 2.8 | 4.2 | 40 | 110 | 5 | MOLECULE: INTEGRASE; |
| 676 | 6bmt-B | 2.8 | 3.4 | 49 | 70 | 4 | MOLECULE: MYELOPEROXIDASE; |
| 677 | 5b1q-A | 2.8 | 8 | 74 | 448 | 8 | MOLECULE: U14 PROTEIN; |
| 678 | 7ou2-B | 2.8 | 9.6 | 80 | 650 | 10 | MOLECULE: DNA MISMATCH REPAIR PROTEIN MUTS; |
| 679 | 6l1k-A | 2.8 | 3.4 | 59 | 384 | 10 | MOLECULE: NADH-DEPENDENT BUTANOL DEHYDROGENASE A; |
| 680 | 6uuj-G | 2.8 | 4.3 | 42 | 74 | 2 | MOLECULE: PE FAMILY IMMUNOMODULATOR PE5; |
| 681 | 3i3l-A | 2.8 | 15.3 | 80 | 550 | 11 | MOLECULE: ALKYLHALIDASE CMLS; |
| 682 | 5gpo-B | 2.8 | 7.5 | 57 | 124 | 7 | MOLECULE: SENSOR PROTEIN CZCS; |
| 683 | 5m4s-A | 2.8 | 3.3 | 64 | 209 | 6 | MOLECULE: TRANSCRIPTION INITIATION FACTOR IIA SUBUNIT 2,TRA |
| 684 | 4v1a-w | 2.8 | 3.9 | 68 | 387 | 12 | MOLECULE: MITORIBOSOMAL PROTEIN ML37, MRPL37; |
| 685 | 5gn1-D | 2.8 | 8.4 | 76 | 312 | 12 | MOLECULE: ATP-DEPENDENT HELICASE FUN30; |
| 686 | 2ipc-A | 2.8 | 4 | 75 | 939 | 8 | MOLECULE: PREPROTEIN TRANSLOCASE SECA SUBUNIT; |
| 687 | 7ac1-A | 2.8 | 2.9 | 59 | 75 | 7 | MOLECULE: TRANSCRIPTION INITIATION FACTOR TFIID SUBUNIT 4; |
| 688 | 7cun-G | 2.8 | 12.3 | 97 | 895 | 8 | MOLECULE: INTEGRATOR COMPLEX SUBUNIT 1; |
| 689 | 6th1-R | 2.8 | 12.8 | 96 | 360 | 7 | MOLECULE: IMMEDIATE EARLY PROTEIN 1; |
| 690 | 4qvh-A | 2.8 | 11.3 | 83 | 596 | 8 | MOLECULE: MALTOSE-BINDING PERIPLASMIC PROTEIN, 4'-PHOSPHOPA |
| 691 | 3lay-J | 2.8 | 2.9 | 42 | 81 | 0 | MOLECULE: ZINC RESISTANCE-ASSOCIATED PROTEIN; |
| 692 | 4uyb-A | 2.8 | 11.9 | 81 | 400 | 11 | MOLECULE: SEC14-LIKE PROTEIN 3; |
| 693 | 6mrr-A | 2.8 | 3.3 | 46 | 68 | 7 | MOLECULE: FOLDIT1; |
| 694 | 6s7t-F | 2.8 | 12.9 | 68 | 250 | 4 | MOLECULE: DOLICHYL-DIPHOSPHOOLIGOSACCHARIDE--PROTEIN |
| 695 | 4djg-A | 2.8 | 4 | 41 | 49 | 7 | MOLECULE: PLECTIN-RELATED PROTEIN; |
| 696 | 4y2i-A | 2.8 | 3.1 | 42 | 65 | 7 | MOLECULE: PUTATIVE METAL-BINDING TRANSPORT PROTEIN; |
| 697 | 1skv-A | 2.8 | 5.2 | 40 | 64 | 10 | MOLECULE: HYPOTHETICAL 7.5 KDA PROTEIN; |

| | | | | | | | |
|---|---|---|---|---|---|---|---|
| 698 | 4uy3-A | 2.8 | 7.1 | 82 | 194 | 11 | MOLECULE: SEPTATION RING FORMATION REGULATOR EZRA; |
| 699 | 6wuc-H | 2.8 | 12.2 | 73 | 179 | 10 | MOLECULE: INNER KINETOCHORE SUBUNIT MCM16; |
| 700 | 6arh-B | 2.8 | 12.2 | 57 | 303 | 11 | MOLECULE: N-ACETYLNEURAMINATE LYASE; |
| 701 | 5oqm-n | 2.8 | 7.3 | 57 | 136 | 5 | MOLECULE: DNA-DIRECTED RNA POLYMERASE II SUBUNIT RPB1; |
| 702 | 6rfq-P | 2.8 | 11.9 | 72 | 123 | 7 | MOLECULE: SUBUNIT NUAM OF NADH:UBIQUINONE OXIDOREDUCTASE (C |
| 703 | 3jbl-A | 2.8 | 16.5 | 82 | 909 | 9 | MOLECULE: NLR FAMILY CARD DOMAIN-CONTAINING PROTEIN 4; |
| 704 | 5h5v-A | 2.8 | 4.8 | 48 | 352 | 8 | MOLECULE: FLAGELLAR HOOK-ASSOCIATED PROTEIN 2; |
| 705 | 2nd2-A | 2.8 | 2.3 | 39 | 44 | 5 | MOLECULE: DE NOVO MINI PROTEIN HHH_06; |
| 706 | 6rre-D | 2.8 | 5.2 | 90 | 427 | 8 | MOLECULE: ADENOSINE MONOPHOSPHATE-PROTEIN HYDROLASE SIDD; |
| 707 | 6ogy-K | 2.8 | 12.6 | 69 | 827 | 6 | MOLECULE: RNA-DEPENDENT RNA POLYMERASE OF ROTAVIRUS A; |
| 708 | 7jhj-R | 2.8 | 4.3 | 64 | 277 | 6 | MOLECULE: GUANINE NUCLEOTIDE-BINDING PROTEIN G(I) SUBUNIT A |
| 709 | 5cd4-I | 2.8 | 8.9 | 69 | 494 | 9 | MOLECULE: CRISPR SYSTEM CASCADE SUBUNIT CASE; |
| 710 | 1qgi-A | 2.8 | 3.1 | 66 | 259 | 9 | MOLECULE: PROTEIN (CHITOSANASE); |
| 711 | 4aim-A | 2.8 | 24.7 | 72 | 698 | 6 | MOLECULE: POLYRIBONUCLEOTIDE NUCLEOTIDYLTRANSFERASE; |
| 712 | 2z3x-A | 2.8 | 3.8 | 44 | 56 | 9 | MOLECULE: SMALL, ACID-SOLUBLE SPORE PROTEIN C; |
| 713 | 4nsm-A | 2.8 | 5.2 | 39 | 71 | 13 | MOLECULE: COLLAGEN-LIKE PROTEIN SCLB; |
| 714 | 6kyf-A | 2.8 | 4.6 | 59 | 137 | 8 | MOLECULE: ACRF11; |
| 715 | 4af8-A | 2.8 | 10.9 | 60 | 333 | 5 | MOLECULE: METACASPASE MCA2; |
| 716 | 1pix-A | 2.8 | 6 | 57 | 586 | 9 | MOLECULE: GLUTACONYL-COA DECARBOXYLASE A SUBUNIT; |
| 717 | 5k12-A | 2.8 | 10.2 | 74 | 294 | 3 | MOLECULE: GLUTAMATE DEHYDROGENASE 1, MITOCHONDRIAL; |
| 718 | 3c3w-A | 2.8 | 8.9 | 66 | 211 | 5 | MOLECULE: TWO COMPONENT TRANSCRIPTIONAL REGULATORY PROTEIN |
| 719 | 4or2-B | 2.8 | 4.9 | 80 | 366 | 3 | MOLECULE: SOLUBLE CYTOCHROME B562, METABOTROPIC GLUTAMATE R |
| 720 | 6c03-A | 2.8 | 5 | 54 | 138 | 6 | MOLECULE: PUTATIVE RNA POLYMERASE ECF-SUBFAMILY SIGMA FACTO |
| 721 | 6r3q-A | 2.8 | 20 | 83 | 841 | 5 | MOLECULE: ADENYLATE CYCLASE 9; |
| 722 | 7onu-F | 2.8 | 11.4 | 80 | 276 | 10 | MOLECULE: 3-HYDROXYACYL-COA DEHYDROGENASE TYPE-2; |
| 723 | 4rta-B | 2.8 | 2.5 | 41 | 71 | 7 | MOLECULE: PROTEIN DPY-30 HOMOLOG; |
| 724 | 6yj4-J | 2.8 | 8.5 | 60 | 185 | 7 | MOLECULE: NADH-UBIQUINONE OXIDOREDUCTASE CHAIN 3; |
| 725 | 5j1f-A | 2.8 | 10.2 | 76 | 188 | 3 | MOLECULE: PLECTIN,PLECTIN; |
| 726 | 5tck-A | 2.8 | 3.7 | 48 | 139 | 8 | MOLECULE: UNCHARACTERIZED PROTEIN; |
| 727 | 2ota-B | 2.8 | 2.5 | 50 | 61 | 12 | MOLECULE: UPF0352 PROTEIN CPS_2611; |
| 728 | 3x38-B | 2.8 | 3.3 | 49 | 79 | 12 | MOLECULE: MITOCHONDRIAL MORPHOGENESIS PROTEIN SLD7; |
| 729 | 6c70-A | 2.8 | 9.1 | 69 | 388 | 10 | MOLECULE: ODORANT RECEPTOR; |
| 730 | 4lid-A | 2.8 | 8.2 | 56 | 86 | 7 | MOLECULE: A-100; |
| 731 | 7p47-B | 2.8 | 4.9 | 45 | 69 | 18 | MOLECULE: STRUCTURAL MAINTENANCE OF CHROMOSOMES PROTEIN 5; |
| 732 | 2c5z-A | 2.8 | 4.8 | 51 | 93 | 8 | MOLECULE: SET DOMAIN PROTEIN 2; |

| | | | | | | | |
|---|---|---|---|---|---|---|---|
| 733 | 2apl-A | 2.8 | 9.2 | 48 | 149 | 8 | MOLECULE: HYPOTHETICAL PROTEIN PG0816; |
| 734 | 6wc3-B | 2.8 | 4.2 | 47 | 94 | 23 | MOLECULE: PROTEIN TRANSPORT PROTEIN TIP20; |
| 735 | 6twr-A | 2.8 | 2.1 | 37 | 169 | 3 | MOLECULE: BETA-GLUCOSIDE BGL OPERON ANTITERMINATOR BGLG FAM |
| 736 | 2l35-A | 2.8 | 3.9 | 44 | 62 | 5 | MOLECULE: DAP12-NKG2C_TM; |
| 737 | 5j2l-A | 2.8 | 5 | 53 | 76 | 13 | MOLECULE: PROTEIN DESIGN 2L4HC2_11; |
| 738 | 4c0j-A | 2.8 | 17.1 | 96 | 404 | 8 | MOLECULE: MITOCHONDRIAL RHO GTPASE; |
| 739 | 5yfp-B | 2.8 | 8 | 72 | 927 | 11 | MOLECULE: EXOCYST COMPLEX COMPONENT SEC3; |
| 740 | 7kpv-B | 2.8 | 11.1 | 81 | 290 | 9 | MOLECULE: MEIOTIC MRNA STABILITY PROTEIN KINASE SSN3; |
| 741 | 6ozu-A | 2.8 | 7.7 | 60 | 235 | 15 | MOLECULE: EUKARYOTIC TRANSLATION INITIATION FACTOR 4 GAMMA |
| 742 | 3gor-A | 2.8 | 4 | 81 | 157 | 4 | MOLECULE: PUTATIVE METAL-DEPENDENT HYDROLASE; |
| 743 | 4ad8-A | 2.8 | 4.3 | 62 | 452 | 5 | MOLECULE: DNA REPAIR PROTEIN RECN; |
| 744 | 4b94-A | 2.8 | 5.4 | 55 | 141 | 11 | MOLECULE: DUAL SPECIFICITY PROTEIN KINASE TTK; |
| 745 | 3pvs-D | 2.8 | 12.5 | 97 | 424 | 7 | MOLECULE: REPLICATION-ASSOCIATED RECOMBINATION PROTEIN A; |
| 746 | 6iva-A | 2.7 | 5.7 | 106 | 420 | 9 | MOLECULE: OXALOACETATE DECARBOXYLASE BETA CHAIN; |
| 747 | 6yiz-A | 2.7 | 4.5 | 46 | 211 | 11 | MOLECULE: TRANSCRIPTIONAL REGULATOR MVFR; |
| 748 | 6hpn-A | 2.7 | 4.3 | 61 | 212 | 5 | MOLECULE: ANTIGEN, P35; |
| 749 | 2rkh-A | 2.7 | 7.1 | 69 | 164 | 7 | MOLECULE: PUTATIVE APHA-LIKE TRANSCRIPTION FACTOR; |
| 750 | 6tj1-C | 2.7 | 5.3 | 43 | 74 | 14 | MOLECULE: DE NOVO DESIGNED WSHC6; |
| 751 | 7mhz-A | 2.7 | 5.7 | 97 | 490 | 7 | MOLECULE: SONIC HEDGEHOG PROTEIN N-PRODUCT PEPTIDE; |
| 752 | 6t8h-B | 2.7 | 7.3 | 106 | 1188 | 9 | MOLECULE: DNA POLYMERASE SLIDING CLAMP; |
| 753 | 6vme-C | 2.7 | 4.8 | 48 | 67 | 6 | MOLECULE: TUMOR SUSCEPTIBILITY GENE 101 PROTEIN; |
| 754 | 7l48-A | 2.7 | 14 | 83 | 522 | 4 | MOLECULE: CAS12F; |
| 755 | 2rq6-A | 2.7 | 10.2 | 58 | 138 | 5 | MOLECULE: ATP SYNTHASE EPSILON CHAIN; |
| 756 | 4jle-B | 2.7 | 3.7 | 47 | 159 | 9 | MOLECULE: PHIST; |
| 757 | 6h0q-A | 2.7 | 2.8 | 55 | 102 | 11 | MOLECULE: TYPE I MODULAR POLYKETIDE SYNTHASE; |
| 758 | 3o0y-A | 2.7 | 3.8 | 64 | 585 | 3 | MOLECULE: LIPOPROTEIN; |
| 759 | 6zbj-B | 2.7 | 16.9 | 100 | 501 | 6 | MOLECULE: PRECURSOR OF THE MAJOR MEROZOITE SURFACE ANTIGENS |
| 760 | 5ean-A | 2.7 | 15.5 | 73 | 1051 | 8 | MOLECULE: DNA REPLICATION ATP-DEPENDENT HELICASE/NUCLEASE D |
| 761 | 6qm7-C | 2.7 | 6.1 | 70 | 276 | 11 | MOLECULE: PROTEASOME ALPHA1 CHAIN; |
| 762 | 2imd-A | 2.7 | 10.2 | 72 | 203 | 6 | MOLECULE: 2-HYDROXYCHROMENE-2-CARBOXYLATE ISOMERASE; |
| 763 | 5yx5-B | 2.7 | 4.4 | 63 | 270 | 8 | MOLECULE: 49 PROTEIN; |
| 764 | 5h11-A | 2.7 | 8.9 | 75 | 494 | 13 | MOLECULE: UNCHARACTERIZED PROTEIN; |
| 765 | 2ptm-A | 2.7 | 7.7 | 43 | 192 | 14 | MOLECULE: HYPERPOLARIZATION-ACTIVATED (IH) CHANNEL; |
| 766 | 4pac-A | 2.7 | 12.1 | 69 | 153 | 9 | MOLECULE: HISTIDINE-CONTAINING PHOSPHOTRANSFER PROTEIN 2; |
| 767 | 4r0g-A | 2.7 | 11.1 | 84 | 253 | 11 | MOLECULE: UNCHARACTERIZED PROTEIN; |

| | | | | | | | |
|---|---|---|---|---|---|---|---|
| 768 | 5b0l-B | 2.7 | 11.1 | 93 | 334 | 3 | MOLECULE: MOEN5,DNA-BINDING PROTEIN 7D; |
| 769 | 6wm6-A | 2.7 | 13.6 | 82 | 536 | 4 | MOLECULE: EXTRACELLULAR SOLUTE-BINDING PROTEIN, FAMILY 5; |
| 770 | 5z79-C | 2.7 | 6.9 | 77 | 599 | 8 | MOLECULE: HYDROXYMETHYLDIHYDROPTERIN PYROPHOSPHOKINASE- |
| 771 | 4tko-B | 2.7 | 6.5 | 61 | 326 | 5 | MOLECULE: EMRA; |
| 772 | 6rd4-4 | 2.7 | 5.4 | 81 | 290 | 12 | MOLECULE: ASA-10: POLYTOMELLA F-ATP SYNTHASE ASSOCIATED SUB |
| 773 | 2khm-A | 2.7 | 3.9 | 50 | 140 | 10 | MOLECULE: FIBROIN-3; |
| 774 | 4u9r-A | 2.7 | 3 | 55 | 147 | 9 | MOLECULE: CZCP CATION EFFLUX P1-ATPASE; |
| 775 | 5z2c-A | 2.7 | 10.1 | 89 | 445 | 3 | MOLECULE: ALPHA-PROTEIN KINASE 1; |
| 776 | 7awt-D | 2.7 | 14.3 | 78 | 589 | 6 | MOLECULE: NADH-QUINONE OXIDOREDUCTASE SUBUNIT B; |
| 777 | 1flc-B | 2.7 | 8.1 | 60 | 162 | 8 | MOLECULE: HAEMAGGLUTININ-ESTERASE-FUSION GLYCOPROTEIN; |
| 778 | 6dmp-B | 2.7 | 4.6 | 49 | 82 | 14 | MOLECULE: DESIGNED ORTHOGONAL PROTEIN DHD13_XAAA_A; |
| 779 | 5nbs-A | 2.7 | 11.4 | 87 | 842 | 2 | MOLECULE: BETA-GLUCOSIDASE; |
| 780 | 5dpl-A | 2.7 | 14.2 | 103 | 521 | 5 | MOLECULE: PROTEIN LYSINE METHYLTRANSFERASE 2; |
| 781 | 6s6b-I | 2.7 | 9.6 | 89 | 284 | 4 | MOLECULE: CRISPR-ASSOCIATED PROTEIN, CMR5 FAMILY; |
| 782 | 4b3i-A | 2.7 | 9.9 | 96 | 730 | 10 | MOLECULE: FATTY ACID BETA-OXIDATION COMPLEX ALPHA-CHAIN FAD |
| 783 | 6kf4-G | 2.7 | 5.5 | 37 | 166 | 8 | MOLECULE: DNA-DIRECTED RNA POLYMERASE SUBUNIT; |
| 784 | 6zzx-F | 2.7 | 5.8 | 45 | 165 | 9 | MOLECULE: PHOTOSYSTEM I P700 CHLOROPHYLL A APOPROTEIN A1; |
| 785 | 6elc-A | 2.7 | 7.9 | 61 | 356 | 10 | MOLECULE: VARIANT SURFACE GLYCOPROTEIN; |
| 786 | 6zdx-A | 2.7 | 3 | 51 | 100 | 8 | MOLECULE: RIFIN; |
| 787 | 5wtn-A | 2.7 | 9.4 | 65 | 268 | 8 | MOLECULE: REPLICATION INITIATION AND MEMBRANE ATTACHMENT PR |
| 788 | 2q0y-A | 2.7 | 3.6 | 40 | 153 | 10 | MOLECULE: GCN5-RELATED N-ACETYLTRANSFERASE; |
| 789 | 2lyi-A | 2.7 | 5.4 | 90 | 167 | 9 | MOLECULE: PROTEIN (ENTITY); |
| 790 | 6p18-Q | 2.7 | 3.6 | 59 | 156 | 3 | MOLECULE: DNA (67-MER) FRAGMENT CARRYING PHAGE-21 PR' PROMO |
| 791 | 5awf-A | 2.7 | 5 | 73 | 385 | 7 | MOLECULE: FES CLUSTER ASSEMBLY PROTEIN SUFB; |
| 792 | 3k33-A | 2.7 | 3.4 | 66 | 124 | 8 | MOLECULE: DEATH ON CURING PROTEIN; |
| 793 | 6yse-A | 2.7 | 4.8 | 37 | 46 | 11 | MOLECULE: GP4; |
| 794 | 4i0x-K | 2.7 | 5.6 | 54 | 77 | 4 | MOLECULE: ESAT-6-LIKE PROTEIN MAB_3112; |
| 795 | 5y6o-D | 2.7 | 15.8 | 50 | 108 | 4 | MOLECULE: DEATH DOMAIN-ASSOCIATED PROTEIN 6,TRANSCRIPTIONAL |
| 796 | 5xj5-A | 2.7 | 10.8 | 71 | 200 | 7 | MOLECULE: GLYCEROL-3-PHOSPHATE ACYLTRANSFERASE; |
| 797 | 7c4o-A | 2.7 | 4 | 39 | 48 | 8 | MOLECULE: TRANSCRIPTION FACTOR HES-1; |
| 798 | 6g2d-C | 2.7 | 20.8 | 95 | 2121 | 5 | MOLECULE: ACETYL-COA CARBOXYLASE 1; |
| 799 | 4rp5-A | 2.7 | 6.3 | 47 | 94 | 4 | MOLECULE: DISKS LARGE 1 TUMOR SUPPRESSOR PROTEIN; |
| 800 | 1iuf-A | 2.7 | 8 | 66 | 144 | 6 | MOLECULE: CENTROMERE ABP1 PROTEIN; |
| 801 | 5idr-D | 2.7 | 16.5 | 64 | 221 | 9 | MOLECULE: DSBA-LIKE PROTEIN; |
| 802 | 3p3o-A | 2.7 | 2.7 | 57 | 416 | 11 | MOLECULE: CYTOCHROME P450; |

| | | | | | | | |
|---|---|---|---|---|---|---|---|
| 803 | 3nmz-A | 2.7 | 17.9 | 78 | 406 | 9 | MOLECULE: APC VARIANT PROTEIN; |
| 804 | 5olk-A | 2.7 | 3.4 | 64 | 395 | 2 | MOLECULE: RIBONUCLEOSIDE-DIPHOSPHATE REDUCTASE, BETA SUBUNI |
| 805 | 6iqc-A | 2.7 | 2.5 | 45 | 103 | 9 | MOLECULE: DNA-BINDING PROTEIN SSO0352; |
| 806 | 5j5l-A | 2.7 | 4.4 | 49 | 152 | 8 | MOLECULE: UNCHARACTERIZED PROTEIN; |
| 807 | 4fbj-A | 2.7 | 3 | 48 | 250 | 13 | MOLECULE: HYPOTHETICAL PROTEIN; |
| 808 | 4myy-B | 2.7 | 2.9 | 54 | 84 | 15 | MOLECULE: CURG, CURH FUSION PROTEIN; |
| 809 | 3bru-A | 2.7 | 12.6 | 73 | 191 | 4 | MOLECULE: REGULATORY PROTEIN, TETR FAMILY; |
| 810 | 3p01-A | 2.7 | 4.3 | 49 | 178 | 8 | MOLECULE: TWO-COMPONENT RESPONSE REGULATOR; |
| 811 | 5f15-A | 2.7 | 5 | 91 | 541 | 2 | MOLECULE: 4-AMINO-4-DEOXY-L-ARABINOSE (L-ARA4N) TRANSFERASE |
| 812 | 2gnx-A | 2.7 | 8.2 | 52 | 288 | 6 | MOLECULE: HYPOTHETICAL PROTEIN; |
| 813 | 6n04-B | 2.7 | 13.6 | 76 | 366 | 9 | MOLECULE: ABSH3; |
| 814 | 6v6b-B | 2.7 | 13 | 59 | 572 | 7 | MOLECULE: GAMMA-TUBULIN COMPLEX COMPONENT 3; |
| 815 | 4x00-A | 2.7 | 4.2 | 64 | 273 | 9 | MOLECULE: PUTATIVE HYDROLASE; |
| 816 | 6n7m-A | 2.7 | 6.2 | 47 | 63 | 6 | MOLECULE: HYPOTHETICAL PROTEIN CD630_05490; |
| 817 | 6m7d-A | 2.7 | 13.6 | 97 | 412 | 5 | MOLECULE: NUCLEOPROTEIN; |
| 818 | 6jo5-F | 2.7 | 5.8 | 45 | 165 | 9 | MOLECULE: PHOTOSYSTEM I P700 CHLOROPHYLL A APOPROTEIN A1; |
| 819 | 6ybw-y | 2.7 | 6.7 | 62 | 162 | 6 | MOLECULE: 40S RIBOSOMAL PROTEIN S4, X ISOFORM; |
| 820 | 6e3d-A | 2.7 | 3.5 | 61 | 510 | 7 | MOLECULE: PERIPLASMIC DIPEPTIDE-BINDING LIPOPROTEIN DPPA; |
| 821 | 6noc-A | 2.7 | 15.6 | 84 | 402 | 10 | MOLECULE: FEM-3 MRNA-BINDING FACTOR 2; |
| 822 | 6k9k-A | 2.7 | 15 | 87 | 2501 | 3 | MOLECULE: SERINE-PROTEIN KINASE ATM; |
| 823 | 1s12-A | 2.7 | 3.6 | 57 | 94 | 9 | MOLECULE: HYPOTHETICAL PROTEIN TM1457; |
| 824 | 5k2a-A | 2.7 | 6.5 | 74 | 398 | 12 | MOLECULE: ADENOSINE RECEPTOR A2A/SOLUBLE CYTOCHROME B562 CH |
| 825 | 4c2e-A | 2.7 | 9.1 | 53 | 435 | 9 | MOLECULE: CARBOXY-TERMINAL PROCESSING PROTEASE CTPB; |
| 826 | 2qup-A | 2.7 | 3.9 | 53 | 119 | 8 | MOLECULE: BH1478 PROTEIN; |
| 827 | 4g80-I | 2.7 | 7.2 | 62 | 139 | 3 | MOLECULE: VOLTAGE-SENSOR CONTAINING PHOSPHATASE; |
| 828 | 3mem-A | 2.7 | 17.1 | 84 | 453 | 7 | MOLECULE: PUTATIVE SIGNAL TRANSDUCTION PROTEIN; |
| 829 | 6ypu-p | 2.7 | 7.4 | 44 | 88 | 14 | MOLECULE: 16S RIBOSOMAL RNA; |
| 830 | 3jac-A | 2.7 | 13 | 67 | 918 | 0 | MOLECULE: PIEZO-TYPE MECHANOSENSITIVE ION CHANNEL COMPONENT |
| 831 | 1qrj-A | 2.7 | 10.5 | 91 | 214 | 10 | MOLECULE: HTLV-I CAPSID PROTEIN; |
| 832 | 5ebz-A | 2.7 | 13.3 | 91 | 655 | 9 | MOLECULE: INHIBITOR OF NUCLEAR FACTOR KAPPA-B KINASE SUBUNI |
| 833 | 5gox-A | 2.7 | 7.2 | 61 | 181 | 11 | MOLECULE: DNA REPAIR PROTEIN RAD50; |
| 834 | 2opd-A | 2.7 | 4.1 | 51 | 121 | 18 | MOLECULE: PILX; |
| 835 | 7eu3-F | 2.7 | 5.2 | 74 | 671 | 9 | MOLECULE: NAD(P)H-QUINONE OXIDOREDUCTASE SUBUNIT 1, CHLOROP |
| 836 | 5xe7-B | 2.7 | 8.3 | 101 | 280 | 6 | MOLECULE: ECF RNA POLYMERASE SIGMA FACTOR SIGJ; |
| 837 | 3ucq-A | 2.7 | 11.7 | 77 | 651 | 9 | MOLECULE: AMYLOSUCRASE; |

| | | | | | | | |
|---|---|---|---|---|---|---|---|
| 838 | 2pih-A | 2.7 | 6.3 | 57 | 123 | 9 | MOLECULE: PROTEIN YMCA; |
| 839 | 6to1-A | 2.7 | 10.5 | 63 | 566 | 6 | MOLECULE: MINOR FIMBRIUM SUBUNIT MFA5; |
| 840 | 4gc5-A | 2.7 | 2.7 | 53 | 318 | 4 | MOLECULE: DIMETHYLADENOSINE TRANSFERASE 1, MITOCHONDRIAL; |
| 841 | 5m0n-A | 2.7 | 8.4 | 69 | 423 | 6 | MOLECULE: TERMINAL OLEFIN-FORMING FATTY ACID DECARBOXYLASE; |
| 842 | 6uak-A | 2.7 | 8.9 | 66 | 298 | 9 | MOLECULE: SAM DEPENDENT METHYLTRANSFERASE LAHSB; |
| 843 | 6xgx-A | 2.7 | 8.4 | 86 | 402 | 7 | MOLECULE: MUTATOR FAMILY TRANSPOSASE; |
| 844 | 4kpk-A | 2.7 | 10.4 | 64 | 265 | 9 | MOLECULE: ENOYL-COA HYDRATASE/ISOMERASE; |
| 845 | 3f1i-H | 2.7 | 8.4 | 56 | 98 | 13 | MOLECULE: HEPATOCYTE GROWTH FACTOR-REGULATED TYROSINE |
| 846 | 6epc-H | 2.7 | 10.8 | 58 | 396 | 3 | MOLECULE: PROTEASOME SUBUNIT ALPHA TYPE-6; |
| 847 | 4i5s-B | 2.7 | 7.4 | 61 | 395 | 5 | MOLECULE: PUTATIVE HISTIDINE KINASE COVS; VICK-LIKE PROTEIN |
| 848 | 2k3n-A | 2.7 | 3.9 | 76 | 160 | 8 | MOLECULE: TUSP1; |
| 849 | 4qlb-D | 2.7 | 5.1 | 59 | 651 | 10 | MOLECULE: PROBABLE GLYCOGEN [STARCH] SYNTHASE; |
| 850 | 5nug-A | 2.7 | 6 | 70 | 2920 | 10 | MOLECULE: CYTOPLASMIC DYNEIN 1 HEAVY CHAIN 1; |
| 851 | 7asl-A | 2.7 | 3.7 | 85 | 235 | 8 | MOLECULE: GAG PROTEIN; |
| 852 | 6vek-A | 2.7 | 9.8 | 72 | 321 | 7 | MOLECULE: CONTACT-DEPENDENT TOXIN CDIA; |
| 853 | 6i3m-E | 2.7 | 4.4 | 51 | 354 | 10 | MOLECULE: TRANSLATION INITIATION FACTOR EIF-2B SUBUNIT ALPH |
| 854 | 4zy8-A | 2.7 | 3.5 | 49 | 146 | 6 | MOLECULE: PROTEIN LST4; |
| 855 | 6qrj-A | 2.7 | 14.5 | 55 | 463 | 4 | MOLECULE: HYBRID KINASE; |
| 856 | 6vgc-D | 2.7 | 8 | 58 | 155 | 9 | MOLECULE: 5-LIPOXYGENASE-ACTIVATING PROTEIN; |
| 857 | 7jtk-M | 2.7 | 2.1 | 36 | 47 | 11 | MOLECULE: FLAGELLAR RADIAL SPOKE PROTEIN 1; |
| 858 | 7njn-C | 2.7 | 12.7 | 73 | 536 | 10 | MOLECULE: ATP SYNTHASE SUBUNIT ALPHA; |
| 859 | 5xqi-B | 2.7 | 8.7 | 78 | 276 | 3 | MOLECULE: PROTEIN ROGDI HOMOLOG; |
| 860 | 5v5t-A | 2.7 | 7 | 77 | 306 | 8 | MOLECULE: CONSERVED DOMAIN PROTEIN; |
| 861 | 6zyx-C | 2.7 | 9.5 | 48 | 278 | 6 | MOLECULE: DYNEIN HEAVY CHAIN, OUTER ARM PROTEIN; |
| 862 | 2i7u-A | 2.7 | 3.5 | 42 | 62 | 5 | MOLECULE: FOUR-ALPHA-HELIX BUNDLE; |
| 863 | 1cm5-A | 2.7 | 3.3 | 59 | 759 | 5 | MOLECULE: PROTEIN (PYRUVATE FORMATE-LYASE); |
| 864 | 5mnt-B | 2.7 | 22.1 | 80 | 421 | 8 | MOLECULE: A2 MATURATION PROTEIN; |
| 865 | 6ys8-C | 2.7 | 5.5 | 43 | 60 | 7 | MOLECULE: GLDM; |
| 866 | 1oks-A | 2.7 | 3.2 | 45 | 53 | 13 | MOLECULE: RNA POLYMERASE ALPHA SUBUNIT; |
| 867 | 6qm5-A | 2.7 | 10.1 | 81 | 673 | 5 | MOLECULE: PREDICTED PROTEIN; |
| 868 | 6cpu-A | 2.7 | 7.5 | 89 | 526 | 4 | MOLECULE: PHOSPHODIESTERASE; |
| 869 | 4xmn-E | 2.7 | 12.8 | 96 | 896 | 4 | MOLECULE: PROTEIN TRANSPORT PROTEIN SEC13; |
| 870 | 3sjr-A | 2.7 | 3.7 | 54 | 126 | 4 | MOLECULE: UNCHARACTERIZED PROTEIN; |
| 871 | 7ah0-A | 2.7 | 6.1 | 44 | 104 | 2 | MOLECULE: 4D2; |
| 872 | 2qsr-A | 2.7 | 3.5 | 53 | 161 | 9 | MOLECULE: TRANSCRIPTION-REPAIR COUPLING FACTOR; |

| | | | | | | | |
|---|---|---|---|---|---|---|---|
| 873 | 4uis-B | 2.7 | 9.7 | 75 | 241 | 3 | MOLECULE: GAMMA-SECRETASE; |
| 874 | 5o4j-A | 2.7 | 7.6 | 63 | 265 | 5 | MOLECULE: HCGC; |
| 875 | 4ql6-B | 2.7 | 3.2 | 60 | 509 | 10 | MOLECULE: CARBOXY-TERMINAL PROCESSING PROTEASE; |
| 876 | 7d3u-E | 2.7 | 4.9 | 52 | 110 | 8 | MOLECULE: MONOVALENT NA+/H+ ANTIPORTER SUBUNIT D; |
| 877 | 6v4m-A | 2.7 | 7.2 | 66 | 163 | 2 | MOLECULE: BCL-2; |
| 878 | 1r71-B | 2.7 | 3.5 | 44 | 116 | 14 | MOLECULE: 5'-D(*AP*(BRU) |
| 879 | 5e6g-B | 2.7 | 3.5 | 49 | 114 | 12 | MOLECULE: DE NOVO DESIGNED PROTEIN CA01; |
| 880 | 6jly-G | 2.7 | 6.1 | 66 | 363 | 6 | MOLECULE: TRANSLATION INITIATION FACTOR EIF-2B SUBUNIT ALPH |
| 881 | 5vp3-A | 2.7 | 3.9 | 58 | 195 | 7 | MOLECULE: MNEMIOPSIN 1; |
| 882 | 6njw-A | 2.7 | 4.1 | 66 | 208 | 9 | MOLECULE: XCC_CTR_PT; |
| 883 | 1q16-A | 2.7 | 5.7 | 83 | 1244 | 5 | MOLECULE: RESPIRATORY NITRATE REDUCTASE 1 ALPHA CHAIN; |
| 884 | 6w1h-A | 2.7 | 11.8 | 78 | 450 | 5 | MOLECULE: HYDROXYGLUTARATE SYNTHASE; |
| 885 | 3sqn-A | 2.7 | 6.8 | 75 | 468 | 11 | MOLECULE: CONSERVED DOMAIN PROTEIN; |
| 886 | 6bs9-A | 2.7 | 4.2 | 49 | 130 | 6 | MOLECULE: STAGE III SPORULATION PROTEIN AB; |
| 887 | 6rql-A | 2.7 | 9 | 86 | 1542 | 9 | MOLECULE: TEMPLATE STRAND; |
| 888 | 4ysx-G | 2.7 | 14 | 77 | 155 | 8 | MOLECULE: SUCCINATE DEHYDROGENASE FLAVOPROTEIN; |
| 889 | 2jvw-A | 2.7 | 4 | 62 | 82 | 10 | MOLECULE: UNCHARACTERIZED PROTEIN; |
| 890 | 5w10-A | 2.7 | 4.3 | 44 | 173 | 9 | MOLECULE: CGMP-SPECIFIC PHOSPHODIESTERASE; |
| 891 | 3k29-A | 2.7 | 6.3 | 55 | 162 | 15 | MOLECULE: PUTATIVE UNCHARACTERIZED PROTEIN; |
| 892 | 5ipx-A | 2.7 | 8.3 | 84 | 282 | 7 | MOLECULE: ORF49 PROTEIN; |
| 893 | 7cm9-A | 2.7 | 9.8 | 85 | 693 | 12 | MOLECULE: DMSP LYASE; |
| 894 | 4yo5-A | 2.7 | 9.4 | 52 | 131 | 4 | MOLECULE: TSSA; |
| 895 | 7oa5-B | 2.7 | 11.1 | 52 | 186 | 6 | MOLECULE: HOLLIDAY JUNCTION ATP-DEPENDENT DNA HELICASE RUVA |
| 896 | 6pw7-A | 2.7 | 10.1 | 62 | 161 | 6 | MOLECULE: STROMAL INTERACTION MOLECULE 1; |
| 897 | 3tvi-D | 2.7 | 16.3 | 77 | 439 | 3 | MOLECULE: ASPARTOKINASE; |
| 898 | 5cqq-B | 2.7 | 4.1 | 51 | 75 | 12 | MOLECULE: REGULATORY PROTEIN ZESTE; |
| 899 | 5tub-A | 2.7 | 4.3 | 81 | 335 | 4 | MOLECULE: SHARK TBC1D15 GTPASE-ACTIVATING PROTEIN; |
| 900 | 6iuy-A | 2.7 | 10.6 | 87 | 585 | 5 | MOLECULE: GLYCEROL-3-PHOSPHATE DEHYDROGENASE [NAD(+)]; |
| 901 | 7dms-A | 2.7 | 3.5 | 34 | 94 | 18 | MOLECULE: FE(II)-BINDING EFFECTOR; |
| 902 | 4hb1-A | 2.7 | 4.6 | 39 | 44 | 5 | MOLECULE: DHP1; |
| 903 | 4uig-A | 2.7 | 8.3 | 55 | 91 | 11 | MOLECULE: COPPER SENSITIVE OPERON REPRESSOR; |
| 904 | 6k2e-A | 2.7 | 7.1 | 49 | 88 | 14 | MOLECULE: CRISPR/CAS2 PROTEIN; |
| 905 | 7mtl-B | 2.7 | 3.5 | 76 | 162 | 8 | MOLECULE: COLIBACTIN SELF-PROTECTION PROTEIN CLBS; |
| 906 | 6qzh-A | 2.7 | 5.4 | 77 | 744 | 10 | MOLECULE: HUMAN CHEMOKINE RECEPTOR 7; |
| 907 | 6n0v-A | 2.7 | 10.3 | 85 | 395 | 7 | MOLECULE: TRNA LIGASE; |

| | | | | | | | |
|---|---|---|---|---|---|---|---|
| 908 | 3mnl-A | 2.7 | 12.2 | 75 | 188 | 8 | MOLECULE: TRANSCRIPTIONAL REGULATORY PROTEIN (PROBABLY TETR |
| 909 | 4geh-D | 2.7 | 5.2 | 51 | 69 | 0 | MOLECULE: PROGRAMMED CELL DEATH PROTEIN 10; |
| 910 | 6hqa-A | 2.6 | 6.3 | 97 | 887 | 9 | MOLECULE: TAF2; |
| 911 | 6mcp-A | 2.6 | 5.5 | 44 | 520 | 7 | MOLECULE: LEGK7; |
| 912 | 4axd-A | 2.6 | 9.4 | 76 | 433 | 1 | MOLECULE: INOSITOL-PENTAKISPHOSPHATE 2-KINASE; |
| 913 | 5v7p-A | 2.6 | 10.4 | 69 | 281 | 6 | MOLECULE: PROTEIN-S-ISOPRENYLCYSTEINE O-METHYLTRANSFERASE; |
| 914 | 4ney-B | 2.6 | 12.1 | 68 | 173 | 10 | MOLECULE: ENGINEERED PROTEIN OR277; |
| 915 | 5ea1-B | 2.6 | 3.2 | 64 | 128 | 11 | MOLECULE: TRANSCRIPTION ACTIVATOR BRG1; |
| 916 | 6z8h-A | 2.6 | 9.3 | 71 | 351 | 8 | MOLECULE: VARIANT SURFACE GLYCOPROTEIN MITAT 1.13; |
| 917 | 4atv-A | 2.6 | 4.8 | 83 | 381 | 5 | MOLECULE: NA(+)/H(+) ANTIPORTER NHAA; |
| 918 | 5y06-A | 2.6 | 7.3 | 60 | 235 | 8 | MOLECULE: MSMEG_4306; |
| 919 | 1otr-A | 2.6 | 2.8 | 42 | 49 | 10 | MOLECULE: PROTEIN CUE2; |
| 920 | 6wti-D | 2.6 | 7.8 | 55 | 99 | 9 | MOLECULE: CYTOCHROME O UBIQUINOL OXIDASE, SUBUNIT I; |
| 921 | 6jbh-C | 2.6 | 15.9 | 87 | 267 | 6 | MOLECULE: TARH; |
| 922 | 6c0f-w | 2.6 | 2.7 | 40 | 70 | 10 | MOLECULE: SACCHAROMYCES CEREVISIAE S288C 35S PRE-RIBOSOMAL |
| 923 | 3a8t-A | 2.6 | 6.3 | 56 | 289 | 7 | MOLECULE: ADENYLATE ISOPENTENYLTRANSFERASE; |
| 924 | 7b9c-A | 2.6 | 9.2 | 94 | 870 | 4 | MOLECULE: SPLICING FACTOR 3B SUBUNIT 3; |
| 925 | 6ff6-A | 2.6 | 8.6 | 86 | 222 | 10 | MOLECULE: BRIC1; |
| 926 | 1cii-A | 2.6 | 2.2 | 54 | 602 | 9 | MOLECULE: COLICIN IA; |
| 927 | 3iuk-A | 2.6 | 7.4 | 71 | 552 | 1 | MOLECULE: UNCHARACTERIZED PROTEIN; |
| 928 | 2ld6-A | 2.6 | 3.8 | 58 | 133 | 5 | MOLECULE: CHEMOTAXIS PROTEIN CHEA; |
| 929 | 4yy2-A | 2.6 | 4.1 | 56 | 99 | 9 | MOLECULE: DTOR_3X33L; |
| 930 | 3jcf-A | 2.6 | 3.1 | 40 | 349 | 5 | MOLECULE: MAGNESIUM TRANSPORT PROTEIN CORA; |
| 931 | 6q6g-O | 2.6 | 5.3 | 84 | 703 | 6 | MOLECULE: CELL DIVISION CYCLE PROTEIN 20 HOMOLOG; |
| 932 | 3dl8-E | 2.6 | 6.2 | 42 | 65 | 7 | MOLECULE: PROTEIN TRANSLOCASE SUBUNIT SECA; |
| 933 | 5hzd-A | 2.6 | 5.3 | 84 | 476 | 10 | MOLECULE: 3' TERMINAL URIDYLYL TRANSFERASE; |
| 934 | 2f4q-A | 2.6 | 6.3 | 61 | 309 | 5 | MOLECULE: TYPE I TOPOISOMERASE, PUTATIVE; |
| 935 | 6wql-A | 2.6 | 4.5 | 44 | 49 | 7 | MOLECULE: SEED PEPTIDE C2 (VBP-1); |
| 936 | 5a5t-M | 2.6 | 13.7 | 74 | 365 | 5 | MOLECULE: EUKARYOTIC TRANSLATION INITIATION FACTOR 3 SUBUNI |
| 937 | 6zui-A | 2.6 | 12.1 | 70 | 420 | 3 | MOLECULE: HTH-TYPE TRANSCRIPTIONAL REPRESSOR NEMR,GREEN FLU |
| 938 | 6ds9-A | 2.6 | 8.3 | 52 | 93 | 2 | MOLECULE: DE NOVO DESIGNED THREE HELIX BUNDLE GRA3D; |
| 939 | 6xl7-A | 2.6 | 4.1 | 61 | 119 | 13 | MOLECULE: SG7.AF; |
| 940 | 3un9-A | 2.6 | 9.9 | 76 | 294 | 8 | MOLECULE: NLR FAMILY MEMBER X1; |
| 941 | 3let-B | 2.6 | 10.9 | 83 | 310 | 5 | MOLECULE: ADENOSINE MONOPHOSPHATE-PROTEIN TRANSFERASE VOPS; |
| 942 | 6thh-C | 2.6 | 6.5 | 69 | 816 | 12 | MOLECULE: SIRV2 ACRID1 (GP02) ANTI-CRISPR PROTEIN; |

| | | | | | | | |
|---|---|---|---|---|---|---|---|
| 943 | 3hkl-B | 2.6 | 5.5 | 56 | 153 | 4 | MOLECULE: MUSCLE, SKELETAL RECEPTOR TYROSINE PROTEIN KINASE |
| 944 | 3pla-A | 2.6 | 10.5 | 68 | 375 | 7 | MOLECULE: PRE MRNA SPLICING PROTEIN; |
| 945 | 5td6-B | 2.6 | 3.4 | 45 | 136 | 4 | MOLECULE: FOG-3 PROTEIN; |
| 946 | 6ka4-A | 2.6 | 5.9 | 66 | 534 | 6 | MOLECULE: F22L4.1 PROTEIN; |
| 947 | 4m70-L | 2.6 | 4 | 56 | 112 | 2 | MOLECULE: RX PROTEIN; |
| 948 | 7a6u-A | 2.6 | 7.2 | 82 | 332 | 11 | MOLECULE: SHORT TRANSIENT RECEPTOR POTENTIAL CHANNEL 6; |
| 949 | 1zw0-D | 2.6 | 4.7 | 38 | 66 | 5 | MOLECULE: TYPE III SECRETION PROTEIN; |
| 950 | 5xah-C | 2.6 | 6.9 | 55 | 376 | 7 | MOLECULE: IMPORTIN-4; |
| 951 | 5kbc-A | 2.6 | 5.5 | 63 | 195 | 5 | MOLECULE: DSBA; |
| 952 | 1gqe-A | 2.6 | 7.9 | 83 | 362 | 11 | MOLECULE: RELEASE FACTOR 2; |
| 953 | 6cjd-A | 2.6 | 10.8 | 58 | 128 | 14 | MOLECULE: PUTATIVE CYTOPLASMIC PROTEIN; |
| 954 | 5orf-A | 2.6 | 10.6 | 87 | 586 | 3 | MOLECULE: SERUM ALBUMIN; |
| 955 | 2p7n-A | 2.6 | 3.6 | 75 | 335 | 4 | MOLECULE: PATHOGENICITY ISLAND 1 EFFECTOR PROTEIN; |
| 956 | 6wqj-A | 2.6 | 2.1 | 32 | 35 | 3 | MOLECULE: VICILIN-BURIED PEPTIDE-10; |
| 957 | 4dci-B | 2.6 | 5.4 | 60 | 148 | 5 | MOLECULE: UNCHARACTERIZED PROTEIN; |
| 958 | 3lqi-C | 2.6 | 3.6 | 66 | 181 | 12 | MOLECULE: MLL1 PHD3-BROMO; |
| 959 | 6n5x-A | 2.6 | 8.5 | 57 | 168 | 7 | MOLECULE: SORTING NEXIN-5,CATION-INDEPENDENT MANNOSE-6-PHOS |
| 960 | 5mmj-c | 2.6 | 7.9 | 51 | 216 | 6 | MOLECULE: 50S RIBOSOMAL PROTEIN L31; |
| 961 | 3jbr-F | 2.6 | 9.9 | 71 | 872 | 7 | MOLECULE: VOLTAGE-DEPENDENT L-TYPE CALCIUM CHANNEL SUBUNIT |
| 962 | 5iy6-U | 2.6 | 12.1 | 85 | 170 | 5 | MOLECULE: DNA-DIRECTED RNA POLYMERASE II SUBUNIT RPB1; |
| 963 | 7jw1-e | 2.6 | 6.4 | 51 | 235 | 2 | MOLECULE: CAPSID PROTEINS; |
| 964 | 6ey4-A | 2.6 | 8.6 | 92 | 483 | 10 | MOLECULE: GLDM; |
| 965 | 6wq2-a | 2.6 | 8.2 | 63 | 202 | 8 | MOLECULE: A-DNA; |
| 966 | 2guz-A | 2.6 | 5.7 | 47 | 71 | 13 | MOLECULE: MITOCHONDRIAL IMPORT INNER MEMBRANE TRANSLOCASE S |
| 967 | 6wej-A | 2.6 | 14.9 | 76 | 513 | 4 | MOLECULE: CYCLIC NUCLEOTIDE-GATED CATION CHANNEL; |
| 968 | 1zbp-A | 2.6 | 9.3 | 60 | 266 | 7 | MOLECULE: HYPOTHETICAL PROTEIN VPA1032; |
| 969 | 5xki-B | 2.6 | 7 | 70 | 259 | 7 | MOLECULE: FAD-LINKED SULFHYDRYL OXIDASE; |
| 970 | 4qvr-A | 2.6 | 4.5 | 65 | 318 | 11 | MOLECULE: UNCHARACTERIZED HYPOTHETICAL PROTEIN FTT_1539C; |
| 971 | 1g8p-A | 2.6 | 4.3 | 71 | 321 | 8 | MOLECULE: MAGNESIUM-CHELATASE 38 KDA SUBUNIT; |
| 972 | 7cma-A | 2.6 | 3.7 | 61 | 155 | 10 | MOLECULE: A151R; |
| 973 | 1z0j-B | 2.6 | 3.8 | 40 | 51 | 3 | MOLECULE: RAS-RELATED PROTEIN RAB-22A; |
| 974 | 7emf-H | 2.6 | 9.5 | 58 | 181 | 14 | MOLECULE: MEDIATOR OF RNA POLYMERASE II TRANSCRIPTION SUBUN |
| 975 | 2xco-A | 2.6 | 14.4 | 66 | 636 | 8 | MOLECULE: DNA GYRASE SUBUNIT B, DNA GYRASE SUBUNIT A; |
| 976 | 6l85-B | 2.6 | 9.9 | 64 | 401 | 3 | MOLECULE: PHOSPHATE TRANSPORTER; |
| 977 | 2h1n-A | 2.6 | 13.6 | 75 | 566 | 9 | MOLECULE: OLIGOENDOPEPTIDASE F; |

| | | | | | | | |
|---|---|---|---|---|---|---|---|
| 978 | 6spb-Y | 2.6 | 5.7 | 42 | 60 | 12 | MOLECULE: 23S RIBOSOMAL RNA; |
| 979 | 6kko-A | 2.6 | 5.4 | 64 | 170 | 5 | MOLECULE: PUTATIVE SERINE PHOSPHATASE; |
| 980 | 4clv-A | 2.6 | 8.5 | 70 | 142 | 6 | MOLECULE: NICKEL-COBALT-CADMIUM RESISTANCE PROTEIN NCCX; |
| 981 | 3a6m-A | 2.6 | 6.5 | 50 | 168 | 6 | MOLECULE: PROTEIN GRPE; |
| 982 | 4an8-A | 2.6 | 10.5 | 89 | 468 | 4 | MOLECULE: CSE1; |
| 983 | 3tsy-A | 2.6 | 8.9 | 71 | 828 | 3 | MOLECULE: FUSION PROTEIN 4-COUMARATE--COA LIGASE 1, RESVERA |
| 984 | 4zi2-C | 2.6 | 5.9 | 55 | 131 | 11 | MOLECULE: ADP-RIBOSYLATION FACTOR-LIKE PROTEIN 3; |
| 985 | 6ww2-R | 2.6 | 4.3 | 63 | 425 | 3 | MOLECULE: ANTI-BRIL FAB HEAVY CHAIN; |
| 986 | 5w99-A | 2.6 | 6.2 | 58 | 319 | 9 | MOLECULE: PBTD; |
| 987 | 1ghe-B | 2.6 | 3.2 | 59 | 171 | 12 | MOLECULE: ACETYLTRANSFERASE; |
| 988 | 5ld2-D | 2.6 | 8.4 | 66 | 605 | 3 | MOLECULE: RECBCD ENZYME SUBUNIT RECB,RECBCD ENZYME SUBUNIT |
| 989 | 6n8a-B | 2.6 | 3.5 | 75 | 194 | 9 | MOLECULE: TRANSCRIPTION REGULATOR ACAB; |
| 990 | 2mqk-A | 2.6 | 2.7 | 45 | 65 | 11 | MOLECULE: ATP-DEPENDENT TARGET DNA ACTIVATOR B; |
| 991 | 6ybv-s | 2.6 | 4.1 | 66 | 138 | 8 | MOLECULE: EUKARYOTIC TRANSLATION INITIATION FACTOR 2 SUBUNI |
| 992 | 7apk-j | 2.6 | 14.2 | 58 | 707 | 16 | MOLECULE: THO COMPLEX SUBUNIT 1; |
| 993 | 2nr5-E | 2.6 | 4.4 | 43 | 65 | 12 | MOLECULE: HYPOTHETICAL PROTEIN SO2669; |
| 994 | 7etw-B | 2.6 | 6.6 | 85 | 483 | 7 | MOLECULE: INSULIN-INDUCED GENE 2 PROTEIN; |
| 995 | 6lqz-A | 2.6 | 3 | 49 | 71 | 10 | MOLECULE: TRANSCRIPTION INITIATION FACTOR TFIID SUBUNIT 14; |
| 996 | 3fd9-A | 2.6 | 12.7 | 80 | 240 | 6 | MOLECULE: UNCHARACTERIZED PROTEIN; |
| 997 | 6njp-G | 2.6 | 5.1 | 43 | 64 | 12 | MOLECULE: TRANSLOCATOR ESCN; |
| 998 | 6xxv-C | 2.6 | 2.5 | 41 | 111 | 12 | MOLECULE: ANTIBODY C57, HEAVY CHAIN; |
| 999 | 6qum-N | 2.6 | 8 | 70 | 649 | 7 | MOLECULE: V-TYPE ATP SYNTHASE ALPHA CHAIN; |
| 1000 | 3nl9-A | 2.6 | 9.1 | 44 | 169 | 11 | MOLECULE: PUTATIVE NTP PYROPHOSPHOHYDROLASE; |
| 1001 | 4rkm-B | 2.6 | 5.2 | 62 | 660 | 10 | MOLECULE: MCCA; |
| 1002 | 3etz-B | 2.6 | 5.2 | 47 | 115 | 11 | MOLECULE: ADHESIN A; |
| 1003 | 2zoz-B | 2.6 | 12.7 | 74 | 181 | 4 | MOLECULE: TRANSCRIPTIONAL REGULATOR; |
| 1004 | 5uac-C | 2.6 | 9.8 | 65 | 1342 | 8 | MOLECULE: DNA-DIRECTED RNA POLYMERASE SUBUNIT ALPHA; |
| 1005 | 5w65-P | 2.6 | 12.3 | 94 | 481 | 6 | MOLECULE: DNA-DIRECTED RNA POLYMERASE I SUBUNIT RPA190; |
| 1006 | 1au1-A | 2.6 | 3.9 | 48 | 166 | 8 | MOLECULE: INTERFERON-BETA; |
| 1007 | 2mx9-B | 2.6 | 4.5 | 71 | 133 | 3 | MOLECULE: MINOR AMPULLATE SPIDROIN; |
| 1008 | 7bin-V | 2.6 | 4.6 | 48 | 133 | 13 | MOLECULE: FLAGELLAR BIOSYNTHETIC PROTEIN FLIP; |
| 1009 | 5n9j-G | 2.6 | 6.9 | 54 | 163 | 15 | MOLECULE: MEDIATOR OF RNA POLYMERASE II TRANSCRIPTION SUBUN |
| 1010 | 5g4y-A | 2.6 | 3.9 | 58 | 147 | 7 | MOLECULE: CHEMOTAXIS PROTEIN; |
| 1011 | 6rxd-A | 2.6 | 10.1 | 76 | 509 | 5 | MOLECULE: HISTIDINE ACID PHOSPHATASE; |
| 1012 | 6okd-C | 2.6 | 3.1 | 36 | 43 | 11 | MOLECULE: TRANSFERRIN RECEPTOR PROTEIN 1; |

| | | | | | | | |
|---|---|---|---|---|---|---|---|
| 1013 | 3otv-C | 2.6 | 12.1 | 67 | 257 | 6 | MOLECULE: PROBABLE CONSERVED TRANSMEMBRANE PROTEIN; |
| 1014 | 1yz7-A | 2.6 | 9 | 66 | 176 | 9 | MOLECULE: PROBABLE TRANSLATION INITIATION FACTOR 2 ALPHA |
| 1015 | 2d32-A | 2.6 | 4.5 | 61 | 512 | 7 | MOLECULE: GLUTAMATE--CYSTEINE LIGASE; |
| 1016 | 5dgk-A | 2.6 | 7.4 | 84 | 519 | 6 | MOLECULE: ACTIVE HELICASE; |
| 1017 | 7cbc-A | 2.6 | 7.7 | 71 | 319 | 6 | MOLECULE: DE NOVO DESIGNED SWITCH PROTEIN CAGING A HEMAGGLU |
| 1018 | 3lph-A | 2.6 | 4.1 | 42 | 62 | 17 | MOLECULE: PROTEIN REV; |
| 1019 | 5eqw-B | 2.6 | 3.8 | 59 | 123 | 8 | MOLECULE: PUTATIVE MAJOR COAT PROTEIN; |
| 1020 | 5ul2-A | 2.6 | 11.7 | 90 | 731 | 4 | MOLECULE: OXSB PROTEIN; |
| 1021 | 2vxo-B | 2.6 | 11.9 | 57 | 658 | 7 | MOLECULE: GMP SYNTHASE [GLUTAMINE-HYDROLYZING]; |
| 1022 | 6gv8-A | 2.6 | 8.7 | 69 | 157 | 12 | MOLECULE: HYPEROSMOLARITY RESISTANCE PROTEIN EMB; |
| 1023 | 7ppo-A | 2.6 | 6 | 76 | 500 | 12 | MOLECULE: UBIQUITINATING/DEUBIQUITINATING ENZYME SDEA; |
| 1024 | 5zk4-C | 2.6 | 3.3 | 47 | 73 | 15 | MOLECULE: DISD PROTEIN; |
| 1025 | 2axt-Z | 2.6 | 5.4 | 41 | 62 | 15 | MOLECULE: PHOTOSYSTEM Q(B) PROTEIN; |
| 1026 | 2mqa-A | 2.6 | 3 | 65 | 125 | 8 | MOLECULE: MINOR AMPULLATE FIBROIN 1; |
| 1027 | 5zr1-B | 2.6 | 11.3 | 68 | 374 | 4 | MOLECULE: ORIGIN RECOGNITION COMPLEX SUBUNIT 1; |
| 1028 | 7ey7-A | 2.6 | 3.7 | 48 | 102 | 8 | MOLECULE: TAIL FIBER PROTEIN; |
| 1029 | 6qv7-A | 2.6 | 12.6 | 72 | 316 | 8 | MOLECULE: UNCHARACTERIZED PROTEIN; |
| 1030 | 6g7c-C | 2.6 | 12.7 | 79 | 239 | 11 | MOLECULE: IMPA-RELATED DOMAIN PROTEIN; |
| 1031 | 6ezn-H | 2.6 | 8.8 | 54 | 259 | 7 | MOLECULE: DOLICHYL-DIPHOSPHOOLIGOSACCHARIDE--PROTEIN |
| 1032 | 4jza-A | 2.6 | 14.6 | 104 | 817 | 8 | MOLECULE: UNCHARACTERIZED PROTEIN; |
| 1033 | 7nj0-C | 2.6 | 4.6 | 66 | 270 | 3 | MOLECULE: SECURIN,SEPARIN; |
| 1034 | 3pyb-B | 2.6 | 4.9 | 80 | 689 | 10 | MOLECULE: ENT-COPALYL DIPHOSPHATE SYNTHASE, CHLOROPLASTIC; |
| 1035 | 7lu4-A | 2.6 | 12.1 | 63 | 978 | 6 | MOLECULE: MULTIFUNCTIONAL FUSION PROTEIN; |
| 1036 | 4q9u-A | 2.6 | 3.8 | 62 | 303 | 3 | MOLECULE: RAB5 GDP/GTP EXCHANGE FACTOR; |
| 1037 | 7cxt-A | 2.6 | 6.2 | 78 | 348 | 5 | MOLECULE: GDP-L-FUCOSE SYNTHASE; |
| 1038 | 5l1a-A | 2.6 | 4.1 | 53 | 108 | 9 | MOLECULE: UNCHARACTERIZED PROTEIN; |
| 1039 | 5yti-A | 2.6 | 18.3 | 69 | 330 | 9 | MOLECULE: FLAGELLAR HOOK ASSOCIATED PROTEIN TYPE 3 FLGL; |
| 1040 | 5yhf-A | 2.6 | 11.3 | 102 | 734 | 6 | MOLECULE: PROTEIN TRANSLOCASE SUBUNIT SECDF; |
| 1041 | 6zls-A | 2.6 | 5.1 | 61 | 292 | 5 | MOLECULE: HISTIDINE KINASE; |
| 1042 | 7ctq-A | 2.6 | 12.5 | 72 | 429 | 7 | MOLECULE: PEPTIDYL TRYPTOPHAN DIHYDROXYLASE; |
| 1043 | 2qzg-B | 2.6 | 3.1 | 56 | 90 | 13 | MOLECULE: CONSERVED UNCHARACTERIZED ARCHAEAL PROTEIN; |
| 1044 | 4l0r-A | 2.6 | 5.3 | 49 | 74 | 6 | MOLECULE: CENTROSOMAL PROTEIN OF 57 KDA; |
| 1045 | 2rgo-A | 2.6 | 5.4 | 64 | 557 | 11 | MOLECULE: ALPHA-GLYCEROPHOSPHATE OXIDASE; |
| 1046 | 7a23-J | 2.6 | 10.6 | 55 | 96 | 2 | MOLECULE: 51KDA; |
| 1047 | 1abz-A | 2.6 | 2.7 | 37 | 38 | 5 | MOLECULE: ALPHA-T-ALPHA; |

| | | | | | | | |
|---|---|---|---|---|---|---|---|
| 1048 | 4kyz-A | 2.6 | 12.5 | 68 | 167 | 10 | MOLECULE: DESIGNED PROTEIN OR327; |
| 1049 | 4xng-D | 2.6 | 10.2 | 65 | 144 | 6 | MOLECULE: UNCHARACTERIZED PROTEIN MG218.1; |
| 1050 | 6tpq-B | 2.6 | 10.2 | 70 | 183 | 10 | MOLECULE: RIBONUCLEASE M5; |
| 1051 | 2keb-A | 2.6 | 3.6 | 53 | 78 | 11 | MOLECULE: DNA POLYMERASE SUBUNIT ALPHA B; |
| 1052 | 3m1c-B | 2.6 | 7.5 | 61 | 146 | 10 | MOLECULE: ENVELOPE GLYCOPROTEIN H; |
| 1053 | 2ve7-B | 2.6 | 12.9 | 70 | 303 | 6 | MOLECULE: KINETOCHORE PROTEIN HEC1, KINETOCHORE PROTEIN SPC |
| 1054 | 6vbu-2 | 2.6 | 16.5 | 69 | 659 | 12 | MOLECULE: BARDET-BIEDL SYNDROME 18 PROTEIN; |
| 1055 | 4c3b-C | 2.6 | 4.3 | 69 | 178 | 12 | MOLECULE: MATRIX PROTEIN 2-1; |
| 1056 | 3c4a-A | 2.6 | 2.8 | 45 | 365 | 11 | MOLECULE: PROBABLE TRYPTOPHAN HYDROXYLASE VIOD; |
| 1057 | 2uwj-E | 2.6 | 5.2 | 48 | 70 | 8 | MOLECULE: TYPE III EXPORT PROTEIN PSCE; |
| 1058 | 4nv0-B | 2.6 | 11.9 | 73 | 300 | 4 | MOLECULE: 7-METHYLGUANOSINE PHOSPHATE-SPECIFIC 5'-NUCLEOTID |
| 1059 | 2cq8-A | 2.6 | 2.8 | 54 | 110 | 13 | MOLECULE: 10-FORMYLTETRAHYDROFOLATE DEHYDROGENASE; |
| 1060 | 6psk-R | 2.6 | 2.7 | 47 | 72 | 2 | MOLECULE: ANTIHOLIN; |
| 1061 | 3e7q-B | 2.6 | 11 | 66 | 210 | 8 | MOLECULE: TRANSCRIPTIONAL REGULATOR; |
| 1062 | 1jud-A | 2.6 | 18.6 | 71 | 220 | 11 | MOLECULE: L-2-HALOACID DEHALOGENASE; |
| 1063 | 6dkm-A | 2.6 | 5.1 | 42 | 77 | 21 | MOLECULE: DHD131_A; |
| 1064 | 3me5-A | 2.6 | 5.6 | 70 | 414 | 7 | MOLECULE: CYTOSINE-SPECIFIC METHYLTRANSFERASE; |
| 1065 | 6o3s-A | 2.6 | 3.7 | 37 | 47 | 8 | MOLECULE: RIBOSOME-INACTIVATING PROTEIN LUFFIN P1; |
| 1066 | 1i49-A | 2.6 | 8.3 | 60 | 201 | 8 | MOLECULE: ARFAPTIN 2; |
| 1067 | 4p6v-E | 2.6 | 3.1 | 45 | 189 | 9 | MOLECULE: NA(+)-TRANSLOCATING NADH-QUINONE REDUCTASE SUBUNI |
| 1068 | 3ang-C | 2.6 | 14.7 | 75 | 201 | 8 | MOLECULE: TRANSCRIPTIONAL REPRESSOR, TETR FAMILY; |
| 1069 | 2oo2-A | 2.6 | 4.7 | 52 | 76 | 4 | MOLECULE: HYPOTHETICAL PROTEIN AF_1782; |
| 1070 | 3few-X | 2.6 | 10.3 | 59 | 431 | 2 | MOLECULE: COLICIN S4; |
| 1071 | 5a60-A | 2.6 | 10.1 | 104 | 429 | 12 | MOLECULE: INORGANIC TRIPHOSPHATASE; |
| 1072 | 3bk6-A | 2.6 | 3.1 | 52 | 170 | 8 | MOLECULE: PH STOMATIN; |
| 1073 | 3wxx-B | 2.6 | 7.5 | 78 | 183 | 10 | MOLECULE: ACRH; |
| 1074 | 3jd5-h | 2.6 | 6.9 | 73 | 103 | 8 | MOLECULE: 28S RIBOSOMAL RNA, MITOCHONDIAL; |
| 1075 | 6cnb-R | 2.6 | 10.7 | 71 | 522 | 11 | MOLECULE: DNA-DIRECTED RNA POLYMERASE III SUBUNIT RPC1; |
| 1076 | 6dxx-A | 2.5 | 3.7 | 47 | 97 | 9 | MOLECULE: N-ACYLETHANOLAMINE-HYDROLYZING ACID AMIDASE SUBUN |
| 1077 | 2mx7-A | 2.5 | 2.9 | 52 | 112 | 10 | MOLECULE: SYNERGIN GAMMA; |
| 1078 | 6awl-A | 2.5 | 11.5 | 88 | 217 | 11 | MOLECULE: UBIQUINONE BIOSYNTHESIS PROTEIN COQ9, MITOCHONDRI |
| 1079 | 6qm7-K | 2.5 | 6.2 | 66 | 206 | 5 | MOLECULE: PROTEASOME ALPHA1 CHAIN; |
| 1080 | 4uet-A | 2.5 | 7.4 | 69 | 170 | 6 | MOLECULE: NEMATODE FATTY ACID RETINOID BINDING PROTEIN; |
| 1081 | 4cid-A | 2.5 | 3.8 | 61 | 497 | 7 | MOLECULE: EH DOMAIN-CONTAINING PROTEIN 2; |
| 1082 | 6nd4-L | 2.5 | 17.6 | 67 | 473 | 7 | MOLECULE: ETS RRNA; |

| | | | | | | | |
|---|---|---|---|---|---|---|---|
| 1083 | 3pwx-A | 2.5 | 4.7 | 47 | 224 | 6 | MOLECULE: PUTATIVE FLAGELLAR HOOK-ASSOCIATED PROTEIN; |
| 1084 | 6dkm-D | 2.5 | 4.6 | 42 | 79 | 0 | MOLECULE: DHD131_A; |
| 1085 | 1qoy-A | 2.5 | 13.5 | 99 | 303 | 5 | MOLECULE: HEMOLYSIN E; |
| 1086 | 2n8p-A | 2.5 | 3.3 | 44 | 53 | 16 | MOLECULE: LACTICIN Q; |
| 1087 | 6mj1-A | 2.5 | 5 | 58 | 200 | 12 | MOLECULE: PROBABLE HTH-TYPE TRANSCRIPTIONAL REGULATOR YTTP; |
| 1088 | 3pux-G | 2.5 | 9.1 | 70 | 293 | 10 | MOLECULE: MALTOSE-BINDING PERIPLASMIC PROTEIN; |
| 1089 | 4ebj-A | 2.5 | 8.2 | 78 | 258 | 5 | MOLECULE: AMINOGLYCOSIDE NUCLEOTIDYLTRANSFERASE; |
| 1090 | 6q6e-A | 2.5 | 6 | 77 | 221 | 8 | MOLECULE: CONDENSIN COMPLEX SUBUNIT 2,STRUCTURAL MAINTENANC |
| 1091 | 2p58-B | 2.5 | 1.9 | 34 | 38 | 9 | MOLECULE: PUTATIVE TYPE III SECRETION PROTEIN YSCE; |
| 1092 | 2np9-A | 2.5 | 5.3 | 59 | 423 | 8 | MOLECULE: DPGC; |
| 1093 | 6ofa-A | 2.5 | 2.3 | 31 | 32 | 10 | MOLECULE: WASABI RECEPTOR TOXIN; |
| 1094 | 5ikn-A | 2.5 | 9 | 87 | 639 | 6 | MOLECULE: DNA-DIRECTED DNA POLYMERASE; |
| 1095 | 1ng6-A | 2.5 | 6.3 | 54 | 148 | 13 | MOLECULE: HYPOTHETICAL PROTEIN YQEY; |
| 1096 | 1w5s-B | 2.5 | 6.2 | 65 | 396 | 6 | MOLECULE: ORIGIN RECOGNITION COMPLEX SUBUNIT 2 ORC2; |
| 1097 | 6g1f-A | 2.5 | 11.8 | 77 | 443 | 6 | MOLECULE: D-PHENYLGLYCINE AMINOTRANSFERASE; |
| 1098 | 6x1g-A | 2.5 | 12.6 | 57 | 216 | 4 | MOLECULE: ULP_PROTEASE DOMAIN-CONTAINING PROTEIN; |
| 1099 | 3oos-A | 2.5 | 11.8 | 57 | 278 | 4 | MOLECULE: ALPHA/BETA HYDROLASE FAMILY PROTEIN; |
| 1100 | 6vtk-A | 2.5 | 5.4 | 54 | 446 | 7 | MOLECULE: ACID-SENSING ION CHANNEL 1; |
| 1101 | 6tgb-B | 2.5 | 10.9 | 69 | 680 | 9 | MOLECULE: DEDICATOR OF CYTOKINESIS PROTEIN 2; |
| 1102 | 5ys9-A | 2.5 | 12.3 | 88 | 692 | 3 | MOLECULE: ACYL-COENZYME A OXIDASE 3; |
| 1103 | 3bos-B | 2.5 | 3.7 | 50 | 231 | 16 | MOLECULE: PUTATIVE DNA REPLICATION FACTOR; |
| 1104 | 2ras-B | 2.5 | 12.1 | 76 | 204 | 11 | MOLECULE: TRANSCRIPTIONAL REGULATOR, TETR FAMILY; |
| 1105 | 5csk-A | 2.5 | 15.4 | 73 | 1996 | 11 | MOLECULE: ACETYL-COA CARBOXYLASE; |
| 1106 | 6ta5-D | 2.5 | 10.2 | 56 | 345 | 5 | MOLECULE: OUTER MEMBRANE PROTEIN OPRM; |
| 1107 | 6ux5-A | 2.5 | 2.9 | 37 | 50 | 5 | MOLECULE: U-ACTITOXIN-AEQ5A; |
| 1108 | 2oeq-C | 2.5 | 5.2 | 49 | 117 | 10 | MOLECULE: PROTEIN OF UNKNOWN FUNCTION, DUF964; |
| 1109 | 1nbw-A | 2.5 | 8 | 93 | 606 | 4 | MOLECULE: GLYCEROL DEHYDRATASE REACTIVASE ALPHA SUBUNIT; |
| 1110 | 4whj-A | 2.5 | 6.4 | 79 | 565 | 9 | MOLECULE: INTERFERON-INDUCED GTP-BINDING PROTEIN MX2; |
| 1111 | 3may-C | 2.5 | 3 | 52 | 97 | 4 | MOLECULE: POSSIBLE EXPORTED PROTEIN; |
| 1112 | 6zyx-Y | 2.5 | 16.8 | 94 | 1067 | 10 | MOLECULE: DYNEIN HEAVY CHAIN, OUTER ARM PROTEIN; |
| 1113 | 3b5o-A | 2.5 | 8.9 | 60 | 230 | 7 | MOLECULE: CADD-LIKE PROTEIN OF UNKNOWN FUNCTION; |
| 1114 | 3nuf-A | 2.5 | 6.5 | 54 | 118 | 15 | MOLECULE: PRD-CONTAINING TRANSCRIPTION REGULATOR; |
| 1115 | 5npl-A | 2.5 | 4.2 | 39 | 181 | 3 | MOLECULE: SIMILAR TO TR\|Q8YYT1\|Q8YYT1; |
| 1116 | 2pgc-A | 2.5 | 14.8 | 55 | 206 | 4 | MOLECULE: UNCHARACTERIZED PROTEIN; |
| 1117 | 4fmt-A | 2.5 | 8.3 | 59 | 209 | 10 | MOLECULE: CHPT PROTEIN; |

| | | | | | | | |
|---|---|---|---|---|---|---|---|
| 1118 | 6jbc-A | 2.5 | 13.6 | 54 | 295 | 6 | MOLECULE: PANTOATE KINASE; |
| 1119 | 2gf4-A | 2.5 | 5.7 | 58 | 89 | 9 | MOLECULE: PROTEIN VNG1086C; |
| 1120 | 6ts2-D | 2.5 | 9.3 | 100 | 1123 | 5 | MOLECULE: UDP-GLUCOSE-GLYCOPROTEIN GLUCOSYLTRANSFERASE- |
| 1121 | 3stq-A | 2.5 | 6.1 | 62 | 86 | 6 | MOLECULE: PUTATIVE UNCHARACTERIZED PROTEIN; |
| 1122 | 3enc-A | 2.5 | 7.4 | 48 | 79 | 4 | MOLECULE: PROTEIN PCC1; |
| 1123 | 6sxf-A | 2.5 | 12.1 | 70 | 263 | 3 | MOLECULE: ION TRANSPORT PROTEIN; |
| 1124 | 5gvx-A | 2.5 | 7.5 | 66 | 386 | 11 | MOLECULE: TREHALOSE-PHOSPHATE PHOSPHATASE; |
| 1125 | 6pwy-B | 2.5 | 4.8 | 89 | 513 | 8 | MOLECULE: ZK177.8; |
| 1126 | 6so5-E | 2.5 | 3.5 | 45 | 103 | 4 | MOLECULE: ATPASE ASNA1; |
| 1127 | 6lk8-J | 2.5 | 9.6 | 60 | 1026 | 2 | MOLECULE: MGC83295 PROTEIN; |
| 1128 | 5yma-A | 2.5 | 7.4 | 57 | 179 | 7 | MOLECULE: PUTATIVE RRNA PROCESSING PROTEIN; |
| 1129 | 1kf6-D | 2.5 | 9.1 | 54 | 119 | 7 | MOLECULE: FUMARATE REDUCTASE FLAVOPROTEIN; |
| 1130 | 5hc9-A | 2.5 | 10.2 | 69 | 425 | 10 | MOLECULE: TRNA NUCLEOTIDYL TRANSFERASE-RELATED PROTEIN; |
| 1131 | 1gvn-A | 2.5 | 6.7 | 51 | 87 | 4 | MOLECULE: EPSILON; |
| 1132 | 5ejk-D | 2.5 | 7 | 72 | 250 | 11 | MOLECULE: GAG-PRO-POL POLYPROTEIN; |
| 1133 | 1ghh-A | 2.5 | 2.5 | 39 | 81 | 10 | MOLECULE: DNA-DAMAGE-INDUCIBLE PROTEIN I; |
| 1134 | 7khw-a | 2.5 | 6.8 | 56 | 174 | 11 | MOLECULE: TRANSLOCON ESPA; |
| 1135 | 7emf-G | 2.5 | 14.4 | 67 | 161 | 3 | MOLECULE: MEDIATOR OF RNA POLYMERASE II TRANSCRIPTION SUBUN |
| 1136 | 6a70-B | 2.5 | 11 | 52 | 704 | 4 | MOLECULE: POLYCYSTIN-2; |
| 1137 | 1ixs-A | 2.5 | 3.2 | 42 | 50 | 19 | MOLECULE: HOLLIDAY JUNCTION DNA HELICASE RUVA; |
| 1138 | 2z51-A | 2.5 | 3.5 | 53 | 154 | 9 | MOLECULE: NIFU-LIKE PROTEIN 2, CHLOROPLAST; |
| 1139 | 6c94-A | 2.5 | 5.9 | 50 | 482 | 10 | MOLECULE: CYTOCHROME P450 4B1; |
| 1140 | 6c14-B | 2.5 | 6 | 59 | 164 | 2 | MOLECULE: PROTOCADHERIN-15; |
| 1141 | 6pe4-A | 2.5 | 7.7 | 73 | 758 | 8 | MOLECULE: V-TYPE PROTON ATPASE SUBUNIT A, VACUOLAR ISOFORM; |
| 1142 | 3cra-A | 2.5 | 11.9 | 48 | 239 | 2 | MOLECULE: PROTEIN MAZG; |
| 1143 | 3gfi-A | 2.5 | 5.9 | 51 | 143 | 6 | MOLECULE: 146AA LONG HYPOTHETICAL TRANSCRIPTIONAL REGULATOR |
| 1144 | 6vfi-A | 2.5 | 4.4 | 75 | 197 | 8 | MOLECULE: O43_DN18B; |
| 1145 | 2ccy-A | 2.5 | 5.3 | 43 | 127 | 7 | MOLECULE: CYTOCHROME C; |
| 1146 | 3ezu-A | 2.5 | 7.5 | 76 | 336 | 5 | MOLECULE: GGDEF DOMAIN PROTEIN; |
| 1147 | 6blj-A | 2.5 | 8 | 70 | 473 | 10 | MOLECULE: SERINE-TRNA LIGASE; |
| 1148 | 4cz8-A | 2.5 | 12.8 | 75 | 422 | 15 | MOLECULE: NA+/H+ ANTIPORTER, PUTATIVE; |
| 1149 | 4y68-A | 2.5 | 13.3 | 75 | 287 | 8 | MOLECULE: PUTATIVE NISIN-RESISTANCE PROTEIN; |
| 1150 | 2q83-A | 2.5 | 6.5 | 69 | 332 | 3 | MOLECULE: YTAA PROTEIN; |
| 1151 | 2p11-A | 2.5 | 10.2 | 70 | 219 | 7 | MOLECULE: HYPOTHETICAL PROTEIN; |
| 1152 | 5htf-B | 2.5 | 16.1 | 80 | 260 | 6 | MOLECULE: FOLDASE PROTEIN PRSA 1; |

| | | | | | | | |
|---|---|---|---|---|---|---|---|
| 1153 | 4ghn-A | 2.5 | 7.8 | 85 | 388 | 9 | MOLECULE: UNCHARACTERIZED PROTEIN; |
| 1154 | 3c98-B | 2.5 | 11.4 | 88 | 230 | 10 | MOLECULE: SYNTAXIN-BINDING PROTEIN 1; |
| 1155 | 3sjb-D | 2.5 | 6.6 | 51 | 72 | 10 | MOLECULE: ATPASE GET3; |
| 1156 | 4noo-B | 2.5 | 4 | 49 | 95 | 8 | MOLECULE: VGRG PROTEIN; |
| 1157 | 3o0q-A | 2.5 | 14 | 61 | 617 | 11 | MOLECULE: RIBONUCLEOSIDE-DIPHOSPHATE REDUCTASE; |
| 1158 | 5twa-A | 2.5 | 10.4 | 47 | 161 | 11 | MOLECULE: BCL-X HOMOLOGOUS PROTEIN, BHP2; |
| 1159 | 1p49-A | 2.5 | 5.6 | 55 | 549 | 4 | MOLECULE: STERYL-SULFATASE; |
| 1160 | 6tgb-A | 2.5 | 17.4 | 90 | 1450 | 4 | MOLECULE: DEDICATOR OF CYTOKINESIS PROTEIN 2; |
| 1161 | 6y07-A | 2.5 | 5.8 | 60 | 154 | 10 | MOLECULE: SOHAIR; |
| 1162 | 2gb7-B | 2.5 | 6.4 | 68 | 295 | 9 | MOLECULE: DNA STRAND 1; |
| 1163 | 5dzr-A | 2.5 | 12.4 | 67 | 965 | 9 | MOLECULE: FRQ-INTERACTING RNA HELICASE; |
| 1164 | 7nvr-n | 2.5 | 8.5 | 58 | 132 | 16 | MOLECULE: TFIIH BASAL TRANSCRIPTION FACTOR COMPLEX HELICASE |
| 1165 | 3wvz-A | 2.5 | 5.4 | 42 | 192 | 0 | MOLECULE: PROTEIN HIKESHI; |
| 1166 | 6sny-A | 2.5 | 4.1 | 50 | 107 | 14 | MOLECULE: SYNTHETIC EPCR BINDING PROTEIN; |
| 1167 | 5z9o-A | 2.5 | 7 | 70 | 373 | 9 | MOLECULE: CYCLOPROPANE-FATTY-ACYL-PHOSPHOLIPID SYNTHASE; |
| 1168 | 2ou3-A | 2.5 | 8.7 | 57 | 160 | 2 | MOLECULE: TELLURITE RESISTANCE PROTEIN OF COG3793; |
| 1169 | 5zt3-A | 2.5 | 3.5 | 51 | 114 | 8 | MOLECULE: WA352; |
| 1170 | 6gwj-B | 2.5 | 3.6 | 47 | 84 | 11 | MOLECULE: EKC/KEOPS COMPLEX SUBUNIT LAGE3; |
| 1171 | 2g0u-A | 2.5 | 6.5 | 62 | 92 | 6 | MOLECULE: TYPE III SECRETION SYSTEM NEEDLE PROTEIN; |
| 1172 | 5vj4-A | 2.5 | 9.7 | 74 | 275 | 9 | MOLECULE: UNCHARACTERIZED PROTEIN; |
| 1173 | 3hr0-A | 2.5 | 5.2 | 56 | 250 | 9 | MOLECULE: COG4; |
| 1174 | 3n71-A | 2.5 | 15.5 | 91 | 469 | 11 | MOLECULE: HISTONE LYSINE METHYLTRANSFERASE SMYD1; |
| 1175 | 5tth-A | 2.5 | 12.4 | 85 | 671 | 7 | MOLECULE: C-TERMINAL SPYCATCHER FUSION OF WILDTYPE ZEBRAFIS |
| 1176 | 7kzn-F | 2.5 | 4.5 | 55 | 100 | 11 | MOLECULE: HEAVY CHAIN ALPHA; |
| 1177 | 5da9-A | 2.5 | 6.5 | 63 | 433 | 5 | MOLECULE: PUTATIVE UNCHARACTERIZED PROTEIN,PUTATIVE UNCHARA |
| 1178 | 3dm0-A | 2.5 | 10.7 | 64 | 675 | 3 | MOLECULE: MALTOSE-BINDING PERIPLASMIC PROTEIN FUSED WITH RA |
| 1179 | 6k0b-G | 2.5 | 5.5 | 46 | 120 | 11 | MOLECULE: RIBONUCLEASE P PROTEIN COMPONENT 2; |
| 1180 | 6vld-B | 2.5 | 5.4 | 52 | 470 | 6 | MOLECULE: ALPHA-(1,6)-FUCOSYLTRANSFERASE; |
| 1181 | 3b40-A | 2.5 | 14.9 | 70 | 400 | 6 | MOLECULE: PROBABLE DIPEPTIDASE; |
| 1182 | 2id3-A | 2.5 | 12.3 | 72 | 191 | 4 | MOLECULE: PUTATIVE TRANSCRIPTIONAL REGULATOR; |
| 1183 | 6so5-C | 2.5 | 7.5 | 58 | 144 | 7 | MOLECULE: ATPASE ASNA1; |
| 1184 | 6jl7-A | 2.5 | 10.1 | 88 | 444 | 8 | MOLECULE: TBC1 DOMAIN FAMILY MEMBER 23; |
| 1185 | 3wqy-A | 2.5 | 12 | 71 | 906 | 11 | MOLECULE: ALANINE--TRNA LIGASE; |
| 1186 | 3r6n-A | 2.5 | 10 | 80 | 450 | 5 | MOLECULE: DESMOPLAKIN; |
| 1187 | 5ot4-A | 2.5 | 10.9 | 75 | 846 | 7 | MOLECULE: INTERAPTIN; |

| | | | | | | | |
|---|---|---|---|---|---|---|---|
| 1188 | 4zsf-A | 2.5 | 10.4 | 72 | 272 | 8 | MOLECULE: BSAWI ENDONUCLEASE; |
| 1189 | 1ls4-A | 2.5 | 4.4 | 76 | 164 | 7 | MOLECULE: APOLIPOPHORIN-III; |
| 1190 | 3zh9-B | 2.5 | 11.8 | 76 | 339 | 17 | MOLECULE: DELTA; |
| 1191 | 7mi4-A | 2.5 | 12.7 | 72 | 554 | 7 | MOLECULE: CRISPR-ASSOCIATED EXONUCLEASE CAS4/ENDONUCLEASE C |
| 1192 | 6h3a-B | 2.5 | 2.5 | 39 | 57 | 8 | MOLECULE: SWI/SNF-RELATED MATRIX-ASSOCIATED ACTIN-DEPENDENT |
| 1193 | 6yj4-p | 2.5 | 6.9 | 52 | 91 | 15 | MOLECULE: NADH-UBIQUINONE OXIDOREDUCTASE CHAIN 3; |
| 1194 | 5kbw-B | 2.5 | 6.2 | 68 | 171 | 0 | MOLECULE: RIBOFLAVIN TRANSPORTER RIBU; |
| 1195 | 7rpk-A | 2.5 | 7.5 | 103 | 953 | 8 | MOLECULE: PROTEIN DISPATCHED HOMOLOG 1; |
| 1196 | 4ua3-B | 2.5 | 11.1 | 58 | 193 | 7 | MOLECULE: UNCHARACTERIZED N-ACETYLTRANSFERASE C825.04C; |
| 1197 | 3e98-A | 2.5 | 4.5 | 71 | 178 | 6 | MOLECULE: GAF DOMAIN OF UNKNOWN FUNCTION; |
| 1198 | 2hkv-A | 2.5 | 3.7 | 72 | 148 | 4 | MOLECULE: HYPOTHETICAL PROTEIN; |
| 1199 | 6zw0-A | 2.5 | 17.1 | 65 | 299 | 11 | MOLECULE: CONNECTASE MJ0548; |
| 1200 | 5ikf-A | 2.5 | 7.2 | 63 | 150 | 6 | MOLECULE: CHROMATIN REMODELING FACTOR MIT1; |
| 1201 | 6y86-A | 2.5 | 5.7 | 75 | 425 | 8 | MOLECULE: MEMBRANE PROTEIN INSERTASE YIDC; |
| 1202 | 4am6-A | 2.5 | 12.7 | 62 | 623 | 13 | MOLECULE: ACTIN-LIKE PROTEIN ARP8; |
| 1203 | 6wi5-A | 2.5 | 3.8 | 55 | 90 | 11 | MOLECULE: DE NOVO DESIGNED PROTEIN FOLDIT4; |
| 1204 | 6zy2-E | 2.5 | 5.6 | 67 | 259 | 4 | MOLECULE: YRBD PROTEIN; |
| 1205 | 6v8o-O | 2.5 | 16.8 | 72 | 384 | 10 | MOLECULE: HIGH TEMPERATURE LETHAL PROTEIN 1; |
| 1206 | 6nqx-A | 2.5 | 5.7 | 82 | 246 | 10 | MOLECULE: FLAGELLAR COILING PROTEIN A; |
| 1207 | 4q5i-B | 2.5 | 16.6 | 80 | 269 | 6 | MOLECULE: FERROUS IRON TRANSPORT PROTEIN B; |
| 1208 | 5b2o-A | 2.5 | 9.5 | 96 | 1455 | 9 | MOLECULE: CRISPR-ASSOCIATED ENDONUCLEASE CAS9; |
| 1209 | 3err-A | 2.5 | 7.8 | 62 | 527 | 10 | MOLECULE: FUSION PROTEIN OF MICROTUBULE BINDING DOMAIN FROM |
| 1210 | 4bpm-A | 2.5 | 2.9 | 51 | 162 | 14 | MOLECULE: PROSTAGLANDIN E SYNTHASE, FUSION PEPTIDE; |
| 1211 | 5fyw-M | 2.5 | 11 | 67 | 231 | 4 | MOLECULE: DNA-DIRECTED RNA POLYMERASE II SUBUNIT RPB1; |
| 1212 | 6yj4-V | 2.5 | 7.1 | 49 | 126 | 6 | MOLECULE: NADH-UBIQUINONE OXIDOREDUCTASE CHAIN 3; |
| 1213 | 5gj8-D | 2.5 | 4.6 | 60 | 121 | 3 | MOLECULE: ACYL-COA DEHYDROGENASE TYPE 2 DOMAIN PROTEIN; |
| 1214 | 6yxa-A | 2.5 | 7.9 | 61 | 536 | 10 | MOLECULE: GTP PYROPHOSPHOKINASE; |
| 1215 | 6j98-A | 2.5 | 2.3 | 48 | 75 | 6 | MOLECULE: P8; |
| 1216 | 7lgu-A | 2.5 | 19.6 | 82 | 680 | 7 | MOLECULE: PRESTIN; |
| 1217 | 5w66-M | 2.5 | 7.7 | 72 | 396 | 8 | MOLECULE: DNA-DIRECTED RNA POLYMERASE I SUBUNIT RPA190; |
| 1218 | 7b1s-B | 2.5 | 5.8 | 82 | 466 | 6 | MOLECULE: ETHYL-COENZYME M REDUCTASE ALPHA SUBUNIT; |
| 1219 | 6vjy-B | 2.5 | 11 | 97 | 280 | 5 | MOLECULE: ERAD-ASSOCIATED E3 UBIQUITIN-PROTEIN LIGASE HRD1; |
| 1220 | 5x6b-J | 2.5 | 8.3 | 74 | 377 | 11 | MOLECULE: O-PHOSPHO-L-SERYL-TRNA:CYS-TRNA SYNTHASE; |
| 1221 | 5ee5-A | 2.5 | 8.2 | 63 | 202 | 8 | MOLECULE: BREFELDIN A-INHIBITED GUANINE NUCLEOTIDE-EXCHANGE |
| 1222 | 7jv7-B | 2.5 | 5.3 | 81 | 310 | 6 | MOLECULE: CTD KINASE SUBUNIT ALPHA; |

| | | | | | | | |
|---|---|---|---|---|---|---|---|
| 1223 | 4avm-A | 2.5 | 9.5 | 78 | 230 | 8 | MOLECULE: BRIDGING INTEGRATOR 2; |
| 1224 | 7mi1-A | 2.5 | 11.4 | 100 | 2628 | 5 | MOLECULE: CHIMERA PROTEIN OF DYNEIN AND ENDOLYSIN; |
| 1225 | 1rso-B | 2.5 | 4.9 | 44 | 56 | 16 | MOLECULE: PRESYNAPTIC PROTEIN SAP97; |
| 1226 | 7jts-s | 2.5 | 5.7 | 58 | 290 | 12 | MOLECULE: RADIAL SPOKE PROTEIN 3; |
| 1227 | 5gad-i | 2.5 | 6.6 | 73 | 450 | 8 | MOLECULE: ESRP 4.5S RNA; |
| 1228 | 6k6l-B | 2.5 | 6.6 | 73 | 263 | 8 | MOLECULE: PSEUDO DEUBIQUITINASE; |
| 1229 | 4nb5-B | 2.5 | 4.8 | 49 | 149 | 4 | MOLECULE: DNA BINDING PROTEIN; |
| 1230 | 5sva-V | 2.5 | 6.9 | 57 | 85 | 11 | MOLECULE: DNA-DIRECTED RNA POLYMERASE II SUBUNIT RPB1; |
| 1231 | 2y1v-A | 2.5 | 8.9 | 73 | 604 | 7 | MOLECULE: CELL WALL SURFACE ANCHOR FAMILY PROTEIN; |
| 1232 | 3s84-B | 2.5 | 9.8 | 78 | 241 | 6 | MOLECULE: APOLIPOPROTEIN A-IV; |
| 1233 | 6w2r-B | 2.5 | 10 | 65 | 221 | 9 | MOLECULE: JUNCTION 19 DHR54-DHR79; |
| 1234 | 1qdb-A | 2.5 | 19.3 | 92 | 473 | 10 | MOLECULE: CYTOCHROME C NITRITE REDUCTASE; |
| 1235 | 5yud-C | 2.5 | 4.8 | 49 | 75 | 12 | MOLECULE: BACULOVIRAL IAP REPEAT-CONTAINING PROTEIN 1E; |
| 1236 | 6ty9-B | 2.5 | 8.8 | 55 | 501 | 4 | MOLECULE: RNA-DEPENDENT RNA POLYMERASE; |
| 1237 | 6qyi-B | 2.5 | 11 | 81 | 519 | 5 | MOLECULE: 4-HYDROXYPHENYLACETATE 3-MONOOXYGENASE |
| 1238 | 7nyx-A | 2.5 | 7.7 | 65 | 1467 | 8 | MOLECULE: CHROMOSOME PARTITION PROTEIN MUKB; |
| 1239 | 1jeq-A | 2.5 | 16.9 | 72 | 548 | 10 | MOLECULE: KU70; |
| 1240 | 3bvo-A | 2.5 | 9.2 | 65 | 197 | 6 | MOLECULE: CO-CHAPERONE PROTEIN HSCB, MITOCHONDRIAL PRECURSO |
| 1241 | 5yz0-A | 2.5 | 13.3 | 84 | 2362 | 8 | MOLECULE: SERINE/THREONINE-PROTEIN KINASE ATR; |
| 1242 | 6cc4-A | 2.5 | 4.9 | 67 | 608 | 4 | MOLECULE: SOLUBLE CYTOCHROME B562, LIPID II FLIPPASE MURJ C |
| 1243 | 7cpx-A | 2.5 | 4.8 | 87 | 2262 | 8 | MOLECULE: LOVASTATIN NONAKETIDE SYNTHASE, POLYKETIDE SYNTHA |
| 1244 | 3n27-B | 2.5 | 4.6 | 43 | 79 | 5 | MOLECULE: FUSION GLYCOPROTEIN F0, LINKER, FUSION GLYCOPROTE |
| 1245 | 4cej-A | 2.5 | 7.6 | 65 | 1177 | 5 | MOLECULE: ATP-DEPENDENT HELICASE/NUCLEASE SUBUNIT A; |
| 1246 | 6v69-J | 2.5 | 7.5 | 81 | 484 | 10 | MOLECULE: GAMMA-TUBULIN COMPLEX COMPONENT 5; |
| 1247 | 7cfa-A | 2.5 | 5.9 | 54 | 234 | 9 | MOLECULE: R.PAB1 FAMILY RESTRICTION ENDONUCLEASE; |
| 1248 | 3di5-A | 2.5 | 3.8 | 76 | 150 | 5 | MOLECULE: DINB-LIKE PROTEIN; |
| 1249 | 6tdx-O | 2.5 | 4.1 | 50 | 81 | 8 | MOLECULE: ATP SYNTHASE F1 SUBUNIT GAMMA; |
| 1250 | 7emf-P | 2.5 | 7.9 | 78 | 766 | 12 | MOLECULE: MEDIATOR OF RNA POLYMERASE II TRANSCRIPTION SUBUN |
| 1251 | 3ph0-A | 2.5 | 5 | 46 | 60 | 7 | MOLECULE: ASCE; |
| 1252 | 6s7j-A | 2.5 | 10.8 | 110 | 497 | 8 | MOLECULE: UNCHARACTERIZED PROTEIN; |
| 1253 | 2is5-C | 2.5 | 3.3 | 55 | 134 | 4 | MOLECULE: HYPOTHETICAL PROTEIN; |
| 1254 | 5a63-C | 2.5 | 11.4 | 84 | 243 | 6 | MOLECULE: NICASTRIN; |
| 1255 | 6dlm-B | 2.5 | 5 | 48 | 72 | 6 | MOLECULE: DHD127_A; |
| 1256 | 4uzx-A | 2.5 | 4.1 | 41 | 67 | 27 | MOLECULE: PROTEIN THO1; |
| 1257 | 6vq6-A | 2.5 | 9.1 | 82 | 600 | 5 | MOLECULE: ATPASE H+-TRANSPORTING V1 SUBUNIT A; |

| | | | | | | | |
|---|---|---|---|---|---|---|---|
| 1258 | 7lkm-A | 2.4 | 2.9 | 58 | 149 | 2 | MOLECULE: PILUS BIOGENESIS PROTEIN; |
| 1259 | 4d2i-A | 2.4 | 22.8 | 93 | 464 | 9 | MOLECULE: HERA; |
| 1260 | 5lde-B | 2.4 | 2.9 | 64 | 225 | 8 | MOLECULE: IMMUNOGLOBULIN G-BINDING PROTEIN G,VIRAL FLICE PR |
| 1261 | 4zhe-D | 2.4 | 10.8 | 86 | 206 | 7 | MOLECULE: ASPR2 PROTEIN; |
| 1262 | 3n0r-A | 2.4 | 11.4 | 69 | 258 | 3 | MOLECULE: RESPONSE REGULATOR; |
| 1263 | 6sp2-A | 2.4 | 7.6 | 83 | 366 | 8 | MOLECULE: MEMBRANE PROTEIN TMS1D; |
| 1264 | 5wd8-A | 2.4 | 5.3 | 44 | 90 | 7 | MOLECULE: UNCHARACTERIZED PROTEIN; |
| 1265 | 5lxj-A | 2.4 | 2.9 | 44 | 53 | 5 | MOLECULE: PHOSPHOPROTEIN; |
| 1266 | 6zka-s | 2.4 | 7.3 | 57 | 122 | 5 | MOLECULE: NADH-UBIQUINONE OXIDOREDUCTASE CHAIN 3; |
| 1267 | 1x2g-C | 2.4 | 2.3 | 46 | 337 | 7 | MOLECULE: LIPOATE-PROTEIN LIGASE A; |
| 1268 | 1x04-A | 2.4 | 7.7 | 73 | 200 | 10 | MOLECULE: SH3-CONTAINING GRB2-LIKE PROTEIN 2; |
| 1269 | 7kyp-F | 2.4 | 5.4 | 65 | 290 | 5 | MOLECULE: MANGANESE ABC TRANSPORTER, ATP-BINDING PROTEIN; |
| 1270 | 6gyr-A | 2.4 | 14.6 | 79 | 589 | 9 | MOLECULE: HISTONE ACETYLTRANSFERASE P300; |
| 1271 | 2x43-S | 2.4 | 4.4 | 48 | 67 | 8 | MOLECULE: SHERP; |
| 1272 | 2rji-A | 2.4 | 2.1 | 41 | 84 | 15 | MOLECULE: ERYTHROCYTE BINDING ANTIGEN 175; |
| 1273 | 1gjs-A | 2.4 | 2.6 | 41 | 65 | 7 | MOLECULE: IMMUNOGLOBULIN G BINDING PROTEIN G; |
| 1274 | 2k3q-A | 2.4 | 4.1 | 64 | 118 | 11 | MOLECULE: TUSP1; |
| 1275 | 6tmh-H | 2.4 | 3.9 | 42 | 71 | 7 | MOLECULE: INHIBITOR OF F1; |
| 1276 | 5m1m-A | 2.4 | 5 | 47 | 154 | 4 | MOLECULE: MATRIX PROTEIN 1; |
| 1277 | 2zc2-A | 2.4 | 2.9 | 46 | 75 | 9 | MOLECULE: DNAD-LIKE REPLICATION PROTEIN; |
| 1278 | 2dhy-A | 2.4 | 3.4 | 45 | 67 | 9 | MOLECULE: CUE DOMAIN-CONTAINING PROTEIN 1; |
| 1279 | 5hsb-A | 2.4 | 8.6 | 46 | 202 | 9 | MOLECULE: RNA POLYMERASE; |
| 1280 | 6nsm-A | 2.4 | 10.4 | 63 | 194 | 3 | MOLECULE: COPPER OUTER MEMBRANE REGULATOR; |
| 1281 | 4i0x-F | 2.4 | 4.3 | 44 | 85 | 9 | MOLECULE: ESAT-6-LIKE PROTEIN MAB_3112; |
| 1282 | 4oge-A | 2.4 | 12.7 | 84 | 977 | 7 | MOLECULE: HNH ENDONUCLEASE DOMAIN PROTEIN; |
| 1283 | 3uo3-B | 2.4 | 6.6 | 53 | 147 | 9 | MOLECULE: J-TYPE CO-CHAPERONE JAC1, MITOCHONDRIAL; |
| 1284 | 2yev-C | 2.4 | 4.7 | 47 | 64 | 11 | MOLECULE: CYTOCHROME C OXIDASE POLYPEPTIDE I+III; |
| 1285 | 6eqo-A | 2.4 | 8.8 | 86 | 1804 | 9 | MOLECULE: ACETYL-COENZYME A SYNTHETASE; |
| 1286 | 3gnw-B | 2.4 | 10.6 | 83 | 568 | 8 | MOLECULE: RNA-DIRECTED RNA POLYMERASE; |
| 1287 | 5gar-O | 2.4 | 4.8 | 44 | 79 | 7 | MOLECULE: V-TYPE ATP SYNTHASE ALPHA CHAIN; |
| 1288 | 6qbi-A | 2.4 | 11.7 | 48 | 102 | 8 | MOLECULE: SURFACE PROTEIN, MLP LIPOPROTEIN FAMILY; |
| 1289 | 3aai-A | 2.4 | 8.2 | 57 | 78 | 14 | MOLECULE: COPPER HOMEOSTASIS OPERON REGULATORY PROTEIN; |
| 1290 | 6row-B | 2.4 | 9.3 | 77 | 619 | 8 | MOLECULE: PUTATIVE ZINC METALLOPEPTIDASE; |
| 1291 | 6ffv-A | 2.4 | 8.4 | 76 | 194 | 9 | MOLECULE: BTUM; |
| 1292 | 6e67-B | 2.4 | 4.2 | 59 | 476 | 7 | MOLECULE: BETA-2 ADRENERGIC RECEPTOR,ENDOLYSIN,GUANINE NUCL |

| | | | | | | | |
|---|---|---|---|---|---|---|---|
| 1293 | 2oh3-A | 2.4 | 6.3 | 63 | 149 | 10 | MOLECULE: COG1633: UNCHARACTERIZED CONSERVED PROTEIN; |
| 1294 | 2pjw-H | 2.4 | 8.3 | 62 | 88 | 3 | MOLECULE: UNCHARACTERIZED PROTEIN YHL002W; |
| 1295 | 3b4q-B | 2.4 | 2.7 | 48 | 90 | 10 | MOLECULE: UNCHARACTERIZED PROTEIN; |
| 1296 | 6n10-A | 2.4 | 11.3 | 52 | 403 | 10 | MOLECULE: DIPHOSPHOMEVALONATE DECARBOXYLASE MVD1, |
| 1297 | 6h4b-B | 2.4 | 8 | 48 | 84 | 6 | MOLECULE: ORF026; |
| 1298 | 5oqj-W | 2.4 | 13 | 62 | 258 | 6 | MOLECULE: DNA-DIRECTED RNA POLYMERASE II SUBUNIT RPB1; |
| 1299 | 6p8r-B | 2.4 | 9.8 | 55 | 168 | 9 | MOLECULE: HORMA DOMAIN CONTAINING PROTEIN; |
| 1300 | 2p1a-B | 2.4 | 3.2 | 53 | 150 | 9 | MOLECULE: HYPOTHETICAL PROTEIN; |
| 1301 | 7ol3-B | 2.4 | 3.2 | 42 | 425 | 14 | MOLECULE: ATLASTIN-1; |
| 1302 | 5l10-A | 2.4 | 8.7 | 53 | 172 | 4 | MOLECULE: N-ACYLHOMOSERINE LACTONE DEPENDENT REGULATORY |
| 1303 | 6rd4-R | 2.4 | 8.2 | 53 | 177 | 6 | MOLECULE: ASA-10: POLYTOMELLA F-ATP SYNTHASE ASSOCIATED SUB |
| 1304 | 6l81-A | 2.4 | 2.7 | 49 | 96 | 14 | MOLECULE: GAMMA-TUBULIN COMPLEX COMPONENT 5; |
| 1305 | 7rj1-A | 2.4 | 11.1 | 77 | 266 | 4 | MOLECULE: CHORISMATE MUTASE; |
| 1306 | 5tue-A | 2.4 | 6.9 | 62 | 388 | 8 | MOLECULE: TETRACYCLINE DESTRUCTASE TET(50); |
| 1307 | 6snh-X | 2.4 | 3.4 | 60 | 479 | 2 | MOLECULE: DOLICHYL PYROPHOSPHATE MAN9GLCNAC2 ALPHA-1,3- |
| 1308 | 7d7q-B | 2.4 | 4.4 | 78 | 310 | 10 | MOLECULE: PHOSPHODIESTERASE; |
| 1309 | 2r01-A | 2.4 | 5.8 | 70 | 195 | 6 | MOLECULE: NITROREDUCTASE FAMILY PROTEIN; |
| 1310 | 1tua-A | 2.4 | 12 | 59 | 189 | 7 | MOLECULE: HYPOTHETICAL PROTEIN APE0754; |
| 1311 | 5oxe-A | 2.4 | 4.9 | 45 | 70 | 4 | MOLECULE: MAJOR VIRION PROTEIN; |
| 1312 | 6rwb-A | 2.4 | 6.3 | 92 | 1873 | 5 | MOLECULE: TOXIN,TOXIN COMPLEX SUBUNIT TCAB,PUTATIVE TOXIN S |
| 1313 | 6xm1-D | 2.4 | 9.7 | 74 | 181 | 9 | MOLECULE: VPS45; |
| 1314 | 5xmk-G | 2.4 | 4.6 | 38 | 54 | 5 | MOLECULE: VACUOLAR PROTEIN SORTING-ASSOCIATED PROTEIN 4; |
| 1315 | 6vk0-C | 2.4 | 7.7 | 68 | 180 | 7 | MOLECULE: U1 SNP1-ASSOCIATING PROTEIN 1; |
| 1316 | 6yeu-A | 2.4 | 3.1 | 58 | 107 | 9 | MOLECULE: CALCIUM-BINDING EF HAND FAMILY PROTEIN; |
| 1317 | 6ysf-F | 2.4 | 10 | 74 | 257 | 1 | MOLECULE: CHEMOTAXIS MOTB PROTEIN; |
| 1318 | 6y1y-B | 2.4 | 6.1 | 56 | 128 | 4 | MOLECULE: CHEA; |
| 1319 | 6jho-A | 2.4 | 5.6 | 55 | 200 | 4 | MOLECULE: CAG PATHOGENICITY ISLAND PROTEIN (CAG6); |
| 1320 | 1vku-A | 2.4 | 2.4 | 49 | 85 | 10 | MOLECULE: ACYL CARRIER PROTEIN; |
| 1321 | 7nhr-A | 2.4 | 7.3 | 75 | 663 | 7 | MOLECULE: PUTATIVE TRANSMEMBRANE PROTEIN WZC; |
| 1322 | 1w8i-A | 2.4 | 3.1 | 44 | 155 | 2 | MOLECULE: PUTATIVE VAPC RIBONUCLEASE AF_1683; |
| 1323 | 2iie-A | 2.4 | 6.2 | 53 | 204 | 15 | MOLECULE: PHAGE P H' SITE; |
| 1324 | 5mmi-Z | 2.4 | 6.7 | 58 | 101 | 9 | MOLECULE: 50S RIBOSOMAL PROTEIN L31; |
| 1325 | 6en8-A | 2.4 | 3.2 | 47 | 191 | 13 | MOLECULE: TRANSCRIPTIONAL REGULATOR TETR FAMILY; |
| 1326 | 6yp7-z | 2.4 | 5.5 | 41 | 62 | 12 | MOLECULE: CHLOROPHYLL A-B BINDING PROTEIN 8, CHLOROPLASTIC; |
| 1327 | 2mmu-A | 2.4 | 4.4 | 40 | 50 | 5 | MOLECULE: CELL DIVISION PROTEIN CRGA; |

| | | | | | | | |
|---|---|---|---|---|---|---|---|
| 1328 | 2ymj-A | 2.4 | 7.2 | 39 | 52 | 8 | MOLECULE: PROTEIN QUAKING-A; |
| 1329 | 2qko-B | 2.4 | 13.1 | 75 | 176 | 7 | MOLECULE: POSSIBLE TRANSCRIPTIONAL REGULATOR, TETR FAMILY P |
| 1330 | 6m6z-A | 2.4 | 7 | 67 | 203 | 6 | MOLECULE: TMH4C4; |
| 1331 | 2ixp-B | 2.4 | 4.2 | 66 | 316 | 8 | MOLECULE: SERINE/THREONINE-PROTEIN PHOSPHATASE 2A ACTIVATOR |
| 1332 | 6g7o-A | 2.4 | 12.8 | 55 | 350 | 4 | MOLECULE: ALKALINE CERAMIDASE 3,SOLUBLE CYTOCHROME B562; |
| 1333 | 5yo8-A | 2.4 | 6.6 | 66 | 349 | 6 | MOLECULE: TETRAPRENYL-BETA-CURCUMENE SYNTHASE; |
| 1334 | 3fx7-B | 2.4 | 6.5 | 52 | 87 | 6 | MOLECULE: PUTATIVE UNCHARACTERIZED PROTEIN; |
| 1335 | 3di3-A | 2.4 | 5.4 | 46 | 118 | 13 | MOLECULE: INTERLEUKIN-7; |
| 1336 | 4kzs-A | 2.4 | 9.6 | 58 | 242 | 12 | MOLECULE: LPP20 LIPOFAMILY PROTEIN; |
| 1337 | 4mou-A | 2.4 | 11.7 | 63 | 261 | 5 | MOLECULE: ENOYL-COA HYDRATASE/ISOMERASE FAMILY PROTEIN; |
| 1338 | 4gmq-A | 2.4 | 2.7 | 54 | 92 | 15 | MOLECULE: PUTATIVE RIBOSOME ASSOCIATED PROTEIN; |
| 1339 | 5ly0-A | 2.4 | 6.4 | 66 | 108 | 3 | MOLECULE: LOB FAMILY TRANSFACTOR RAMOSA2.1; |
| 1340 | 5c37-A | 2.4 | 8 | 66 | 632 | 14 | MOLECULE: FATTY ACID SYNTHASE; |
| 1341 | 2g7s-A | 2.4 | 11.2 | 72 | 190 | 6 | MOLECULE: TRANSCRIPTIONAL REGULATOR, TETR FAMILY; |
| 1342 | 5gm2-B | 2.4 | 13 | 72 | 283 | 7 | MOLECULE: O-METHYLRANSFERASE; |
| 1343 | 3i5q-A | 2.4 | 4.4 | 45 | 248 | 16 | MOLECULE: NUCLEOPORIN NUP170; |
| 1344 | 2zdi-C | 2.4 | 5.7 | 49 | 148 | 6 | MOLECULE: PREFOLDIN SUBUNIT BETA; |
| 1345 | 6dtd-A | 2.4 | 4.3 | 77 | 1053 | 6 | MOLECULE: NUCLEASE; |
| 1346 | 6pl5-A | 2.4 | 8.5 | 89 | 336 | 15 | MOLECULE: PEPTIDOGLYCAN GLYCOSYLTRANSFERASE RODA; |
| 1347 | 7lg5-A | 2.4 | 5.4 | 64 | 871 | 9 | MOLECULE: CYANOPHYCIN SYNTHASE; |
| 1348 | 2byd-A | 2.4 | 5.6 | 77 | 283 | 6 | MOLECULE: HSPC223; |
| 1349 | 5zr1-E | 2.4 | 9.6 | 71 | 460 | 7 | MOLECULE: ORIGIN RECOGNITION COMPLEX SUBUNIT 1; |
| 1350 | 2job-A | 2.4 | 2.9 | 54 | 102 | 9 | MOLECULE: ANTILIPOPOLYSACCHARIDE FACTOR; |
| 1351 | 7asm-W | 2.4 | 4.3 | 43 | 66 | 14 | MOLECULE: 50S RIBOSOMAL PROTEIN L19; |
| 1352 | 3e7g-C | 2.4 | 12.7 | 89 | 423 | 2 | MOLECULE: NITRIC OXIDE SYNTHASE, INDUCIBLE; |
| 1353 | 5a1u-E | 2.4 | 17.7 | 75 | 822 | 9 | MOLECULE: ADP-RIBOSYLATION FACTOR 1; |
| 1354 | 6eml-p | 2.4 | 3 | 48 | 185 | 6 | MOLECULE: PRE-18S RIBOSOMAL RNA; |
| 1355 | 3hh0-A | 2.4 | 2 | 34 | 134 | 18 | MOLECULE: TRANSCRIPTIONAL REGULATOR, MERR FAMILY; |
| 1356 | 7jfs-A | 2.4 | 13 | 74 | 958 | 8 | MOLECULE: F5/8 TYPE C DOMAIN PROTEIN; |
| 1357 | 6eud-A | 2.4 | 14.8 | 87 | 808 | 11 | MOLECULE: ATP-DEPENDENT RNA HELICASE HRPB; |
| 1358 | 5jno-A | 2.4 | 2.9 | 52 | 97 | 4 | MOLECULE: BEN DOMAIN-CONTAINING PROTEIN 3; |
| 1359 | 6ck0-B | 2.4 | 4.3 | 66 | 210 | 3 | MOLECULE: BIOTIN ACETYL COENZYME A CARBOXYLASE SYNTHETASE; |
| 1360 | 6ejq-B | 2.4 | 9.7 | 51 | 141 | 8 | MOLECULE: TERMINASE SMALL SUBUNIT; |
| 1361 | 2zop-A | 2.4 | 3.6 | 51 | 113 | 12 | MOLECULE: PUTATIVE UNCHARACTERIZED PROTEIN TTHB164; |
| 1362 | 6zr2-Y | 2.4 | 7.4 | 69 | 140 | 10 | MOLECULE: NADH-UBIQUINONE OXIDOREDUCTASE CHAIN 3; |

| | | | | | | | |
|---|---|---|---|---|---|---|---|
| 1363 | 3c2b-A | 2.4 | 2.7 | 48 | 200 | 13 | MOLECULE: TRANSCRIPTIONAL REGULATOR, TETR FAMILY; |
| 1364 | 4y2f-A | 2.4 | 3.7 | 43 | 143 | 12 | MOLECULE: SENSOR PROTEIN KDPD; |
| 1365 | 3v9r-A | 2.4 | 11 | 62 | 88 | 2 | MOLECULE: UNCHARACTERIZED PROTEIN YOL086W-A; |
| 1366 | 5i2l-A | 2.4 | 2.9 | 44 | 105 | 5 | MOLECULE: EF-HAND DOMAIN-CONTAINING PROTEIN D2; |
| 1367 | 2ch5-A | 2.4 | 8.3 | 90 | 344 | 7 | MOLECULE: NAGK PROTEIN; |
| 1368 | 6xmv-A | 2.4 | 3.5 | 61 | 307 | 8 | MOLECULE: HEMOLYSIN,ENDOLYSIN; |
| 1369 | 5lut-F | 2.4 | 3.6 | 48 | 55 | 6 | MOLECULE: BLM HELICASE; |
| 1370 | 6ywd-C | 2.4 | 8.2 | 45 | 75 | 7 | MOLECULE: ANTIBODY MOTA, HEAVY CHAIN; |
| 1371 | 6vy1-A | 2.4 | 4.7 | 43 | 121 | 26 | MOLECULE: PREFOLDIN SUBUNIT ALPHA 2; |
| 1372 | 5nh2-B | 2.4 | 3.3 | 38 | 56 | 8 | MOLECULE: ADENOSINE MONOPHOSPHATE-PROTEIN TRANSFERASE; |
| 1373 | 5z62-B | 2.4 | 9.7 | 64 | 227 | 9 | MOLECULE: CYTOCHROME C OXIDASE SUBUNIT 1; |
| 1374 | 2i2x-B | 2.4 | 8.5 | 73 | 258 | 7 | MOLECULE: METHYLTRANSFERASE 1; |
| 1375 | 3v7o-B | 2.4 | 3.5 | 65 | 203 | 14 | MOLECULE: MINOR NUCLEOPROTEIN VP30; |
| 1376 | 5dic-A | 2.4 | 4.1 | 50 | 115 | 4 | MOLECULE: ODORANT-BINDING PROTEIN; |
| 1377 | 5ndd-A | 2.4 | 6.1 | 69 | 567 | 7 | MOLECULE: LYSOZYME,PROTEINASE-ACTIVATED RECEPTOR 2,SOLUBLE |
| 1378 | 1wpk-A | 2.4 | 10.2 | 69 | 146 | 6 | MOLECULE: ADA REGULATORY PROTEIN; |
| 1379 | 5hus-A | 2.4 | 12.3 | 61 | 293 | 7 | MOLECULE: TREHALOSE SYNTHASE REGULATORY PROTEIN; |
| 1380 | 2o57-A | 2.4 | 6.5 | 69 | 282 | 7 | MOLECULE: PUTATIVE SARCOSINE DIMETHYLGLYCINE METHYLTRANSFER |
| 1381 | 4jcs-A | 2.4 | 9.3 | 61 | 265 | 3 | MOLECULE: ENOYL-COA HYDRATASE/ISOMERASE; |
| 1382 | 2y31-A | 2.4 | 12.5 | 67 | 242 | 4 | MOLECULE: PUTATIVE REPRESSOR SIMREG2; |
| 1383 | 6qi8-E | 2.4 | 8.4 | 67 | 335 | 13 | MOLECULE: RUVB-LIKE 1; |
| 1384 | 6nnw-A | 2.4 | 5.7 | 50 | 208 | 12 | MOLECULE: TETRONASIN; |
| 1385 | 4u67-P | 2.4 | 7 | 63 | 130 | 6 | MOLECULE: 50S RIBOSOMAL PROTEIN L2; |
| 1386 | 5kd2-A | 2.4 | 13.9 | 89 | 591 | 7 | MOLECULE: METALLOPEPTIDASE; |
| 1387 | 6p8v-C | 2.4 | 9.6 | 53 | 303 | 8 | MOLECULE: ATPASE, AAA FAMILY; |
| 1388 | 2fna-A | 2.4 | 12.7 | 67 | 352 | 7 | MOLECULE: CONSERVED HYPOTHETICAL PROTEIN; |
| 1389 | 6gef-A | 2.4 | 3.9 | 63 | 388 | 0 | MOLECULE: TYPE IV SECRETION SYSTEM PROTEIN DOTB; |
| 1390 | 5t58-N | 2.4 | 3.8 | 54 | 196 | 6 | MOLECULE: KLLA0F02343P; |
| 1391 | 3cqx-D | 2.4 | 4.6 | 39 | 84 | 3 | MOLECULE: HEAT SHOCK COGNATE 71 KDA PROTEIN; |
| 1392 | 6wq0-A | 2.4 | 9.4 | 56 | 131 | 4 | MOLECULE: DNA (301-MER); |
| 1393 | 6zie-A | 2.4 | 4.6 | 53 | 123 | 9 | MOLECULE: CMPX-383B; |
| 1394 | 3mka-J | 2.4 | 8.3 | 75 | 252 | 5 | MOLECULE: PROTEASOME SUBUNIT ALPHA; |
| 1395 | 4y9j-B | 2.4 | 8.7 | 81 | 593 | 9 | MOLECULE: PROTEIN ACDH-11, ISOFORM B; |
| 1396 | 4ip8-A | 2.4 | 5.6 | 52 | 105 | 4 | MOLECULE: SERUM AMYLOID A-1 PROTEIN; |
| 1397 | 6scj-A | 2.4 | 7.1 | 89 | 2551 | 8 | MOLECULE: THYROGLOBULIN; |

| | | | | | | | |
|---|---|---|---|---|---|---|---|
| 1398 | 4fz4-A | 2.4 | 9 | 76 | 154 | 13 | MOLECULE: UNCHARACTERIZED PROTEIN CONSERVED IN BACTERIA; |
| 1399 | 1e1d-A | 2.4 | 10.2 | 78 | 553 | 8 | MOLECULE: HYDROXYLAMINE REDUCTASE; |
| 1400 | 3s1e-A | 2.4 | 14.1 | 74 | 499 | 8 | MOLECULE: CYTOKININ DEHYDROGENASE 1; |
| 1401 | 6o3v-B | 2.4 | 5.9 | 79 | 831 | 8 | MOLECULE: PROTEIN VP3; |
| 1402 | 6tnn-I | 2.4 | 3.9 | 73 | 133 | 5 | MOLECULE: 50S RIBOSOMAL PROTEIN L10; |
| 1403 | 4abx-A | 2.4 | 4.1 | 55 | 167 | 7 | MOLECULE: DNA REPAIR PROTEIN RECN; |
| 1404 | 6ewz-A | 2.4 | 11.4 | 66 | 202 | 8 | MOLECULE: GTP PYROPHOSPHOKINASE; |
| 1405 | 1xbn-A | 2.4 | 3.9 | 71 | 195 | 8 | MOLECULE: METHYL-ACCEPTING CHEMOTAXIS PROTEIN; |
| 1406 | 2uui-A | 2.4 | 9.2 | 60 | 155 | 2 | MOLECULE: LEUKOTRIENE C4 SYNTHASE; |
| 1407 | 6edw-B | 2.4 | 14.5 | 81 | 746 | 7 | MOLECULE: ISOCITRATE LYASE 2; |
| 1408 | 5me8-A | 2.4 | 6.2 | 53 | 107 | 9 | MOLECULE: INHIBITOR OF GROWTH PROTEIN 5; |
| 1409 | 7ahd-A | 2.4 | 6.4 | 71 | 547 | 8 | MOLECULE: ABC-TYPE PROLINE/GLYCINE BETAINE TRANSPORT SYSTEM |
| 1410 | 5dku-B | 2.4 | 12 | 73 | 581 | 8 | MOLECULE: PREX DNA POLYMERASE; |
| 1411 | 3d1b-A | 2.4 | 3.6 | 49 | 111 | 8 | MOLECULE: RNA-INDUCED TRANSCRIPTIONAL SILENCING COMPLEX PRO |
| 1412 | 6sku-A | 2.4 | 14.4 | 82 | 785 | 10 | MOLECULE: PHOSPHOCHOLINE TRANSFERASE ANKX; |
| 1413 | 3eps-A | 2.4 | 14.3 | 94 | 566 | 6 | MOLECULE: ISOCITRATE DEHYDROGENASE KINASE/PHOSPHATASE; |
| 1414 | 5hwy-A | 2.4 | 4.2 | 63 | 300 | 5 | MOLECULE: UNCHARACTERIZED MEMBRANE PROTEIN MJ0091; |
| 1415 | 6al9-B | 2.4 | 6 | 68 | 91 | 7 | MOLECULE: CHORISMATE MUTASE; |
| 1416 | 4j41-B | 2.4 | 5.3 | 53 | 86 | 4 | MOLECULE: SECRETED PROTEIN ESXB; |
| 1417 | 2ajq-A | 2.4 | 6.7 | 58 | 704 | 9 | MOLECULE: DNA PRIMER; |
| 1418 | 6ty9-A | 2.4 | 13.2 | 99 | 1208 | 5 | MOLECULE: RNA-DEPENDENT RNA POLYMERASE; |
| 1419 | 2au5-A | 2.4 | 3.7 | 67 | 129 | 10 | MOLECULE: CONSERVED DOMAIN PROTEIN; |
| 1420 | 4f0x-A | 2.4 | 6.4 | 74 | 456 | 3 | MOLECULE: MALONYL-COA DECARBOXYLASE, MITOCHONDRIAL; |
| 1421 | 5cr4-B | 2.4 | 8.5 | 63 | 226 | 10 | MOLECULE: SLEEPING BEAUTY TRANSPOSASE, SB100X; |
| 1422 | 4c00-A | 2.4 | 15.3 | 67 | 544 | 12 | MOLECULE: TRANSLOCATION AND ASSEMBLY MODULE TAMA; |
| 1423 | 5mqf-M | 2.4 | 4.7 | 60 | 706 | 5 | MOLECULE: PRE-MRNA-PROCESSING-SPLICING FACTOR 8; |
| 1424 | 4hga-A | 2.4 | 8.1 | 48 | 207 | 15 | MOLECULE: DEATH DOMAIN-ASSOCIATED PROTEIN 6; |
| 1425 | 5mpd-Z | 2.4 | 15 | 89 | 906 | 2 | MOLECULE: 26S PROTEASOME REGULATORY SUBUNIT RPN10; |
| 1426 | 2raj-A | 2.4 | 5.9 | 91 | 382 | 7 | MOLECULE: SORTING NEXIN-9; |
| 1427 | 6zsi-D | 2.4 | 4.7 | 45 | 133 | 9 | MOLECULE: RAS-RELATED PROTEIN RAB-8A; |
| 1428 | 5ecj-A | 2.4 | 11.1 | 55 | 263 | 4 | MOLECULE: PR DOMAIN ZINC FINGER PROTEIN 14,PROTEIN CBFA2T2; |
| 1429 | 7bgy-C | 2.4 | 3.6 | 45 | 190 | 11 | MOLECULE: POTASSIUM-TRANSPORTING ATPASE POTASSIUM-BINDING S |
| 1430 | 6s1k-A | 2.4 | 13 | 58 | 383 | 7 | MOLECULE: CHEMOTAXIS PROTEIN CHEA; |
| 1431 | 2fm8-C | 2.4 | 14.1 | 58 | 220 | 10 | MOLECULE: SURFACE PRESENTATION OF ANTIGENS PROTEIN SPAK; |
| 1432 | 5xu0-B | 2.4 | 5 | 58 | 231 | 5 | MOLECULE: MEMBRANE-FUSION PROTEIN; |

| | | | | | | | |
|---|---|---|---|---|---|---|---|
| 1433 | 3boy-A | 2.4 | 5.1 | 66 | 150 | 8 | MOLECULE: 5'-R(*UP*UP*UP*AP*GP*UP*UP*UP*UP*UP*AP*GP*UP*UP*U |
| 1434 | 7krw-A | 2.4 | 12.3 | 73 | 608 | 10 | MOLECULE: CHAPERONE PROTEIN DNAK FUSED WITH SUBSTRATE PEPTI |
| 1435 | 3u24-A | 2.4 | 10.3 | 80 | 546 | 5 | MOLECULE: PUTATIVE LIPOPROTEIN; |
| 1436 | 4qes-A | 2.4 | 4.6 | 61 | 441 | 5 | MOLECULE: NON-HAEM BROMOPEROXIDASE BPO-A2, MATRIX PROTEIN 1 |
| 1437 | 7k5c-A | 2.4 | 12.8 | 88 | 650 | 10 | MOLECULE: INTERNAL VIRION PROTEIN GP15; |
| 1438 | 6zyv-A | 2.4 | 7.1 | 57 | 238 | 9 | MOLECULE: CIR PROTEIN; |
| 1439 | 4gl6-B | 2.4 | 3.8 | 48 | 241 | 10 | MOLECULE: HYPOTHETICAL PROTEIN; |
| 1440 | 5lnk-M | 2.4 | 9.7 | 83 | 459 | 14 | MOLECULE: MITOCHONDRIAL COMPLEX I, 51 KDA SUBUNIT; |
| 1441 | 2ex3-B | 2.4 | 11.3 | 50 | 196 | 8 | MOLECULE: DNA POLYMERASE; |
| 1442 | 6f2d-F | 2.4 | 7 | 62 | 258 | 3 | MOLECULE: FLAGELLAR BIOSYNTHETIC PROTEIN FLIP; |
| 1443 | 6yaq-A | 2.4 | 10.5 | 71 | 499 | 8 | MOLECULE: CYTOKININ DEHYDROGENASE 8; |
| 1444 | 7emf-J | 2.4 | 8 | 61 | 122 | 13 | MOLECULE: MEDIATOR OF RNA POLYMERASE II TRANSCRIPTION SUBUN |
| 1445 | 6rax-N | 2.4 | 11 | 69 | 207 | 9 | MOLECULE: DNA REPLICATION LICENSING FACTOR MCM2; |
| 1446 | 6f7s-C | 2.4 | 16.8 | 96 | 319 | 8 | MOLECULE: SERRATE RNA EFFECTOR MOLECULE HOMOLOG; |
| 1447 | 1y6d-A | 2.4 | 3.5 | 59 | 114 | 5 | MOLECULE: PHOSPHORELAY PROTEIN LUXU; |
| 1448 | 7rtm-A | 2.4 | 20.1 | 71 | 571 | 7 | MOLECULE: ELECTRONEUTRAL SODIUM BICARBONATE EXCHANGER 1; |
| 1449 | 3fhf-A | 2.4 | 12.2 | 55 | 214 | 5 | MOLECULE: N-GLYCOSYLASE/DNA LYASE; |
| 1450 | 6grd-A | 2.4 | 14 | 78 | 407 | 14 | MOLECULE: NUCLEASE-LIKE PROTEIN; |
| 1451 | 2hsb-A | 2.4 | 5.9 | 48 | 126 | 13 | MOLECULE: HYPOTHETICAL UPF0332 PROTEIN AF0298; |
| 1452 | 1ij6-A | 2.4 | 3.4 | 54 | 305 | 6 | MOLECULE: PLASMODIAL SPECIFIC LAV1-2 PROTEIN; |
| 1453 | 4cht-B | 2.4 | 7.3 | 58 | 215 | 7 | MOLECULE: DNA TOPOISOMERASE 3-ALPHA; |
| 1454 | 6wg3-A | 2.4 | 29 | 49 | 561 | 2 | MOLECULE: STRUCTURAL MAINTENANCE OF CHROMOSOMES PROTEIN 1A; |
| 1455 | 1sfx-A | 2.4 | 4.2 | 37 | 109 | 8 | MOLECULE: CONSERVED HYPOTHETICAL PROTEIN AF2008; |
| 1456 | 4zua-A | 2.4 | 4.2 | 54 | 155 | 9 | MOLECULE: EXOENZYME S SYNTHESIS REGULATORY PROTEIN EXSA; |
| 1457 | 3bo0-A | 2.4 | 13.9 | 77 | 442 | 4 | MOLECULE: 23S RIBOSOMAL RNA; |
| 1458 | 3p9y-A | 2.4 | 10.3 | 49 | 198 | 6 | MOLECULE: CG14216; |
| 1459 | 4wid-A | 2.4 | 8.5 | 66 | 353 | 11 | MOLECULE: RHUL123; |
| 1460 | 5lsw-A | 2.4 | 8.2 | 75 | 273 | 7 | MOLECULE: CELL DIFFERENTIATION PROTEIN RCD1 HOMOLOG; |
| 1461 | 5ewp-A | 2.4 | 13.1 | 84 | 241 | 6 | MOLECULE: ARO (ARMADILLO REPEATS ONLY PROTEIN); |
| 1462 | 5jp6-A | 2.4 | 5.2 | 56 | 339 | 5 | MOLECULE: PUTATIVE POLYSACCHARIDE DEACETYLASE; |
| 1463 | 3mx2-B | 2.4 | 22.4 | 79 | 517 | 8 | MOLECULE: NUCLEOPROTEIN; |
| 1464 | 6tdv-H | 2.4 | 9.1 | 58 | 388 | 14 | MOLECULE: ATPTB1; |
| 1465 | 3rc8-A | 2.4 | 8.2 | 88 | 609 | 8 | MOLECULE: ATP-DEPENDENT RNA HELICASE SUPV3L1, MITOCHONDRIAL |
| 1466 | 6rth-A | 2.4 | 12 | 80 | 504 | 4 | MOLECULE: RTX TOXIN AND CA2+-BINDING PROTEIN; |
| 1467 | 2ve7-D | 2.4 | 5 | 73 | 242 | 3 | MOLECULE: KINETOCHORE PROTEIN HEC1, KINETOCHORE PROTEIN SPC |

| | | | | | | | |
|---|---|---|---|---|---|---|---|
| 1468 | 2i7x-A | 2.4 | 4.6 | 61 | 499 | 5 | MOLECULE: PROTEIN CFT2; |
| 1469 | 2xeq-C | 2.4 | 12.2 | 68 | 241 | 21 | MOLECULE: PAT1 HOMOLOG 1,; |
| 1470 | 6l82-A | 2.4 | 7.3 | 65 | 98 | 12 | MOLECULE: SPINDLE POLE BODY COMPONENT; |
| 1471 | 2jdi-H | 2.4 | 6.5 | 45 | 88 | 7 | MOLECULE: ATP SYNTHASE SUBUNIT ALPHA HEART ISOFORM; |
| 1472 | 5ijz-K | 2.4 | 7.3 | 65 | 304 | 8 | MOLECULE: NADP-SPECIFIC GLUTAMATE DEHYDROGENASE; |
| 1473 | 6ubz-A | 2.3 | 8.4 | 76 | 792 | 5 | MOLECULE: UNCHARACTERIZED PROTEIN GOXA; |
| 1474 | 4ok7-A | 2.3 | 7.3 | 88 | 223 | 6 | MOLECULE: ENDOLYSIN; |
| 1475 | 2pbi-A | 2.3 | 14.7 | 86 | 415 | 12 | MOLECULE: REGULATOR OF G-PROTEIN SIGNALING 9; |
| 1476 | 2lck-A | 2.3 | 15.7 | 99 | 296 | 5 | MOLECULE: MITOCHONDRIAL UNCOUPLING PROTEIN 2; |
| 1477 | 1qdm-A | 2.3 | 4.1 | 56 | 430 | 4 | MOLECULE: PROPHYTEPSIN; |
| 1478 | 6bbm-D | 2.3 | 8.7 | 65 | 455 | 6 | MOLECULE: REPLICATIVE DNA HELICASE; |
| 1479 | 3wvo-A | 2.3 | 10 | 78 | 543 | 4 | MOLECULE: CRISPR-ASSOCIATED PROTEIN, CSE1 FAMILY; |
| 1480 | 4cth-A | 2.3 | 11.8 | 83 | 698 | 11 | MOLECULE: NEPRILYSIN; |
| 1481 | 6wcw-A | 2.3 | 11.3 | 90 | 247 | 2 | MOLECULE: RAS-RELATED PROTEIN RAB-7A; |
| 1482 | 2ncj-A | 2.3 | 4.2 | 54 | 171 | 19 | MOLECULE: UNCHARACTERIZED PROTEIN; |
| 1483 | 6tpk-A | 2.3 | 4.1 | 69 | 461 | 7 | MOLECULE: OXYTOCIN RECEPTOR; |
| 1484 | 1sf9-A | 2.3 | 5.8 | 65 | 118 | 8 | MOLECULE: YFHH HYPOTHETICAL PROTEIN; |
| 1485 | 1knz-A | 2.3 | 7.1 | 53 | 154 | 6 | MOLECULE: 5'-R(*UP*GP*AP*CP*C)-3'; |
| 1486 | 6hgc-A | 2.3 | 10.4 | 62 | 304 | 5 | MOLECULE: UBIQUITIN CARBOXYL-TERMINAL HYDROLASE CALYPSO,UBI |
| 1487 | 6jbn-A | 2.3 | 7.4 | 66 | 391 | 5 | MOLECULE: PEROXIDASE EFEB; |
| 1488 | 6hk5-B | 2.3 | 4.8 | 36 | 66 | 11 | MOLECULE: COOJ; |
| 1489 | 2q06-A | 2.3 | 5.8 | 71 | 467 | 6 | MOLECULE: NUCLEOPROTEIN; |
| 1490 | 7c79-B | 2.3 | 4 | 54 | 793 | 7 | MOLECULE: RIBONUCLEASE MRP RNA SUBUNIT NME1; |
| 1491 | 6ymw-A | 2.3 | 10.7 | 87 | 913 | 14 | MOLECULE: MITOCHONDRIAL TRANSCRIPTION FACTOR 1; |
| 1492 | 2a3l-A | 2.3 | 5.5 | 50 | 616 | 6 | MOLECULE: AMP DEAMINASE; |
| 1493 | 4aup-B | 2.3 | 4.6 | 54 | 123 | 7 | MOLECULE: PHOSPHOLIPASE A2 GROUP XIII; |
| 1494 | 2n39-A | 2.3 | 4.7 | 60 | 108 | 8 | MOLECULE: CHROMODOMAIN-HELICASE-DNA-BINDING PROTEIN 1; |
| 1495 | 6z6e-A | 2.3 | 9.4 | 57 | 104 | 9 | MOLECULE: TERMINASE SMALL SUBUNIT; |
| 1496 | 2xwb-F | 2.3 | 15.5 | 71 | 714 | 6 | MOLECULE: COMPLEMENT C3B BETA CHAIN; |
| 1497 | 6bfi-B | 2.3 | 6.5 | 79 | 806 | 5 | MOLECULE: VIN1; |
| 1498 | 4cgz-A | 2.3 | 10.9 | 61 | 629 | 8 | MOLECULE: BLOOM'S SYNDROME HELICASE; |
| 1499 | 1rfz-A | 2.3 | 3.8 | 70 | 164 | 9 | MOLECULE: HYPOTHETICAL PROTEIN APC35681; |
| 1500 | 3ont-A | 2.3 | 8.1 | 65 | 112 | 8 | MOLECULE: SPOT 14 PROTEIN; |
| 1501 | 3jd5-b | 2.3 | 13.1 | 51 | 135 | 8 | MOLECULE: 28S RIBOSOMAL RNA, MITOCHONDIAL; |
| 1502 | 6jhm-A | 2.3 | 11.5 | 97 | 494 | 5 | MOLECULE: CHLOROPHENOL MONOOXYGENASE; |

| | | | | | | | |
|---|---|---|---|---|---|---|---|
| 1503 | 3l9f-A | 2.3 | 6.6 | 62 | 170 | 5 | MOLECULE: PUTATIVE UNCHARACTERIZED PROTEIN SMU.1604C; |
| 1504 | 5uke-A | 2.3 | 7.6 | 78 | 124 | 10 | MOLECULE: INTERLEUKIN-1 RECEPTOR-ASSOCIATED KINASE 3; |
| 1505 | 5ip0-D | 2.3 | 3.3 | 41 | 108 | 15 | MOLECULE: PHA GRANULE-ASSOCIATED PROTEIN; |
| 1506 | 7die-A | 2.3 | 3.4 | 46 | 164 | 2 | MOLECULE: FERRITIN; |
| 1507 | 5kko-C | 2.3 | 6.3 | 46 | 55 | 7 | MOLECULE: UNCHARACTERISED PROTEIN; |
| 1508 | 6vin-A | 2.3 | 11.6 | 83 | 363 | 5 | MOLECULE: THREONINE ASPARTASE 1; |
| 1509 | 4x8d-A | 2.3 | 4.3 | 66 | 429 | 5 | MOLECULE: SULFOXIDE SYNTHASE EGTB; |
| 1510 | 1tu3-J | 2.3 | 3.2 | 35 | 53 | 11 | MOLECULE: RAS-RELATED PROTEIN RAB-5A; |
| 1511 | 6lea-C | 2.3 | 4.8 | 54 | 126 | 13 | MOLECULE: FLAGELLAR SECRETION CHAPERONE FLIS; |
| 1512 | 5nmo-A | 2.3 | 5.5 | 57 | 162 | 14 | MOLECULE: CHROMOSOME PARTITION PROTEIN SMC,CHROMOSOME |
| 1513 | 2p22-C | 2.3 | 5.5 | 49 | 186 | 10 | MOLECULE: SUPPRESSOR PROTEIN STP22 OF TEMPERATURE-SENSITIVE |
| 1514 | 1w6j-A | 2.3 | 14.8 | 95 | 727 | 5 | MOLECULE: LANOSTEROL SYNTHASE; |
| 1515 | 3c46-A | 2.3 | 7.3 | 102 | 1095 | 10 | MOLECULE: VIRION RNA POLYMERASE; |
| 1516 | 6uxe-B | 2.3 | 4.2 | 63 | 85 | 6 | MOLECULE: CYSTEINE DESULFURASE, MITOCHONDRIAL; |
| 1517 | 4dnr-A | 2.3 | 10.6 | 103 | 1031 | 8 | MOLECULE: CATION EFFLUX SYSTEM PROTEIN CUSB; |
| 1518 | 6ef3-A | 2.3 | 13.2 | 68 | 247 | 12 | MOLECULE: PROTEASOME SUBUNIT BETA TYPE-1; |
| 1519 | 3kyj-A | 2.3 | 4.3 | 64 | 129 | 11 | MOLECULE: PUTATIVE HISTIDINE PROTEIN KINASE; |
| 1520 | 6wq2-A | 2.3 | 6.8 | 60 | 154 | 10 | MOLECULE: A-DNA; |
| 1521 | 5cwb-A | 2.3 | 3.8 | 70 | 197 | 13 | MOLECULE: DESIGNED HELICAL REPEAT PROTEIN; |
| 1522 | 6zr2-d | 2.3 | 4.9 | 56 | 120 | 2 | MOLECULE: NADH-UBIQUINONE OXIDOREDUCTASE CHAIN 3; |
| 1523 | 5y69-A | 2.3 | 8.8 | 67 | 147 | 10 | MOLECULE: CHAIN A; |
| 1524 | 2pyq-A | 2.3 | 8 | 47 | 114 | 6 | MOLECULE: UNCHARACTERIZED PROTEIN; |
| 1525 | 6csv-A | 2.3 | 5.1 | 52 | 90 | 8 | MOLECULE: CENTROSOMAL PROTEIN OF 63 KDA,CENTROSOMAL PROTEIN |
| 1526 | 6oap-A | 2.3 | 10.5 | 76 | 305 | 8 | MOLECULE: DUAL SENSOR HISTIDINE KINASE; |
| 1527 | 6rkw-A | 2.3 | 10.4 | 66 | 828 | 3 | MOLECULE: DNA GYRASE SUBUNIT A; |
| 1528 | 6ptg-B | 2.3 | 6.9 | 49 | 86 | 4 | MOLECULE: DNAK SUPPRESSOR; |
| 1529 | 3q4i-A | 2.3 | 12.9 | 69 | 204 | 9 | MOLECULE: PHOSPHOHYDROLASE (MUTT/NUDIX FAMILY PROTEIN); |
| 1530 | 7lmz-C | 2.3 | 10.7 | 66 | 752 | 5 | MOLECULE: TRANSITIONAL ENDOPLASMIC RETICULUM ATPASE; |
| 1531 | 6jx7-A | 2.3 | 9.1 | 80 | 1245 | 11 | MOLECULE: FELINE INFECTIOUS PERITONITIS VIRUS SPIKE PROTEIN |
| 1532 | 7bwk-A | 2.3 | 7 | 57 | 113 | 4 | MOLECULE: ICMO (DOTL); |
| 1533 | 3ke3-A | 2.3 | 9.8 | 58 | 371 | 12 | MOLECULE: PUTATIVE SERINE-PYRUVATE AMINOTRANSFERASE; |
| 1534 | 7b6d-D | 2.3 | 3.1 | 47 | 219 | 2 | MOLECULE: TRAFFICKING PROTEIN PARTICLE COMPLEX SUBUNIT; |
| 1535 | 3j9a-A | 2.3 | 7.3 | 66 | 81 | 9 | MOLECULE: CAPSID PROTEIN VP26 HOMOLOG; |
| 1536 | 3fey-A | 2.3 | 5.2 | 76 | 761 | 8 | MOLECULE: NUCLEAR CAP-BINDING PROTEIN SUBUNIT 1; |
| 1537 | 7ljn-A | 2.3 | 12.6 | 80 | 342 | 4 | MOLECULE: CD-NTASE; |

| | | | | | | | |
|---|---|---|---|---|---|---|---|
| 1538 | 3l0i-C | 2.3 | 4.2 | 49 | 322 | 8 | MOLECULE: DRRA; |
| 1539 | 6rwx-A | 2.3 | 7.6 | 59 | 189 | 14 | MOLECULE: PROTEIN MXIG; |
| 1540 | 6sjl-A | 2.3 | 5.4 | 53 | 324 | 8 | MOLECULE: PUTATIVE TYPE VI SECRETION PROTEIN; |
| 1541 | 4xwj-A | 2.3 | 3.8 | 66 | 153 | 9 | MOLECULE: REGULATOR OF SIGMA D; |
| 1542 | 6l80-C | 2.3 | 7.5 | 49 | 107 | 6 | MOLECULE: GAMMA-TUBULIN COMPLEX SUBUNIT MOD21; |
| 1543 | 5j6f-A | 2.3 | 6.5 | 77 | 352 | 9 | MOLECULE: 3-DEOXY-D-ARABINO-HEPTULOSONATE 7-PHOSPHATE SYNTH |
| 1544 | 6r6n-A | 2.3 | 3.7 | 58 | 111 | 7 | MOLECULE: SMALL SOLUBLE CYT C; |
| 1545 | 3ck9-B | 2.3 | 7.4 | 87 | 515 | 5 | MOLECULE: SUSD; |
| 1546 | 1zs3-A | 2.3 | 10.3 | 66 | 171 | 8 | MOLECULE: LACTOCOCCUS LACTIS MG1363 DPSA; |
| 1547 | 1yoz-B | 2.3 | 6 | 48 | 116 | 0 | MOLECULE: HYPOTHETICAL PROTEIN AF0941; |
| 1548 | 6ahp-A | 2.3 | 2.3 | 46 | 110 | 9 | MOLECULE: FLAGELLAR PROTEIN FLIL; |
| 1549 | 2db7-A | 2.3 | 4 | 36 | 57 | 6 | MOLECULE: HAIRY/ENHANCER-OF-SPLIT RELATED WITH YRPW MOTIF |
| 1550 | 4r04-A | 2.3 | 20.2 | 92 | 1793 | 9 | MOLECULE: TOXIN A; |
| 1551 | 7ls0-B | 2.3 | 4.5 | 69 | 362 | 10 | MOLECULE: ALK TYROSINE KINASE RECEPTOR FUSED WITH ALK AND L |
| 1552 | 6ero-A | 2.3 | 3.7 | 64 | 296 | 9 | MOLECULE: DIMETHYLADENOSINE TRANSFERASE 2, MITOCHONDRIAL, |
| 1553 | 4xk8-k | 2.3 | 2.6 | 36 | 46 | 6 | MOLECULE: PHOTOSYSTEM I P700 CHLOROPHYLL A APOPROTEIN A1; |
| 1554 | 3izq-1 | 2.3 | 13.1 | 55 | 516 | 5 | MOLECULE: PROTEIN DOM34; |
| 1555 | 7kfl-A | 2.3 | 4.4 | 66 | 356 | 14 | MOLECULE: MYOSIN-17; |
| 1556 | 3t3o-A | 2.3 | 5.9 | 57 | 553 | 11 | MOLECULE: METAL DEPENDENT HYDROLASE; |
| 1557 | 5xmg-A | 2.3 | 7.7 | 64 | 344 | 6 | MOLECULE: UNCHARACTERIZED PROTEIN; |
| 1558 | 1g4u-S | 2.3 | 10.3 | 70 | 360 | 13 | MOLECULE: PROTEIN TYROSINE PHOSPHATASE SPTP; |
| 1559 | 4roe-A | 2.3 | 10.6 | 60 | 307 | 7 | MOLECULE: TRANSCRIPTION FACTOR IIIB 50 KDA SUBUNIT; |
| 1560 | 5azm-A | 2.3 | 10.8 | 62 | 254 | 5 | MOLECULE: N-ACETYLGLUCOSAMINIDASE; |
| 1561 | 6v6d-A | 2.3 | 4.2 | 60 | 209 | 10 | MOLECULE: PANNEXIN-1; |
| 1562 | 7ole-K | 2.3 | 5 | 44 | 394 | 5 | MOLECULE: RUVB-LIKE 1; |
| 1563 | 1zpv-C | 2.3 | 4.9 | 48 | 88 | 10 | MOLECULE: ACT DOMAIN PROTEIN; |
| 1564 | 3b77-A | 2.3 | 8.9 | 54 | 188 | 6 | MOLECULE: UNCHARACTERIZED PROTEIN; |
| 1565 | 5k7l-A | 2.3 | 13 | 51 | 701 | 6 | MOLECULE: POTASSIUM VOLTAGE-GATED CHANNEL SUBFAMILY H MEMBE |
| 1566 | 1zp2-A | 2.3 | 4.6 | 70 | 227 | 9 | MOLECULE: RNA POLYMERASE II HOLOENZYME CYCLIN-LIKE SUBUNIT; |
| 1567 | 1gh6-A | 2.3 | 5.2 | 54 | 114 | 9 | MOLECULE: LARGE T ANTIGEN; |
| 1568 | 2js1-A | 2.3 | 5.3 | 51 | 80 | 8 | MOLECULE: UNCHARACTERIZED PROTEIN YVFG; |
| 1569 | 6m3q-F | 2.3 | 11.1 | 86 | 301 | 8 | MOLECULE: ANKYRIN-2; |
| 1570 | 2qby-B | 2.3 | 7.5 | 61 | 368 | 10 | MOLECULE: CELL DIVISION CONTROL PROTEIN 6 HOMOLOG 1; |
| 1571 | 5tsz-A | 2.3 | 4.9 | 54 | 130 | 4 | MOLECULE: PV CELL-TRAVERSAL PROTEIN; |
| 1572 | 6vwb-A | 2.3 | 3.5 | 50 | 90 | 8 | MOLECULE: CROSSOVER JUNCTION ENDONUCLEASE MUS81; |

| | | | | | | | |
|---|---|---|---|---|---|---|---|
| 1573 | 3nqw-A | 2.3 | 5.1 | 46 | 178 | 9 | MOLECULE: CG11900; |
| 1574 | 6o9l-3 | 2.3 | 4.3 | 54 | 309 | 6 | MOLECULE: DNA-DIRECTED RNA POLYMERASE II SUBUNIT RPB1; |
| 1575 | 5w7p-A | 2.3 | 14.5 | 73 | 397 | 7 | MOLECULE: OXAC; |
| 1576 | 4wai-A | 2.3 | 4.3 | 49 | 91 | 14 | MOLECULE: COMF OPERON PROTEIN 2; |
| 1577 | 2pff-A | 2.3 | 8.5 | 68 | 1683 | 7 | MOLECULE: FATTY ACID SYNTHASE SUBUNIT ALPHA; |
| 1578 | 5lsk-D | 2.3 | 3.7 | 44 | 178 | 5 | MOLECULE: PROTEIN MIS12 HOMOLOG; |
| 1579 | 4eek-A | 2.3 | 6 | 64 | 229 | 5 | MOLECULE: BETA-PHOSPHOGLUCOMUTASE-RELATED PROTEIN; |
| 1580 | 5l4k-P | 2.3 | 15.7 | 78 | 456 | 5 | MOLECULE: 26S PROTEASOME NON-ATPASE REGULATORY SUBUNIT 4; |
| 1581 | 2rdc-B | 2.3 | 5.3 | 64 | 135 | 8 | MOLECULE: UNCHARACTERIZED PROTEIN; |
| 1582 | 2e87-A | 2.3 | 6.6 | 82 | 356 | 9 | MOLECULE: HYPOTHETICAL PROTEIN PH1320; |
| 1583 | 5i6c-A | 2.3 | 7.1 | 98 | 480 | 10 | MOLECULE: URIC ACID-XANTHINE PERMEASE; |
| 1584 | 4ozq-B | 2.3 | 11.1 | 81 | 687 | 6 | MOLECULE: CHIMERA OF MALTOSE-BINDING PERIPLASMIC PROTEIN AN |
| 1585 | 4y7s-A | 2.3 | 3.2 | 60 | 111 | 7 | MOLECULE: SURFACE ANTIGEN PROTEIN 2; |
| 1586 | 5j9q-D | 2.3 | 8.3 | 52 | 120 | 8 | MOLECULE: HISTONE ACETYLTRANSFERASE ESA1; |
| 1587 | 5xsj-L | 2.3 | 5.9 | 58 | 122 | 9 | MOLECULE: PERIPLASMIC BINDING PROTEIN/LACI TRANSCRIPTIONAL |
| 1588 | 6lkc-B | 2.3 | 14.1 | 86 | 533 | 6 | MOLECULE: POLYUNSATURATED FATTY ACID SYNTHASE PFAD; |
| 1589 | 5jcp-A | 2.3 | 15.3 | 82 | 360 | 6 | MOLECULE: ARF-GAP WITH RHO-GAP DOMAIN, ANK REPEAT AND PH DO |
| 1590 | 5x3q-A | 2.3 | 8 | 72 | 313 | 6 | MOLECULE: ENVELOPE GLYCOPROTEIN; |
| 1591 | 2ffl-A | 2.3 | 13.5 | 87 | 732 | 6 | MOLECULE: DICER; |
| 1592 | 2f22-B | 2.3 | 3.8 | 60 | 143 | 2 | MOLECULE: BH3987; |
| 1593 | 2qnd-B | 2.3 | 2.7 | 39 | 143 | 13 | MOLECULE: FMR1 PROTEIN; |
| 1594 | 1sed-A | 2.3 | 2.9 | 45 | 112 | 16 | MOLECULE: HYPOTHETICAL PROTEIN YHAI; |
| 1595 | 6tdw-H | 2.3 | 9.1 | 58 | 249 | 14 | MOLECULE: ATPTB1; |
| 1596 | 5jbr-A | 2.3 | 4.2 | 41 | 149 | 7 | MOLECULE: UNCHARACTERIZED PROTEIN BCAV_2135; |
| 1597 | 2m5z-A | 2.3 | 2.9 | 40 | 44 | 13 | MOLECULE: ENTEROCIN JSA; |
| 1598 | 5tj5-A | 2.3 | 6.8 | 56 | 570 | 4 | MOLECULE: V-TYPE PROTON ATPASE SUBUNIT A; |
| 1599 | 5lox-1 | 2.3 | 15.2 | 68 | 241 | 0 | MOLECULE: PEPTIDASE; |
| 1600 | 4ycz-B | 2.3 | 7.7 | 81 | 590 | 7 | MOLECULE: FUSION PROTEIN OF SEC13 AND NUP145C; |
| 1601 | 5oeu-A | 2.3 | 8.3 | 100 | 460 | 7 | MOLECULE: GLUTATHIONE SYNTHETASE-LIKE EFFECTOR 22 (GPA-GSS2 |
| 1602 | 3ct9-A | 2.3 | 5 | 61 | 346 | 8 | MOLECULE: ACETYLORNITHINE DEACETYLASE; |
| 1603 | 6xwx-B | 2.3 | 5.9 | 53 | 84 | 6 | MOLECULE: UNCHARACTERIZED PROTEIN,UNCHARACTERIZED PROTEIN; |
| 1604 | 5vyk-C | 2.3 | 8 | 59 | 203 | 12 | MOLECULE: CHIMERA PROTEIN OF BRS DOMAIN OF BRAF AND CC-SAM |
| 1605 | 4hlq-C | 2.3 | 4.1 | 66 | 281 | 9 | MOLECULE: TBC1 DOMAIN FAMILY MEMBER 20; |
| 1606 | 6au8-A | 2.3 | 14.6 | 65 | 280 | 5 | MOLECULE: GOLGI TO ER TRAFFIC PROTEIN 4 HOMOLOG; |
| 1607 | 4zvc-A | 2.3 | 6.1 | 61 | 116 | 8 | MOLECULE: DIGUANYLATE CYCLASE DOSC; |

| | | | | | | | |
|---|---|---|---|---|---|---|---|
| 1608 | 6cs2-C | 2.3 | 11.8 | 83 | 893 | 13 | MOLECULE: SPIKE GLYCOPROTEIN,FIBRITIN; |
| 1609 | 6hua-B | 2.3 | 5.8 | 47 | 229 | 9 | MOLECULE: UNCHARACTERIZED PROTEIN; |
| 1610 | 2fbq-A | 2.3 | 11.5 | 57 | 213 | 7 | MOLECULE: PROBABLE TRANSCRIPTIONAL REGULATOR; |
| 1611 | 7p5v-B | 2.3 | 4.8 | 75 | 732 | 5 | MOLECULE: VOLUME-REGULATED ANION CHANNEL SUBUNIT LRRC8A; |
| 1612 | 5tfp-A | 2.3 | 6.3 | 47 | 59 | 4 | MOLECULE: HISTONE-LYSINE N-METHYLTRANSFERASE SETDB2; |
| 1613 | 5kp7-B | 2.3 | 2.8 | 43 | 79 | 21 | MOLECULE: CURD; |
| 1614 | 3ogi-B | 2.3 | 5.3 | 55 | 89 | 5 | MOLECULE: PUTATIVE ESAT-6-LIKE PROTEIN 6; |
| 1615 | 2x0l-A | 2.3 | 4.9 | 55 | 670 | 9 | MOLECULE: LYSINE-SPECIFIC HISTONE DEMETHYLASE 1; |
| 1616 | 6j1x-B | 2.3 | 8 | 71 | 472 | 10 | MOLECULE: NEDD4-LIKE E3 UBIQUITIN-PROTEIN LIGASE WWP1; |
| 1617 | 6psi-A | 2.3 | 10.2 | 70 | 280 | 7 | MOLECULE: CHAPERONE PROTEIN DNAJ 2; |
| 1618 | 6wuc-I | 2.3 | 8.2 | 77 | 713 | 4 | MOLECULE: INNER KINETOCHORE SUBUNIT MCM16; |
| 1619 | 7ar9-J | 2.3 | 4 | 47 | 145 | 9 | MOLECULE: ND3; |
| 1620 | 2pms-C | 2.3 | 5 | 46 | 109 | 13 | MOLECULE: LACTOTRANSFERRIN; |
| 1621 | 2pc1-A | 2.3 | 10 | 58 | 173 | 7 | MOLECULE: ACETYLTRANSFERASE, GNAT FAMILY; |
| 1622 | 4wfc-B | 2.3 | 10.6 | 69 | 118 | 9 | MOLECULE: EXOSOME COMPLEX EXONUCLEASE RRP6; |
| 1623 | 3bzk-A | 2.3 | 5.4 | 48 | 728 | 8 | MOLECULE: TEX; |
| 1624 | 5od9-A | 2.3 | 8.8 | 54 | 95 | 7 | MOLECULE: MID1SC9; |
| 1625 | 5an3-B | 2.3 | 6.4 | 53 | 136 | 9 | MOLECULE: SGT1; |
| 1626 | 1kdo-A | 2.3 | 7.8 | 60 | 223 | 8 | MOLECULE: CYTIDYLATE KINASE; |
| 1627 | 5l09-B | 2.3 | 4.2 | 43 | 164 | 12 | MOLECULE: QUORUM-SENSING TRANSCRIPTIONAL ACTIVATOR; |
| 1628 | 6w1s-Y | 2.3 | 9.4 | 61 | 132 | 8 | MOLECULE: MEDIATOR OF RNA POLYMERASE II TRANSCRIPTION SUBUN |
| 1629 | 6cuq-B | 2.3 | 3.3 | 47 | 117 | 6 | MOLECULE: MACROPHAGE MIGRATION INHIBITORY FACTOR-LIKE PROTE |
| 1630 | 4fym-D | 2.3 | 12.2 | 65 | 227 | 12 | MOLECULE: OROTATE PHOSPHORIBOSYLTRANSFERASE; |
| 1631 | 5wah-A | 2.3 | 6.4 | 67 | 99 | 9 | MOLECULE: IGA FC RECEPTOR; |
| 1632 | 6l7r-A | 2.3 | 8.8 | 61 | 105 | 5 | MOLECULE: PUTATIVE SPINDLE POLE BODY COMPONENT ALP6 PROTEIN |
| 1633 | 4l8k-A | 2.3 | 8.4 | 86 | 315 | 6 | MOLECULE: PUTATIVE PEPTIDASE; |
| 1634 | 2ex5-A | 2.3 | 14.9 | 57 | 207 | 11 | MOLECULE: I-CEUI DNA TARGET SITE; |
| 1635 | 3fmt-A | 2.3 | 3.3 | 47 | 162 | 15 | MOLECULE: PROTEIN SEQA; |
| 1636 | 3ay5-A | 2.3 | 10.7 | 96 | 298 | 9 | MOLECULE: CYCLIN-D1-BINDING PROTEIN 1; |
| 1637 | 4oyd-B | 2.3 | 3.9 | 46 | 117 | 9 | MOLECULE: APOPTOSIS REGULATOR BHRF1; |
| 1638 | 1jr3-E | 2.3 | 14.7 | 69 | 334 | 10 | MOLECULE: DNA POLYMERASE III SUBUNIT GAMMA; |
| 1639 | 6btm-C | 2.3 | 4.4 | 62 | 457 | 10 | MOLECULE: ALTERNATIVE COMPLEX III SUBUNIT A; |
| 1640 | 4hr1-B | 2.3 | 4.7 | 61 | 119 | 10 | MOLECULE: PUTATIVE UNCHARACTERIZED PROTEIN; |
| 1641 | 6zfw-A | 2.3 | 3.1 | 39 | 70 | 3 | MOLECULE: PEROXIN-14; |
| 1642 | 6tg9-D | 2.3 | 2.9 | 46 | 69 | 0 | MOLECULE: FORMATE DEHYDROGENASE SUBUNIT ALPHA; |

| | | | | | | | |
|---|---|---|---|---|---|---|---|
| 1643 | 4ec5-A | 2.3 | 13.2 | 69 | 218 | 10 | MOLECULE: GENERAL SECRETION PATHWAY PROTEIN D; |
| 1644 | 5nv9-A | 2.3 | 4.6 | 55 | 480 | 2 | MOLECULE: PUTATIVE SODIUM:SOLUTE SYMPORTER; |
| 1645 | 1t7s-A | 2.3 | 7.8 | 68 | 129 | 6 | MOLECULE: BAG-1 COCHAPERONE; |
| 1646 | 5a0j-A | 2.3 | 9.3 | 76 | 311 | 8 | MOLECULE: LABDANE-RELATED DITERPENE SYNTHASE; |
| 1647 | 5b86-A | 2.3 | 8.4 | 85 | 579 | 9 | MOLECULE: TUMOR NECROSIS FACTOR ALPHA-INDUCED PROTEIN 2; |
| 1648 | 6u45-A | 2.3 | 15 | 99 | 726 | 9 | MOLECULE: ELONGATION FACTOR 2; |
| 1649 | 6mdm-D | 2.3 | 11.1 | 80 | 713 | 4 | MOLECULE: VESICLE-FUSING ATPASE; |
| 1650 | 6yxq-A | 2.3 | 5.6 | 81 | 188 | 4 | MOLECULE: ACTIVATING SIGNAL COINTEGRATOR 1 COMPLEX SUBUNIT |
| 1651 | 4hte-A | 2.3 | 12.2 | 71 | 340 | 10 | MOLECULE: NICKING ENZYME; |
| 1652 | 1ii0-B | 2.3 | 3.5 | 53 | 553 | 4 | MOLECULE: ARSENICAL PUMP-DRIVING ATPASE; |
| 1653 | 1fxk-C | 2.3 | 6.5 | 60 | 133 | 7 | MOLECULE: PREFOLDIN; |
| 1654 | 3g9g-A | 2.3 | 10.7 | 66 | 250 | 12 | MOLECULE: SUPPRESSOR OF YEAST PROFILIN DELETION; |
| 1655 | 3trc-A | 2.3 | 4.1 | 53 | 168 | 9 | MOLECULE: PHOSPHOENOLPYRUVATE-PROTEIN PHOSPHOTRANSFERASE; |
| 1656 | 3fp5-A | 2.3 | 3.3 | 63 | 106 | 5 | MOLECULE: ACYL-COA BINDING PROTEIN; |
| 1657 | 3plt-A | 2.3 | 2.9 | 51 | 214 | 14 | MOLECULE: SPHINGOLIPID LONG CHAIN BASE-RESPONSIVE PROTEIN L |
| 1658 | 7c7e-A | 2.3 | 5 | 61 | 142 | 5 | MOLECULE: PUTATIVE DNA-BINDING TRANSCRIPTIONAL REGULATOR; |
| 1659 | 5aww-G | 2.3 | 5.6 | 46 | 75 | 9 | MOLECULE: PROTEIN TRANSLOCASE SUBUNIT SECY; |
| 1660 | 6r7i-D | 2.3 | 13.1 | 84 | 407 | 7 | MOLECULE: COP9 SIGNALOSOME COMPLEX SUBUNIT 1; |
| 1661 | 3i0o-A | 2.3 | 8.9 | 82 | 329 | 4 | MOLECULE: SPECTINOMYCIN PHOSPHOTRANSFERASE; |
| 1662 | 6w17-H | 2.3 | 6.6 | 54 | 132 | 7 | MOLECULE: ACTIN-RELATED PROTEIN 3; |
| 1663 | 7nsc-H | 2.3 | 9.7 | 83 | 158 | 10 | MOLECULE: RAN-BINDING PROTEIN 9; |
| 1664 | 5h69-A | 2.3 | 5 | 48 | 252 | 4 | MOLECULE: CHROMOSOME PARTITION PROTEIN SMC; |
| 1665 | 5n3u-B | 2.3 | 10.7 | 59 | 170 | 12 | MOLECULE: PHYCOCYANOBILIN LYASE SUBUNIT ALPHA; |
| 1666 | 3bqz-A | 2.3 | 12.1 | 71 | 186 | 8 | MOLECULE: HTH-TYPE TRANSCRIPTIONAL REGULATOR QACR; |
| 1667 | 6o3q-A | 2.3 | 4.9 | 39 | 49 | 8 | MOLECULE: VICILIN; |
| 1668 | 7d3u-F | 2.3 | 4.5 | 46 | 85 | 9 | MOLECULE: MONOVALENT NA+/H+ ANTIPORTER SUBUNIT D; |
| 1669 | 3jvo-G | 2.3 | 3.9 | 49 | 97 | 6 | MOLECULE: GP6; |
| 1670 | 1osd-A | 2.3 | 3.8 | 44 | 72 | 7 | MOLECULE: HYPOTHETICAL PROTEIN MERP; |
| 1671 | 6es9-A | 2.3 | 11.2 | 84 | 545 | 8 | MOLECULE: ACYL-COA DEHYDROGENASE; |
| 1672 | 6qg0-G | 2.3 | 11.7 | 62 | 355 | 5 | MOLECULE: TRANSLATION INITIATION FACTOR EIF-2B SUBUNIT ALPH |
| 1673 | 4bjm-A | 2.3 | 8.4 | 66 | 226 | 6 | MOLECULE: AVRM; |
| 1674 | 5n9y-A | 2.3 | 3.3 | 41 | 327 | 10 | MOLECULE: ZINC TRANSPORT PROTEIN ZNTB; |
| 1675 | 5xq3-A | 2.3 | 17.1 | 78 | 901 | 5 | MOLECULE: PCRGLX PROTEIN; |
| 1676 | 6f2d-G | 2.3 | 3.3 | 44 | 89 | 5 | MOLECULE: FLAGELLAR BIOSYNTHETIC PROTEIN FLIP; |
| 1677 | 1e7p-A | 2.3 | 10.3 | 67 | 655 | 1 | MOLECULE: FUMARATE REDUCTASE FLAVOPROTEIN SUBUNIT; |

| | | | | | | | |
|---|---|---|---|---|---|---|---|
| 1678 | 2kru-A | 2.3 | 3.8 | 47 | 63 | 13 | MOLECULE: LIGHT-INDEPENDENT PROTOCHLOROPHYLLIDE REDUCTASE S |
| 1679 | 6nd4-Q | 2.3 | 8 | 63 | 862 | 2 | MOLECULE: ETS RRNA; |
| 1680 | 5z51-A | 2.3 | 6.1 | 46 | 161 | 9 | MOLECULE: DNA PRIMASE; |
| 1681 | 5k94-A | 2.3 | 11.3 | 57 | 506 | 14 | MOLECULE: MALTOSE-BINDING PERIPLASMIC PROTEIN,PROTEIN TRANS |
| 1682 | 7pow-B | 2.3 | 4.9 | 60 | 205 | 5 | MOLECULE: CDP-DIACYLGLYCEROL--SERINE O-PHOSPHATIDYLTRANSFER |
| 1683 | 7ar9-M | 2.3 | 10.3 | 93 | 438 | 10 | MOLECULE: ND3; |
| 1684 | 3gwl-A | 2.3 | 5.3 | 59 | 106 | 7 | MOLECULE: FAD-LINKED SULFHYDRYL OXIDASE; |
| 1685 | 5zki-B | 2.3 | 8.8 | 49 | 304 | 8 | MOLECULE: NUCLEASE EXOG, MITOCHONDRIAL; |
| 1686 | 1a8r-A | 2.3 | 11.9 | 76 | 221 | 7 | MOLECULE: GTP CYCLOHYDROLASE I; |
| 1687 | 6xmg-A | 2.3 | 9.7 | 77 | 638 | 6 | MOLECULE: CRISPR-CAS; |
| 1688 | 5lc5-n | 2.3 | 10.3 | 63 | 166 | 8 | MOLECULE: NADH-UBIQUINONE OXIDOREDUCTASE CHAIN 3; |
| 1689 | 7ns4-b | 2.3 | 8 | 59 | 340 | 3 | MOLECULE: E3 UBIQUITIN-PROTEIN LIGASE RMD5; |
| 1690 | 7a46-A | 2.3 | 9.7 | 66 | 223 | 11 | MOLECULE: PUTATIVE TRANSPORT PROTEIN; |
| 1691 | 6swy-5 | 2.3 | 13.3 | 91 | 789 | 9 | MOLECULE: VACUOLAR IMPORT AND DEGRADATION PROTEIN 28; |
| 1692 | 5izo-A | 2.3 | 7.2 | 61 | 290 | 10 | MOLECULE: BIFUNCTIONAL OLIGORIBONUCLEASE AND PAP PHOSPHATAS |
| 1693 | 7f3h-D | 2.3 | 3.4 | 61 | 464 | 8 | MOLECULE: BIFUNCTIONAL CYTOCHROME P450/NADPH--P450 REDUCTAS |
| 1694 | 5mmc-A | 2.3 | 2.2 | 35 | 70 | 3 | MOLECULE: PEROXIN 14; |
| 1695 | 6z7p-A | 2.3 | 8.5 | 75 | 1025 | 4 | MOLECULE: S-LAYER PROTEIN; |
| 1696 | 4ily-B | 2.3 | 12.4 | 76 | 250 | 7 | MOLECULE: CHITOSANASE; |
| 1697 | 6s6y-L | 2.3 | 4.1 | 61 | 311 | 3 | MOLECULE: FORMYLMETHANOFURAN DEHYDROGENASE SUBUNIT A; |
| 1698 | 6a5n-A | 2.3 | 14 | 84 | 502 | 4 | MOLECULE: HISTONE-LYSINE N-METHYLTRANSFERASE, H3 LYSINE-9 S |
| 1699 | 6jo5-K | 2.3 | 3.3 | 41 | 45 | 2 | MOLECULE: PHOTOSYSTEM I P700 CHLOROPHYLL A APOPROTEIN A1; |
| 1700 | 6oce-A | 2.3 | 8.3 | 82 | 689 | 7 | MOLECULE: STRESS-GATED CATION CHANNEL 1.2; |
| 1701 | 5i0n-A | 2.3 | 16.8 | 100 | 489 | 6 | MOLECULE: PHOSPHATIDYLINOSITOL 4-KINASE TYPE 2-ALPHA,LYSOZY |
| 1702 | 4o5p-A | 2.2 | 10.7 | 74 | 742 | 1 | MOLECULE: UNCHARACTERIZED PROTEIN; |
| 1703 | 1u96-A | 2.2 | 3 | 45 | 69 | 9 | MOLECULE: CYTOCHROME C OXIDASE COPPER CHAPERONE; |
| 1704 | 2qnl-A | 2.2 | 6.1 | 59 | 162 | 5 | MOLECULE: UNCHARACTERIZED PROTEIN; |
| 1705 | 1z0x-A | 2.2 | 12.8 | 67 | 216 | 10 | MOLECULE: TRANSCRIPTIONAL REGULATOR, TETR FAMILY; |
| 1706 | 6pvg-A | 2.2 | 7.2 | 75 | 445 | 5 | MOLECULE: FAD MONOOXYGENASE; |
| 1707 | 1w36-C | 2.2 | 22.3 | 142 | 1121 | 8 | MOLECULE: EXODEOXYRIBONUCLEASE V BETA CHAIN; |
| 1708 | 2gsc-B | 2.2 | 7.7 | 58 | 117 | 2 | MOLECULE: PUTATIVE UNCHARACTERIZED PROTEIN XCC0516; |
| 1709 | 7cgp-J | 2.2 | 5 | 57 | 83 | 5 | MOLECULE: MITOCHONDRIAL IMPORT INNER MEMBRANE TRANSLOCASE S |
| 1710 | 6tdv-A | 2.2 | 22.4 | 92 | 486 | 5 | MOLECULE: ATPTB1; |
| 1711 | 3bpx-A | 2.2 | 4.3 | 49 | 147 | 4 | MOLECULE: TRANSCRIPTIONAL REGULATOR; |
| 1712 | 5xyf-A | 2.2 | 5.5 | 45 | 192 | 7 | MOLECULE: TERF1-INTERACTING NUCLEAR FACTOR 2; |

| | | | | | | | |
|---|---|---|---|---|---|---|---|
| 1713 | 7cae-E | 2.2 | 10.9 | 64 | 443 | 6 | MOLECULE: ABC TRANSPORTER, ATP-BINDING PROTEIN SUGC; |
| 1714 | 1zs4-A | 2.2 | 7.2 | 46 | 82 | 4 | MOLECULE: DNA - 27MER; |
| 1715 | 6s3l-K | 2.2 | 6.5 | 49 | 193 | 6 | MOLECULE: FLAGELLAR BIOSYNTHETIC PROTEIN FLIP; |
| 1716 | 6rmo-I | 2.2 | 5 | 75 | 432 | 3 | MOLECULE: IMP-SPECIFIC 5'-NUCLEOTIDASE, PUTATIVE; |
| 1717 | 2oap-1 | 2.2 | 12 | 53 | 498 | 9 | MOLECULE: TYPE II SECRETION SYSTEM PROTEIN; |
| 1718 | 1vp7-B | 2.2 | 7 | 52 | 77 | 17 | MOLECULE: EXODEOXYRIBONUCLEASE VII SMALL SUBUNIT; |
| 1719 | 5u30-A | 2.2 | 8.7 | 72 | 1085 | 10 | MOLECULE: CRISPR-ASSOCIATED ENDONUCLEASE C2C1; |
| 1720 | 6ekr-A | 2.2 | 5 | 76 | 303 | 4 | MOLECULE: TYPE II SITE-SPECIFIC DEOXYRIBONUCLEASE; |
| 1721 | 3b5m-B | 2.2 | 13.3 | 64 | 195 | 9 | MOLECULE: UNCHARACTERIZED PROTEIN; |
| 1722 | 4bjt-A | 2.2 | 4.8 | 63 | 152 | 10 | MOLECULE: DNA-BINDING PROTEIN RAP1; |
| 1723 | 7kuw-A | 2.2 | 7.3 | 56 | 62 | 7 | MOLECULE: SEQUENCE-BASED DESIGNED PROTEIN NMT_0994_GUIDED_0 |
| 1724 | 7b54-X | 2.2 | 4.2 | 87 | 1732 | 5 | MOLECULE: VAR2CSA IN PRESENCE OF PLCS, DBL1-DBL4,ERYTHROCYT |
| 1725 | 2guz-B | 2.2 | 5.5 | 43 | 65 | 9 | MOLECULE: MITOCHONDRIAL IMPORT INNER MEMBRANE TRANSLOCASE S |
| 1726 | 2n5j-A | 2.2 | 4.4 | 43 | 49 | 5 | MOLECULE: RIBONUCLEASE ZC3H12A; |
| 1727 | 3u8p-A | 2.2 | 8.6 | 70 | 336 | 9 | MOLECULE: CYTOCHROME B562 INTEGRAL FUSION WITH ENHANCED GRE |
| 1728 | 2p5t-E | 2.2 | 5.8 | 62 | 95 | 10 | MOLECULE: FRAGMENT OF PEZA HELIX-TURN-HELIX MOTIF; |
| 1729 | 6toa-D | 2.2 | 3.6 | 44 | 195 | 11 | MOLECULE: ADAPTOR PROTEIN RCC01688; |
| 1730 | 5lj3-G | 2.2 | 7 | 50 | 97 | 10 | MOLECULE: U5 SNRNA (SMALL NUCLEAR RNA); |
| 1731 | 7bqr-A | 2.2 | 2.8 | 48 | 137 | 21 | MOLECULE: MUSSOC; |
| 1732 | 4gfq-A | 2.2 | 3.1 | 46 | 186 | 9 | MOLECULE: RIBOSOME-RECYCLING FACTOR; |
| 1733 | 4a9a-A | 2.2 | 13.8 | 62 | 356 | 10 | MOLECULE: RIBOSOME-INTERACTING GTPASE 1; |
| 1734 | 5e5w-B | 2.2 | 8 | 51 | 149 | 18 | MOLECULE: HEMAGGLUTININ-ESTERASE; |
| 1735 | 2kw2-A | 2.2 | 4.6 | 54 | 101 | 7 | MOLECULE: SPECIALIZED ACYL CARRIER PROTEIN; |
| 1736 | 3gac-C | 2.2 | 3.3 | 47 | 117 | 13 | MOLECULE: MACROPHAGE MIGRATION INHIBITORY FACTOR-LIKE |
| 1737 | 4y13-A | 2.2 | 11.6 | 68 | 244 | 4 | MOLECULE: TRANSCRIPTIONAL REGULATOR OF FTSQAZ GENE CLUSTER; |
| 1738 | 6fnp-A | 2.2 | 13.5 | 65 | 158 | 3 | MOLECULE: MEMBRANE PROTEIN; |
| 1739 | 6wg3-E | 2.2 | 14 | 57 | 1226 | 7 | MOLECULE: STRUCTURAL MAINTENANCE OF CHROMOSOMES PROTEIN 1A; |
| 1740 | 6bv7-A | 2.2 | 3.9 | 41 | 57 | 12 | MOLECULE: SODIUM/CALCIUM EXCHANGER 1; |
| 1741 | 7d3u-D | 2.2 | 7.7 | 83 | 534 | 10 | MOLECULE: MONOVALENT NA+/H+ ANTIPORTER SUBUNIT D; |
| 1742 | 6qly-A | 2.2 | 13.7 | 87 | 330 | 8 | MOLECULE: E3 UBIQUITIN-PROTEIN LIGASE MYLIP; |
| 1743 | 6rjb-B | 2.2 | 10.3 | 57 | 622 | 2 | MOLECULE: TRANSKETOLASE; |
| 1744 | 4ckg-A | 2.2 | 12.1 | 85 | 368 | 11 | MOLECULE: ARF-GAP WITH COILED-COIL, ANK REPEAT AND PH DOMAI |
| 1745 | 6fon-A | 2.2 | 2.7 | 40 | 252 | 15 | MOLECULE: COPPER CHAPERONE FOR SUPEROXIDE DISMUTASE; |
| 1746 | 7tbs-A | 2.2 | 14.6 | 64 | 221 | 16 | MOLECULE: GLUTAREDOXIN 2; |
| 1747 | 2jxu-A | 2.2 | 4.1 | 56 | 153 | 7 | MOLECULE: TERB; |

| | | | | | | | |
|---|---|---|---|---|---|---|---|
| 1748 | 4uzz-A | 2.2 | 4 | 52 | 108 | 2 | MOLECULE: INTRAFLAGELLAR TRANSPORT COMPLEX B PROTEIN 46 |
| 1749 | 4fyg-A | 2.2 | 8.9 | 72 | 743 | 7 | MOLECULE: SIDF, INHIBITOR OF GROWTH FAMILY, MEMBER 3; |
| 1750 | 5a5t-L | 2.2 | 10.9 | 90 | 372 | 2 | MOLECULE: EUKARYOTIC TRANSLATION INITIATION FACTOR 3 SUBUNI |
| 1751 | 7apk-i | 2.2 | 10.8 | 59 | 275 | 8 | MOLECULE: THO COMPLEX SUBUNIT 1; |
| 1752 | 3u9j-A | 2.2 | 3.3 | 68 | 157 | 7 | MOLECULE: F-BOX/LRR-REPEAT PROTEIN 5; |
| 1753 | 6dku-A | 2.2 | 2.9 | 43 | 125 | 7 | MOLECULE: VP35; |
| 1754 | 7cm3-A | 2.2 | 8.7 | 100 | 1278 | 3 | MOLECULE: SODIUM LEAK CHANNEL NON-SELECTIVE PROTEIN; |
| 1755 | 6ar7-C | 2.2 | 4.5 | 63 | 207 | 8 | MOLECULE: UNCHARACTERIZED PROTEIN; |
| 1756 | 3mse-B | 2.2 | 5.5 | 62 | 168 | 2 | MOLECULE: CALCIUM-DEPENDENT PROTEIN KINASE, PUTATIVE; |
| 1757 | 6zr2-e | 2.2 | 13.2 | 68 | 105 | 9 | MOLECULE: NADH-UBIQUINONE OXIDOREDUCTASE CHAIN 3; |
| 1758 | 6msr-A | 2.2 | 5.6 | 54 | 76 | 17 | MOLECULE: PRO-2.5; |
| 1759 | 1fe8-C | 2.2 | 11.2 | 59 | 191 | 2 | MOLECULE: VON WILLEBRAND FACTOR; |
| 1760 | 6o84-A | 2.2 | 7.6 | 95 | 415 | 11 | MOLECULE: LOC100127796 PROTEIN,LOC100127796 PROTEIN,OTOP3, |
| 1761 | 5im3-A | 2.2 | 10.1 | 69 | 874 | 7 | MOLECULE: RIBONUCLEOSIDE-DIPHOSPHATE REDUCTASE; |
| 1762 | 2kw7-A | 2.2 | 3 | 52 | 157 | 15 | MOLECULE: CONSERVED DOMAIN PROTEIN; |
| 1763 | 3ub0-D | 2.2 | 6.8 | 63 | 194 | 16 | MOLECULE: NON-STRUCTURAL PROTEIN 6, NSP6,; |
| 1764 | 6fvb-A | 2.2 | 6.7 | 86 | 1001 | 6 | MOLECULE: IMPORTIN BETA-LIKE PROTEIN KAP120; |
| 1765 | 6f2p-A | 2.2 | 15 | 80 | 1040 | 4 | MOLECULE: PAENIBACILLUS XANTHAN LYASE; |
| 1766 | 3mk7-C | 2.2 | 8.1 | 56 | 303 | 9 | MOLECULE: CYTOCHROME C OXIDASE, CBB3-TYPE, SUBUNIT N; |
| 1767 | 4iw7-A | 2.2 | 10.1 | 68 | 335 | 9 | MOLECULE: 8-AMINO-7-OXONONANOATE SYNTHASE; |
| 1768 | 4lmg-B | 2.2 | 3.9 | 50 | 121 | 10 | MOLECULE: IRON-REGULATED TRANSCRIPTIONAL ACTIVATOR AFT2; |
| 1769 | 3l09-C | 2.2 | 4.5 | 81 | 253 | 5 | MOLECULE: PUTATIVE TRANSCRIPTIONAL REGULATOR; |
| 1770 | 2vz9-B | 2.2 | 4.3 | 82 | 2086 | 5 | MOLECULE: FATTY ACID SYNTHASE; |
| 1771 | 6nbn-A | 2.2 | 4 | 65 | 123 | 5 | MOLECULE: AAEL005772-PA; |
| 1772 | 6kth-A | 2.2 | 8 | 57 | 190 | 7 | MOLECULE: JUVENILE HORMONE DIOL KINASE; |
| 1773 | 7nnl-B | 2.2 | 9.1 | 74 | 682 | 5 | MOLECULE: POTASSIUM-TRANSPORTING ATPASE POTASSIUM-BINDING S |
| 1774 | 5d06-A | 2.2 | 8 | 114 | 1526 | 15 | MOLECULE: UNCHARACTERIZED PROTEIN; |
| 1775 | 3k1r-A | 2.2 | 5.6 | 59 | 192 | 5 | MOLECULE: HARMONIN; |
| 1776 | 2a7o-A | 2.2 | 3.8 | 51 | 100 | 8 | MOLECULE: HUNTINGTIN INTERACTING PROTEIN B; |
| 1777 | 6xz4-A | 2.2 | 16.4 | 78 | 304 | 8 | MOLECULE: TALIN ROD DOMAIN-CONTAINING PROTEIN 1; |
| 1778 | 7a23-X | 2.2 | 10.9 | 47 | 103 | 4 | MOLECULE: 51KDA; |
| 1779 | 4v1f-A | 2.2 | 4.7 | 45 | 86 | 2 | MOLECULE: F0F1 ATP SYNTHASE SUBUNIT C; |
| 1780 | 5lm2-B | 2.2 | 7.7 | 71 | 340 | 14 | MOLECULE: TYROSINE-PROTEIN PHOSPHATASE NON-RECEPTOR TYPE 23 |
| 1781 | 2o3f-A | 2.2 | 3.3 | 50 | 83 | 14 | MOLECULE: PUTATIVE HTH-TYPE TRANSCRIPTIONAL REGULATOR YBBH; |
| 1782 | 6c4v-A | 2.2 | 2.6 | 45 | 81 | 9 | MOLECULE: POLYKETIDE SYNTHASE PKS13; |

| | | | | | | | |
|---|---|---|---|---|---|---|---|
| 1783 | 3pas-B | 2.2 | 10.8 | 59 | 192 | 8 | MOLECULE: TETR FAMILY TRANSCRIPTION REGULATOR; |
| 1784 | 7lga-D | 2.2 | 3.7 | 49 | 91 | 10 | MOLECULE: RETROTRANSPOSON-DERIVED PROTEIN PEG10; |
| 1785 | 4nqi-D | 2.2 | 6.3 | 71 | 232 | 11 | MOLECULE: SH3 DOMAIN-CONTAINING PROTEIN; |
| 1786 | 3rzs-A | 2.2 | 3.5 | 59 | 119 | 7 | MOLECULE: OBP14; |
| 1787 | 2gd5-D | 2.2 | 12 | 61 | 162 | 8 | MOLECULE: CHARGED MULTIVESICULAR BODY PROTEIN 3; |
| 1788 | 6u4k-A | 2.2 | 7.2 | 58 | 391 | 12 | MOLECULE: TALIN-2; |
| 1789 | 6v9z-A | 2.2 | 10.9 | 54 | 715 | 4 | MOLECULE: ABC-TYPE BACTERIOCIN TRANSPORTER; |
| 1790 | 3utk-A | 2.2 | 3.1 | 44 | 95 | 7 | MOLECULE: LIPOPROTEIN OUTS; |
| 1791 | 2hf6-A | 2.2 | 9.6 | 54 | 149 | 11 | MOLECULE: COATOMER SUBUNIT ZETA-1; |
| 1792 | 3mn2-A | 2.2 | 5.1 | 63 | 108 | 11 | MOLECULE: PROBABLE ARAC FAMILY TRANSCRIPTIONAL REGULATOR; |
| 1793 | 6iie-A | 2.2 | 5.5 | 50 | 87 | 12 | MOLECULE: DIACYLGLYCEROL KINASE ALPHA; |
| 1794 | 4c5f-A | 2.2 | 3.3 | 61 | 330 | 5 | MOLECULE: MEMBRANE-BOUND LYTIC MUREIN TRANSGLYCOSYLASE C; |
| 1795 | 6v4l-A | 2.2 | 11.9 | 70 | 479 | 7 | MOLECULE: TRK SYSTEM POTASSIUM UPTAKE PROTEIN TRKH; |
| 1796 | 2dzn-F | 2.2 | 3 | 44 | 69 | 9 | MOLECULE: PROBABLE 26S PROTEASOME REGULATORY SUBUNIT P28; |
| 1797 | 5u70-A | 2.2 | 9.5 | 81 | 901 | 10 | MOLECULE: POTASSIUM CHANNEL SUBFAMILY T MEMBER 1; |
| 1798 | 2bw3-A | 2.2 | 14 | 63 | 518 | 16 | MOLECULE: TRANSPOSASE; |
| 1799 | 3aei-A | 2.2 | 4.5 | 40 | 94 | 10 | MOLECULE: PREFOLDIN BETA SUBUNIT 2; |
| 1800 | 1hyn-Q | 2.2 | 12 | 59 | 302 | 2 | MOLECULE: BAND 3 ANION TRANSPORT PROTEIN; |
| 1801 | 3zc7-B | 2.2 | 7 | 40 | 63 | 5 | MOLECULE: ADENOSINE MONOPHOSPHATE-PROTEIN TRANSFERASE VBHT; |
| 1802 | 3h4z-B | 2.2 | 10 | 71 | 563 | 8 | MOLECULE: MALTOSE-BINDING PERIPLASMIC PROTEIN FUSED WITH AL |
| 1803 | 2mbg-A | 2.2 | 4.4 | 60 | 265 | 5 | MOLECULE: RALA-BINDING PROTEIN 1; |
| 1804 | 6ww7-C | 2.2 | 8.9 | 67 | 209 | 6 | MOLECULE: ER MEMBRANE PROTEIN COMPLEX SUBUNIT 1; |
| 1805 | 1og6-C | 2.2 | 12.8 | 56 | 298 | 9 | MOLECULE: HYPOTHETICAL OXIDOREDUCTASE YDHF; |
| 1806 | 1zae-A | 2.2 | 8.1 | 52 | 70 | 13 | MOLECULE: EARLY PROTEIN GP16.7; |
| 1807 | 2p0n-A | 2.2 | 2.7 | 52 | 161 | 6 | MOLECULE: HYPOTHETICAL PROTEIN NMB1532; |
| 1808 | 6wb9-6 | 2.2 | 4 | 47 | 98 | 11 | MOLECULE: ENDOPLASMIC RETICULUM MEMBRANE PROTEIN COMPLEX SU |
| 1809 | 3vw4-A | 2.2 | 2.2 | 44 | 122 | 9 | MOLECULE: REP; |
| 1810 | 5k8c-A | 2.2 | 5.4 | 76 | 358 | 4 | MOLECULE: 3-DEOXY-ALPHA-D-MANNO-OCTULOSONATE 8-OXIDASE; |
| 1811 | 5a34-C | 2.2 | 5.9 | 74 | 170 | 7 | MOLECULE: BIFUNCTIONAL GLUTAMATE/PROLINE--TRNA LIGASE; |
| 1812 | 5h0j-A | 2.2 | 10.4 | 61 | 234 | 7 | MOLECULE: UNCHARACTERIZED PROTEIN; |
| 1813 | 7n8n-B | 2.2 | 3.3 | 55 | 202 | 7 | MOLECULE: HISTONE H4-H3 DOUBLET; |
| 1814 | 3tdw-A | 2.2 | 8.8 | 76 | 302 | 4 | MOLECULE: GENTAMICIN RESISTANCE PROTEIN; |
| 1815 | 2ymb-D | 2.2 | 13.6 | 65 | 231 | 12 | MOLECULE: MIT DOMAIN-CONTAINING PROTEIN 1; |
| 1816 | 7my4-A | 2.2 | 9.4 | 48 | 66 | 15 | MOLECULE: SPERM AUTOANTIGENIC PROTEIN 17; |
| 1817 | 3ed5-A | 2.2 | 21 | 65 | 232 | 11 | MOLECULE: YFNB; |

| | | | | | | | |
|---|---|---|---|---|---|---|---|
| 1818 | 1wud-A | 2.2 | 3 | 41 | 77 | 12 | MOLECULE: ATP-DEPENDENT DNA HELICASE RECQ; |
| 1819 | 6iu4-A | 2.2 | 9.7 | 68 | 225 | 10 | MOLECULE: VIT1; |
| 1820 | 3o3u-N | 2.2 | 11.2 | 76 | 580 | 4 | MOLECULE: MALTOSE-BINDING PERIPLASMIC PROTEIN, ADVANCED GLY |
| 1821 | 2omo-A | 2.2 | 11.3 | 52 | 105 | 6 | MOLECULE: DUF176; |
| 1822 | 4xa6-A | 2.2 | 7.1 | 55 | 168 | 2 | MOLECULE: GP7-MYH7(1777-1855)-EB1 CHIMERA PROTEIN; |
| 1823 | 5kkp-A | 2.2 | 16.9 | 67 | 509 | 1 | MOLECULE: PSEUDOURIDYLATE SYNTHASE 7; |
| 1824 | 1v32-A | 2.2 | 8.4 | 55 | 101 | 9 | MOLECULE: HYPOTHETICAL PROTEIN RAFL09-47-K03; |
| 1825 | 2ntz-A | 2.2 | 11.6 | 87 | 183 | 11 | MOLECULE: 5'- |
| 1826 | 6mrs-A | 2.2 | 2.6 | 38 | 77 | 18 | MOLECULE: PEAK6; |
| 1827 | 3c7n-A | 2.2 | 7.9 | 76 | 648 | 3 | MOLECULE: HEAT SHOCK PROTEIN HOMOLOG SSE1; |
| 1828 | 6jmg-A | 2.2 | 9.2 | 65 | 267 | 14 | MOLECULE: DNAJ HOMOLOG SUBFAMILY C MEMBER 27-A; |
| 1829 | 2ptf-A | 2.2 | 11.7 | 52 | 200 | 12 | MOLECULE: UNCHARACTERIZED PROTEIN MTH_863; |
| 1830 | 6wqz-A | 2.2 | 4.5 | 57 | 536 | 9 | MOLECULE: AUTOPHAGY-RELATED PROTEIN 9A; |
| 1831 | 3ziu-A | 2.2 | 8.4 | 74 | 621 | 12 | MOLECULE: LEUCYL-TRNA SYNTHETASE; |
| 1832 | 3twb-C | 2.2 | 4.4 | 67 | 417 | 4 | MOLECULE: PUTATIVE DEHYDRATASE; |
| 1833 | 5h3i-D | 2.2 | 4.9 | 53 | 94 | 9 | MOLECULE: PUTATIVE ACYL-COA-BINDING PROTEIN; |
| 1834 | 3fvq-B | 2.2 | 13.2 | 80 | 349 | 3 | MOLECULE: FE(3+) IONS IMPORT ATP-BINDING PROTEIN FBPC; |
| 1835 | 4eyy-Q | 2.2 | 13.3 | 63 | 188 | 8 | MOLECULE: ICMR; |
| 1836 | 5mc1-B | 2.2 | 11.6 | 87 | 487 | 7 | MOLECULE: XAA-PRO DIPEPTIDASE; |
| 1837 | 3iwf-B | 2.2 | 4.8 | 56 | 90 | 4 | MOLECULE: TRANSCRIPTION REGULATOR RPIR FAMILY; |
| 1838 | 6psd-A | 2.2 | 2.3 | 37 | 73 | 0 | MOLECULE: EF-HAND CALCIUM-BINDING DOMAIN-CONTAINING PROTEIN |
| 1839 | 3uw3-B | 2.2 | 7.6 | 68 | 375 | 9 | MOLECULE: ASPARTATE-SEMIALDEHYDE DEHYDROGENASE; |
| 1840 | 2dae-A | 2.2 | 3 | 44 | 75 | 14 | MOLECULE: KIAA0733 PROTEIN; |
| 1841 | 2cru-A | 2.2 | 10.6 | 60 | 118 | 10 | MOLECULE: PROGRAMMED CELL DEATH PROTEIN 5; |
| 1842 | 6ek4-A | 2.2 | 12.7 | 69 | 342 | 7 | MOLECULE: PAXB; |
| 1843 | 7ad3-A | 2.2 | 5.2 | 98 | 298 | 7 | MOLECULE: PHEROMONE ALPHA FACTOR RECEPTOR; |
| 1844 | 2vsg-A | 2.2 | 5.7 | 67 | 358 | 4 | MOLECULE: VARIANT SURFACE GLYCOPROTEIN ILTAT 1.24; |
| 1845 | 2fbi-A | 2.2 | 6 | 53 | 136 | 11 | MOLECULE: PROBABLE TRANSCRIPTIONAL REGULATOR; |
| 1846 | 5uyo-A | 2.2 | 4.7 | 34 | 44 | 9 | MOLECULE: HEEH_RD4_0097; |
| 1847 | 2y9z-B | 2.2 | 14.4 | 82 | 595 | 6 | MOLECULE: IMITATION SWITCH PROTEIN 1 (DEL_ATPASE); |
| 1848 | 2nb2-A | 2.2 | 2.5 | 37 | 38 | 11 | MOLECULE: NIGELLIN-1.1; |
| 1849 | 3ce8-A | 2.2 | 4.7 | 46 | 90 | 7 | MOLECULE: PUTATIVE PII-LIKE NITROGEN REGULATORY PROTEIN; |
| 1850 | 6zka-K | 2.2 | 5.6 | 50 | 98 | 4 | MOLECULE: NADH-UBIQUINONE OXIDOREDUCTASE CHAIN 3; |
| 1851 | 7f9m-A | 2.2 | 14.9 | 71 | 158 | 4 | MOLECULE: RIFIN; |
| 1852 | 5bs1-D | 2.2 | 6.5 | 69 | 121 | 6 | MOLECULE: CRRBCX-IIA; |

| | | | | | | | |
|---|---|---|---|---|---|---|---|
| 1853 | 3c4r-C | 2.2 | 4.3 | 84 | 127 | 10 | MOLECULE: UNCHARACTERIZED PROTEIN; |
| 1854 | 6rd4-2 | 2.2 | 13.3 | 90 | 441 | 4 | MOLECULE: ASA-10: POLYTOMELLA F-ATP SYNTHASE ASSOCIATED SUB |
| 1855 | 4pl0-B | 2.2 | 11.9 | 77 | 576 | 3 | MOLECULE: MICROCIN-J25 EXPORT ATP-BINDING/PERMEASE PROTEIN |
| 1856 | 1i7q-A | 2.2 | 4.2 | 64 | 517 | 13 | MOLECULE: ANTHRANILATE SYNTHASE; |
| 1857 | 5fd4-A | 2.2 | 13.7 | 89 | 309 | 8 | MOLECULE: COMR; |
| 1858 | 2gta-A | 2.2 | 5.7 | 56 | 98 | 11 | MOLECULE: HYPOTHETICAL PROTEIN YPJD; |
| 1859 | 3qth-B | 2.2 | 4 | 75 | 164 | 8 | MOLECULE: UNCHARACTERIZED PROTEIN; |
| 1860 | 2p62-A | 2.2 | 10.1 | 69 | 241 | 7 | MOLECULE: HYPOTHETICAL PROTEIN PH0156; |
| 1861 | 2iho-A | 2.2 | 5 | 56 | 292 | 4 | MOLECULE: LECTIN; |
| 1862 | 2w96-A | 2.2 | 10.9 | 76 | 249 | 7 | MOLECULE: CELL DIVISION PROTEIN KINASE 4; |
| 1863 | 6lcu-A | 2.2 | 14 | 79 | 756 | 15 | MOLECULE: MTSASE; |
| 1864 | 5adz-A | 2.2 | 10.6 | 67 | 566 | 4 | MOLECULE: ALKYLDIHYDROXYACETONEPHOSPHATE SYNTHASE, |
| 1865 | 6ygh-D | 2.2 | 3 | 46 | 193 | 7 | MOLECULE: CAPSID PROTEIN; |
| 1866 | 6vmb-g | 2.2 | 5.2 | 49 | 323 | 4 | MOLECULE: ATP SYNTHASE SUBUNIT ALPHA, CHLOROPLASTIC; |
| 1867 | 1rj1-A | 2.2 | 5.9 | 66 | 148 | 2 | MOLECULE: INVERTASE INHIBITOR; |
| 1868 | 3hc1-A | 2.2 | 13.1 | 78 | 298 | 9 | MOLECULE: UNCHARACTERIZED HDOD DOMAIN PROTEIN; |
| 1869 | 2owy-A | 2.2 | 7.5 | 60 | 306 | 12 | MOLECULE: RECOMBINATION-ASSOCIATED PROTEIN RDGC; |
| 1870 | 7ef9-A | 2.2 | 12.5 | 64 | 414 | 9 | MOLECULE: ADENINE DNA GLYCOSYLASE; |
| 1871 | 7m4m-A | 2.2 | 4.4 | 47 | 173 | 13 | MOLECULE: E3 UBIQUITIN-PROTEIN LIGASE RNF216; |
| 1872 | 6dlm-A | 2.2 | 4 | 38 | 74 | 8 | MOLECULE: DHD127_A; |
| 1873 | 5zhy-B | 2.2 | 4.6 | 40 | 89 | 8 | MOLECULE: SPIKE GLYCOPROTEIN, SPIKE GLYCOPROTEIN; |
| 1874 | 7etw-A | 2.2 | 5.2 | 92 | 197 | 4 | MOLECULE: INSULIN-INDUCED GENE 2 PROTEIN; |
| 1875 | 6tky-A | 2.2 | 10.5 | 65 | 417 | 9 | MOLECULE: DEDICATOR OF CYTOKINESIS PROTEIN 10; |
| 1876 | 6w1s-J | 2.2 | 2.9 | 53 | 167 | 8 | MOLECULE: MEDIATOR OF RNA POLYMERASE II TRANSCRIPTION SUBUN |
| 1877 | 3vej-B | 2.2 | 3 | 37 | 41 | 8 | MOLECULE: UBIQUITIN-LIKE PROTEIN MDY2; |
| 1878 | 5eik-A | 2.2 | 3.8 | 68 | 234 | 10 | MOLECULE: UNCHARACTERIZED PROTEIN Y57A10A.28; |
| 1879 | 7auf-A | 2.2 | 2.7 | 47 | 89 | 15 | MOLECULE: SIMILAR TO ACYL CARRIER PROTEIN; |
| 1880 | 2x3m-A | 2.2 | 4.7 | 58 | 166 | 5 | MOLECULE: HYPOTHETICAL PROTEIN ORF239; |
| 1881 | 2i15-A | 2.2 | 5.5 | 40 | 122 | 5 | MOLECULE: HYPOTHETICAL PROTEIN MG296 HOMOLOG; |
| 1882 | 4art-A | 2.2 | 14.8 | 75 | 237 | 4 | MOLECULE: STRUCTURAL PROTEIN ORF273; |
| 1883 | 7lma-A | 2.2 | 4.9 | 74 | 1012 | 3 | MOLECULE: TELOMERASE LA-RELATED PROTEIN P65; |
| 1884 | 4dvy-P | 2.2 | 7.3 | 57 | 656 | 9 | MOLECULE: CYTOTOXICITY-ASSOCIATED IMMUNODOMINANT ANTIGEN; |
| 1885 | 6hyp-A | 2.2 | 12.1 | 80 | 2272 | 6 | MOLECULE: MIDASIN,MIDASIN; |
| 1886 | 2bf0-X | 2.2 | 3.8 | 52 | 131 | 4 | MOLECULE: PCF11; |
| 1887 | 6fsf-A | 2.2 | 3.9 | 68 | 218 | 15 | MOLECULE: GTPASE-ACTIVATING PROTEIN BEM3; |

| | | | | | | | |
|---|---|---|---|---|---|---|---|
| 1888 | 6p4o-F | 2.2 | 18.3 | 65 | 481 | 11 | MOLECULE: DNA-DEPENDENT ATPASE XPBII; |
| 1889 | 2fuq-A | 2.2 | 14.6 | 83 | 747 | 6 | MOLECULE: HEPARINASE II PROTEIN; |
| 1890 | 6khj-E | 2.2 | 4.5 | 54 | 101 | 13 | MOLECULE: NAD(P)H-QUINONE OXIDOREDUCTASE SUBUNIT 1; |
| 1891 | 5cuy-D | 2.2 | 13.4 | 81 | 396 | 9 | MOLECULE: ACIDOCALCISOMAL PYROPHOSPHATASE; |
| 1892 | 6f0k-H | 2.2 | 4.5 | 75 | 156 | 8 | MOLECULE: CYTOCHROME C FAMILY PROTEIN; |
| 1893 | 6qeq-A | 2.2 | 3.3 | 39 | 111 | 5 | MOLECULE: PCFF; |
| 1894 | 3dsq-B | 2.2 | 3.1 | 47 | 282 | 6 | MOLECULE: PYRROLYSYL-TRNA SYNTHETASE; |
| 1895 | 6e4j-A | 2.2 | 3.9 | 54 | 72 | 13 | MOLECULE: UNCHARACTERIZED PROTEIN PF2048.1; |
| 1896 | 5ted-B | 2.2 | 3.3 | 41 | 210 | 7 | MOLECULE: LMO0488 PROTEIN; |
| 1897 | 2vrz-A | 2.2 | 7.7 | 52 | 98 | 10 | MOLECULE: VIRULENCE FACTOR ESXA; |
| 1898 | 6zh3-A | 2.2 | 8.6 | 57 | 170 | 5 | MOLECULE: VACUOLAR PROTEIN-SORTING-ASSOCIATED PROTEIN 24; |
| 1899 | 6ec8-A | 2.2 | 7 | 65 | 803 | 5 | MOLECULE: LANTIBIOTIC DEHYDRATASE DOMAIN PROTEIN; |
| 1900 | 4lur-A | 2.2 | 8.2 | 69 | 309 | 7 | MOLECULE: INTERPHOTORECEPTOR RETINOID-BINDING PROTEIN(IRBP) |
| 1901 | 7eev-B | 2.2 | 17.6 | 61 | 399 | 7 | MOLECULE: GTP CYCLOHYDROLASE II; |
| 1902 | 5uay-A | 2.2 | 7.1 | 73 | 302 | 4 | MOLECULE: PROTEIN TOC75-3, CHLOROPLASTIC; |
| 1903 | 6rwy-g | 2.2 | 4.3 | 42 | 86 | 7 | MOLECULE: INNER ROD PROTEIN; |
| 1904 | 4oh3-B | 2.2 | 17.7 | 98 | 537 | 4 | MOLECULE: NITRATE TRANSPORTER 1.1; |
| 1905 | 3bhp-C | 2.2 | 5.6 | 43 | 54 | 5 | MOLECULE: UPF0291 PROTEIN YNZC; |
| 1906 | 5lsk-N | 2.2 | 8.7 | 62 | 173 | 6 | MOLECULE: PROTEIN MIS12 HOMOLOG; |
| 1907 | 2oev-A | 2.2 | 5.8 | 64 | 697 | 6 | MOLECULE: PROGRAMMED CELL DEATH 6-INTERACTING PROTEIN; |
| 1908 | 1u2m-C | 2.2 | 9.4 | 60 | 143 | 2 | MOLECULE: HISTONE-LIKE PROTEIN HLP-1; |
| 1909 | 5fjd-B | 2.2 | 4.2 | 49 | 111 | 6 | MOLECULE: COPPER STORAGE PROTEIN 1; |
| 1910 | 5gpa-A | 2.2 | 7 | 42 | 189 | 2 | MOLECULE: TRANSCRIPTIONAL REGULATOR (TETR/ACRR FAMILY); |
| 1911 | 7m7a-A | 2.2 | 8.1 | 74 | 538 | 11 | MOLECULE: PHOSPHOINOSITIDE 3-KINASE MAVQ; |
| 1912 | 3lk7-A | 2.2 | 14.3 | 79 | 448 | 11 | MOLECULE: UDP-N-ACETYLMURAMOYLALANINE--D-GLUTAMATE LIGASE; |
| 1913 | 4fhn-B | 2.2 | 8.9 | 62 | 1022 | 8 | MOLECULE: NUCLEOPORIN NUP37; |
| 1914 | 5uaw-B | 2.2 | 15.4 | 59 | 281 | 3 | MOLECULE: PYRROLINE-5-CARBOXYLATE REDUCTASE 1, MITOCHONDRIA |
| 1915 | 3u8v-A | 2.2 | 4.4 | 52 | 83 | 13 | MOLECULE: METAL-BINDING PROTEIN SMBP; |
| 1916 | 1z9h-A | 2.2 | 10.1 | 72 | 274 | 4 | MOLECULE: MEMBRANE-ASSOCIATED PROSTAGLANDIN E SYNTHASE-2; |
| 1917 | 5mmj-n | 2.2 | 8.9 | 53 | 99 | 6 | MOLECULE: 50S RIBOSOMAL PROTEIN L31; |
| 1918 | 5wu1-A | 2.2 | 4.2 | 67 | 493 | 7 | MOLECULE: SPECKLE TARGETED PIP5K1A-REGULATED POLY(A) POLYME |
| 1919 | 4ps2-A | 2.2 | 4.2 | 46 | 79 | 9 | MOLECULE: PUTATIVE TYPE VI SECRETION PROTEIN; |
| 1920 | 1z0p-A | 2.2 | 5.5 | 47 | 73 | 6 | MOLECULE: HYPOTHETICAL PROTEIN SPY1572; |
| 1921 | 3zym-C | 2.2 | 11 | 65 | 280 | 3 | MOLECULE: PHOSPHATIDYLINOSITOL-BINDING CLATHRIN ASSEMBLY PR |
| 1922 | 5vtl-A | 2.2 | 7.5 | 65 | 210 | 14 | MOLECULE: TB427.07.360- PUTATIVE UNCHARACTERIZED METACYCLIC |

| | | | | | | | |
|---|---|---|---|---|---|---|---|
| 1923 | 6h02-A | 2.2 | 20.6 | 85 | 1334 | 5 | MOLECULE: MEDIATOR OF RNA POLYMERASE II TRANSCRIPTION SUBUN |
| 1924 | 5ca9-A | 2.2 | 12.8 | 79 | 656 | 8 | MOLECULE: PROTEIN SEY1; |
| 1925 | 5xsi-A | 2.2 | 5.1 | 49 | 333 | 4 | MOLECULE: PHOSPHODIESTERASE ACTING ON CYCLIC DINUCLEOTIDES; |
| 1926 | 5n5e-A | 2.2 | 4.1 | 55 | 98 | 9 | MOLECULE: PFC_05175; |
| 1927 | 4ryk-A | 2.2 | 9.7 | 68 | 295 | 9 | MOLECULE: LMO0325 PROTEIN; |
| 1928 | 5jqz-A | 2.2 | 4.8 | 42 | 75 | 7 | MOLECULE: DE NOVO DESIGNED HOMOTETRAMER; |
| 1929 | 2kob-A | 2.2 | 8 | 43 | 108 | 9 | MOLECULE: UNCHARACTERIZED PROTEIN; |
| 1930 | 6ioh-A | 2.2 | 5.5 | 77 | 375 | 6 | MOLECULE: HOMOSERINE O-ACETYLTRANSFERASE; |
| 1931 | 5cet-A | 2.2 | 11.2 | 52 | 328 | 10 | MOLECULE: BIFUNCTIONAL OLIGORIBONUCLEASE AND PAP PHOSPHATAS |
| 1932 | 4dnh-A | 2.2 | 3.2 | 42 | 386 | 7 | MOLECULE: UNCHARACTERIZED PROTEIN; |
| 1933 | 1w0b-A | 2.2 | 6.5 | 57 | 102 | 9 | MOLECULE: ALPHA-HEMOGLOBIN STABILIZING PROTEIN; |
| 1934 | 5ydn-A | 2.2 | 3.9 | 45 | 109 | 13 | MOLECULE: GENE PRODUCT J; |
| 1935 | 2cs1-A | 2.2 | 5.4 | 43 | 92 | 7 | MOLECULE: PMS1 PROTEIN HOMOLOG 1; |
| 1936 | 5mlq-A | 2.2 | 6.5 | 86 | 226 | 6 | MOLECULE: CDPS; |
| 1937 | 6g0z-A | 2.2 | 7.1 | 71 | 290 | 10 | MOLECULE: RIBOSOME BIOGENESIS GTPASE A; |
| 1938 | 3f0c-A | 2.2 | 11.9 | 71 | 193 | 8 | MOLECULE: TRANSCRIPTIONAL REGULATOR; |
| 1939 | 3k2j-A | 2.2 | 3.2 | 60 | 121 | 5 | MOLECULE: PROTEIN POLYBROMO-1; |
| 1940 | 6s8g-G | 2.2 | 3.8 | 50 | 244 | 2 | MOLECULE: LIPOPOLYSACCHARIDE ABC TRANSPORTER, ATP-BINDING P |
| 1941 | 5lnk-Y | 2.2 | 15.2 | 64 | 171 | 9 | MOLECULE: MITOCHONDRIAL COMPLEX I, 51 KDA SUBUNIT; |
| 1942 | 2dsy-D | 2.2 | 4.4 | 46 | 81 | 9 | MOLECULE: HYPOTHETICAL PROTEIN TTHA0281; |
| 1943 | 6drh-E | 2.2 | 4.5 | 81 | 366 | 11 | MOLECULE: ADP-RIBOSYL-(DINITROGEN REDUCTASE) HYDROLASE; |
| 1944 | 2eyq-A | 2.2 | 10.3 | 72 | 1146 | 8 | MOLECULE: TRANSCRIPTION-REPAIR COUPLING FACTOR; |
| 1945 | 7pt0-A | 2.2 | 12 | 69 | 201 | 12 | MOLECULE: TETR FAMILY TRANSCRIPTIONAL REGULATOR; |
| 1946 | 4x5m-B | 2.2 | 2.6 | 46 | 93 | 11 | MOLECULE: UNCHARACTERIZED PROTEIN; |
| 1947 | 5eyb-A | 2.2 | 4.2 | 59 | 340 | 7 | MOLECULE: DNA-BINDING PROTEIN REB1; |
| 1948 | 2n2h-B | 2.2 | 5.8 | 47 | 125 | 6 | MOLECULE: SIN3 HISTONE DEACETYLASE COREPRESSOR COMPLEX COMP |
| 1949 | 2b8i-A | 2.2 | 3.2 | 50 | 77 | 4 | MOLECULE: PAS FACTOR; |
| 1950 | 6iv9-A | 2.2 | 6.4 | 77 | 874 | 4 | MOLECULE: CAS13D; |
| 1951 | 7mdy-B | 2.2 | 6.3 | 78 | 411 | 3 | MOLECULE: LIPOPROTEIN TRANSPORTER SUBUNIT LOLE; |
| 1952 | 7bin-g | 2.2 | 5.2 | 52 | 260 | 2 | MOLECULE: FLAGELLAR BIOSYNTHETIC PROTEIN FLIP; |
| 1953 | 6nr8-2 | 2.2 | 6.3 | 52 | 103 | 4 | MOLECULE: PREFOLDIN SUBUNIT 1; |
| 1954 | 7jsr-A | 2.2 | 10.2 | 56 | 1496 | 4 | MOLECULE: NAD-SPECIFIC GLUTAMATE DEHYDROGENASE; |
| 1955 | 7pw5-B | 2.2 | 11.2 | 84 | 702 | 6 | MOLECULE: SMG1,SERINE/THREONINE-PROTEIN KINASE SMG1,SMG1, |
| 1956 | 4a12-B | 2.2 | 4.1 | 63 | 186 | 8 | MOLECULE: TRANSCRIPTION FACTOR FAPR; |
| 1957 | 1k7y-A | 2.2 | 15.2 | 64 | 577 | 6 | MOLECULE: METHIONINE SYNTHASE; |

| | | | | | | | |
|---|---|---|---|---|---|---|---|
| 1958 | 5y77-A | 2.2 | 12.9 | 55 | 456 | 9 | MOLECULE: KYNURENINE 3-MONOOXYGENASE; |
| 1959 | 5e1l-A | 2.2 | 8 | 56 | 173 | 9 | MOLECULE: CELL DIVISION PROTEIN ZAPC; |
| 1960 | 2v57-B | 2.2 | 4.3 | 51 | 176 | 6 | MOLECULE: TETR FAMILY TRANSCRIPTIONAL REPRESSOR LFRR; |
| 1961 | 6vr7-A | 2.2 | 5.7 | 69 | 485 | 9 | MOLECULE: MUR LIGASE MIDDLE DOMAIN PROTEIN; |
| 1962 | 3mzy-A | 2.2 | 3.5 | 47 | 123 | 15 | MOLECULE: RNA POLYMERASE SIGMA-H FACTOR; |
| 1963 | 6qct-A | 2.2 | 4.7 | 70 | 722 | 6 | MOLECULE: POLYMERASE ACIDIC PROTEIN; |
| 1964 | 2yqy-A | 2.2 | 3.4 | 53 | 126 | 6 | MOLECULE: HYPOTHETICAL PROTEIN TTHA0303; |
| 1965 | 3okg-B | 2.2 | 4.3 | 50 | 391 | 14 | MOLECULE: RESTRICTION ENDONUCLEASE S SUBUNITS; |
| 1966 | 1vyi-A | 2.2 | 3.2 | 66 | 111 | 6 | MOLECULE: RNA POLYMERASE ALPHA SUBUNIT; |
| 1967 | 3vou-A | 2.2 | 4 | 58 | 139 | 3 | MOLECULE: ION TRANSPORT 2 DOMAIN PROTEIN, VOLTAGE-GATED SOD |
| 1968 | 3eo8-A | 2.2 | 10.3 | 51 | 219 | 6 | MOLECULE: BLUB-LIKE FLAVOPROTEIN; |
| 1969 | 6aun-A | 2.2 | 8.1 | 88 | 618 | 5 | MOLECULE: PLA2G6, IPLA2BETA; |
| 1970 | 3lpz-A | 2.2 | 11.4 | 59 | 314 | 7 | MOLECULE: UNCHARACTERIZED PROTEIN; |
| 1971 | 6nzk-A | 2.2 | 6.8 | 86 | 1175 | 8 | MOLECULE: SPIKE SURFACE GLYCOPROTEIN; |
| 1972 | 5c0w-K | 2.2 | 15.8 | 93 | 576 | 8 | MOLECULE: EXOSOME COMPLEX COMPONENT RRP45; |
| 1973 | 5a1v-K | 2.2 | 9.6 | 67 | 1125 | 6 | MOLECULE: ADP-RIBOSYLATION FACTOR 1; |
| 1974 | 5o9z-L | 2.2 | 9.6 | 57 | 459 | 12 | MOLECULE: PRE-MRNA-PROCESSING-SPLICING FACTOR 8; |
| 1975 | 4xh3-A | 2.1 | 11.1 | 59 | 214 | 2 | MOLECULE: ACTIN-BINDING PROTEIN ANILLIN; |
| 1976 | 3rh2-A | 2.1 | 10.6 | 74 | 210 | 9 | MOLECULE: HYPOTHETICAL TETR-LIKE TRANSCRIPTIONAL REGULATOR; |
| 1977 | 3e8p-A | 2.1 | 4.3 | 57 | 153 | 4 | MOLECULE: UNCHARACTERIZED PROTEIN; |
| 1978 | 1f51-B | 2.1 | 7.3 | 63 | 182 | 10 | MOLECULE: SPORULATION INITIATION PHOSPHOTRANSFERASE B; |
| 1979 | 2xex-B | 2.1 | 16.2 | 93 | 674 | 13 | MOLECULE: ELONGATION FACTOR G; |
| 1980 | 4lws-B | 2.1 | 5.4 | 51 | 88 | 8 | MOLECULE: UNCHARACTERIZED PROTEIN; |
| 1981 | 6u8y-c | 2.1 | 6.2 | 52 | 112 | 4 | MOLECULE: MONOVALENT CATION/H+ ANTIPORTER SUBUNIT E; |
| 1982 | 3a8r-B | 2.1 | 10.9 | 80 | 167 | 10 | MOLECULE: PUTATIVE UNCHARACTERIZED PROTEIN; |
| 1983 | 4ayb-K | 2.1 | 4.1 | 58 | 84 | 7 | MOLECULE: DNA-DIRECTED RNA POLYMERASE; |
| 1984 | 5of3-C | 2.1 | 9.5 | 46 | 107 | 7 | MOLECULE: DNA PRIMASE SMALL SUBUNIT PRIS; |
| 1985 | 7dqk-B | 2.1 | 3.7 | 56 | 479 | 7 | MOLECULE: PROTEIN DETOXIFICATION; |
| 1986 | 4gu7-A | 2.1 | 5.2 | 69 | 310 | 12 | MOLECULE: PUTATIVE UNCHARACTERIZED PROTEIN SCO7193; |
| 1987 | 6c2s-A | 2.1 | 5.6 | 83 | 142 | 14 | MOLECULE: TRANSCRIPTIONAL REGULATOR, MARR FAMILY; |
| 1988 | 3k1l-A | 2.1 | 13 | 62 | 376 | 8 | MOLECULE: FANCL; |
| 1989 | 4r6i-A | 2.1 | 14.7 | 85 | 472 | 9 | MOLECULE: ANTHRAX TOXIN EXPRESSION TRANS-ACTING POSITIVE RE |
| 1990 | 4n78-A | 2.1 | 6.5 | 71 | 1184 | 14 | MOLECULE: CYTOPLASMIC FMR1-INTERACTING PROTEIN 1; |
| 1991 | 2pnk-A | 2.1 | 9.7 | 48 | 424 | 4 | MOLECULE: BH0493 PROTEIN; |
| 1992 | 2p6v-A | 2.1 | 9.4 | 55 | 97 | 7 | MOLECULE: TRANSCRIPTION INITIATION FACTOR TFIID SUBUNIT 4; |

| | | | | | | | |
|---|---|---|---|---|---|---|---|
| 1993 | 5mlp-A | 2.1 | 5.7 | 64 | 255 | 9 | MOLECULE: UNCHARACTERIZED PROTEIN; |
| 1994 | 7crc-B | 2.1 | 13.3 | 73 | 1112 | 10 | MOLECULE: NAD+ HYDROLASE (NADASE); |
| 1995 | 3dao-B | 2.1 | 18.5 | 63 | 274 | 11 | MOLECULE: PUTATIVE PHOSPHATE; |
| 1996 | 2p01-A | 2.1 | 9.9 | 74 | 323 | 7 | MOLECULE: ALPHA-2-MACROGLOBULIN RECEPTOR-ASSOCIATED |
| 1997 | 5woe-A | 2.1 | 3.9 | 64 | 125 | 8 | MOLECULE: SORTING NEXIN-25; |
| 1998 | 6wpm-A | 2.1 | 10.1 | 57 | 401 | 0 | MOLECULE: SUBSTRATE-BINDING PROTEIN; |
| 1999 | 6xp5-G | 2.1 | 4.9 | 43 | 122 | 5 | MOLECULE: MEDIATOR OF RNA POLYMERASE II TRANSCRIPTION SUBUN |
| 2000 | 2yb5-F | 2.1 | 7.1 | 65 | 215 | 9 | MOLECULE: PUTATIVE FUSIDIC ACID RESISTANCE PROTEIN; |
| 2001 | 2rq5-A | 2.1 | 4.8 | 41 | 121 | 7 | MOLECULE: PROTEIN JUMONJI; |
| 2002 | 6whj-B | 2.1 | 8.3 | 58 | 335 | 9 | MOLECULE: RIBOKINASE; |
| 2003 | 1gp8-A | 2.1 | 5.8 | 36 | 40 | 11 | MOLECULE: PROTEIN (SCAFFOLDING PROTEIN); |
| 2004 | 7jr7-A | 2.1 | 5.5 | 87 | 588 | 7 | MOLECULE: ATP-BINDING CASSETTE SUB-FAMILY G MEMBER 5; |
| 2005 | 7oq4-Z | 2.1 | 3.5 | 48 | 118 | 23 | MOLECULE: DNA-DIRECTED RNA POLYMERASE SUBUNIT A'; |
| 2006 | 2jnk-A | 2.1 | 6.7 | 69 | 140 | 12 | MOLECULE: HYALURONONGLUCOSAMINIDASE; |
| 2007 | 5kzb-A | 2.1 | 7.3 | 49 | 104 | 16 | MOLECULE: VIRUS MATRIX PROTEIN; |
| 2008 | 1l6n-A | 2.1 | 5.6 | 76 | 288 | 5 | MOLECULE: GAG POLYPROTEIN; |
| 2009 | 7e4h-B | 2.1 | 7.5 | 100 | 317 | 10 | MOLECULE: SORTING ASSEMBLY MACHINERY 50 KDA SUBUNIT; |
| 2010 | 4j19-A | 2.1 | 2.9 | 46 | 77 | 9 | MOLECULE: HOMEOBOX-CONTAINING PROTEIN 1; |
| 2011 | 4yut-A | 2.1 | 12.3 | 76 | 351 | 14 | MOLECULE: FAMILY 3 ADENYLATE CYCLASE; |
| 2012 | 6ph4-B | 2.1 | 4.6 | 50 | 460 | 10 | MOLECULE: BLUE-LIGHT-ACTIVATED HISTIDINE KINASE; |
| 2013 | 6h0g-B | 2.1 | 8.4 | 70 | 401 | 9 | MOLECULE: DNA DAMAGE-BINDING PROTEIN 1,DNA DAMAGE-BINDING P |
| 2014 | 7ob9-M | 2.1 | 6.9 | 63 | 388 | 13 | MOLECULE: DNA-DIRECTED RNA POLYMERASE I SUBUNIT RPA1; |
| 2015 | 6yx5-B | 2.1 | 5.9 | 91 | 339 | 4 | MOLECULE: RAS-RELATED PROTEIN RAB-8A; |
| 2016 | 7bin-L | 2.1 | 5.1 | 47 | 75 | 11 | MOLECULE: FLAGELLAR BIOSYNTHETIC PROTEIN FLIP; |
| 2017 | 6u4o-A | 2.1 | 10.7 | 116 | 479 | 5 | MOLECULE: FUMARATE HYDRATASE; |
| 2018 | 6rfq-9 | 2.1 | 7.1 | 57 | 86 | 5 | MOLECULE: SUBUNIT NUAM OF NADH:UBIQUINONE OXIDOREDUCTASE (C |
| 2019 | 5mlt-A | 2.1 | 15.8 | 71 | 454 | 13 | MOLECULE: ABC TRANSPORTER, SUBSTRATE-BINDING PROTEIN; |
| 2020 | 6u8t-B | 2.1 | 11.7 | 66 | 221 | 9 | MOLECULE: PFHB2; |
| 2021 | 5zxd-B | 2.1 | 10.9 | 76 | 491 | 5 | MOLECULE: ATP-BINDING CASSETTE SUB-FAMILY F MEMBER 1; |
| 2022 | 2pjw-V | 2.1 | 8 | 53 | 91 | 9 | MOLECULE: UNCHARACTERIZED PROTEIN YHL002W; |
| 2023 | 3l76-A | 2.1 | 20.9 | 91 | 589 | 4 | MOLECULE: ASPARTOKINASE; |
| 2024 | 5ayk-A | 2.1 | 9 | 74 | 744 | 7 | MOLECULE: DNAJ HOMOLOG SUBFAMILY C MEMBER 10; |
| 2025 | 5a31-N | 2.1 | 10.7 | 94 | 703 | 5 | MOLECULE: ANAPHASE-PROMOTING COMPLEX SUBUNIT 1; |
| 2026 | 5fg0-A | 2.1 | 4 | 50 | 407 | 6 | MOLECULE: E3 UBIQUITIN-PROTEIN LIGASE LISTERIN; |
| 2027 | 6l30-A | 2.1 | 3.7 | 46 | 518 | 20 | MOLECULE: PROTEIN ECT2; |

| | | | | | | | |
|---|---|---|---|---|---|---|---|
| 2028 | 6rbf-B | 2.1 | 14.8 | 79 | 364 | 8 | MOLECULE: MUCIN-2; |
| 2029 | 5uft-A | 2.1 | 6.6 | 56 | 144 | 7 | MOLECULE: NITROGEN-FIXING NIFU-LIKE, N-TERMINAL; |
| 2030 | 4jd9-G | 2.1 | 3.8 | 66 | 120 | 5 | MOLECULE: 14.5 KDA SALIVARY PROTEIN; |
| 2031 | 1v74-B | 2.1 | 3.5 | 44 | 87 | 14 | MOLECULE: COLICIN D; |
| 2032 | 7bc4-B | 2.1 | 8.3 | 87 | 2054 | 9 | MOLECULE: FATTY ACID SYNTHASE SUBUNIT ALPHA; |
| 2033 | 6hd8-A | 2.1 | 9.5 | 47 | 461 | 4 | MOLECULE: NANOBODY,MALTOSE/MALTODEXTRIN-BINDING PERIPLASMIC |
| 2034 | 7ml7-A | 2.1 | 16.2 | 95 | 1216 | 7 | MOLECULE: TOXIN B; |
| 2035 | 6egc-A | 2.1 | 5.6 | 61 | 146 | 10 | MOLECULE: SC_2L4HC2_23; |
| 2036 | 4yxw-G | 2.1 | 5.8 | 52 | 213 | 10 | MOLECULE: ATP SYNTHASE SUBUNIT ALPHA, MITOCHONDRIAL; |
| 2037 | 6vw7-D | 2.1 | 9.6 | 63 | 514 | 6 | MOLECULE: NAD-DEPENDENT FORMATE DEHYDROGENASE GAMMA |
| 2038 | 4gr6-B | 2.1 | 13.6 | 65 | 105 | 3 | MOLECULE: ATRBCX2; |
| 2039 | 5wlq-A | 2.1 | 7.1 | 53 | 104 | 4 | MOLECULE: CAPSID ASSEMBLY SCAFFOLDING PROTEIN,MYOSIN-7,MICR |
| 2040 | 6nbu-A | 2.1 | 4 | 59 | 126 | 15 | MOLECULE: CRISPR-ASSOCIATED PROTEIN; |
| 2041 | 2uxw-A | 2.1 | 12.8 | 111 | 567 | 4 | MOLECULE: VERY-LONG-CHAIN SPECIFIC ACYL-COA DEHYDROGENASE; |
| 2042 | 6fea-C | 2.1 | 4.5 | 54 | 113 | 15 | MOLECULE: NITROGENASE PROTEIN ALPHA CHAIN; |
| 2043 | 2h8r-A | 2.1 | 5.8 | 81 | 176 | 7 | MOLECULE: HEPATOCYTE NUCLEAR FACTOR 1-BETA; |
| 2044 | 1vi7-A | 2.1 | 11.1 | 68 | 206 | 3 | MOLECULE: HYPOTHETICAL PROTEIN YIGZ; |
| 2045 | 3gn4-A | 2.1 | 8.1 | 61 | 129 | 10 | MOLECULE: MYOSIN-VI; |
| 2046 | 1fse-B | 2.1 | 2.5 | 44 | 70 | 5 | MOLECULE: GERE; |
| 2047 | 3j25-A | 2.1 | 15.4 | 100 | 638 | 7 | MOLECULE: TETRACYCLINE RESISTANCE PROTEIN TETM; |
| 2048 | 2jes-A | 2.1 | 10.3 | 95 | 370 | 9 | MOLECULE: PORTAL PROTEIN; |
| 2049 | 3f46-A | 2.1 | 13.1 | 57 | 345 | 5 | MOLECULE: 5,10-METHENYLTETRAHYDROMETHANOPTERIN |
| 2050 | 7cq3-A | 2.1 | 4.9 | 80 | 298 | 3 | MOLECULE: STRUCTURE-SPECIFIC ENDONUCLEASE SUBUNIT SLX1; |
| 2051 | 4bxx-X | 2.1 | 8.4 | 58 | 120 | 12 | MOLECULE: DNA-DIRECTED RNA POLYMERASE II SUBUNIT RPB1; |
| 2052 | 3npe-A | 2.1 | 4.1 | 60 | 522 | 12 | MOLECULE: 9-CIS-EPOXYCAROTENOID DIOXYGENASE 1, CHLOROPLASTI |
| 2053 | 1nkl-A | 2.1 | 2.9 | 46 | 78 | 13 | MOLECULE: NK-LYSIN; |
| 2054 | 3td7-A | 2.1 | 4.8 | 82 | 253 | 9 | MOLECULE: FAD-LINKED SULFHYDRYL OXIDASE R596; |
| 2055 | 2jek-A | 2.1 | 4.2 | 65 | 140 | 2 | MOLECULE: RV1873; |
| 2056 | 4zmk-A | 2.1 | 4.7 | 47 | 71 | 6 | MOLECULE: TELOMERE LENGTH REGULATOR TAZ1; |
| 2057 | 6lk8-A | 2.1 | 10.7 | 85 | 1409 | 4 | MOLECULE: MGC83295 PROTEIN; |
| 2058 | 3lf9-A | 2.1 | 3.9 | 48 | 120 | 8 | MOLECULE: 4E10_D0_1IS1A_001_C (T161); |
| 2059 | 2nrk-A | 2.1 | 3 | 46 | 165 | 2 | MOLECULE: HYPOTHETICAL PROTEIN GRPB; |
| 2060 | 5z68-C | 2.1 | 4.4 | 73 | 373 | 7 | MOLECULE: DNA REPLICATION AND REPAIR PROTEIN RECF; |
| 2061 | 2r11-D | 2.1 | 7.8 | 47 | 288 | 4 | MOLECULE: CARBOXYLESTERASE NP; |
| 2062 | 7c9m-C | 2.1 | 11.7 | 65 | 260 | 5 | MOLECULE: D-HISTIDINE 2-AMINOBUTANOYLTRANSFERASE; |

| | | | | | | | |
|---|---|---|---|---|---|---|---|
| 2063 | 2f66-B | 2.1 | 4 | 44 | 111 | 7 | MOLECULE: SUPPRESSOR PROTEIN STP22 OF TEMPERATURE- |
| 2064 | 1hrk-A | 2.1 | 12.2 | 53 | 359 | 8 | MOLECULE: FERROCHELATASE; |
| 2065 | 6pns-A | 2.1 | 13.1 | 87 | 1291 | 8 | MOLECULE: RNA-DIRECTED RNA POLYMERASE; |
| 2066 | 4av2-A | 2.1 | 5.2 | 52 | 274 | 6 | MOLECULE: TYPE IV PILUS BIOGENESIS AND COMPETENCE PROTEIN P |
| 2067 | 7amy-A | 2.1 | 4.2 | 59 | 353 | 8 | MOLECULE: FLAGELLAR BIOSYNTHESIS PROTEIN FLHA; |
| 2068 | 6o8b-B | 2.1 | 9.7 | 81 | 657 | 4 | MOLECULE: STIMULATOR OF INTERFERON GENES PROTEIN; |
| 2069 | 4ilo-A | 2.1 | 6 | 52 | 236 | 10 | MOLECULE: CT398; |
| 2070 | 6lum-D | 2.1 | 9.3 | 63 | 146 | 8 | MOLECULE: SUCCINATE DEHYDROGENASE SUBUNIT C; |
| 2071 | 3lyq-B | 2.1 | 9.3 | 58 | 186 | 12 | MOLECULE: IPGB2; |
| 2072 | 6oew-B | 2.1 | 12.5 | 68 | 239 | 4 | MOLECULE: CYTIDYLYLTRANSFERASE; |
| 2073 | 6cn0-B | 2.1 | 3.9 | 74 | 272 | 7 | MOLECULE: 16S RRNA (GUANINE(1405)-N(7))-METHYLTRANSFERASE; |
| 2074 | 1u9d-A | 2.1 | 9.8 | 59 | 122 | 15 | MOLECULE: HYPOTHETICAL PROTEIN VC0714; |
| 2075 | 3tix-C | 2.1 | 4.9 | 59 | 147 | 10 | MOLECULE: UBIQUITIN-LIKE PROTEIN SMT3,RNA-INDUCED TRANSCRIP |
| 2076 | 4asv-A | 2.1 | 5.1 | 54 | 79 | 6 | MOLECULE: SMALL GLUTAMINE-RICH TETRATRICOPEPTIDE REPEAT-CON |
| 2077 | 3rkl-A | 2.1 | 6.5 | 52 | 80 | 10 | MOLECULE: STIV-A81; |
| 2078 | 2i0z-A | 2.1 | 12.5 | 82 | 416 | 9 | MOLECULE: NAD(FAD)-UTILIZING DEHYDROGENASES; |
| 2079 | 1pd3-A | 2.1 | 3.8 | 36 | 54 | 6 | MOLECULE: NONSTRUCTURAL PROTEIN NS2; |
| 2080 | 6w6v-E | 2.1 | 5.1 | 61 | 169 | 10 | MOLECULE: RNA COMPONENT OF RNASE MRP NME1; |
| 2081 | 2ksf-A | 2.1 | 4.2 | 50 | 107 | 10 | MOLECULE: SENSOR PROTEIN KDPD; |
| 2082 | 6pe4-Q | 2.1 | 4.1 | 66 | 369 | 12 | MOLECULE: V-TYPE PROTON ATPASE SUBUNIT A, VACUOLAR ISOFORM; |
| 2083 | 2o4c-A | 2.1 | 3.6 | 53 | 380 | 4 | MOLECULE: ERYTHRONATE-4-PHOSPHATE DEHYDROGENASE; |
| 2084 | 3rfy-A | 2.1 | 4 | 61 | 356 | 7 | MOLECULE: PEPTIDYL-PROLYL CIS-TRANS ISOMERASE CYP38, CHLORO |
| 2085 | 5muu-D | 2.1 | 8.3 | 55 | 148 | 4 | MOLECULE: MAJOR INNER PROTEIN P1; |
| 2086 | 5mz2-D | 2.1 | 12.3 | 69 | 482 | 6 | MOLECULE: RUBISCO LARGE SUBUNIT; |
| 2087 | 6hmj-A | 2.1 | 4.4 | 53 | 359 | 6 | MOLECULE: PUTATIVE PAS/PAC SENSOR PROTEIN; |
| 2088 | 2i8e-A | 2.1 | 2.6 | 39 | 88 | 0 | MOLECULE: HYPOTHETICAL PROTEIN; |
| 2089 | 6f7j-A | 2.1 | 12.9 | 47 | 97 | 11 | MOLECULE: SERRATE RNA EFFECTOR MOLECULE HOMOLOG; |
| 2090 | 3af5-A | 2.1 | 5.6 | 58 | 638 | 2 | MOLECULE: PUTATIVE UNCHARACTERIZED PROTEIN PH1404; |
| 2091 | 7eep-M | 2.1 | 3.8 | 46 | 182 | 7 | MOLECULE: PAM1 PORTAL PROTEINS; |
| 2092 | 6h86-A | 2.1 | 8 | 52 | 76 | 2 | MOLECULE: SYNAPTONEMAL COMPLEX CENTRAL ELEMENT PROTEIN 3; |
| 2093 | 7mpy-B | 2.1 | 7.1 | 71 | 430 | 10 | MOLECULE: FOLATE SYNTHESIS BIFUNCTIONAL PROTEIN; |
| 2094 | 6yvd-D | 2.1 | 4.8 | 74 | 414 | 9 | MOLECULE: CONDENSIN COMPLEX SUBUNIT 2; |
| 2095 | 6wge-B | 2.1 | 9.1 | 76 | 490 | 12 | MOLECULE: STRUCTURAL MAINTENANCE OF CHROMOSOMES PROTEIN 1A; |
| 2096 | 2aj0-A | 2.1 | 3.2 | 41 | 71 | 7 | MOLECULE: PROBABLE CADMIUM-TRANSPORTING ATPASE; |
| 2097 | 4v1a-m | 2.1 | 7.4 | 71 | 109 | 7 | MOLECULE: MITORIBOSOMAL PROTEIN ML37, MRPL37; |

| | | | | | | | |
|---|---|---|---|---|---|---|---|
| 2098 | 7fas-A | 2.1 | 8.1 | 68 | 1229 | 6 | MOLECULE: ERYTHROCYTE MEMBRANE PROTEIN 1, PFEMP1; |
| 2099 | 6ici-A | 2.1 | 18.1 | 84 | 585 | 2 | MOLECULE: [F-ACTIN]-MONOOXYGENASE MICAL3; |
| 2100 | 6lo8-E | 2.1 | 4.4 | 50 | 80 | 12 | MOLECULE: MITOCHONDRIAL IMPORT INNER MEMBRANE TRANSLOCASE S |
| 2101 | 5iqc-A | 2.1 | 8 | 80 | 301 | 9 | MOLECULE: BIFUNCTIONAL AAC/APH; |
| 2102 | 1l8q-A | 2.1 | 8.9 | 55 | 321 | 15 | MOLECULE: CHROMOSOMAL REPLICATION INITIATOR PROTEIN DNAA; |
| 2103 | 2my1-A | 2.1 | 15.2 | 54 | 159 | 11 | MOLECULE: PRE-MRNA-SPLICING FACTOR BUD31; |
| 2104 | 5d1w-D | 2.1 | 3 | 48 | 196 | 6 | MOLECULE: RV3249C TRANSCRIPTIONAL REGULATOR; |
| 2105 | 7aqo-J | 2.1 | 7.1 | 55 | 230 | 4 | MOLECULE: THO COMPLEX SUBUNIT 2; |
| 2106 | 1p68-A | 2.1 | 5.4 | 45 | 102 | 4 | MOLECULE: DE NOVO DESIGNED PROTEIN S-824; |
| 2107 | 1u5k-A | 2.1 | 12.1 | 69 | 242 | 9 | MOLECULE: HYPOTHETICAL PROTEIN; |
| 2108 | 6l7o-C | 2.1 | 11.2 | 53 | 121 | 9 | MOLECULE: NAD(P)H-QUINONE OXIDOREDUCTASE SUBUNIT 1; |
| 2109 | 6t5a-A | 2.1 | 4.7 | 43 | 96 | 9 | MOLECULE: TEGUMENT PROTEIN UL51; |
| 2110 | 5k04-A | 2.1 | 4.9 | 59 | 732 | 8 | MOLECULE: UNCHARACTERIZED PROTEIN; |
| 2111 | 2k19-A | 2.1 | 3 | 50 | 98 | 6 | MOLECULE: PUTATIVE PISCICOLIN 126 IMMUNITY PROTEIN; |
| 2112 | 2ouw-B | 2.1 | 14 | 52 | 136 | 6 | MOLECULE: ALKYLHYDROPEROXIDASE AHPD CORE; |
| 2113 | 2euf-A | 2.1 | 4.1 | 64 | 244 | 9 | MOLECULE: VIRAL CYCLIN; |
| 2114 | 3at7-A | 2.1 | 3.9 | 79 | 251 | 9 | MOLECULE: ALGINATE-BINDING FLAGELLIN; |
| 2115 | 5zib-A | 2.1 | 5.5 | 87 | 579 | 6 | MOLECULE: ALPHA-1,6-MANNOSYLGLYCOPROTEIN 6-BETA-N- |
| 2116 | 7ogp-A | 2.1 | 8.6 | 71 | 361 | 7 | MOLECULE: PHIKZ055,PHIKZ056.1; |
| 2117 | 1le6-B | 2.1 | 4.2 | 57 | 123 | 9 | MOLECULE: GROUP X SECRETORY PHOSPHOLIPASE A2; |
| 2118 | 4hkr-A | 2.1 | 6 | 56 | 165 | 9 | MOLECULE: CALCIUM RELEASE-ACTIVATED CALCIUM CHANNEL PROTEIN |
| 2119 | 6nhh-A | 2.1 | 12.2 | 77 | 431 | 6 | MOLECULE: CYTOCHROME B; |
| 2120 | 6f70-A | 2.1 | 8.5 | 84 | 242 | 11 | MOLECULE: GLUTATHIONE TRANSFERASE; |
| 2121 | 6k93-A | 2.1 | 7.3 | 69 | 239 | 7 | MOLECULE: TYPE III EFFECTOR XOPAI; |
| 2122 | 2hsz-A | 2.1 | 11.1 | 60 | 225 | 10 | MOLECULE: NOVEL PREDICTED PHOSPHATASE; |
| 2123 | 6hu9-c | 2.1 | 4.8 | 80 | 269 | 8 | MOLECULE: CYTOCHROME B-C1 COMPLEX SUBUNIT 1, MITOCHONDRIAL; |
| 2124 | 5u0p-2 | 2.1 | 7.7 | 40 | 78 | 8 | MOLECULE: MEDIATOR COMPLEX SUBUNIT 14; |
| 2125 | 4oph-A | 2.1 | 6.2 | 49 | 202 | 10 | MOLECULE: NONSTRUCTURAL PROTEIN 1; |
| 2126 | 2c0s-A | 2.1 | 4.5 | 39 | 64 | 5 | MOLECULE: CONSERVED DOMAIN PROTEIN; |
| 2127 | 5n77-A | 2.1 | 8 | 83 | 257 | 5 | MOLECULE: MAGNESIUM TRANSPORT PROTEIN CORA; |
| 2128 | 3c18-A | 2.1 | 11.5 | 78 | 290 | 8 | MOLECULE: NUCLEOTIDYLTRANSFERASE-LIKE PROTEIN; |
| 2129 | 6eys-B | 2.1 | 10.2 | 63 | 478 | 10 | MOLECULE: PVDP; |
| 2130 | 4om3-D | 2.1 | 5.6 | 49 | 122 | 4 | MOLECULE: TRANSDUCIN-LIKE ENHANCER PROTEIN 1; |
| 2131 | 6fxd-A | 2.1 | 9.4 | 55 | 128 | 9 | MOLECULE: MUPZ; |
| 2132 | 1eqf-A | 2.1 | 12.4 | 83 | 267 | 5 | MOLECULE: RNA POLYMERASE II TRANSCRIPTION INITIATION |

| | | | | | | | |
|---|---|---|---|---|---|---|---|
| 2133 | 6lcc-A | 2.1 | 8.8 | 58 | 375 | 5 | MOLECULE: AATPS; |
| 2134 | 6ndu-A | 2.1 | 5 | 58 | 170 | 5 | MOLECULE: PARKIN COREGULATED GENE PROTEIN; |
| 2135 | 2nyf-A | 2.1 | 8.1 | 80 | 511 | 11 | MOLECULE: NOSTOC PUNCTIFORME PHENYLALANINE AMMONIA LYASE; |
| 2136 | 5flv-M | 2.1 | 4.7 | 58 | 229 | 10 | MOLECULE: HOMEOBOX PROTEIN NKX-2.5, T-BOX TRANSCRIPTION FAC |
| 2137 | 4p17-B | 2.1 | 9.2 | 76 | 290 | 9 | MOLECULE: RABGAP/TBC PROTEIN; |
| 2138 | 3j7a-E | 2.1 | 9 | 69 | 185 | 1 | MOLECULE: 18S RIBOSOMAL RNA; |
| 2139 | 4ow5-A | 2.1 | 9.3 | 61 | 307 | 8 | MOLECULE: FUSOLIN; |
| 2140 | 4q28-A | 2.1 | 3.1 | 43 | 110 | 7 | MOLECULE: PERIPLAKIN; |
| 2141 | 1of9-A | 2.1 | 2.7 | 47 | 77 | 11 | MOLECULE: PORE-FORMING PEPTIDE AMEOBAPORE A; |
| 2142 | 3rg2-C | 2.1 | 4.4 | 51 | 613 | 6 | MOLECULE: ENTEROBACTIN SYNTHASE COMPONENT E (ENTE), 2,3-DIH |
| 2143 | 4tr2-B | 2.1 | 2.9 | 43 | 474 | 9 | MOLECULE: SUBTILISIN-LIKE 1 SERINE PROTEASE; |
| 2144 | 5gko-A | 2.1 | 6.5 | 70 | 650 | 1 | MOLECULE: MACROLIDE EXPORT ATP-BINDING/PERMEASE PROTEIN MAC |
| 2145 | 6s18-A | 2.1 | 4.6 | 44 | 143 | 2 | MOLECULE: AROMATIC ACID CHEMORECEPTOR; |
| 2146 | 5imj-A | 2.1 | 4.1 | 67 | 246 | 7 | MOLECULE: CELL DIVISION PROTEIN ZAPD; |
| 2147 | 3qr0-A | 2.1 | 15.9 | 81 | 781 | 14 | MOLECULE: PHOSPHOLIPASE C-BETA (PLC-BETA); |
| 2148 | 1paq-A | 2.1 | 4.1 | 61 | 161 | 15 | MOLECULE: TRANSLATION INITIATION FACTOR EIF-2B EPSILON |
| 2149 | 6j9r-B | 2.1 | 5.2 | 41 | 126 | 12 | MOLECULE: BRAIN TUMOR PROTEIN; |
| 2150 | 2y3a-B | 2.1 | 9.5 | 63 | 236 | 10 | MOLECULE: PHOSPHATIDYLINOSITOL-4,5-BISPHOSPHATE 3-KINASE CA |
| 2151 | 1zke-D | 2.1 | 4.2 | 41 | 83 | 15 | MOLECULE: HYPOTHETICAL PROTEIN HP1531; |
| 2152 | 7obb-B | 2.1 | 11 | 78 | 1130 | 6 | MOLECULE: DNA-DIRECTED RNA POLYMERASE I SUBUNIT RPA1; |
| 2153 | 3bit-A | 2.1 | 15.8 | 77 | 447 | 5 | MOLECULE: FACT COMPLEX SUBUNIT SPT16; |
| 2154 | 1upt-B | 2.1 | 6 | 37 | 58 | 11 | MOLECULE: ADP-RIBOSYLATION FACTOR-LIKE PROTEIN 1; |
| 2155 | 6ej7-A | 2.1 | 12.6 | 72 | 706 | 3 | MOLECULE: XYLOSYLTRANSFERASE 1; |
| 2156 | 3c8g-C | 2.1 | 4 | 64 | 167 | 8 | MOLECULE: PUTATIVE TRANSCRIPTIONAL REGULATOR; |
| 2157 | 5h9c-A | 2.1 | 4.8 | 39 | 78 | 10 | MOLECULE: ENVELOPE GLYCOPROTEIN GP95; |
| 2158 | 6ghc-B | 2.1 | 14.2 | 68 | 286 | 6 | MOLECULE: 5-METHYLCYTOSINE-SPECIFIC RESTRICTION ENZYME A; |
| 2159 | 5yh1-A | 2.1 | 13.5 | 65 | 439 | 11 | MOLECULE: MEMBER OF S1P FAMILY OF RIBOSOMAL PROTEINS; |
| 2160 | 4qmf-B | 2.1 | 2.6 | 41 | 175 | 15 | MOLECULE: KRR1 SMALL SUBUNIT PROCESSOME COMPONENT; |
| 2161 | 2kjf-A | 2.1 | 2.6 | 43 | 60 | 7 | MOLECULE: CARNOCYCLIN-A; |
| 2162 | 6hs6-A | 2.1 | 7.7 | 59 | 69 | 2 | MOLECULE: TYPE VI SECRETION PROTEIN IMPA; |
| 2163 | 6tdx-H | 2.1 | 10.9 | 59 | 160 | 10 | MOLECULE: ATP SYNTHASE F1 SUBUNIT GAMMA; |
| 2164 | 2hvr-A | 2.1 | 4.6 | 44 | 319 | 16 | MOLECULE: 5'- |
| 2165 | 5whm-C | 2.1 | 5.1 | 53 | 264 | 17 | MOLECULE: ICLR FAMILY TRANSCRIPTIONAL REGULATOR; |
| 2166 | 5mx5-L | 2.1 | 4.7 | 64 | 214 | 8 | MOLECULE: PROTEASOME ACTIVATOR COMPLEX SUBUNIT 1; |
| 2167 | 2mpk-A | 2.1 | 4.2 | 44 | 74 | 7 | MOLECULE: CHITIN SYNTHASE 1; |

| | | | | | | | |
|---|---|---|---|---|---|---|---|
| 2168 | 6pwd-A | 2.1 | 12.2 | 68 | 176 | 9 | MOLECULE: TYPE III EFFECTOR HOPBF1; |
| 2169 | 6pl5-B | 2.1 | 3.8 | 70 | 577 | 4 | MOLECULE: PEPTIDOGLYCAN GLYCOSYLTRANSFERASE RODA; |
| 2170 | 6pwn-A | 2.1 | 5.9 | 53 | 280 | 6 | MOLECULE: SMALL-CONDUCTANCE MECHANOSENSITIVE CHANNEL; |
| 2171 | 6fml-G | 2.1 | 9.5 | 64 | 731 | 5 | MOLECULE: RUVB-LIKE HELICASE; |
| 2172 | 7bc6-A | 2.1 | 5 | 98 | 432 | 4 | MOLECULE: PROTON-COUPLED FOLATE TRANSPORTER; |
| 2173 | 6hum-L | 2.1 | 5.4 | 40 | 76 | 5 | MOLECULE: NAD(P)H-QUINONE OXIDOREDUCTASE SUBUNIT 1; |
| 2174 | 3bkx-A | 2.1 | 12.6 | 58 | 274 | 5 | MOLECULE: SAM-DEPENDENT METHYLTRANSFERASE; |
| 2175 | 6orb-A | 2.1 | 39 | 78 | 3352 | 5 | MOLECULE: MIDASIN; |
| 2176 | 3c9p-A | 2.1 | 3.7 | 50 | 122 | 12 | MOLECULE: UNCHARACTERIZED PROTEIN SP1917; |
| 2177 | 6m17-A | 2.1 | 6.5 | 81 | 606 | 1 | MOLECULE: SODIUM-DEPENDENT NEUTRAL AMINO ACID TRANSPORTER B |
| 2178 | 5iy7-M | 2.1 | 14.6 | 58 | 310 | 5 | MOLECULE: DNA-DIRECTED RNA POLYMERASE II SUBUNIT RPB1; |
| 2179 | 4kxr-A | 2.1 | 4.4 | 43 | 85 | 7 | MOLECULE: PE25; |
| 2180 | 6neq-g | 2.1 | 9.1 | 100 | 351 | 13 | MOLECULE: 28S RIBOSOMAL RNA, MITOCHONDRIAL; |
| 2181 | 3bh6-B | 2.1 | 12.1 | 66 | 314 | 11 | MOLECULE: ADP-RIBOSYLATION FACTOR-LIKE PROTEIN 3; |
| 2182 | 4gou-A | 2.1 | 6.4 | 75 | 507 | 9 | MOLECULE: EHRGS-RHOGEF; |
| 2183 | 1snl-A | 2.1 | 10.9 | 55 | 99 | 7 | MOLECULE: NUCLEOBINDIN 1; |
| 2184 | 5uif-A | 2.1 | 3.4 | 50 | 112 | 8 | MOLECULE: PS01740; |
| 2185 | 7pmk-D | 2.1 | 10.9 | 57 | 243 | 5 | MOLECULE: DNA REPLICATION LICENSING FACTOR MCM2; |
| 2186 | 6jy0-A | 2.1 | 7.3 | 57 | 410 | 9 | MOLECULE: FLAGELLIN; |
| 2187 | 4rs7-B | 2.1 | 6 | 50 | 88 | 6 | MOLECULE: PARB-C; |
| 2188 | 1f93-A | 2.1 | 3.2 | 49 | 103 | 4 | MOLECULE: DIMERIZATION COFACTOR OF HEPATOCYTE NUCLEAR |
| 2189 | 5b6b-G | 2.1 | 3.8 | 42 | 74 | 17 | MOLECULE: MOB KINASE ACTIVATOR 1B; |
| 2190 | 7lb8-B | 2.1 | 8.7 | 82 | 630 | 7 | MOLECULE: IRON(3+)-HYDROXAMATE IMPORT SYSTEM PERMEASE PROTE |
| 2191 | 7et3-1 | 2.1 | 4.1 | 71 | 285 | 6 | MOLECULE: TRIPLEX CAPSID PROTEIN 2; |
| 2192 | 5wwx-A | 2.1 | 3.4 | 41 | 84 | 10 | MOLECULE: RNA-BINDING E3 UBIQUITIN-PROTEIN LIGASE MEX3C; |
| 2193 | 6z8k-A | 2.1 | 16.4 | 83 | 2158 | 11 | MOLECULE: LA CROSSE VIRUS 5' VRNA 1-10; |
| 2194 | 5x9a-B | 2.1 | 18.6 | 64 | 198 | 6 | MOLECULE: CALAXIN; |
| 2195 | 6x6n-A | 2.1 | 3.5 | 51 | 134 | 8 | MOLECULE: OUTER MEMBRANE VIRULENCE PROTEIN YOPE; |
| 2196 | 2lxe-A | 2.1 | 6.6 | 51 | 109 | 6 | MOLECULE: HISTONE-LYSINE N-METHYLTRANSFERASE SUVR4; |
| 2197 | 2bl8-A | 2.1 | 3.7 | 50 | 93 | 18 | MOLECULE: ENTEROCINE A IMMUNITY PROTEIN; |
| 2198 | 5b7j-A | 2.1 | 5.1 | 65 | 110 | 11 | MOLECULE: SWITCH-ACTIVATING PROTEIN 1; |
| 2199 | 1n93-X | 2.1 | 13.8 | 76 | 335 | 11 | MOLECULE: P40 NUCLEOPROTEIN; |
| 2200 | 6r5k-A | 2.1 | 10.2 | 68 | 1040 | 15 | MOLECULE: PAN2-PAN3 DEADENYLATION COMPLEX CATALYTIC SUBUNIT |
| 2201 | 3urz-B | 2.1 | 4.8 | 46 | 206 | 11 | MOLECULE: UNCHARACTERIZED PROTEIN; |
| 2202 | 4kqt-A | 2.1 | 4.2 | 55 | 169 | 7 | MOLECULE: PUTATIVE OUTER MEMBRANE CHAPERONE (OMPH-LIKE); |

| | | | | | | | |
|---|---|---|---|---|---|---|---|
| 2203 | 7am1-B | 2.1 | 6.7 | 73 | 811 | 15 | MOLECULE: PROTEIN SSD1; |
| 2204 | 6w32-B | 2.1 | 8.7 | 92 | 288 | 11 | MOLECULE: PHOSPHATIDYLINOSITOL TRANSFER PROTEIN SFH5; |
| 2205 | 3l0m-A | 2.1 | 6 | 63 | 328 | 10 | MOLECULE: DRRA; |
| 2206 | 3ueb-E | 2.1 | 3.4 | 47 | 104 | 11 | MOLECULE: PUTATIVE UNCHARACTERIZED PROTEIN; |
| 2207 | 4od8-D | 2.1 | 5.7 | 40 | 51 | 13 | MOLECULE: URACIL-DNA GLYCOSYLASE; |
| 2208 | 4rs0-A | 2.1 | 10.9 | 84 | 557 | 12 | MOLECULE: PROSTAGLANDIN G/H SYNTHASE 2; |
| 2209 | 2vwa-A | 2.1 | 3.8 | 56 | 99 | 16 | MOLECULE: PUTATIVE UNCHARACTERIZED PROTEIN PF13_0012; |
| 2210 | 7d5i-A | 2.1 | 7 | 70 | 443 | 9 | MOLECULE: CYTOCHROME D UBIQUINOL OXIDASE SUBUNIT 1; |
| 2211 | 6c26-3 | 2.1 | 8.6 | 58 | 126 | 10 | MOLECULE: DOLICHYL-DIPHOSPHOOLIGOSACCHARIDE--PROTEIN |
| 2212 | 6fkg-C | 2.1 | 2.9 | 49 | 111 | 6 | MOLECULE: RV1989C (MBCT); |
| 2213 | 4cpg-A | 2.1 | 5.4 | 44 | 69 | 11 | MOLECULE: SMALL GLUTAMINE-RICH TETRATRICOPEPTIDE REPEAT-CON |
| 2214 | 6gyt-B | 2.1 | 3.5 | 61 | 168 | 5 | MOLECULE: HISTONE ACETYLTRANSFERASE P300; |
| 2215 | 6qq4-A | 2.1 | 3.9 | 50 | 121 | 0 | MOLECULE: GENERAL ODORANT-BINDING PROTEIN 28A; |
| 2216 | 3ezx-A | 2.1 | 4.3 | 58 | 212 | 3 | MOLECULE: MONOMETHYLAMINE CORRINOID PROTEIN 1; |
| 2217 | 3pm2-A | 2.1 | 3.6 | 58 | 173 | 7 | MOLECULE: ODORANT BINDING PROTEIN (AGAP007287-PA); |
| 2218 | 4aqi-A | 2.1 | 3.3 | 52 | 96 | 6 | MOLECULE: PROTEIN S100-A7A; |
| 2219 | 1ypy-A | 2.1 | 3.2 | 64 | 182 | 6 | MOLECULE: VIRION MEMBRANE PROTEIN; |
| 2220 | 6agb-K | 2.1 | 5.4 | 49 | 128 | 8 | MOLECULE: CHROMOSOME V, COMPLETE SEQUENCE; |
| 2221 | 4j8f-A | 2.1 | 6.5 | 75 | 551 | 8 | MOLECULE: HEAT SHOCK 70 KDA PROTEIN 1A/1B, HSC70-INTERACTIN |
| 2222 | 4hh5-A | 2.1 | 3.9 | 74 | 163 | 5 | MOLECULE: PUTATIVE TYPE VI SECRETION PROTEIN; |
| 2223 | 5mss-A | 2.1 | 9.1 | 85 | 717 | 7 | MOLECULE: THIOESTER REDUCTASE DOMAIN-CONTAINING PROTEIN; |
| 2224 | 6qld-N | 2.1 | 13.3 | 94 | 371 | 6 | MOLECULE: INNER KINETOCHORE SUBUNIT MIF2; |
| 2225 | 5xu6-A | 2.1 | 9.8 | 65 | 367 | 5 | MOLECULE: INOSITOL-PENTAKISPHOSPHATE 2-KINASE; |
| 2226 | 6y92-A | 2.1 | 9.4 | 56 | 178 | 9 | MOLECULE: B-LYMPHOCYTE ANTIGEN CD20; |
| 2227 | 7pgs-N | 2.1 | 10.5 | 78 | 1787 | 5 | MOLECULE: NEUROFIBROMIN; |
| 2228 | 7bkb-e | 2.1 | 4.4 | 72 | 411 | 6 | MOLECULE: COB--COM HETERODISULFIDE REDUCTASE IRON-SULFUR SU |
| 2229 | 4kqe-A | 2.1 | 7.3 | 65 | 597 | 11 | MOLECULE: GLYCINE--TRNA LIGASE; |
| 2230 | 2lm9-A | 2.1 | 9.2 | 51 | 96 | 12 | MOLECULE: BLO T 21 ALLERGEN; |
| 2231 | 6pd1-C | 2.1 | 5.4 | 60 | 617 | 5 | MOLECULE: NUCLEOTIDYL TRANSFERASE/AMINOTRANSFERASE, CLASS V |
| 2232 | 2q01-B | 2.1 | 5 | 51 | 480 | 12 | MOLECULE: URONATE ISOMERASE; |
| 2233 | 5i6r-A | 2.1 | 13.2 | 82 | 419 | 6 | MOLECULE: SLIT-ROBO RHO GTPASE-ACTIVATING PROTEIN 2; |
| 2234 | 1x4q-A | 2.1 | 4.7 | 50 | 92 | 8 | MOLECULE: U4/U6 SMALL NUCLEAR RIBONUCLEOPROTEIN PRP3; |
| 2235 | 6l4o-A | 2.1 | 9.7 | 69 | 429 | 4 | MOLECULE: APOPTOSIS INHIBITOR 5; |
| 2236 | 1lxj-A | 2.1 | 3.1 | 45 | 104 | 18 | MOLECULE: HYPOTHETICAL 11.5KDA PROTEIN IN HTB2-NTH2 INTERGE |
| 2237 | 2ibo-A | 2.1 | 2.8 | 46 | 90 | 11 | MOLECULE: HYPOTHETICAL PROTEIN SP2199; |

| | | | | | | | |
|---|---|---|---|---|---|---|---|
| 2238 | 1dd3-A | 2.1 | 6.6 | 49 | 128 | 8 | MOLECULE: 50S RIBOSOMAL PROTEIN L7/L12; |
| 2239 | 5t58-D | 2.1 | 7.2 | 50 | 201 | 10 | MOLECULE: KLLA0F02343P; |
| 2240 | 5a2g-A | 2.1 | 15 | 96 | 501 | 5 | MOLECULE: CARBOXYLIC ESTER HYDROLASE; |
| 2241 | 6rd4-X | 2.1 | 10.2 | 89 | 542 | 4 | MOLECULE: ASA-10: POLYTOMELLA F-ATP SYNTHASE ASSOCIATED SUB |
| 2242 | 6ayi-A | 2.1 | 12.2 | 64 | 183 | 6 | MOLECULE: HTH-TYPE TRANSCRIPTIONAL REGULATOR UIDR; |
| 2243 | 1v33-A | 2.1 | 10.8 | 66 | 346 | 8 | MOLECULE: DNA PRIMASE SMALL SUBUNIT; |
| 2244 | 6tm2-A | 2.1 | 4.2 | 79 | 708 | 6 | MOLECULE: MUCIN-2; |
| 2245 | 1i5p-A | 2.1 | 20.9 | 89 | 633 | 8 | MOLECULE: PESTICIDIAL CRYSTAL PROTEIN CRY2AA; |
| 2246 | 6yxu-H | 2.1 | 12.9 | 71 | 691 | 8 | MOLECULE: DNA-DIRECTED RNA POLYMERASE SUBUNIT ALPHA; |
| 2247 | 2d9d-A | 2.1 | 9.2 | 54 | 89 | 4 | MOLECULE: BAG FAMILY MOLECULAR CHAPERONE REGULATOR 5; |
| 2248 | 4acl-A | 2.1 | 5 | 50 | 155 | 14 | MOLECULE: TSSL; |
| 2249 | 6j52-A | 2.1 | 9.6 | 67 | 94 | 12 | MOLECULE: CASPASE RECRUITMENT DOMAIN-ONLY PROTEIN; |
| 2250 | 7emf-K | 2 | 8.4 | 61 | 112 | 5 | MOLECULE: MEDIATOR OF RNA POLYMERASE II TRANSCRIPTION SUBUN |
| 2251 | 1x9b-A | 2 | 8.5 | 42 | 53 | 7 | MOLECULE: HYPOTHETICAL MEMBRANE PROTEIN TA0354_69_121; |
| 2252 | 6kn5-A | 2 | 4.3 | 51 | 225 | 10 | MOLECULE: AF4/FMR2 FAMILY MEMBER 4; |
| 2253 | 6bwi-A | 2 | 13 | 71 | 816 | 11 | MOLECULE: TRANSIENT RECEPTOR POTENTIAL CATION CHANNEL SUBFA |
| 2254 | 6ijf-A | 2 | 3 | 49 | 101 | 6 | MOLECULE: TAI4; |
| 2255 | 3zhe-A | 2 | 17.4 | 70 | 399 | 4 | MOLECULE: NONSENSE-MEDIATED MRNA DECAY PROTEIN; |
| 2256 | 2roe-A | 2 | 2.5 | 39 | 66 | 10 | MOLECULE: HEAVY METAL BINDING PROTEIN; |
| 2257 | 6ras-I | 2 | 16.3 | 59 | 433 | 8 | MOLECULE: DNA (5'- |
| 2258 | 4g6d-B | 2 | 8.7 | 60 | 198 | 5 | MOLECULE: RNA POLYMERASE SIGMA FACTOR RPOD; |
| 2259 | 6s6v-A | 2 | 11.8 | 51 | 386 | 16 | MOLECULE: NUCLEASE SBCCD SUBUNIT D; |
| 2260 | 3jcm-B | 2 | 15.3 | 80 | 428 | 6 | MOLECULE: PRE-MRNA-SPLICING FACTOR 8; |
| 2261 | 5wjt-A | 2 | 6.1 | 77 | 302 | 5 | MOLECULE: FLAGELLIN; |
| 2262 | 5zdh-A | 2 | 12.4 | 88 | 497 | 10 | MOLECULE: TYPE II SECRETION SYSTEM PROTEIN D; |
| 2263 | 6d5f-a | 2 | 8.5 | 56 | 131 | 11 | MOLECULE: FIMBRIAL PROTEIN; |
| 2264 | 4noo-A | 2 | 3.4 | 61 | 200 | 7 | MOLECULE: VGRG PROTEIN; |
| 2265 | 6nf1-A | 2 | 9.1 | 80 | 550 | 4 | MOLECULE: PROTO-ONCOGENE VAV; |
| 2266 | 6wu0-B | 2 | 18.2 | 83 | 854 | 4 | MOLECULE: HOPANOID BIOSYNTHESIS ASSOCIATED RND TRANSPORTER |
| 2267 | 5z7c-A | 2 | 7.6 | 82 | 414 | 9 | MOLECULE: 3'3'-CGAMP-SPECIFIC PHOSPHODIESTERASE 3; |
| 2268 | 2k3o-A | 2 | 3.6 | 64 | 129 | 9 | MOLECULE: TUSP1; |
| 2269 | 5ok8-B | 2 | 5.6 | 50 | 115 | 2 | MOLECULE: LPP20 LIPOPROTEIN; |
| 2270 | 2rd9-A | 2 | 9.8 | 77 | 189 | 6 | MOLECULE: BH0186 PROTEIN; |
| 2271 | 5u9m-D | 2 | 4.3 | 49 | 234 | 4 | MOLECULE: SUPEROXIDE DISMUTASE [CU-ZN]; |
| 2272 | 1win-A | 2 | 2.5 | 50 | 143 | 20 | MOLECULE: FLOTILLIN 2; |

| | | | | | | | |
|---|---|---|---|---|---|---|---|
| 2273 | 4igg-B | 2 | 5.2 | 81 | 771 | 9 | MOLECULE: CATENIN ALPHA-1; |
| 2274 | 5eom-C | 2 | 4 | 72 | 353 | 11 | MOLECULE: PROTEIN MAB-21-LIKE 1; |
| 2275 | 4ycz-A | 2 | 4.8 | 68 | 713 | 4 | MOLECULE: FUSION PROTEIN OF SEC13 AND NUP145C; |
| 2276 | 4kf8-A | 2 | 12.9 | 77 | 344 | 3 | MOLECULE: NUP188; |
| 2277 | 3k1z-A | 2 | 20.4 | 78 | 241 | 8 | MOLECULE: HALOACID DEHALOGENASE-LIKE HYDROLASE DOMAIN-CONTA |
| 2278 | 2fu2-A | 2 | 3.1 | 45 | 78 | 18 | MOLECULE: HYPOTHETICAL PROTEIN SPY2152; |
| 2279 | 5v8z-C | 2 | 5.6 | 50 | 98 | 10 | MOLECULE: ENDOPLASMIC RETICULUM RESIDENT PROTEIN 29; |
| 2280 | 5da9-C | 2 | 6.2 | 50 | 85 | 6 | MOLECULE: PUTATIVE UNCHARACTERIZED PROTEIN,PUTATIVE UNCHARA |
| 2281 | 7pmk-X | 2 | 13.2 | 83 | 665 | 8 | MOLECULE: DNA REPLICATION LICENSING FACTOR MCM2; |
| 2282 | 1k99-A | 2 | 13.5 | 45 | 91 | 9 | MOLECULE: UPSTREAM BINDING FACTOR 1; |
| 2283 | 6x0l-P | 2 | 19.6 | 84 | 474 | 11 | MOLECULE: HISTONE PARYLATION FACTOR 1; |
| 2284 | 6lul-A | 2 | 5.4 | 52 | 223 | 2 | MOLECULE: RESPIRATORY SUPERCOMPLEX FACTOR 2, MITOCHONDRIAL; |
| 2285 | 6w17-G | 2 | 3.6 | 72 | 150 | 8 | MOLECULE: ACTIN-RELATED PROTEIN 3; |
| 2286 | 7o3x-B | 2 | 9.9 | 66 | 215 | 12 | MOLECULE: PROTEIN SLL0617; |
| 2287 | 1u8s-B | 2 | 11.6 | 58 | 172 | 5 | MOLECULE: GLYCINE CLEAVAGE SYSTEM TRANSCRIPTIONAL |
| 2288 | 3cxj-A | 2 | 9.6 | 67 | 144 | 7 | MOLECULE: UNCHARACTERIZED PROTEIN; |
| 2289 | 6xfl-A | 2 | 9.2 | 52 | 79 | 10 | MOLECULE: TYPE 3 SECRETION SYSTEM PILOTIN; |
| 2290 | 7ar9-X | 2 | 3.2 | 45 | 99 | 4 | MOLECULE: ND3; |
| 2291 | 5x4j-A | 2 | 2.2 | 40 | 471 | 8 | MOLECULE: UNCHARACTERIZED PROTEIN; |
| 2292 | 1uus-A | 2 | 8.7 | 99 | 465 | 6 | MOLECULE: STAT PROTEIN; |
| 2293 | 3zta-A | 2 | 4.4 | 58 | 139 | 12 | MOLECULE: ANTI-SIGMA-FACTOR ANTAGONIST (STAS) DOMAIN PROTEI |
| 2294 | 6nt7-A | 2 | 16.4 | 54 | 294 | 0 | MOLECULE: STIMULATOR OF INTERFERON GENES PROTEIN; |
| 2295 | 5c00-A | 2 | 5.1 | 55 | 202 | 4 | MOLECULE: MDBA PROTEIN; |
| 2296 | 2l22-A | 2 | 10.4 | 63 | 183 | 8 | MOLECULE: MUPIROCIN DIDOMAIN ACYL CARRIER PROTEIN; |
| 2297 | 2ets-A | 2 | 4.9 | 50 | 115 | 8 | MOLECULE: HYPOTHETICAL PROTEIN; |
| 2298 | 2q0o-D | 2 | 6.1 | 48 | 87 | 8 | MOLECULE: PROBABLE TRANSCRIPTIONAL ACTIVATOR PROTEIN TRAR; |
| 2299 | 4p6v-B | 2 | 6.4 | 66 | 348 | 6 | MOLECULE: NA(+)-TRANSLOCATING NADH-QUINONE REDUCTASE SUBUNI |
| 2300 | 6giy-G | 2 | 7.1 | 67 | 444 | 6 | MOLECULE: TSSF; |
| 2301 | 1mqs-A | 2 | 11.3 | 86 | 588 | 13 | MOLECULE: SLY1 PROTEIN; |
| 2302 | 3cit-A | 2 | 4.8 | 50 | 155 | 12 | MOLECULE: SENSOR HISTIDINE KINASE; |
| 2303 | 2wop-A | 2 | 12.7 | 71 | 555 | 7 | MOLECULE: CLAVULANIC ACID BIOSYNTHESIS OLIGOPEPTIDE |
| 2304 | 5gai-K | 2 | 3.5 | 51 | 146 | 14 | MOLECULE: PORTAL PROTEIN; |
| 2305 | 1urf-A | 2 | 6.1 | 50 | 81 | 12 | MOLECULE: PROTEIN KINASE C-LIKE 1; |
| 2306 | 4ng2-E | 2 | 5.9 | 50 | 94 | 4 | MOLECULE: TRANSCRIPTIONAL ACTIVATOR PROTEIN LASR; |
| 2307 | 5ehk-A | 2 | 12.5 | 68 | 1036 | 10 | MOLECULE: LANTIBIOTIC DEHYDRATASE; |

| | | | | | | | |
|---|---|---|---|---|---|---|---|
| 2308 | 6fqb-A | 2 | 6.2 | 73 | 383 | 4 | MOLECULE: MUR LIGASE FAMILY PROTEIN; |
| 2309 | 6q0x-A | 2 | 9.7 | 65 | 373 | 11 | MOLECULE: SORTING NEXIN MVP1; |
| 2310 | 3veb-B | 2 | 9.9 | 59 | 150 | 5 | MOLECULE: MACRODOMAIN TER PROTEIN; |
| 2311 | 5aa5-C | 2 | 9.9 | 69 | 561 | 7 | MOLECULE: NIFE-HYDROGENASE SMALL SUBUNIT, HOFK; |
| 2312 | 6f5d-G | 2 | 4.8 | 56 | 277 | 7 | MOLECULE: ATP SYNTHASE ALPHA CHAIN, MITOCHONDRIAL; |
| 2313 | 3ce9-A | 2 | 6.9 | 100 | 349 | 6 | MOLECULE: GLYCEROL DEHYDROGENASE; |
| 2314 | 5e8j-A | 2 | 6.2 | 56 | 311 | 13 | MOLECULE: MRNA CAP GUANINE-N7 METHYLTRANSFERASE; |
| 2315 | 2jmh-A | 2 | 5.6 | 56 | 117 | 9 | MOLECULE: MITE ALLERGEN BLO T 5; |
| 2316 | 5yz0-C | 2 | 5.6 | 91 | 362 | 9 | MOLECULE: SERINE/THREONINE-PROTEIN KINASE ATR; |
| 2317 | 6nps-A | 2 | 5.7 | 88 | 968 | 10 | MOLECULE: AXYAGU115A; |
| 2318 | 7clv-A | 2 | 8.2 | 76 | 222 | 4 | MOLECULE: TIM23 ISOFORM 1; |
| 2319 | 6gdj-A | 2 | 4.1 | 38 | 71 | 5 | MOLECULE: MTO2; |
| 2320 | 6rj9-C | 2 | 18.9 | 74 | 836 | 3 | MOLECULE: ACRIIA6; |
| 2321 | 5v2o-A | 2 | 3.8 | 44 | 60 | 9 | MOLECULE: TP2; |
| 2322 | 2yn7-A | 2 | 6 | 77 | 214 | 17 | MOLECULE: OUTER SURFACE PROTEIN; |
| 2323 | 6oit-F | 2 | 4.2 | 40 | 342 | 13 | MOLECULE: PROTEIN RDM1; |
| 2324 | 2rp4-A | 2 | 5.7 | 44 | 71 | 11 | MOLECULE: TRANSCRIPTION FACTOR P53; |
| 2325 | 4rbr-A | 2 | 2.5 | 43 | 131 | 14 | MOLECULE: HTH-TYPE TRANSCRIPTIONAL REGULATOR ROT; |
| 2326 | 6lw5-A | 2 | 5.1 | 75 | 427 | 12 | MOLECULE: SOLUBLE CYTOCHROME B562,N-FORMYL PEPTIDE RECEPTOR |
| 2327 | 2pzi-A | 2 | 10.4 | 91 | 654 | 8 | MOLECULE: PROBABLE SERINE/THREONINE-PROTEIN KINASE PKNG; |
| 2328 | 5xtc-j | 2 | 11.6 | 56 | 115 | 5 | MOLECULE: NADH DEHYDROGENASE [UBIQUINONE] IRON-SULFUR PROTE |
| 2329 | 3av0-B | 2 | 9.8 | 63 | 365 | 6 | MOLECULE: DNA DOUBLE-STRAND BREAK REPAIR PROTEIN MRE11; |
| 2330 | 5jjh-A | 2 | 11.9 | 87 | 704 | 6 | MOLECULE: 4-ALPHA-GLUCANOTRANSFERASE; |
| 2331 | 4zcf-A | 2 | 7.7 | 72 | 616 | 8 | MOLECULE: RESTRICTION ENDONUCLEASE ECOP15I, MODIFICATION SU |
| 2332 | 5nik-D | 2 | 4.1 | 48 | 340 | 19 | MOLECULE: OUTER MEMBRANE PROTEIN TOLC; |
| 2333 | 5a7d-R | 2 | 10.5 | 81 | 311 | 14 | MOLECULE: PINS; |
| 2334 | 1te2-A | 2 | 8.1 | 73 | 218 | 4 | MOLECULE: 2-DEOXYGLUCOSE-6-P PHOSPHATASE; |
| 2335 | 4cej-B | 2 | 16.2 | 97 | 1156 | 6 | MOLECULE: ATP-DEPENDENT HELICASE/NUCLEASE SUBUNIT A; |
| 2336 | 5j1s-A | 2 | 9.9 | 68 | 277 | 12 | MOLECULE: TORSIN-1A; |
| 2337 | 6g2a-A | 2 | 10.8 | 73 | 361 | 4 | MOLECULE: [PROTEIN ADP-RIBOSYLARGININE] HYDROLASE; |
| 2338 | 6f1y-j | 2 | 10.5 | 53 | 303 | 6 | MOLECULE: CYTOPLASMIC DYNEIN 1 HEAVY CHAIN 1,DYNEIN HEAVY C |
| 2339 | 3din-E | 2 | 5.6 | 43 | 65 | 9 | MOLECULE: PROTEIN TRANSLOCASE SUBUNIT SECA; |
| 2340 | 1in0-A | 2 | 4.8 | 67 | 162 | 7 | MOLECULE: YAJQ PROTEIN; |
| 2341 | 6x0u-D | 2 | 6.1 | 60 | 158 | 15 | MOLECULE: GAMMA-TUBULIN COMPLEX COMPONENT 3; |
| 2342 | 2oeb-A | 2 | 5.1 | 57 | 152 | 12 | MOLECULE: HYPOTHETICAL PROTEIN; |

| | | | | | | | |
|---|---|---|---|---|---|---|---|
| 2343 | 6iqs-C | 2 | 7.3 | 64 | 541 | 5 | MOLECULE: LIPOPROTEIN NLPI; |
| 2344 | 7pmk-Y | 2 | 6 | 57 | 98 | 16 | MOLECULE: DNA REPLICATION LICENSING FACTOR MCM2; |
| 2345 | 3qnq-D | 2 | 10.5 | 75 | 436 | 5 | MOLECULE: PTS SYSTEM, CELLOBIOSE-SPECIFIC IIC COMPONENT; |
| 2346 | 2m5o-A | 2 | 15.1 | 56 | 97 | 7 | MOLECULE: NFU1 IRON-SULFUR CLUSTER SCAFFOLD HOMOLOG, MITOCH |
| 2347 | 6j6g-C | 2 | 17.2 | 95 | 920 | 11 | MOLECULE: PRE-MRNA-SPLICING FACTOR 8; |
| 2348 | 5a1v-Q | 2 | 11.4 | 71 | 292 | 6 | MOLECULE: ADP-RIBOSYLATION FACTOR 1; |
| 2349 | 5ame-A | 2 | 2.9 | 42 | 123 | 2 | MOLECULE: BROMODOMAIN-CONTAINING PROTEIN 1; |
| 2350 | 3bf5-A | 2 | 10.3 | 53 | 284 | 8 | MOLECULE: RIBOKINASE RELATED PROTEIN; |
| 2351 | 2kg7-B | 2 | 6.7 | 53 | 97 | 2 | MOLECULE: UNCHARACTERIZED PROTEIN ESXG (PE FAMILY PROTEIN); |
| 2352 | 5xjt-2 | 2 | 6.1 | 50 | 217 | 12 | MOLECULE: GEM-ASSOCIATED PROTEIN 2; |
| 2353 | 6u0o-B | 2 | 7.4 | 66 | 288 | 8 | MOLECULE: LYSOSTAPHIN RESISTANCE PROTEIN A; |
| 2354 | 5l4k-V | 2 | 9.3 | 67 | 293 | 7 | MOLECULE: 26S PROTEASOME NON-ATPASE REGULATORY SUBUNIT 4; |
| 2355 | 2lyy-A | 2 | 3.3 | 50 | 96 | 6 | MOLECULE: UNCHARACTERIZED PROTEIN; |
| 2356 | 2iaz-B | 2 | 15 | 55 | 113 | 5 | MOLECULE: HYPOTHETICAL PROTEIN SP1372; |
| 2357 | 5a63-D | 2 | 7.9 | 54 | 100 | 4 | MOLECULE: NICASTRIN; |
| 2358 | 2kna-A | 2 | 5.4 | 51 | 104 | 4 | MOLECULE: BACULOVIRAL IAP REPEAT-CONTAINING PROTEIN 4; |
| 2359 | 1khm-A | 2 | 3.4 | 43 | 89 | 12 | MOLECULE: PROTEIN (HNRNP K); |
| 2360 | 5n9q-A | 2 | 14 | 72 | 128 | 4 | MOLECULE: INACTIVE POLY [ADP-RIBOSE] POLYMERASE RCD1; |
| 2361 | 6fpd-A | 2 | 6.3 | 50 | 191 | 10 | MOLECULE: PROTEIN AB21; |
| 2362 | 5ldi-A | 2 | 17.2 | 55 | 175 | 11 | MOLECULE: RNA 2',3'-CYCLIC PHOSPHODIESTERASE; |
| 2363 | 6xrx-A | 2 | 12.5 | 88 | 551 | 7 | MOLECULE: MALTOSE/MALTODEXTRIN-BINDING PERIPLASMIC PROTEIN, |
| 2364 | 5f1q-A | 2 | 12.8 | 71 | 515 | 7 | MOLECULE: PERIPLASMIC DIPEPTIDE TRANSPORT PROTEIN; |
| 2365 | 6v85-A | 2 | 16.1 | 76 | 1901 | 3 | MOLECULE: RNA-DIRECTED RNA POLYMERASE L; |
| 2366 | 1ss3-A | 2 | 13.9 | 45 | 50 | 9 | MOLECULE: POLLEN ALLERGEN OLE E 6; |
| 2367 | 2pmy-A | 2 | 3.1 | 39 | 73 | 10 | MOLECULE: RAS AND EF-HAND DOMAIN-CONTAINING PROTEIN; |
| 2368 | 7bfq-B | 2 | 6.7 | 60 | 590 | 0 | MOLECULE: INTEGRATOR COMPLEX SUBUNIT 9; |
| 2369 | 4bi8-A | 2 | 4.3 | 49 | 94 | 8 | MOLECULE: RAP1A; |
| 2370 | 1r9q-A | 2 | 3.1 | 49 | 310 | 4 | MOLECULE: GLYCINE BETAINE-BINDING PERIPLASMIC PROTEIN; |
| 2371 | 2lu1-A | 2 | 3.3 | 43 | 89 | 14 | MOLECULE: SUBTILASE; |
| 2372 | 2r10-B | 2 | 12.9 | 61 | 298 | 11 | MOLECULE: CHROMATIN STRUCTURE-REMODELING COMPLEX PROTEIN |
| 2373 | 1yvw-A | 2 | 5.8 | 57 | 92 | 4 | MOLECULE: PHOSPHORIBOSYL-ATP PYROPHOSPHATASE; |
| 2374 | 7ktr-A | 2 | 10 | 98 | 3042 | 7 | MOLECULE: TRANSFORMATION/TRANSCRIPTION DOMAIN-ASSOCIATED PR |
| 2375 | 6ulg-G | 2 | 12.1 | 59 | 310 | 7 | MOLECULE: FOLLICULIN; |
| 2376 | 3t57-A | 2 | 14.3 | 52 | 288 | 8 | MOLECULE: UDP-N-ACETYLGLUCOSAMINE O-ACYLTRANSFERASE DOMAIN- |
| 2377 | 7b2b-A | 2 | 3.9 | 48 | 104 | 10 | MOLECULE: AMINO ACID ADENYLATION DOMAIN-CONTAINING PROTEIN; |

| | | | | | | | |
|---|---|---|---|---|---|---|---|
| 2378 | 7nvr-3 | 2 | 4.5 | 42 | 214 | 12 | MOLECULE: TFIIH BASAL TRANSCRIPTION FACTOR COMPLEX HELICASE |
| 2379 | 3d6j-A | 2 | 7.3 | 59 | 210 | 7 | MOLECULE: PUTATIVE HALOACID DEHALOGENASE-LIKE HYDROLASE; |
| 2380 | 5uxg-A | 2 | 4.5 | 76 | 212 | 3 | MOLECULE: ALDEHYDE DEFORMYLATING OXYGENASE; |
| 2381 | 6n9j-B | 2 | 11.8 | 50 | 356 | 4 | MOLECULE: CLOSTRIPAIN-RELATED PROTEIN; |
| 2382 | 6swy-9 | 2 | 8.2 | 48 | 157 | 0 | MOLECULE: VACUOLAR IMPORT AND DEGRADATION PROTEIN 28; |
| 2383 | 3nyb-A | 2 | 11.2 | 72 | 323 | 7 | MOLECULE: POLY(A) RNA POLYMERASE PROTEIN 2; |
| 2384 | 6nr8-3 | 2 | 4.2 | 44 | 132 | 11 | MOLECULE: PREFOLDIN SUBUNIT 1; |
| 2385 | 3kkz-A | 2 | 10 | 69 | 257 | 6 | MOLECULE: UNCHARACTERIZED PROTEIN Q5LES9; |
| 2386 | 1gak-A | 2 | 4.7 | 57 | 137 | 11 | MOLECULE: FERTILIZATION PROTEIN; |
| 2387 | 3bqp-B | 2 | 2.5 | 47 | 80 | 11 | MOLECULE: PROACTIVATOR POLYPEPTIDE; |
| 2388 | 1oao-A | 2 | 7 | 68 | 673 | 6 | MOLECULE: CARBON MONOXIDE DEHYDROGENASE/ACETYL-COA |
| 2389 | 1ijl-B | 2 | 4.4 | 59 | 123 | 10 | MOLECULE: PHOSPHOLIPASE A2; |
| 2390 | 5ux5-A | 2 | 12.3 | 60 | 963 | 13 | MOLECULE: BIFUNCTIONAL PROTEIN PROLINE UTILIZATION A (PUTA) |
| 2391 | 1h21-A | 2 | 8.2 | 64 | 240 | 8 | MOLECULE: SPLIT-SORET CYTOCHROME C; |
| 2392 | 3hi0-A | 2 | 12 | 88 | 501 | 9 | MOLECULE: PUTATIVE EXOPOLYPHOSPHATASE; |
| 2393 | 6s3f-A | 2 | 3.5 | 43 | 65 | 14 | MOLECULE: 2S ALBUMIN; |
| 2394 | 6mn5-D | 2 | 13.5 | 65 | 260 | 8 | MOLECULE: AMINOGLYCOSIDE N(3)-ACETYLTRANSFERASE, AAC(3)-IVA |
| 2395 | 1q1v-A | 2 | 5.6 | 49 | 70 | 12 | MOLECULE: DEK PROTEIN; |
| 2396 | 1vzs-A | 2 | 10.6 | 46 | 76 | 11 | MOLECULE: ATP SYNTHASE COUPLING FACTOR 6, MITOCHONDRIAL |
| 2397 | 5d18-A | 2 | 11 | 76 | 202 | 11 | MOLECULE: TETR FAMILY TRANSCRIPTIONAL REGULATOR; |
| 2398 | 7kah-D | 2 | 6.6 | 52 | 493 | 10 | MOLECULE: PROTEIN TRANSPORT PROTEIN SEC61; |
| 2399 | 1e7d-A | 2 | 2.9 | 54 | 157 | 4 | MOLECULE: RECOMBINATION ENDONUCLEASE VII; |
| 2400 | 1wte-A | 2 | 3.6 | 56 | 272 | 4 | MOLECULE: 5'-D(*AP*CP*CP*GP*GP*GP*CP*CP*CP*TP*GP*CP*C)-3'; |
| 2401 | 5but-I | 2 | 7.3 | 68 | 431 | 7 | MOLECULE: KTR SYSTEM POTASSIUM UPTAKE PROTEIN A,KTR SYSTEM |
| 2402 | 3s63-B | 2 | 4.3 | 43 | 90 | 9 | MOLECULE: SAPOSIN-LIKE PROTEIN; |
| 2403 | 1eyy-A | 2 | 7.3 | 70 | 504 | 10 | MOLECULE: ALDEHYDE DEHYDROGENASE; |
| 2404 | 7ctp-A | 2 | 4.4 | 65 | 553 | 11 | MOLECULE: PROTEIN NIBAN 2; |
| 2405 | 5z08-D | 2 | 5.5 | 36 | 44 | 8 | MOLECULE: CENP-I; |
| 2406 | 5g4d-A | 2 | 3.4 | 45 | 79 | 7 | MOLECULE: CRISPR-ASSOCIATED ENDORIBONUCLEASE CAS2; |
| 2407 | 4wbt-A | 2 | 17.5 | 83 | 369 | 4 | MOLECULE: PROBABLE HISTIDINOL-PHOSPHATE AMINOTRANSFERASE; |
| 2408 | 5fhp-C | 2 | 10.8 | 73 | 209 | 7 | MOLECULE: NICR; |
| 2409 | 6gyb-C | 2 | 7.7 | 53 | 205 | 6 | MOLECULE: VIRB7; |
| 2410 | 7e1w-O | 2 | 10.7 | 71 | 223 | 3 | MOLECULE: CYTOCHROME C OXIDASE SUBUNIT 2; |
| 2411 | 1n1c-A | 2 | 8.2 | 68 | 199 | 4 | MOLECULE: TORA SPECIFIC CHAPERONE; |
| 2412 | 7bk0-a | 2 | 14.2 | 45 | 250 | 16 | MOLECULE: FLAGELLAR M-RING PROTEIN; |

| | | | | | | | |
|---|---|---|---|---|---|---|---|
| 2413 | 1gmu-C | 2 | 11.1 | 58 | 140 | 3 | MOLECULE: UREE; |
| 2414 | 3dee-A | 2 | 12.4 | 50 | 200 | 8 | MOLECULE: PUTATIVE REGULATORY PROTEIN; |
| 2415 | 4od4-A | 2 | 5.1 | 70 | 275 | 7 | MOLECULE: 4-HYDROXYBENZOATE OCTAPRENYLTRANSFERASE; |
| 2416 | 7e40-D | 2 | 10 | 95 | 345 | 12 | MOLECULE: PROTEIN PHOSPHATE STARVATION RESPONSE 2; |

**S2 Table.** Worm strains used in this study

| Strain name | Genotype (transgene or mutant alelle details) | Figures | Source | Notes |
|---|---|---|---|---|
| ERT054 | *jyIs8[pals-5p::gfp, myo-2p::mCherry] X* | 4 | Bakowski et al., 2012 [1] | |
| ERT1053 | *mgTi54[rpl-28p::mScarlet::drh-1]; drh-1(jy110) IV* | 5A/B, S5 | Mao et al., 2020 [2] | GR3325 from Mao et al., 2020 was crossed with *drh-1(jy110)* in this |
| ERT1076 | *jyEx302[vha-6p::drh-1-NTD::wrmScarlet::3xFLAG::unc-54 3'UTR, myo-3p::mCherry]; rde-1(ne219) V; jyIs8 X* | 1C-E, S3B | This study | |
| ERT1151 | *mgTi54[rpl-28p::mScarlet::drh-1]; drh-1(jy110) IV; rde-1(ne219) V* | 5C | Mao et al., 2020 [2] | |
| ERT1182 | *jyEx305[vha-6p::drh-1-NTD::wrmScarlet::3xFLAG::unc-54 3'UTR]; rde-1(ne219) V; jyIs8 X* | 1C/D, 2A/B, S1B | This study | |
| ERT1199 | *jyEx305; rde-1(ne219) V* | 1F, 3A/B | This study | |
| ERT1205 | *jyEx305; zip-1(jy14) III; rde-1(ne219) V; jyIs8 X* | 2A/B | This study | |
| ERT1207 | *jyEx304[vha-6p::drh-1-NTD::wrmScarlet::3xFLAG::unc-54 3'UTR]; rde-1(ne219) V; jyIs8 X* | 1C/D | This study | |
| ERT1208 | *jyEx304; rde-1(ne219) V* | 3A-E, S3A, S4 | This study | |
| ERT1209 | *jyEx302; rde-1(ne219) V* | 3A/B, S3C | This study | |
| ERT1215 | *frSi17[mtl-2p::rde-1 3'UTR] II; rde-1(ne300) V; jyIs8 X* | 4 | Watts et al., 2020 [3] | IG1839 from Watts et al., 2020 was crossed with ERT054 in this study |
| ERT1216 | *frSi21[col-62p::rde-1 3'UTR] II; rde-1(ne300) V; jyIs8 X* | 4 | Watts et al., 2020 [3] | IG1846 from Watts et al., 2020 was crossed with ERT054 in this study |
| ERT1257 | *jyEx305; zip-1::gfp(jy132) III; rde-1(ne219) V* | S2 | This study | |
| ERT1262 | *jyEx336[vha-6p::wrmScarlet::3xFLAG 3'UTR]; rde-1(ne219) V; jyIs8 X* | S1A | This study | |
| ERT711 | *rde-1(ne219) V; jyIs8 X* | 1C/D, 2A/B, S1B | This study | |
| ERT781 | *drh-1(jy110)* | S5 | Sowa et al., 2020 [4] | |
| GR3325 | *mgTi54[rpl-28p::mScarlet::drh-1]* | S5 | Mao et al., 2020 [2] | |
| N2 | wild type | S5 | Caenorhabditis Genetics | |
| WM27 | *rde-1(ne219) V* | 1F | Caenorhabditis Genetics | |

**S3 Table.** Constructs used in this study

| Construct name | Description | Source |
|---|---|---|
| pET636 | *vha-6p::pals-22 cDNA no stop::gfp_SBP_3xFLAG::unc-54 3'UTR* in pCFJ151 | [1] |
| pET770 | *vha-6p::drh-1-NTD::wrmScarlet::3xFLAG::unc-54 3'UTR, unc-119(+)* | This study |
| pET786 | *vha-6p::wrmScarlet::3xFLAG::unc-54 3'UTR, unc-119(+)* | This study |
| pET788 | *vha-6p::drh-1::wrmScarlet::3XFLAG::unc-54 3'UTR, unc-119(+)* | This study |
| Tian233 | *rpl-28p::mScarlet::drh-1::unc-119(+), Mos1* | [2] |

**S4 Table.** Primers used in this study

| Primer name | Sequence | Purpose |
|---|---|---|
| LEB001 | TACAAATCAGGATCAGGATCAGC | Cloning for intestinal expression of *drh-1(2CARD)* |
| LEB002 | CCCGGGTTTATGGGTTTTGG | Cloning for intestinal expression of *drh-1(2CARD)* |
| LEB003 | ATGGTCAGCAAGGGAGAGG | Cloning for empty vector control |
| LEB004 | CCCGGGTTTATGGGTTTTGG | Cloning for empty vector control |
| LEB005 | TTCAGGGCGACTAAAACCTACCAAAACCCATAAACCCGGGAGGAAAAAGCAGTGTTCTTC | Cloning for intestinal expression of *drh-1* |
| LEB006 | GAACTCCTTGATAACTGCCTCTCCCTTGCTGACCATTGCTTCTCTGATTAAATTGACTAC | Cloning for intestinal expression of *drh-1* |
| snb-1 F | CCGGATAAGACCATCTTGACG | qPCR |
| snb-1 R | GACGACTTCATCAACCTGAGC | qPCR |
| pals-5 F | CATTGGAAAGCGATATTGGA | qPCR |
| pals-5 R | TCTCCAGGCACCTATCTTGTAG | qPCR |
| F26F2.1 F | TGGAACCAGGTCAGAGACAC | qPCR |
| F26F2.1 R | TTGTGAGAATTTCCGCGATA | qPCR |
| skr-5 F | CGAAGAGCAAGATGTCAAAATTG | qPCR |
| skr-5 R | AGAAGCTTGGATTGATTGGCA | qPCR |
| cul-6 F | CTGGGCTTACTCACAATGCC | qPCR |
| cul-6 R | GCAGAGTTGGCTTGCTGTAA | qPCR |
| eol-1 F | GAAGGAGGTGGCGATGTTTAT | qPCR |
| eol-1 R | CGGCGTCGATTGTCTCTTT | qPCR |
| RNA1 F | ACCTCACAACTGCCATCTACA | qPCR |
| RNA1 R | GACGCTTCCAAGATTGGTATTGGT | qPCR |
